# *Drosophila pseudoobscura* third chromosome inversion arrangements have temperature-dependent and sex-specific effects on life history traits

**DOI:** 10.64898/2026.04.06.716560

**Authors:** Gabriel A. Reyes Castellon, Ghada Aimadeddine, Caleb R. Chiao, Sadhana Guruprasad, Paris E. Halbert, Syed Ali Hassan, Michelle Q. Luong, Kushani S. Mailanperuma Arachchillage, Yazmin Martinez, Maryam Mukhtarov, Gopika Nair, Ethan N. Nguyen, Chidinma L. Onochie, Om Patel, Jennifer T. Than, Yesbol Manat, IISAGE Consortium, Richard P. Meisel

## Abstract

Chromosomal inversions can create polymorphic “supergenes” consisting of many alleles in linkage disequilibrium that affect constellations of traits. The coadaptation hypothesis supposes that many alleles within an inversion haplotype have synergistic epistatic interactions, which are maintained by suppressed recombination in inversion heterozygotes, heterozygote advantage, and context-dependent fitness effects. The fitness benefits of chromosomal inversions frequently arise from effects on life history syndromes, including ecotypes and reproductive strategies. Life history syndromes can also be caused by individual pleiotropic alleles in ways that resemble the effects of chromosomal inversions. It is therefore possible that the fitness effects of chromosomal inversions arise because of pleiotropic alleles and not coadaptation. These two hypotheses can be distinguished because pleiotropy predicts greater genetic correlations of traits than coadaptation. To test the coadaptation and pleiotropy hypotheses, we measured three life history traits (lifespan, development time, and body size) in *Drosophila pseudoobscura* with six different chromosomal inversion genotypes. We assayed males and females raised at multiple temperatures in order to expose phenotype variation. Temperature affected lifespan and development more than any other factor, but we also observed differences in mortality rates and development times that depended on chromosomal inversions and sex. Despite these context-dependent effects on life history traits, we failed to identify genetic correlations that would be evidence for pleiotropic alleles. Our results therefore suggest that the coadaptation hypothesis is better supported.

## Introduction

Chromosomal inversion polymorphisms can play important roles in local adaptation, speciation, and the evolution of specialized traits (Davis *et al*., 2011; Joron *et al*., 2011; Fuller *et al*., 2019; Villoutreix *et al*., 2021; Berdan *et al*., 2023). An important component of all these effects is that recombination is suppressed between inverted and non-inverted chromosomes (Navarro *et al*., 1997; Fuller *et al*., 2020). Suppressed recombination allows the inverted and non-inverted arrangements to each accumulate alleles at multiple loci that have synergistic epistatic interactions (Charlesworth, 1974; Charlesworth & Flatt, 2021; Kapun *et al*., 2023), contributing to “coadapation” across loci within the inverted region (Dobzhansky, 1949). The coadapted alleles within inversions often have context-dependent fitness effects, such that the the frequencies of inversion arrangements differ across populations or form geographic clines (e.g., Dobzhansky & Sturtevant, 1938; Lowry & Willis, 2010; Durmaz *et al*., 2018).

Among the fitness effects of chromosomal inversions, life history traits appear to be particularly important for local adaptation. For example, chromosomal inversions have been associated with life history trade-offs in insects (Durmaz *et al*., 2018; Mérot *et al*., 2020), plant and animal ecotypes (Lowry & Willis, 2010; Meyer *et al*., 2024), and migratory tendencies in fish (Pearse *et al*., 2014, 2019). Each of these inversion polymorphisms is thought to affect life history syndromes via the compound action of multiple coadapted alleles held in linkage disequilibrium within the inverted region (Schaal *et al*., 2022; Lappalainen *et al*., 2024). The fitness effects of these traits can depend on environmental context, resulting in genotype-environment interactions (GEIs) that cause the frequencies of inversions to vary across geographic regions where ecological factors differ (Wellenreuther & Bernatchez, 2018; Berdan *et al*., 2023; Akopyan *et al*., 2026). The importance of GEIs can be illustrated by the *Drosophila melanogaster In(3R)Payne* inversion, where temperature modulates the extent of life history traits that are affected by inversion genotypes (Weeks *et al*., 2002; Rako *et al*., 2006; Kapun *et al*., 2016b; Durmaz *et al*., 2018; Paris *et al*., 2026). The fitness effects of inversion genotypes can additionally differ between males and females, generating sexually antagonistic effects over life history syndromes (Pearse *et al*., 2019; McAllester & Pool, 2025).

Life history syndromes can also emerge independently of chromosomal inversions. One such way is by pleiotropic effects of individual alleles (Stearns, 1992; Hughes & Leips, 2017; Healy *et al*., 2019). Pleiotropic alleles can create genetic correlations across traits, resulting in “pace of life” syndromes that concordantly affect life history timing, physiological processes, morphology, and behavior (Réale *et al*., 2010; Dammhahn *et al*., 2018; Arnqvist & Rowe, 2023). These correlations can generate life history trade-offs if alleles have discordant fitness effects across traits, leading to antagonistic pleiotropy that could be responsible for aging and senescence (Williams, 1957). The optimal pace may additionally differ between the sexes—males are often expected to be under selection for a “live fast, die young” strategy where they reach sexual maturity earlier but have reduced lifespan, which could differ from the female optimum—meaning that pleiotropic alleles that affect life history traits can also have sexually antagonistic effects (Promislow, 2003; Bonduriansky *et al*., 2008). Moreover, the genetic architecture underlying complex traits can vary across environments as a result of GEIs (Mackay *et al*., 2009; Haselhorst *et al*., 2011; Anderson *et al*., 2014). As an example of all of these factors acting together, *D. melanogaster* alleles that affect lifespan have pleiotropic effects that depend on sex and temperature (Huang *et al*., 2020).

There are important similarities and differences between the fitness effects of chromosomal inversions and the genetic architecture of life history traits more broadly. In both cases, a life history syndrome can emerge from a constellation of individual traits, with fitness effects that depend on sex and GEIs. In the case of chromosomal inversions, it is hypothesized that trait values arise via coadpation—the combined effects of multiple alleles at genetically linked loci. In contrast, other life history syndromes may emerge if a single allele at one locus has pleiotropic effects. Distinguishing between linked alleles and pleiotropy is an important challenge in quantitative genetics (Lande, 1984; Chebib & Guillaume, 2021), which can confound our understanding of how chromosomal inversions affect life history traits. For example, alleles of the *D. melanogaster Insulin-like Receptor* (*InR*) gene have pleiotropic effects on life history traits and are distributed along geographic clines (Paaby *et al*., 2010, 2014), consistent with natural variation in life history syndromes that arise from a single, large-effect locus. However, *InR* is located within the *In(3R)P* inversion, and *InR* is one of multiple loci with allelic differentiation between inverted and non-inverted chromosomes (Fabian *et al*., 2012; Kapun *et al*., 2016a). The extent to which the fitness effects of *In(3R)P* can be explained by pleiotropic *InR* alleles as opposed to coadapted alleles across the inverted region is not yet resolved. More generally, differentiating between coadaptation and pleiotropy remains an important problem in the study of the phenotypic and fitness effects of polymorphic chromosomal inversions.

To address that problem, we used *Drosophila pseudoobscura* to investigate how chromosomal inversions affect life history syndromes. Natural populations of *D. pseudoobscura* segregate for a well-documented inversion polymorphism on the third chromosome, with over 30 different inversion arrangements that have ecologically-relevant fitness effects (Dobzhansky & Sturtevant, 1938; Anderson *et al*., 1991; Fuller *et al*., 2019). The chromosomal inversion arrangements form stable clines across longitudinal, latitudinal, and altitudinal gradients; they cycle across seasons; and they can respond to selection in the laboratory (Dobzhansky, 1943, 1944, 1947, 1948; Wright & Dobzhansky, 1946). These patterns are consistent with GEIs that generate fitness effects which depend on ecological factors, including temperature variation (Schaeffer, 2008). Moreover, there is natural variation for lifespan, and selection on life history traits can cause changes in *D. pseudoobscura* arrangement frequencies (Taylor & Condra, 1980; Taylor *et al*., 1981), consistent with the inversions affecting life history syndromes. While there is some indirect evidence for coadaptation within the *D. pseudoobscura* third chromosome inversions (Schaeffer *et al*., 2003, 2024; Fuller *et al*., 2017), the pleiotropy hypothesis has yet to be tested.

We determined how *D. pseudoobscura* third chromosome inversion arrangements affected life history traits in order to test if those effects were consistent with coadaptation or pleiotropy. Trait correlations are expected to be stronger when pleiotropic alleles underlie life history syndromes than if multiple linked loci are responsible for a complex trait (Chebib & Guillaume, 2021). We measured lifespan, development time, and body size in thousands of *D. pseudoobscura* males and females, each of which carried one of six different chromosomal inversion genotypes. We performed the lifespan and development time assays at multiple temperatures to test for GEIs. The lifespan measurements were also used to calculate mortality rates and rates of aging, or the increase in mortality rate over time (Bronikowski & Promislow, 2005; Baudisch, 2011). We tested if the chromosomal inversions affected each trait, and we tested for correlations amongst traits, between sexes, and across temperatures in order to evaluate the coadaptation and pleiotropy hypotheses.

## Methods

### Strains and rearing conditions

We sampled flies from 28 strains of *D. pseudoobscura*, each of which carries one of six third chromosome inversion arrangements: Arrowhead (AR), Chiricahua (CH), Cuernavaca (CU), Pikes Peak (PP), Standard (ST), and Tree Line (TL). The chromosomal arrangements are related to each other through a series of overlapping inversions on the third chromosome, and the network of relationships is shown in **Figure 1A** (Wallace *et al*., 2011). Each strain in our sample was homozygous for a third chromosome inversion arrangement that was independently derived from a single female from a natural population, which was extracted onto a laboratory derived genetic background (Fuller *et al*., 2016, 2017). Most of the 28 strains were sampled in all experiments, but some strains were not sampled in some experiments because they were not available for that experiment (**Supplementary Table S1)**.

**Figure 1.**
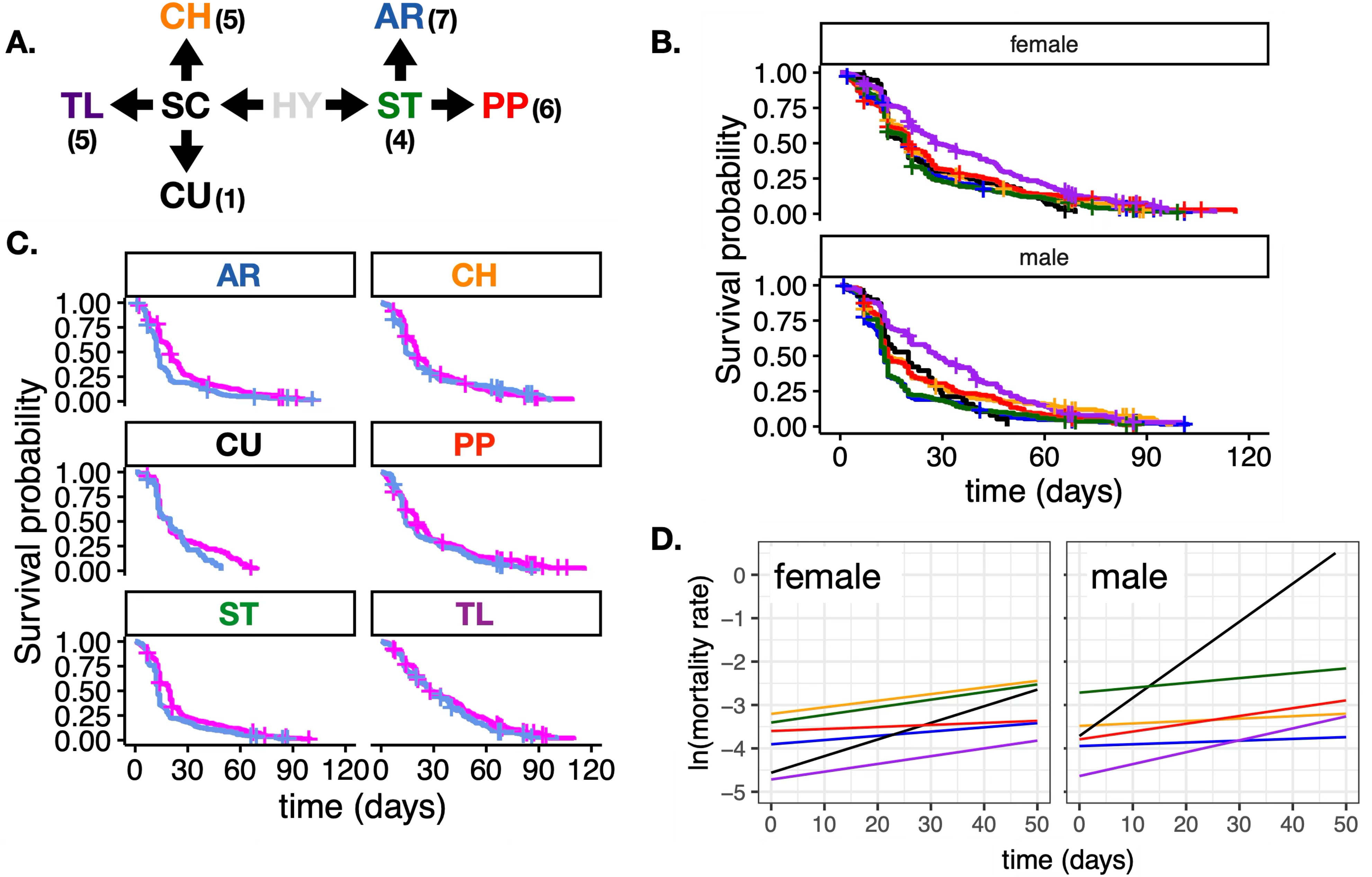
Lifespan and aging of *Drosophila pseudoobscura* carrying one of six inversion arrangements of the third chromosome at 22°C in trial 1. **A.** Network of the evolutionary relationships of the chromosomal inversions, with the common ancestor (Hypothetical, HY) shown. Each of the sampled inversion arrangements is indicated by its two letter code, and the number in parentheses show the amount of independent strains we sampled for a given arrangement. Each arrow represents a single chromosomal inversion. **B.** Survivorship curves for female (top) and male (bottom) flies carrying the six different inversion arrangements (colors match the two letter codes from **A**). **C.** Survivorship curves comparing female (magenta) and male (blue) flies with each of the six inversion arrangements. In **B** and **C**, the X-axis shows the days since emergence from pupa (lifespan), and the Y-axis shows the proportion of **D.** Each line shows the age-specific mortality rate for female (left) or male (right) *D. pseudoobscura* carrying one of six different third chromosome arrangements (color-coded from **A**). Mortality rates were estimated using a Gompertz model separately for each arrangement in each sex. The X-axis shows the days since emergence from pupa, and the Y-axis shows the natural logarithm of the mortality rate, which linearizes the exponential Gompertz function.

Flies from each strain were raised at 18°C, 22°C, or 25°C within reach-in incubators with a 12:12 hour light:dark cycle. Flies were kept in 25×95 mm vials containing standard *Drosophila* media consisting of cornmeal, active dry yeast, sugar, agar, Tegosept, propanoic acid, and water. Fly densities were maintained to prevent larval over-crowding (∼5–20 adult flies per vial), and females were allowed to lay eggs for 3–5 days prior to being transferred to a new vial or discarded. We determined through experimentation that adult female *D. pseudoobscura* collected within 24 h of eclosion did not lay fertilized eggs, and we therefore considered any fly collected within 24 h of emergence to be unmated.

### Lifespan measurements

To assay lifespan, individual flies were collected within 24 h of pupal emergence and stored in vials with standard *Drosophila* medium. Each fly was kept at the same temperature as the vial from which it was collected, and each fly was transferred to a new vial every ∼7 days until it died. The date of emergence from the pupal case (birth) and death were recorded for each fly. These dates were used to calculate the age at death (in days since emergence from the pupal case). Observations were predominantly made on weekdays. If a fly died over an unobserved weekend or holiday, the midpoint of the unobserved period was recorded as the age at death. The one exception to this is if a fly died over the winter holidays, in which case the fly was treated as right censored data on the day before winter holiday (last observation alive). Flies that escaped during transfer to a new vial were also scored as right censored data, with the censoring date recorded as the date of escape. In order to assess the replicability of our assay, two experimental trials were carried out at 22°C. Single trials were performed at 18°C and 25°C.

### Development time assays

We measured egg-to-adult development time for each strain at each temperature. To do so, five female flies were placed into a vial with standard *Drosophila* media, allowed to lay eggs for 24 h, and then removed. Flies were collected daily from the vial once emergence from pupae began, and the sex of each fly was recorded. Egg-to-adult development time was calculated as the difference in days from when the five females (i.e., mothers) were placed in the vial and the date when each fly emerged. Assays were performed at 18°C, 22°C, or 25°C. Note that the development time assays were performed on different flies from the lifespan assays, but the flies were all sampled from the same strains. We could therefore calculate genetic correlations by comparing across strains.

### Body size measurements

We measured the body size of flies from each strain at 22°C. Five females from a given strain were placed into a vial with standard *Drosophila* media and allowed to lay eggs for 24 h, as done with the development time assays. Adult offspring were collected daily starting ∼12–24 h after emergence began. The body length of each fly was measured using an Olympus SZX10 stereomicroscope with the Olympus cellSens software.

### Data analysis

We analyzed the lifespan data using Cox proportional hazards and Gompertz models implemented in the survival and flexsurv packages in the R statistical programming environment (Andersen & Gill, 1982; Therneau & Grambsch, 2010; Jackson, 2016; R Core Team, 2025). The hazard ratio (HR) in a Cox proportional hazards model estimates the relative mortality rate between two groups. The Gompertz model provides an initial mortality rate and a measure of the change in the mortality rate over time, the latter of which is interpreted as the rate of aging (Bronikowski & Promislow, 2005; Baudisch, 2011). To evaluate which factors affected the mortality rate, we compared nested models with inversion arrangement, sex, and temperature as predictors. We extracted the HR and mortality rates for specific levels of factors (e.g., females and males within sex) when the had a significant effect on the response.

We used linear models to analyze the development time and body size data, which were implemented using the nlme and lme4 packages (Bates *et al*., 2015; Pinheiro *et al*., 2023). In our analysis, we compared the fit of nested models in order to identify the predictors (e.g., inversion arrangement, sex, temperature) that significantly affect development time or body size. We also used these models to estimate the effects of specific factors on development time or body size.

Additional analyses were performed to calculate correlations of lifespan and development time. To those ends, we extract the median or mean trait value for each sex within each strain, and we constructed linear models in which one sex-specific trait value was a response and predictors included a sex-specific trait value, chromosomal inversion arrangement, or temperature treatment. In all analyses, multiple testing corrections were applied to control for the familywise error rate when independent statistical tests were performed to assess differences on the same dataset. However, most of our statistical tests were not in the same family of tests, and therefore they did not require any multiple testing corrections. Details of the statistical analyses are provided in the Supplementary Materials.

## Results

### Genotype and sex affect lifespan and aging

We measured lifespan (time from pupal emergence until death) at 22°C in 1,412 female and 887 male *D. pseudoobscura* from third chromosome extraction strains, each of which carried one of six different inversion arrangement genotypes on a common set of genetic backgrounds (**Figure 1B**). We analyzed the lifespan measurements using nested Cox proportional hazards models to assess if sex, inversion genotype, and their interactions affected the mortality rate. A model including both sex and inversion genotype, but not their interaction, fit the data best (Supplementary File S1). However, we observed that many flies died within one day of emergence (likely as a result of handling effects). If we excluded those one-day-old deaths, a sex-inversion interaction term improved model fit (Supplementary File S1).

We used the Cox proportional hazards models to estimate the male:female (m:f) HR, or relative mortality rate. Regardless of which version of the data we analyzed, males had a higher mortality rate than females (m:f HR=1.4, *p*=10^-11^), which resulted in a longer average female lifespan (**Figure 1C**). When we analyzed each inversion genotype separately, the sex difference in mortality rate was greatest in flies carrying the AR arrangement (m:f HR=1.7, *p*=10^-7^), and it was also significant in flies with PP (m:f HR=1.3, *p*=0.02) and ST (m:f HR=1.4, *p*=0.002) arrangements (**Figure 1C**). There was not a significant sex difference in mortality rate for flies with the CH, CU, or TL arrangements. We also analyzed each sex separately, finding that females with the TL arrangement had a reduced mortality rate when compared to females carrying other inversion arrangements (2.3<HR<5.8, *p*<10^-5^), but there was not a significant difference across inversion arrangements in males (**Figure 1B**).

We next tested if there were age-dependent changes in the mortality rate by fitting Gompertz models to our lifespan data. The Gompertz model includes both an initial mortality rate and a shape parameter, the latter of which measures the change in mortality rate with age. The shape parameter can be interpreted as the rate of aging (Baudisch, 2011). We tested if sex or inversion genotype affected the initial mortality rate or shape parameter by comparing the fit of nested Gompertz models, similar to our analysis of the Cox proportional hazards.

In order to test for sex differences in the initial mortality rate or shape parameter, we compared males and females carrying the same inversion arrangement. In this analysis, a model with a sex-specific initial mortality rate fit the data best for flies with either the AR or ST arrangement (Supplementary File S1). This result is consistent with the significant sex differences in mortality rate we observed for AR and ST flies in the Cox proportional hazards models. A sex-specific initial mortality rate did not improve model fit for flies with other inversion genotypes. Likewise, adding a sex-specific shape parameter did not improve model fit (Supplementary File S1), suggesting that there were inversion-dependent sex differences in the initial mortality rate but not in the rate of aging.

We then tested for inversion effects on the initial mortality rate and shape parameter within each sex. To those ends, we compared inversion genotypes within males and females separately. In both males and females, a Gompertz model with inversion-specific initial mortality rates and shape parameters fit the data best (Supplementary File S1). The effect of the shape parameter can be visualized by taking the logarithm of the Gompertz function, where the shape parameter is the slope of the line relating log of the mortality rate to age. We can therefore compare rates of aging by contrasting slopes across inversion genotypes (**Figure 1D**). In each sex, flies carrying the CU arrangement had steeper slope, indicating a faster increase in the mortality rate over time. In other words, the CU arrangement conferred a higher rate of aging. In contrast, flies with an AR arrangement had much smaller increase in mortality rate with age, meaning that the AR arrangement conferred a reduced rate of aging relative to other inversion genotypes.

### Temperature modulates the effects of genotype and sex on lifespan and aging

We next performed the same lifespan assay described above, using males and females from the same strains, but at three different temperatures. The data set included 1,185 flies at 18°C, 1,296 flies at 22°C, and 995 flies at 25°C (Supplementary File S2). The experiments in our first trial (reported above) were performed at 22°C, which allowed us to evaluate the reproducibility of our lifespan measurements by contrasting with the second trial at 22°C. To test if our assay was reproducible, we compared two measures of average lifespan (mean and median) for each strain between the two 22°C trials (**Figure 2A**). In a model that included both sex and inversion arrangement, there was a positive correlation of mean lifespan per strain between the two trials (slope=0.84, *R*^2^=0.62, *p*=10^-7^), and a similar result using the median (Supplementary File S2). We therefore conclude that our lifespan assay is reproducible across strains and sexes within a temperature.

**Figure 2.**
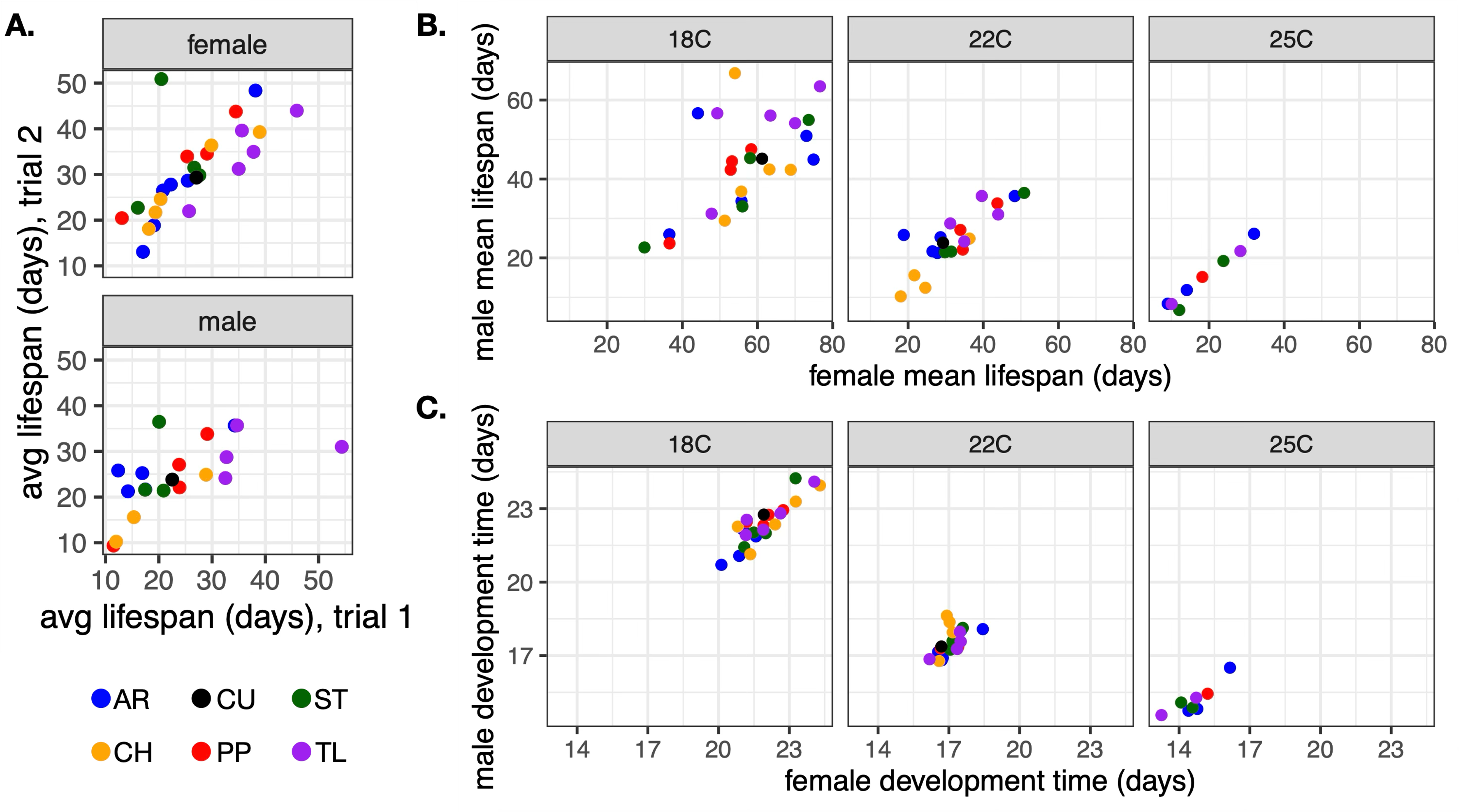
Correlation of lifespan and development time across replicates and sexes. Each dot shows the mean value for a given strain, and the dots are colored based on the third chromosome inversion arrangement carried by the strain. Lifespan was measured as the time from pupal emergence until death. A. Graphs show the correlation of lifespan across strains measured in two experimental trials at 22°C. Each dot shows the mean lifespan of a strain measured in trial 1 (X-axis) and trial 2 (Y-axis). Data are shown for females (top) and males (bottom). B-C. Graphs show the inter-sexual correlations of lifespan and development time. Each dot shows the mean (A) lifespan or (B) egg-to-adult development time of a strain measured in females (X-axis) and males (Y-axis). Assays in B-C were performed at 18°C, 22°C, and 25°C.

We also measured the intersexual correlation (*r*_MF_) of mean lifespan across strains at each temperature. There was a strong positive correlation of lifespan between females and males at 18°C (*r*_MF_=0.55, *p*=0.004), 22°C (*r*_MF_=0.59, *p*=10^-6^), and 25°C (*r*_MF_=0.80, *p*=0.0009) when comparing across strains (**Figure 2B**; Supplementary File S3). We observed a similar positive intersexual correlation when analyzing all three temperatures in a single model (*r*_MF_=0.72, *p*=10^-16^). These results demonstrate an inter-sexual genetic correlation for lifespan, but they do not exclude the effect of sex on lifespan or aging.

We then tested if sex and inversion genotype affected mortality rate at each temperature. A Cox proportional hazard model with sex and inversion, but not their interaction, fit the survival data best for each temperature (Supplementary File S2). Similar to our observations in the first trial at 22°C, males had a higher mortality rate than females at 18°C (m:f HR=1.83, *p*=10^-15^), 22°C (m:f HR=1.63, *p*=10^-15^), and 25°C (m:f HR=1.53, *p*=10^-9^). The effect of this difference in mortality rate is that females had longer average lifespans than males across all three temperatures (**Figure 3A**). However, when considering each inversion arrangement at each temperature, the evidence for sex differences was weaker (Supplementary File S2). For example, including a sex effect on the initial mortality rate significantly improved the fit of a Gompertz model for most inversion arrangements at 18°C, but the effect of sex was not as strong at 22°C or 25°C (Supplementary File S2). Including sex as a shape parameter (i.e., rate of aging) did not significantly improve model fit at any temperature or for any inversion arrangement, similar to what we observed in our initial trial at 22°C. In addition, including inversion genotype as a rate parameter only improved model fit for males at 18°C, but not at other temperatures and not for females. Therefore, while there was evidence for sex and inversion effects on mortality rates at each temperature, the effects were most pronounced at the lowest temperature.

**Figure 3.**
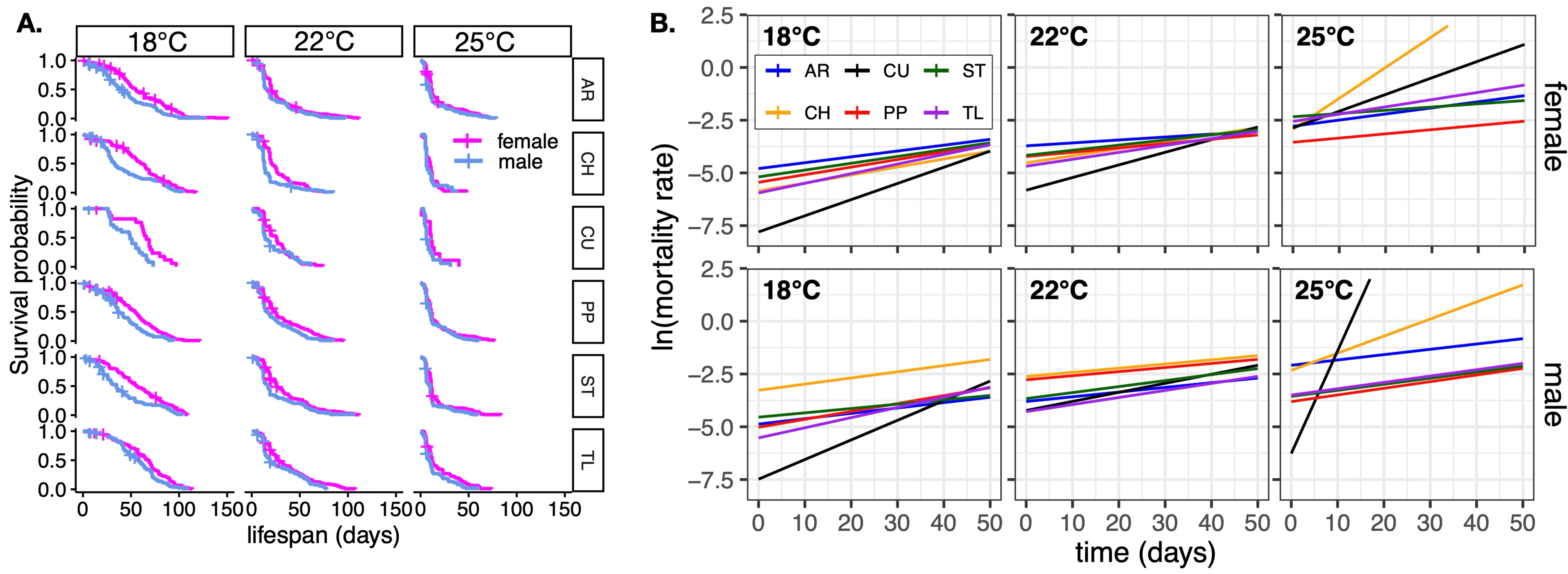
Sex-specific lifespan and aging across genotypes and temperatures. **A.** Survivorship curves in each panel compare female (magenta) and male (blue) flies with one of six inversion arrangements at one of three temperatures. The X-axis shows the days since emergence from pupa (lifespan), and the Y-axis shows the proportion of surviving flies. **B.** Graphs show the age-specific mortality rates for males and females with different third chromosome arrangement genotypes raised across three temperatures. Each line shows the natural logarithm of the age-specific mortality rate for females (top) or males (bottom) carrying one of six different third chromosome arrangements. Flies were raised at 18°C, 22°C, or 25°C. Mortality rates were estimated using a Gompertz model separately for each arrangement in each sex. The X-axis shows the days since emergence from pupa, and the Y-axis shows the natural logarithm of the mortality rate, which linearizes the exponential Gompertz function.

We tested for effects of inversion arrangements within each sex and temperature using TL as a reference because it tended to be the longest lived (**Figure 3A**). We found that PP had a significantly higher mortality rate than TL in males at 18°C (HR=2.2, *p=*0.03), but this difference was no longer significant after applying a multiple testing correction. No other inversion arrangements significantly differed from TL in either sex at any temperature (Supplementary File S2).

To further investigate how temperature modulates the effects of inversion genotype and sex on mortality rate, we analyzed data from all three temperatures together in a single model. The best fitting Cox proportional hazards model included inversion genotype, sex, temperature, and the inversion-temperature interaction as predictors (Supplementary File S2). No other interaction terms improved model fit. The mortality rate increased as temperature increased, in both males and females, causing shorter average lifespans at warmer temperatures (**Figure 3B**). In addition, both males and females carrying the PP arrangement had significantly higher mortality rates than flies carrying the TL arrangement (HR=1.9, *p* < 0.05), when we considered all temperatures in a single model. There was also a significant positive interaction between temperature and whether female flies carried the CH arrangement, meaning increasing temperature increased the mortality rate more when females had the CH arrangement genotype. In contrast, the mortality rate in PP females was less affected by the increase in temperature from 18°C to 22°C.

We determined that temperature affected the rate of aging (i.e., shape parameter, or increase in mortality rate with age) in a Gompertz model for flies carrying some inversion arrangements, and in a sex-specific manner (**Figure 3B**; Supplementary File S2). For example, in AR and ST males and both males and females carrying the CH arrangement, a model including a temperature-specific initial mortality rate fit the data best; a temperature-specific shape parameter did not improve model fit for these inversion-sex combinations. In contrast, a model with both a temperature-specific initial mortality rate and a temperature-specific shape parameter fit the lifespan data best for both males and females carrying either the PP and TL arrangements, as well as females with the AR and ST arrangement genotypes. In all cases where the model included a temperature-specific shape parameter, the effect was to reduce the rate of aging as temperature increased. Therefore, higher temperatures universally increased the initial mortality rate across all inversion genotypes and in both sexes, but higher temperatures also decreased the rate of aging in some genotype-sex combinations (**Figure 3B**).

### Sex, genotype, and temperature affect development time

We next measured the egg-to-adult development time in *D. pseudoobscura* males and females carrying the six different inversions across three temperatures. We measured the development time of 2,799 flies at 18°C, 3,022 flies at 22°C, and 929 flies at 25°C. The data were analyzed using linear models with development time as the response variable (Supplementary File S3). There was a significant positive correlation of average development time between females and males across strains at 18°C (*r*_MF_=0.74, *p*=10^-7^), 22°C (*r*_MF_=0.74, *p*=0.006), 25°C (*r*_MF_=0.78, *p*=0.04), and when considering all temperatures together (*r*_MF_=0.97, *p*=10^-15^). These results demonstrate an inter-sexual correlation for egg-to-adult development time (**Figure 2C**)

We then tested if inversion genotype and sex affected development time at each of the three temperatures. To those ends, we fit linear models to predict egg-to-adult development time at each temperature using sex and inversion genotype as predictors. At each temperature, there was a significant effect of sex, and there was a sex-inversion interaction at 25°C (Supplementary File S3). Males emerged approximately a half day later than females at 18°C (*F*=145, *p*<0.0001), 22°C (*F*=167, *p*<0.0001), and 25°C (*F*=55, *p*<0.0001) when considering all inversion genotypes (**Figure 4A**). At 25°C, where there was a sex-inversion interaction, the sex difference in development time was greatest in flies with the CU arrangement; CU males took >1 day longer than CU females to complete egg-to-adult development (**Figure 4A**). In contrast, the sex difference in development time was much smaller (0.12 days) for flies with the CH arrangement. An alternative explanation of these results is that XX eggs were laid prior to XY eggs within a given 24 h period, which created the appearance of faster female development. We lack the resolution to differentiate faster development from sex differences in egg laying order.

**Figure 4.**
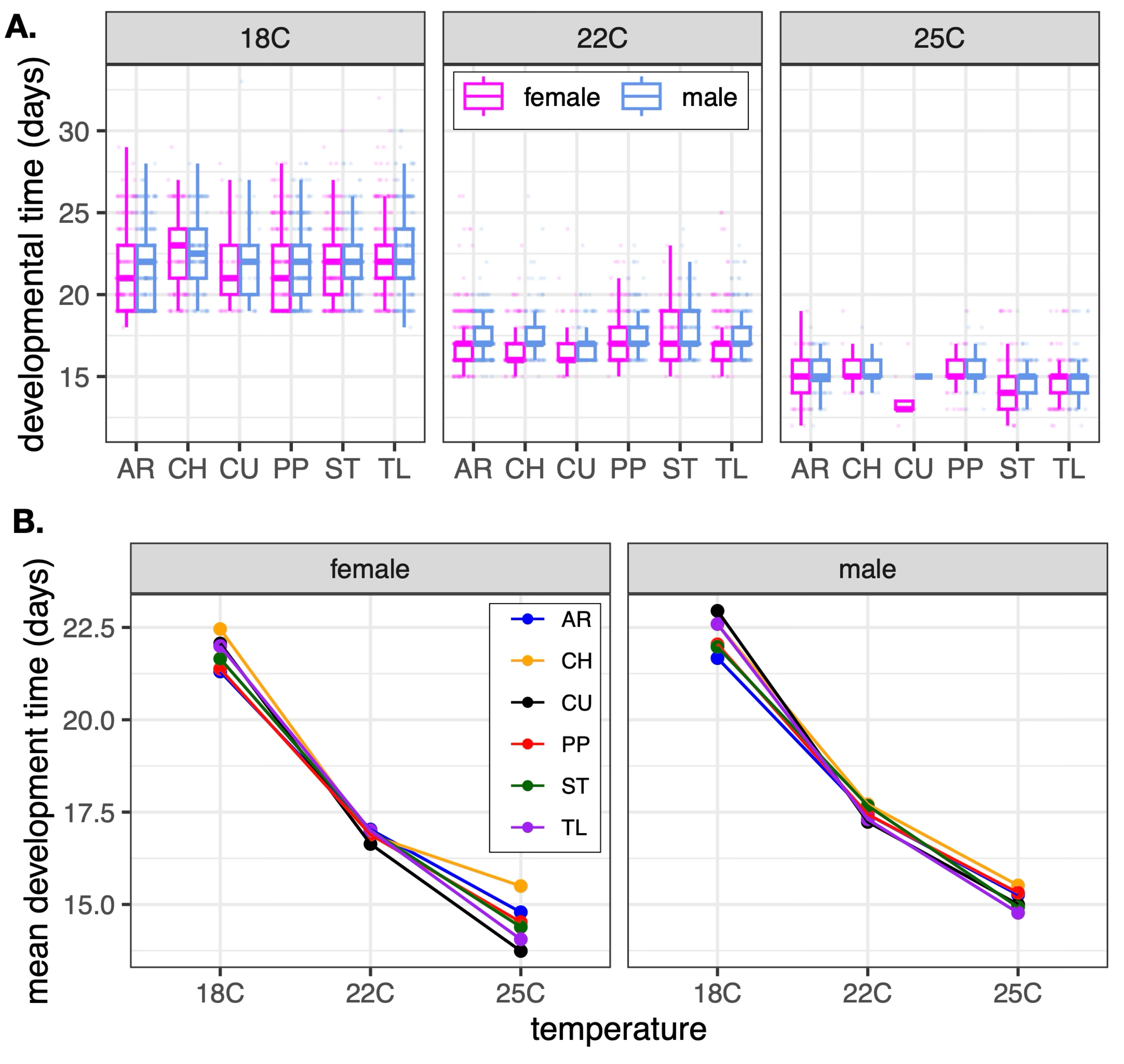
Egg-to-adult development time in females and males carrying different third chromosome inversion arrangements at three temperatures. **A.** Boxplots show the distributions of development times for female (magenta) and male (blue) flies with one of six inversion arrangements at one of three temperatures. Each point in the figure represents the measurement for an individual fly. **B.** Graphs show the mean development time for female (left) and male (right) flies with each of the chromosomal inversions at each of the three temperatures. Each dot connected by a line is colored according to third chromosome arrangement.

We next directly tested for the effect of temperature by analyzing data from all three temperatures in a single model. We found that our data were best explained by a model that included the following predictors of development time: temperature, sex, and an inversion-temperature interaction (Supplementary File S3). Inversion genotype on its own had a mild effect whose significance depended on how model fit was measured. The temperature effect resulted in each 1°C increase in temperature associated with 0.7–1.0 day faster development. In addition, males took approximately a half day longer to complete egg-to-adult development than females, similar to what we saw above when we analyzed each temperature separately. Notably, flies carrying the CU or TL arrangement had the most accelerated development time as temperature increased, while the development of AR flies was least accelerated by the increase in temperature (**Figure 4B**). Our results therefore suggest that the developmental time of flies with the CU arrangement was most sensitive to temperature and sex. In contrast, AR flies developed quickly at colder temperatures but were not as accelerated by increases in temperature. However, these inversion-temperature interactions had much smaller effects than the direct effects of sex and temperature on development time (**Figure 4**).

We tested for a genetic correlation between development time and lifespan at each temperature. Our development time measurements were performed on different flies as those used in the lifespan assays; however, all flies were sampled from the same strains, allowing us to test for genetic correlations by fitting linear models that included the average lifespan of a strain as a response and the following predictors: average development time of each strain, the inversion genotype of the strain, and sex (Supplementary File S3). There was a nearly significant negative correlation between development time and lifespan at 22°C (*β* =-1.9, *F*=3.24, *p* = 0.08), but the correlations were not significant at 18°C or 25°C (**Figure 5**). We next considered all three temperatures together in a single model. Sex, temperature, and development time were significant predictors of lifespan, as was the development time by sex interaction (Supplementary File S3). The interaction between development time and sex arises because development time was negatively correlated with male lifespan, not female lifespan (**Figure 5**); we found that sex significantly affected the relationship between development time and lifespan (*F*=6.35, *p*=0.01). When we accounted for the sex difference in development time, we found that males were expected to live >22 days longer than females based on their egg-to-adult development. We therefore conclude that there was a weak correlation between development time and lifespan, and the sign and significance depended on temperature and sex.

**Figure 5.**
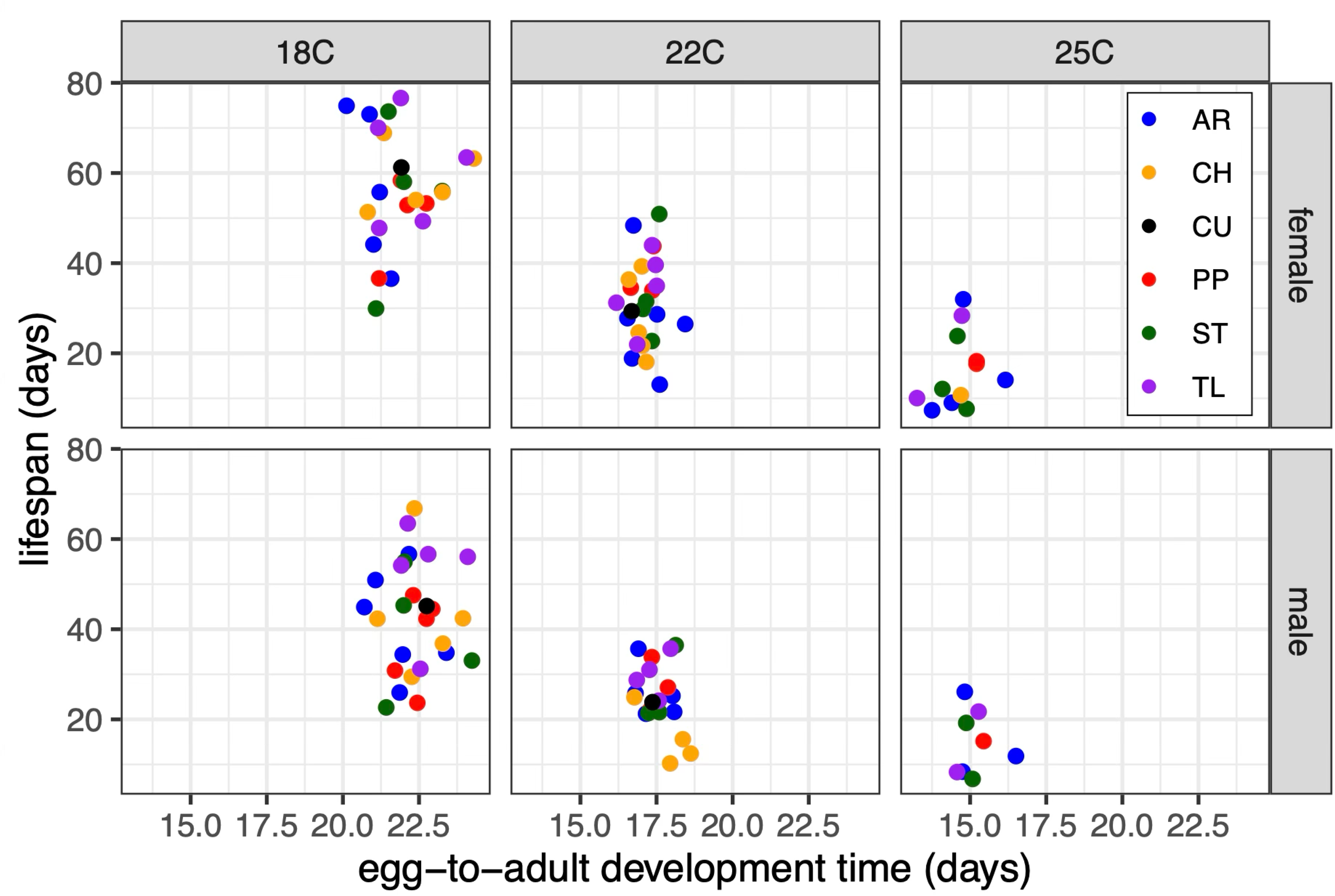
Correlation of lifespan and development time in each sex at three temperatures. Lifespan was measured as the time from pupal emergence until death. Each dot shows the mean lifespan (measured on Y-axis) and egg-to-adult development time (measured on X-axis) of a strain measured in females (top) or males (bottom) at each temperature (18°C, 22°C, or 25°C.).

### Sex differences in body size but no genotype effect

We lastly tested for sex and genotype effects on body size across 1,380 females and 1,235 males from the *D. pseudoobscura* strains at 22°C. There was a significant effect of sex on body length (*F*=851.6, *p*<0.0001), but no effect of inversion genotype (**Figure 6**; Supplementary File S4). We estimated that males were 282–306 µm smaller than females.

**Figure 6.**
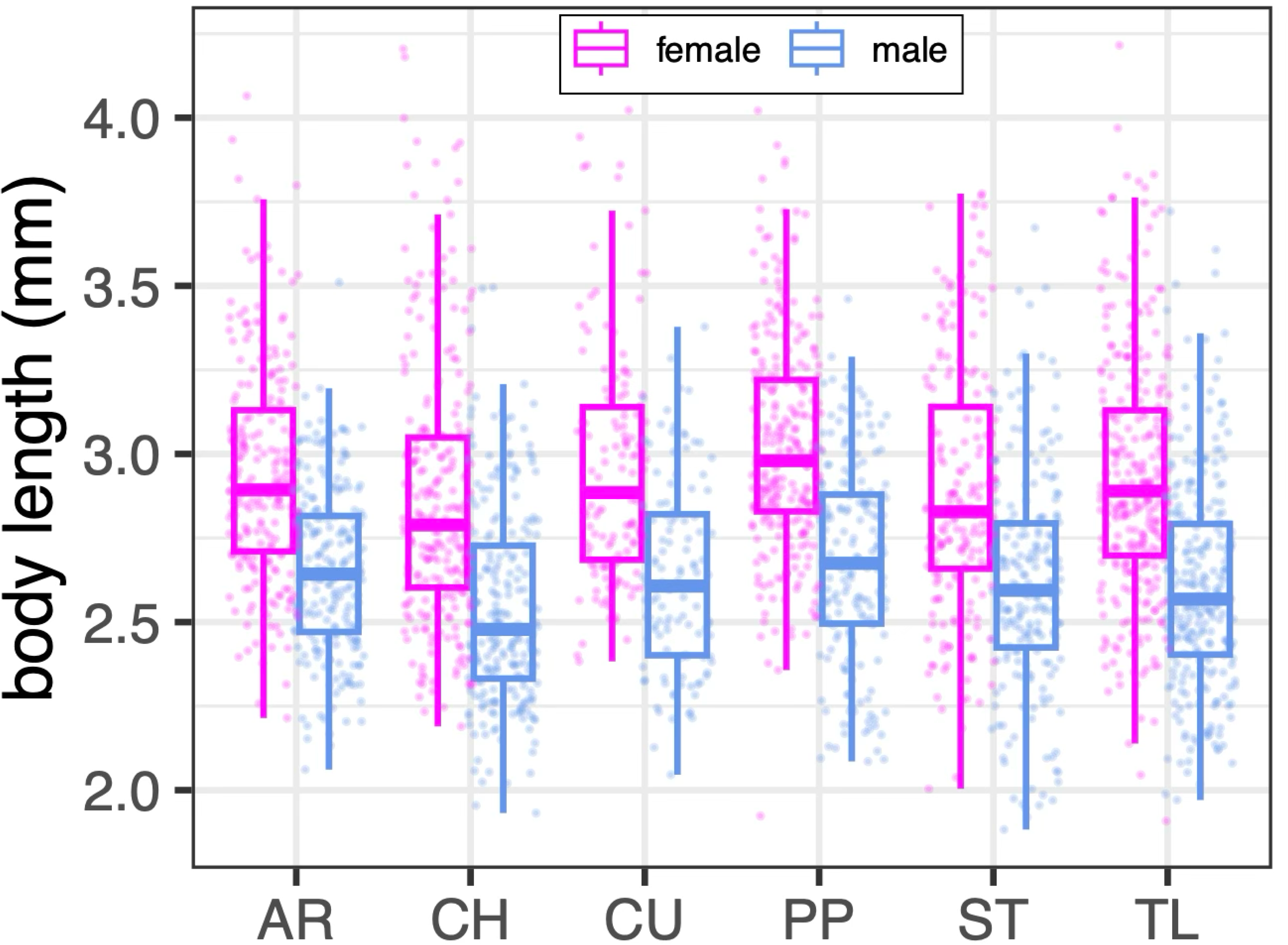
Body size of males and females with different chromosomal inversion arrangements. Boxplots show the distributions of body length (mm) for female (magenta) and male (blue) flies with one of six inversion arrangements (X-axis) measured at 22°C. Each point in the figure represents the measurement for an individual fly.

## Discussion

We aimed to test if coadaption of many linked alleles or pleiotropic effects of individual mutations could better explain the effects of *D. pseudoobscura* chromosomal inversions on life history traits. Individual pleiotropic alleles are expected to result in stronger genetic correlations than multiple linked alleles (Chebib & Guillaume, 2021). Differentiating between the coadaptation and pleiotropy hypotheses requires evaluating trait correlations across environments (such as different temperatures) because genetic correlations may be context-dependent (Sgrò & Hoffmann, 2004; Kapun *et al*., 2016b). In addition, by comparing males and females, we could test for inter-sexual differences and correlations. Sex-differences can provide evidence for sexual antagonism, be useful for understanding life history trade-offs, and allow for an even more extensive test of context-dependent trait correlations (Promislow, 2003; Bonduriansky *et al*., 2008; Pearse *et al*., 2019; McAllester & Pool, 2025). To those ends, we measured lifespan, development time, and body size at multiple temperatures in *D. pseudoobscura* males and females carrying six different chromosomal inversion arrangements.

### Temperature, sex, and genotype affect aging and development time

We detected modest genotype effects on the traits we examined, but strong effects of sex and temperature. For example, higher temperatures universally increased mortality rate, decreased average lifespan, and accelerated development time, as expected for an ectotherm (Angilletta, 2009). Despite elevating the initial mortality rate across all flies, increasing temperatures reduced the rate of aging in some inversion-sex combinations. We also observed substantial sex differences in lifespan, development time, and body size. Notably, our results confirmed previous observations that *D. pseudoobscura* females live longer than males (Taylor & Condra, 1980; Yoon *et al*., 1990). However, the sex differences in mortality rates and development time often depended on inversion genotype and temperature. Similarly, direct effects of inversion genotype on mortality rate and development time depended greatly on sex and temperature, but there was no direct inversion effect on body size.

We observed multiple examples of sex differences in aging depending on the combined effects of inversion genotype and temperature. For example, sex differences in mortality rates were greatest in flies carrying the AR arrangement, which resulted in a separation of the sex-specific survival curves (**Figure 1C**). In addition, within each inversion genotype, we detected more sex differences in mortality rates at 18°C than at the higher temperatures, causing greater separation of the sex-specific survival curves at lower temperatures (**Figure 3A**). Higher temperatures also increased the initial mortality rate—the intercept of the age-dependent mortality rate—in all sex-genotype combinations (**Figure 3B**). The extent of the temperature-dependent increase in mortality rate was largest when females had the CH arrangement genotype. In contrast, the rate of aging (slope) decreased with increasing temperatures in other genotype-sex combinations.

To further illustrate the temperature-dependent effects of inversion genotypes on sex-specific aging, we will contrast the mortality rates of flies with the PP and TL arrangements. Males with the PP arrangement had higher mortality rates than TL males, especially at cooler temperatures (compare intercepts between PP and TL in **Figures 1D and 3B**). While mortality rates generally increased with increasing temperature, the mortality rate of females with the PP arrangement was less affected by the increase in temperature from 18°C to 22°C (compare intercepts in **Figure 3B**). Increasing temperature also decreased the rate of aging (slope of the line in **Figure 3B**) in both males and females with the PP arrangement genotype. Therefore, while PP genotypes had a higher mortality rate than TL flies, the survival of the PP flies was generally less affected by increases in temperature. These results demonstrate the importance of considering environmental context when evaluating genetic effects on life history traits.

We also observed significant effects of both sex and temperature on egg-to-adult development time (**Figure 4A**), but the effects of inversion genotype on development were much weaker. When we did observe inversion effects on development time, they primarily acted to modulate the effects of sex and temperature (**Figure 4B**). For example, flies with the CU arrangement had the largest sex difference in development time and experienced the greatest acceleration of development with increasing temperature. In contrast, there were very small sex differences in development time for flies carrying the CH arrangement at most temperatures. These minimal effects of inversion genotypes on development time are in contrast to the effects of the inversions on aging described above.

### The pace of life and decoupling of life history traits

We observed variation in the degree to which the three traits were affected by inversion genotype. As described above, the *D. pseudoobscura* chromosomal inversions affected sex differences in aging, but the effects of inversion genotypes on development time were much weaker. We additionally observed no effect of inversion genotype on body size. These varying effects of the *D. pseudoobscura* chromosomal inversions on different traits provides some evidence against the pleiotropy hypothesis. Testing the pleiotropy hypothesis, however, requires assessing if the traits are genetically correlated.

The test for genetic correlations across life history traits, we directly compared lifespan and development time across the fly strains we assayed. Life history timing is often positively correlated across stages, such that acceleration of one stage is accompanied by other stages progressing faster. These correlations can result in a consistent pace of life for a given species, producing correlations when comparing across species (Healy *et al*., 2019). It is also predicted that pace of life traits could be genetically correlated within species, and pleiotropic alleles could be responsible for within population variation in pace of life (Réale *et al*., 2010; Dammhahn *et al*., 2018; Arnqvist & Rowe, 2023). We focused our analysis on two non-overlapping life history stages where we identified genotype effects: egg-to-adult development and lifespan (time from pupal emergence until death).

In contrast to the pace of life expectation, we failed to observe positive correlations between development time and adult lifespan. Because we did not measure lifespan and development time in the same animals, we were limited in our ability to examine trait covariance (Lande, 1979; Arnold *et al*., 2008). Nonetheless, we did sample individuals from inbred lines, which allowed us to examine inter-trait genetic correlations. Across all strains and temperatures, there was a negative association between development time and lifespan for males, and the negative relationship was especially pronounced at 22°C (**Figure 5**). In other words, genotypes that were associated with shorter male development time were also associated with longer male lifespans. These negative correlations are consistent with a genetic decoupling of the pace of life between life history stages (Moran, 1994; Collet & Fellous, 2019), providing additional evidence against the pleiotropy hypothesis (Arnqvist & Rowe, 2023).

The genetic decoupling can also be seen within genotypes by considering the effects of specific inversion arrangements on development and aging. For example, flies carrying an AR chromosome developed faster than other inversion genotypes (**Figure 4**). The pace of life hypothesis would predict that AR flies should have a corresponding higher mortality rate or faster rate of aging. However, AR flies had a lower rate of aging than other genotypes, which can be visualized as a flatter slope of age-specific mortality line (**Figures 1D and 3B**). In addition, development of flies with the CU arrangement was accelerated by increasing temperatures relative to other arrangement genotypes (**Figure 4B**). While there was a dramatic acceleration of the rate of aging in CU males at 25°C (steep slope in **Figure 3B**), most of our analyses did not detect a significant difference in the effect of temperature on aging in CU flies relative to other inversion arrangements. In contrast, flies with the TL or PP arrangement had decreased rates of aging as temperature increased (**Figure 3B**), but their development was not differentially affected by temperature when compared to other arrangements (**Figure 4B**). The lack of consistent chromosomal inversion effects on pace of life traits is also evidence against the pleiotropy hypothesis.

Similar to the expected shared pace of life across development and aging, there is an expected correlation between body size and life history traits. The general pattern we would expect is for a slower pace of life to be associated with larger animals (Roff, 1981; Stearns, 1992). In contrast to this expectation, we failed to observe any differences in body sizes across inversion genotypes, despite substantial sex effects on body size (**Figure 6**). This result is consistent with a prior experiment in which two different life history syndromes were selected for in *D. pseudoobscura*, and there was no corresponding evolution of body size (Taylor & Condra, 1980). Notably, the previous experiment did discover changes in third chromosome inversion frequencies in response to selection on life history, consistent with our observation that the third chromosome arrangements affect life history timing but not body size. In summary, the *D. pseudoobscura* inversion arrangements do not have correlated effects on any of the traits we measured in directions that would be consistent with a pace of life or covarying life history traits, providing evidence against the pleiotropy hypothesis.

### Sexual antagonism, pleiotropy, and context-dependent traits

Comparing life history traits between males and females is an alternative approach for identifying genetic correlations. Males and females largely share the same genome, which can generate inter-sexual trait correlations (Lande, 1980). Sexual antagonism can arise if the optimal trait value differs between males and females because alleles that increase fitness in one sex could decrease fitness in the other sex (Bonduriansky & Chenoweth, 2009; van Doorn, 2009). Opposing effects of sexually antagonistic alleles has been proposed explanation for the evolution of senescence (Promislow, 2003; Bonduriansky *et al*., 2008; Adler & Bonduriansky, 2014), and this hypothesis is supported by observed inter-sexual correlations of life history traits within insect species (Archer *et al*., 2012; Berg & Maklakov, 2012). We observed similar strong positive inter-sexual correlations amongst strains for both development time and lifespan in *D. pseudoobscura* (**Figure 2B**). These correlations, along with the intra-sexual replicability of our experiments (**Figure 2A**), are evidence of a genetic effect for the trait variation we observed. However, these inter-sexual correlations alone are not evidence for pleiotropy.

A promising way forward to test for pleiotropy is to consider how environmental variation affects the traits under consideration (Schluter *et al*., 1991; Connallon *et al*., 2018). In the context of our results, the environmental factor that we manipulated was temperature. Sokoloff (1966) argued that raising *D. pseudoobscura* in stressful environments may bring out more phenotypic differences between genotypes, which could then allow for more powerful examinations of trait correlations. In contrast to that prediction, we found that sex and arrangement effects on mortality rate were most pronounced at the lowest temperature we considered (**Figure 3**), which is the least stressful for *D. pseudoobscura* (Druger, 1962). Therefore, considering how extreme temperatures affect genotype-to-phenotype mapping did not provide evidence for pleiotropy.

Comparing our results with prior experiments allows us to identify possible trade-offs, which are indirect evidence for pleiotropy (Paaby & Rockman, 2013). For example, many prior results suggest that the ST arrangement confers traits that increase fitness at warmer temperatures. The ST arrangement, for instance, was observed more often at lower elevations, increased in frequency in summer months, and had higher fitness in a laboratory population at 25°C (Wright & Dobzhansky, 1946; Dobzhansky, 1948). We found that increasing temperature reduced the rate of aging in ST females (flatter slope at higher temperatures in **Figure 3B**), consistent with the ST chromosome conferring a fitness benefit at warmer temperatures. However, we did not observe that effect in males, suggesting that beneficial fitness effects of the ST arrangement at warmer temperatures may be female-specific.

The CH arrangement was similarly hypothesized to increase fitness at warm temperatures. For instance, CH homozygotes had higher egg production and pupal hatching rates than other genotypes at warmer temperature and higher humidity (Heuts, 1947; Tantawy, 1961). In contrast, we observed that increasing temperatures increased the mortality rate of CH females (decreased intercept in **Figure 3B**), providing evidence for a fitness cost at higher temperatures. Consistent with such a temperature-dependent cost, multiple previous studies found that the CH arrangement decreased in frequency in experimental populations starting with equal numbers of AR and CH flies at 25°C (Levine, 1955; Levene *et al*., 1958; Beardmore *et al*., 1960). These results therefore suggest that there are temperature-dependent trade-offs associated with the CH genotype. While trade-offs are consistent with pleiotropic effects (Paaby & Rockman, 2013), they are not direct evidence for pleiotropy.

The AR arrangement provides additional evidence for trade-offs. AR is the youngest (i.e., most recently derived) inversion arrangement we sampled, and it has likely increased in frequency rapidly such that it is the most common across much of the species’ range (Wallace *et al*., 2011; Fuller *et al*., 2017). It is therefore reasonable to assume that flies carrying at least one AR chromosome have high fitness in many environments in which *D. pseudoobscura* is found (Schaeffer, 2008). As described above, the AR arrangement increased in frequency in experimental populations starting with equal numbers of AR and CH flies at 25°C (Levine, 1955; Levene *et al*., 1958; Beardmore *et al*., 1960). We observed that the effects of the AR arrangement on mortality rate differed between females and males (**Figure 1**), with AR females outliving males. We further observed that increasing temperature reduced the rate of aging for AR females but not males (flatter slopes for AR females as temperature increased in **Figure 3B**), suggesting that the AR arrangement may confer a female-specific benefit at warmer temperatures. However, other work found that AR is associated with decreased fitness at warmer temperatures, and there is evidence for benefits of the AR arrangement at cooler, drier temperatures. For instance, the AR arrangement is most abundant at higher elevations and decreases in frequency in summer in natural populations (Dobzhansky, 1948). In addition, a study of AR and CH homo- and hetero-zygotes found that AR homozygotes had the highest egg production at 15°C and were outperformed by CH homozygotes at warmer temperatures (Tantawy, 1961). Notably, AR/CH heterozygotes had high egg production at warmer temperatures, suggesting a complex relationship between dominance and temperature on the phenotype. Furthermore, the pupal hatching rate of flies with the AR arrangement was lower than other genotypes at higher humidities (Heuts, 1947). These conflicting results are consistent with temperature- and sex-dependent trade-offs in the effects of the AR arrangement across life history traits. As mentioned above, trade-offs are consistent with the pleiotropy hypothesis, but they are not direct evidence for pleiotropy.

In contrast to the AR arrangement, TL is one of the oldest inversion arrangements that we sampled (Wallace *et al*., 2011), and TL chromosomes are rare across most of the species’ range (Dobzhansky, 1944; Anderson *et al*., 1991). Given the low frequency of TL, we may expect TL flies to have reduced fitness in many environmental conditions. It is therefore surprising that TL flies had the lowest mortality rate and longest lifespan at 18°C and 22°C (**Figures 1 and 2**). We also found that the TL arrangement was associated with faster development—in particular, an acceleration of development at warmer temperatures (**Figure 4**). Similarly, a previous laboratory experiment that selected for faster developing life history syndromes observed an increase in the frequency of the TL arrangement across multiple replicates (Taylor & Condra, 1980). Given these results, it is possible that the TL chromosome confers a temperature-dependent “live fast” strategy, but we have yet to discover any measurable “die young” costs to this strategy. Additional work is therefore needed to test for life-history trade-offs that could explain the low frequency of the TL arrangement given the fitness benefits that it appears to confer.

### Evaluating the coadaptation and pleiotropy hypotheses

We aimed to test the coadaptation and pleiotropy hypotheses for the effects of *D. pseudoobscura* chromosomal inversions on life history traits and organismal fitness more generally. As described above, we did not detect genetic correlations that would be consistent with the pleiotropy hypothesis. However, by contrasting our results with prior work we did identify potential life history trade-offs, which provide indirect evidence for pleiotropy. In comparison, other previous work identified linkage disequilibrium within *D. pseudoobscura* inversion haplotypes that is suggestive of epistatic selection, providing evidence for the coadaptation hypothesis (Schaeffer *et al*., 2003, 2024; Fuller *et al*., 2017).

There are multiple explanations for why we did not identify genetic correlations that directly supported the pleiotropy hypothesis. For one, life history traits affected by the *D. pseudoobscura* chromosomal inversions may indeed not be correlated. However, there may be an inherent independence of traits separated by metamorphosis (Moran, 1994; Collet & Fellous, 2019), which prevents genetic correlation between larval and adult life history timing. We tested for genetic correlations between one larval trait and one adult trait, which may not be an appropriate method for identifying genetic correlations if larval and adult timing decoupled for other reasons. We therefore may need to consider more trait correlations within each developmental stage to test the pleiotropy hypothesis.

Another reason we may have not detected evidence for pleiotropy is that we only measured traits in inversion homozygotes. Extensive prior work has shown that the *D. pseudoobscura* inversions have non-additive phenotypic and fitness effects that depend on the diploid genotype (Wright & Dobzhansky, 1946; Levene *et al*., 1954; Tantawy, 1961). Importantly genetic variation underlying antagonistic pleiotropy is more likely to persist in a population if the alleles have different dominance coefficients in two contexts, in what is known as dominance reversal (Curtsinger *et al*., 1994; Grieshop *et al*., 2024). Recent work has identified examples of dominance reversals in natural populations (Grieshop & Arnqvist, 2018), including sex- and environment-dependent dominance of the fitness effects of chromosomal inversions (Pearse *et al*., 2019; Mérot *et al*., 2020; Durmaz Mitchell *et al*., 2025). We therefore may not have detected genetic correlations associated with pleiotropy because we did not test for dominance reversal in heterozygotes.

However, non-additive fitness effects are also consistent with the coadaptation hypothesis. When Dobzhansky (1949) presented the coadaptation hypothesis to explain the maintenance of the *D. pseudoobscura* inversion polymorphism, a key component was the increased fitness of inversion heterozygotes. Subsequent work has indeed shown that when an inversion polymorphism is maintained by coadaptation, there is a heterozygote advantage created because recombination is only suppressed in inversion heterozygotes (Charlesworth, 1974; Charlesworth & Flatt, 2021; Berdan *et al*., 2023). Moreover, Wright and Dobzhansky (1946) showed that sex-specific dominance can further contribute to the maintenance of the *D. pseudoobscura* inversion polymorphism under the coadaptation hypothesis. Therefore, non-additive dominance may be consistent with both the coadaptation and pleiotropy hypotheses.

We thus conclude that we identified no direct evidence that the pleiotropy hypothesis can explain the phenotypic and fitness effects of the *D. pseudoobscura* chromosomal inversions. However, we did detect some trade-offs that are consistent with the pleiotropy hypothesis when comparing our results with previously published work, meaning that the pleiotropy hypothesis cannot be completely rejected. Additional experiments are thus needed to further differentiate between the coadaptation and pleiotropy hypotheses for the fitness effects of *D. pseudoobscura* inversions.

Future work should focus on two specific questions. First, most of the previous phenotypic and fitness assays in *D. pseudobscura* were performed in different laboratories using a variety wild-derived isolates. This limits our ability to conduct an integrated analysis of inter-sexual trait covariance that would be required to test for pleiotropy or antagonistic effects (Lande, 1980; Fry, 1993; Bonduriansky & Chenoweth, 2009). Formal tests of trait covariance are required to test the pleiotropy hypothesis using trait measurements from the same individuals from the same genotypes sampled in the same laboratory environments (Chebib & Guillaume, 2017). Second, these measurements should consider both homo- and hetero-karyotypic individuals to test for heterozygote advantages and dominance reversals. Therefore, future tests of the pleiotropy hypothesis should examine multiple phenotypes in homo- and hetero-karyotypic individuals across the same wild-derived chromosomes.

## Acknowledgements

Stephen Schaeffer kindly shared the fly strains used in these experiments. Salam Azim assisted with the experiments to measure lifespan. We received valuable feedback on experimental design, data analysis, and writing from members of the IISAGE consortium.

## Funding

This work was supported by the National Science Foundation under grant 2213824 to Nicole Riddle.

## Conflict of Interest Statement

The authors declare that they have no conflicts of interest.

## Data Availability

All code and data to repeat the analysis are available as Supplementary Files at https://doi.org/10.18738/T8/EKYDQQ

## Supplementary Tables

**Supplementary Table 1.** Sample size of flies from each strain for each assay.

| str<sup>1</sup> | arr<sup>2</sup> | lifespan<br>22C (trial 1) |  | lifespan<br>18C |  | lifespan<br>22C (trial 2) |  | lifespan<br>25C |  | dev time<br>18C |  | dev time<br>22C |  | dev time<br>25C |  | body size |  |
| --- | --- | --- | --- | --- | --- | --- | --- | --- | --- | --- | --- | --- | --- | --- | --- | --- | --- |
|  |  | f | m | f | m | f | m | f | m | f | m | f | m | f | m | f | m |
| 3 | AR | 35 | 27 | 9 | 15 | 29 | 21 | 41 | 36 | 12 | 17 | 37 | 36 | 34 | 32 | 32 | 25 |
| 4 | AR | 36 | 24 | 20 | 17 | 21 | 18 | 10 | 3 | 14 | 24 | 86 | 84 | 20 | 15 | 46 | 54 |
| 5 | AR | 50 | 23 | 36 | 26 | 40 | 35 | 0 | 0 | 17 | 22 | 49 | 36 | 0 | 0 | 12 | 16 |
| 6 | AR | 32 | 12 | 0 | 0 | 0 | 0 | 0 | 0 | 0 | 0 | 0 | 0 | 0 | 0 | 0 | 0 |
| 7 | AR | 24 | 14 | 27 | 12 | 15 | 7 | 11 | 12 | 63 | 51 | 25 | 34 | 15 | 16 | 25 | 21 |
| 8 | AR | 80 | 45 | 52 | 44 | 39 | 23 | 47 | 39 | 140 | 151 | 106 | 68 | 22 | 17 | 60 | 60 |
| 14 | AR | 65 | 48 | 16 | 21 | 37 | 21 | 24 | 20 | 29 | 40 | 86 | 91 | 0 | 0 | 51 | 47 |
| 17 | PP | 90 | 47 | 54 | 33 | 40 | 30 | 50 | 84 | 72 | 74 | 196 | 200 | 61 | 77 | 67 | 50 |
| 18 | PP | 22 | 11 | 21 | 10 | 16 | 14 | 12 | 6 | 30 | 48 | 0 | 0 | 0 | 0 | 2 | 49 |
| 19 | PP | 96 | 64 | 25 | 15 | 29 | 18 | 33 | 14 | 77 | 85 | 47 | 24 | 12 | 8 | 75 | 18 |
| 20 | PP | 0 | 0 | 9 | 7 | 10 | 9 | 17 | 11 | 65 | 69 | 0 | 0 | 0 | 0 | 19 | 74 |
| 22 | PP | 0 | 0 | 7 | 10 | 7 | 7 | 0 | 0 | 22 | 15 | 0 | 0 | 0 | 0 | 5 | 23 |
| 24 | PP | 98 | 51 | 40 | 18 | 37 | 31 | 16 | 11 | 32 | 20 | 41 | 35 | 5 | 6 | 104 | 54 |
| 26 | ST | 82 | 51 | 21 | 25 | 38 | 21 | 13 | 14 | 96 | 105 | 47 | 60 | 11 | 10 | 24 | 39 |
| 28 | ST | 66 | 54 | 34 | 20 | 38 | 32 | 60 | 54 | 114 | 84 | 114 | 101 | 55 | 64 | 56 | 88 |
| 31 | ST | 22 | 24 | 19 | 17 | 17 | 9 | 18 | 2 | 58 | 62 | 31 | 27 | 40 | 36 | 31 | 41 |
| 33 | ST | 87 | 60 | 27 | 21 | 49 | 28 | 36 | 48 | 50 | 37 | 120 | 139 | 4 | 9 | 86 | 16 |
| 34 | CH | 67 | 43 | 14 | 16 | 28 | 8 | 9 | 8 | 50 | 59 | 44 | 41 | 5 | 6 | 54 | 39 |
| 35 | CH | 25 | 25 | 16 | 27 | 16 | 12 | 0 | 0 | 20 | 13 | 12 | 16 | 0 | 0 | 15 | 65 |
| 36 | CH | 11 | 7 | 31 | 26 | 15 | 12 | 0 | 0 | 25 | 30 | 12 | 23 | 7 | 3 | 34 | 1 |
| 38 | CH | 58 | 29 | 17 | 19 | 26 | 22 | 16 | 18 | 41 | 37 | 27 | 25 | 6 | 8 | 64 | 73 |
| 42 | CH | 61 | 47 | 27 | 21 | 37 | 43 | 11 | 7 | 21 | 21 | 74 | 62 | 13 | 10 | 87 | 78 |
| 43 | TL | 87 | 53 | 20 | 23 | 23 | 31 | 38 | 25 | 111 | 121 | 96 | 109 | 116 | 113 | 82 | 48 |
| 46 | TL | 31 | 14 | 24 | 23 | 28 | 18 | 6 | 8 | 64 | 69 | 74 | 56 | 0 | 0 | 39 | 63 |
| 47 | TL | 35 | 21 | 18 | 15 | 23 | 16 | 8 | 4 | 29 | 29 | 32 | 32 | 14 | 10 | 60 | 45 |
| 48 | TL | 14 | 13 | 26 | 14 | 19 | 20 | 11 | 5 | 33 | 34 | 32 | 21 | 0 | 0 | 64 | 37 |
| 49 | TL | 48 | 33 | 24 | 12 | 24 | 9 | 32 | 19 | 36 | 31 | 52 | 51 | 18 | 15 | 54 | 112 |
| 53 | CU | 74 | 39 | 18 | 26 | 46 | 34 | 9 | 19 | 65 | 65 | 103 | 108 | 12 | 4 | 133 | 0 |

1. Strain identification number. 2. Chromosomal inversion arrangement in the strain genotype. - 3.

## Notes

https://doi.org/10.18738/T8/EKYDQQ

### Competing Interest Statement

The authors have declared no competing interest.

### Summary of Updates

The manuscript has been revised to better motivate they study and explain the meaning of the results.

