## Supplemental File S1 for "*Drosophila pseudoobscura* third chromosome inversion arrangements have temperature-dependent and sex-specific effects on life history traits"

### Supplementary File S1: Analyze lifespan data from *Drosophila pseudoobscura* at 22C

2026-09-03

#### Overview

The analysis described below was performed on survival data from *Drosophila pseudoobscura* collected in 2024. Males and females carrying one of six different third chromosome inversions arrangements were sampled. The flies were sampled from 26 different strains, each of which is homozygous for one of six different chromosomal inversion arrangements.

Individual flies were collected upon emergence and stored in vials with standard *Drosophila* medium until they died. Each fly was transferred to a new vial every ~7 days until it died. The date of emergence from the pupal case (birth) and death were recorded for each fly. These dates were used to calculate the age at death (in days since emergence from the pupal case). Observations were predominately made on weekdays. If a fly died over an unobserved weekend or holiday, the midpoint of the unobserved period was recorded as the age at death.
Flies that escaped were marked as censored data.

#### Install and Load Required Packages

```
if (!require("tidyr")) install.packages("tidyr")
if (!require("ggplot2")) install.packages("ggplot2")
if (!require("ggfortify")) install.packages("ggfortify")
if (!require("survival")) install.packages("survival")
if (!require("survminer")) install.packages("survminer")
if (!require("flexsurv")) install.packages("flexsurv")
if (!require("lme4")) install.packages("lme4")

library(tidyr)
library(ggplot2)
library(ggfortify)
library(survival)
library(survminer)
library(flexsurv)
library(lme4)
```

#### Load, Prepare, and QC the Datasel

Load the survival data from file.

```
aging <- read.delim("Dpse_aging_strains_2025_01_10.txt")
head(aging)
```

```
##   Strain    sex     Birth     Death Age.at.death
## 1      6 female 10-Sep-24 11-Sep-24            1
## 2      6 female 10-Sep-24 11-Sep-24            1
## 3      6 female 10-Sep-24 11-Sep-24            1
```

```
## 4     19   male 10-Sep-24 11-Sep-24            1
## 5      4   male 10-Sep-24 12-Sep-24            2
## 6     19 female 10-Sep-24 12-Sep-24            2
##                                  notes censrec winter_cens winter_death
## 1 stuck in food -- killed before awake?       1          NA           NA
## 2 stuck in food -- killed before awake?       1          NA           NA
## 3 stuck in food -- killed before awake?       1          NA           NA
## 4 stuck in food -- killed before awake?       1          NA           NA
## 5                                             1          NA           NA
## 6                                             1          NA           NA
```

Each row in the aging dataframe is a single fly, and the columns contain information about the fly. The columns are strain ID, sex of the fly, birth date, death date, age at death, any notes (related to censoring), if they fly is censored (0) or not (1), and information about censored data over winter holidays (see below).

Build a dataframe out of the birth dates to count how many flies have each birth date.

```
births <- levels(factor(aging$Birth))

Births.df <- data.frame(
    birth = births,
    count = NA
)
for(i in 1:length(births)){
    Births.df$count[i] <- length(subset(aging, Birth==births[i])[,1])
}
Births.df
```

```
##        birth count
## 1   1-Oct-24   137
## 2  10-Oct-24    88
## 3  10-Sep-24    76
## 4  11-Sep-24    92
## 5  12-Sep-24    99
## 6  13-Sep-24    96
## 7  15-Oct-24   231
## 8  16-Oct-24   163
## 9  17-Oct-24   145
## 10 17-Sep-24    50
## 11 18-Oct-24   192
## 12 18-Sep-24    59
## 13 19-Sep-24    41
## 14  2-Oct-24   111
## 15 20-Sep-24   121
## 16 24-Sep-24    99
## 17 25-Sep-24    93
## 18 26-Sep-24   131
## 19 27-Sep-24   102
## 20  3-Oct-24   175
```

Make a bar graph of these data to show sampling from each birth date. Birth dates will be treated as batches in the statistical analysis.

```
Births.df %>%
  separate_wider_delim(birth,
    delim="-", names=c("day", "month", "year"), cols_remove = FALSE
  ) -> Birth.graph
```

```r
Birth.graph$month[Birth.graph$month=="Sep"] <- " Sep"
Birth.graph$days <- as.numeric(Birth.graph$day)
Birth.graph$day[Birth.graph$days < 10] <- paste(" ", Birth.graph$day[Birth.graph$days<10], sep="")
Birth.graph$date <- paste(Birth.graph$month, Birth.graph$day)

ggplot(Birth.graph, aes(x=date, y=count)) +
    geom_bar(stat="identity") +
    scale_x_discrete(guide = guide_axis(angle = 90), name=NULL) +
    scale_y_continuous(name="number of flies") +
    theme_bw()
```

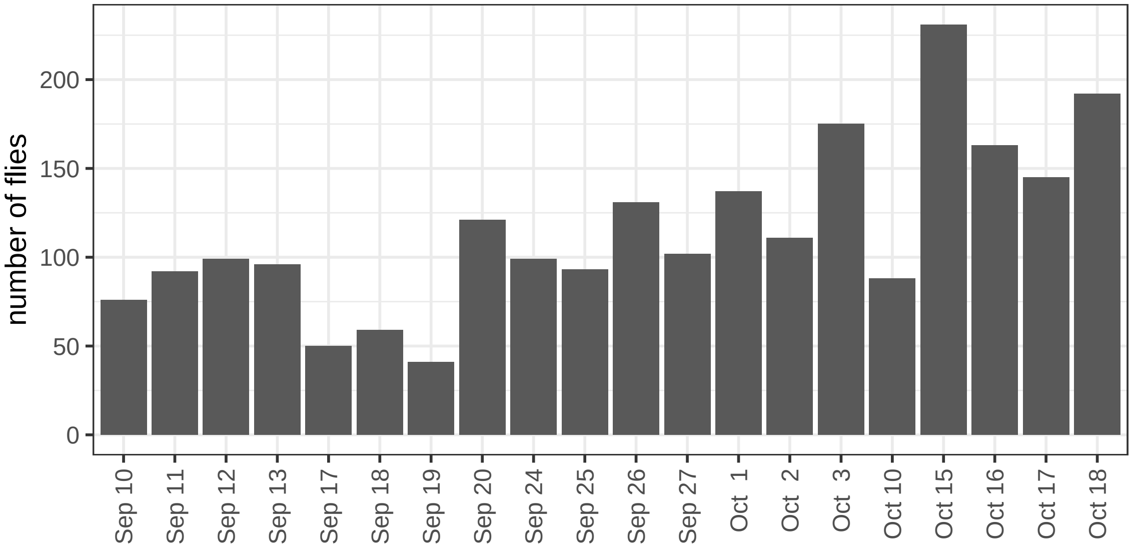


Next, load the information about the strains from a file.

```r
strains <- read.delim("Dpse-strains.tsv")
head(strains)
```

```
##   ID           accession arrangement isogen balancer
## 1  3 DSSC 14011_0121.225          AR DM1015        L
## 2  4 DSSC 14011_0121.226          AR DM1050        L
## 3  5 DSSC 14011_0121.227          AR DM1056        L
## 4  6 DSSC 14011_0121.228          AR DM1088        L
## 5  7 DSSC 14011_0121.231          AR  KB635        L
## 6  8 DSSC 14011_0121.232          AR  KB652        L
##                                strain count
## 1 3 DSSC 14011_0121.225 AR_DM1015_L     1
## 2 4 DSSC 14011_0121.226 AR_DM1050_L     2
## 3 5 DSSC 14011_0121.227 AR_DM1056_L     3
## 4 6 DSSC 14011_0121.228 AR_DM1088_L     4
## 5  7 DSSC 14011_0121.231 AR_KB635_L     5
## 6  8 DSSC 14011_0121.232 AR_KB652_L     6
```

The `strains` object has the strain ID, which will be used to match the strain information to the individual flies in the `aging` data. The remaining columns have the accession ID for the strain, the chromosomal arrangement (inversion name) for the strain, strain name (isogen), and the balancer chromosome that was used to extract the arrangement from the originating isofemale line and homozygose it.

Attach the arrangement, isogen, and balancer information for each fly in the `aging` data frame, using the `match()` function.

```
aging$arr <- strains$arrangement[match(aging$Strain, strains$ID)]
aging$isogen <- strains$isogen[match(aging$Strain, strains$ID)]
aging$bal <- strains$balancer[match(aging$Strain, strains$ID)]
```

Next, evaluate how well each strain is represented in the data. To do so, first create a table with all strains assayed.

```
aging_strains <- data.frame(
    strains = levels(factor(aging$Strain))
)
aging_strains$arr <- strains$arrangement[match(aging_strains$strains, strains$ID)]
aging_strains$isogen <- strains$isogen[match(aging_strains$strains, strains$ID)]
aging_strains$bal <- strains$balancer[match(aging_strains$strains, strains$ID)]
for(i in 1:length(aging_strains$strains)){
    aging_strains$females[i] <- length(subset(aging,
                                    Strain==aging_strains$strains[i] & sex=="female")[,1])
    aging_strains$males[i] <- length(subset(aging,
                                    Strain==aging_strains$strains[i] & sex=="male")[,1])
}
aging_strains %>% pivot_longer(
      cols = c("females", "males"),
      names_to = "sex",
      values_to = "count"
    ) %>%
ggplot(aes(x=strains, y=count, fill=sex)) +
    geom_bar(stat="identity") +
    scale_x_discrete() +
    scale_y_continuous(name="number of flies") +
    scale_fill_manual(values = c("females" = "magenta", "males" = "cornflowerblue")) +
  facet_wrap(~sex, ncol=1) +
    theme_bw() +
  theme(legend.position="none")
```

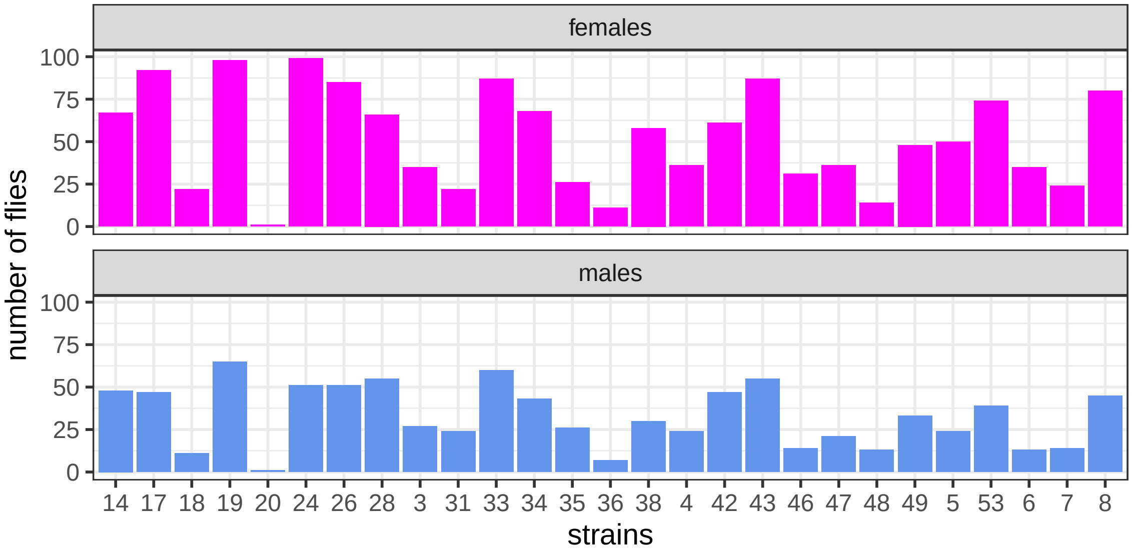


Note that there are very few individuals from strain 20 (only sampled 2 flies total). We will remove strain 20 from our analysis.

```r
aging <- subset(aging, Strain!=20)
```

Add some annotations to the data in order to handle some odd features of our data. In particular, there was a period over the winter holidays where the flies were not observed. To handle any unobserved deaths, treat any fly that died over the winter holiday as censored data. Do this by creating a new `aging_winterCens` object, and mark flies with 'winter' in their `notes` column as censored (coded with 0). The age at censoring is the day before winter holiday (last observation alive).

```r
aging_winterCens <- aging[,c(1:5,7,10:12)]
aging_winterCens[(grepl("winter", aging$notes)),5] <- aging[(grepl("winter", aging$notes)),8]
aging_winterCens[(grepl("winter", aging$notes)),6] <- 0
subset(aging_winterCens, grepl('23-Dec-24', Death))
```

```
##      Strain    sex     Birth     Death Age.at.death censrec arr     isogen bal
## 2171      8   male 11-Sep-24 23-Dec-24          102       1  AR      KB652   L
## 2172     46   male 13-Sep-24 23-Dec-24          100       1  TL    SCI12-2   B
## 2173     34   male 25-Sep-24 23-Dec-24           88       1  CH       JR32   B
## 2174     42 female 25-Sep-24 23-Dec-24           88       1  CH     MSH202   B
## 2175     48   male  1-Oct-24 23-Dec-24           82       1  TL SPE123_5-1   B
## 2176     19   male  3-Oct-24 23-Dec-24           80       1  PP     DM1038   B
## 2177     33 female 10-Oct-24 23-Dec-24           73       1  ST     MSH217   B
## 2178     31 female 10-Oct-24 23-Dec-24           73       1  ST       JR91   L
## 2179     19 female 15-Oct-24 23-Dec-24           68       1  PP     DM1038   B
## 2180     34 female 15-Oct-24 23-Dec-24           68       1  CH       JR32   B
## 2181     24 female 16-Oct-24 23-Dec-24           67       1  PP     DM1085   B
## 2182     49 female 16-Oct-24 23-Dec-24           67       1  TL SPE123_6-3   B
## 2183     33 female 16-Oct-24 23-Dec-24           67       1  ST     MSH217   B
## 2184     19   male 16-Oct-24 23-Dec-24           67       1  PP     DM1038   B
## 2185     14 female 17-Oct-24 23-Dec-24           66       1  AR     MSH126   L
## 2186     19   male 18-Oct-24 23-Dec-24           65       1  PP     DM1038   B
## 2187     26 female 18-Oct-24 23-Dec-24           65       1  ST      JR138   L
```

Examine the distribution of age at death to test for additional peculiarities in the data. Plot a histogram of age at death.

```r
hist(aging_winterCens$Age.at.death,
     breaks=max(aging_winterCens$Age.at.death),
     ylab="number of flies",
     xlab="age at death (days)",
     main = "")
```

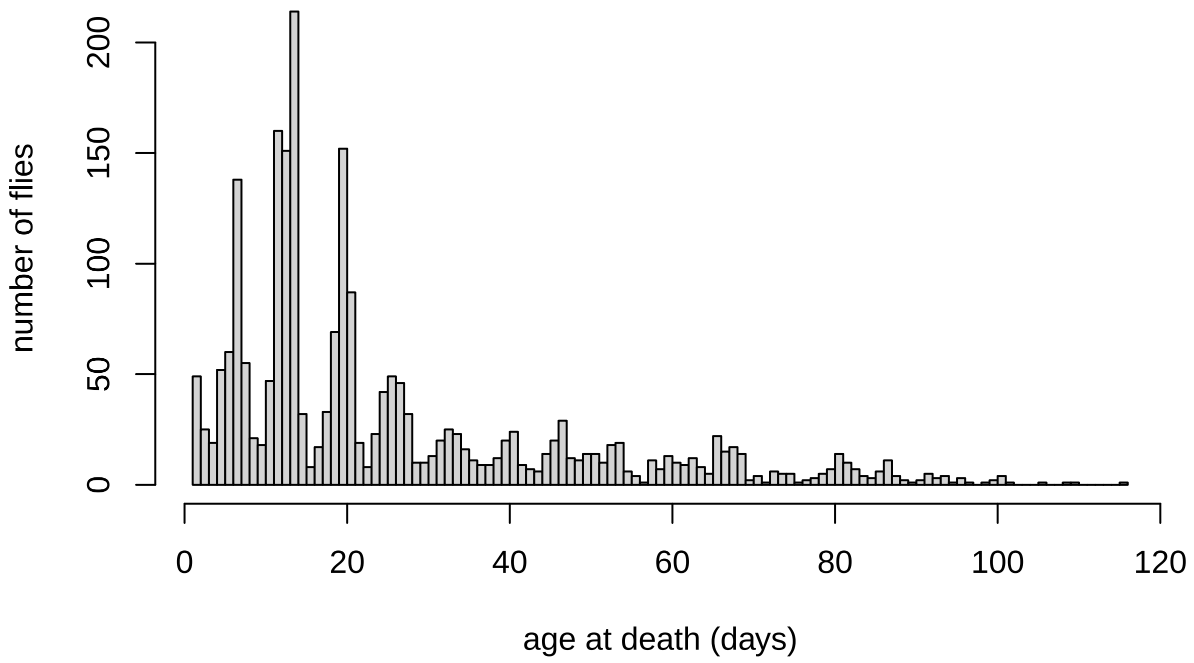


The histogram above reveals an oscillating pattern of ages at death, which is likely attributable to not recording deaths on weekends (so that there is an excess of counts on Monday) and deaths shortly following transfer to a new vial. These effects should create noise in the data that reduces any signal of genotype or sex on aging. However, we do not expect this noise to bias our results because there should not be any relationship between the bias and our predictors (sex or genotype). In addition, there is a small peak of flies dead at 1 day old (i.e., a lot of flies died one day after collection), suggesting that they may have been collected in poor condition. To evaluate if this affects the analysis, create a second version of the data that excludes flies that died at 1 day old by subsetting for individuals that lived > 1 day (or `Age.at.death >= 2`).

```
aging_winterCens_no1 <- subset(aging_winterCens, Age.at.death >= 2)
```

Now there are two versions of the data to analyze going forward (including or excluding flies that died after one day).

##### Use Cox proportional hazard model to test for genotype and sex effects

Write function to compare different Cox proportional hazard models. Model birth (emergence) date as a frailty factor, and consider the effect of strain (isogen). Use `anova()` to compare how well nested models fit the data.

```
coxph.frailty <- function(agingData){
  sex <- coxph(
    Surv(Age.at.death, censrec) ~ sex + frailty(Birth),
    agingData)

  arr <- coxph(
    Surv(Age.at.death, censrec) ~ arr + arr:isogen + frailty(Birth),
    agingData)

    sex_arr <- coxph(
      Surv(Age.at.death, censrec) ~ sex + arr + arr:isogen + frailty(Birth),
      agingData)

    sex_arr_int <- coxph(
```

```
      Surv(Age.at.death, censrec) ~ sex + arr + sex*arr + arr:isogen + frailty(Birth),
      agingData)

    data.frame(
      factor = c("sex", "arr", "sex-by-arr"),
      pval = c(
        anova(sex_arr, arr)$`Pr(>|Chi|)`[2],
        anova(sex_arr, sex)$`Pr(>|Chi|)`[2],
        anova(sex_arr_int, sex_arr)$`Pr(>|Chi|)`[2]
      )
    )
}
```

Compare model fit for complete dataset.

```
coxph_frailty_winterCens <- coxph.frailty(aging_winterCens)
coxph_frailty_winterCens
```

```
##       factor         pval
## 1        sex 7.565240e-03
## 2        arr 1.040675e-59
## 3 sex-by-arr 2.643030e-02
```

Compare model fit for data excluding 1 day old deaths.

```
coxph_frailty_winterCens_no1 <- coxph.frailty(aging_winterCens_no1)
coxph_frailty_winterCens_no1
```

```
##       factor         pval
## 1        sex 2.173126e-11
## 2        arr 1.037002e-60
## 3 sex-by-arr 6.689470e-01
```

There is strong evidence that both sex and arrangement improve the fit of the model. The interaction effect (between sex and arrangement) is only significant when we include the 1 day old deaths.

We can extract the coefficients of the best fitting models:

```
aging_winterCens_sex_arr <- coxph(
  Surv(Age.at.death, censrec) ~ sex + arr + arr:isogen + frailty(Birth),
  aging_winterCens)

aging_winterCens_no1_sex_arr <- coxph(
  Surv(Age.at.death, censrec) ~ sex + arr + arr:isogen + frailty(Birth),
  aging_winterCens_no1)

aging_winterCens_sex_arr_int <- coxph(
  Surv(Age.at.death, censrec) ~ sex + arr + sex*arr + arr:isogen + frailty(Birth),
  aging_winterCens)

aging_winterCens_no1_sex_arr_int <- coxph(
  Surv(Age.at.death, censrec) ~ sex + arr + sex*arr + arr:isogen + frailty(Birth),
  aging_winterCens_no1)
```

Subset only the top seven rows of the table to view the effects of sex and arrangements.

```
summary(aging_winterCens_sex_arr)$coefficients[1:7,]
```

```
##                       coef   se(coef)        se2     Chisq      DF             p
## sexmale        0.3021189 0.04540204 0.04529193  44.27973  1.0000  2.846481e-11
## arrCH          3.9155310 0.13677886 0.13659574 819.48949  1.0000 3.124332e-180
## arrCU          3.5401367 0.13617764 0.13570151 675.81600  1.0000 5.430088e-149
## arrPP          3.5430584 0.13075437 0.13037821 734.25046  1.0000 1.066516e-161
## arrST          3.5994424 0.12798443 0.12761692 790.96307  1.0000 4.975841e-174
## arrTL          3.6727260 0.14958501 0.14918947 602.83843  1.0000 4.040354e-133
## frailty(Birth)        NA         NA         NA  67.13210 15.4442  2.049409e-08
```

Subset only the top twelve rows of the table to view the effects of sex, arrangements, and interactions.

```
summary(aging_winterCens_sex_arr_int)$coefficients[1:12,]
```

```
##                        coef   se(coef)        se2      Chisq       DF
## sexmale           0.5669903 0.09566421 0.09545504  35.127915  1.00000
## arrCH             3.8314931 0.15159938 0.15135344 638.765197  1.00000
## arrCU             3.4106775 0.15916936 0.15879646 459.158238  1.00000
## arrPP             3.3858502 0.14311093 0.14281405 559.744587  1.00000
## arrST             3.5028790 0.14052671 0.14017575 621.344556  1.00000
## arrTL             3.5806239 0.16259205 0.16220432 484.974387  1.00000
## frailty(Birth)           NA         NA         NA  71.862653 15.77695
## sexmale:arrCH    -0.4288189 0.14531566 0.14510398   8.708092  1.00000
## sexmale:arrCU    -0.3465319 0.22427685 0.22355444   2.387358  1.00000
## sexmale:arrPP    -0.2698573 0.13688153 0.13673688   3.886679  1.00000
## sexmale:arrST    -0.2579478 0.13808227 0.13782422   3.489697  1.00000
## sexmale:arrTL    -0.4259727 0.15046506 0.15013890   8.014791  1.00000
##                             p
## sexmale           3.087430e-09
## arrCH            6.201433e-141
## arrCU            7.329488e-102
## arrPP            9.553886e-124
## arrST            3.813482e-137
## arrTL            1.767419e-107
## frailty(Birth)    3.927007e-09
## sexmale:arrCH     3.168006e-03
## sexmale:arrCU     1.223202e-01
## sexmale:arrPP     4.867058e-02
## sexmale:arrST     6.175189e-02
## sexmale:arrTL     4.639684e-03
```

We can extract the arrangement effects within each sex by analyzing only the data from each separately. First, we will set TL as a reference strain because it is the longest lived (see graphs below).

```
aging_winterCens$arr_factor <- factor(aging_winterCens$arr,
                                      levels = c("TL", "AR", "CH", "CU", "PP", "ST"))
```

Use the TL reference for the model.

```
aging_winterCens_female_arr <- coxph(
  Surv(Age.at.death, censrec) ~ arr_factor + arr:isogen + frailty(Birth),
  subset(aging_winterCens, sex=="female")
)
summary(aging_winterCens_female_arr)$coefficients[1:6,]
```

```
##                     coef  se(coef)       se2    Chisq       DF            p
## arr_factorAR    1.646970 0.1955360 0.1945928 70.94438  1.00000 3.674382e-17
## arr_factorCH    1.765824 0.2013252 0.2004767 76.93051  1.00000 1.770799e-18
```

```
## arr_factorCU   1.350275 0.1919607 0.1912555 49.47891  1.00000 2.005160e-12
## arr_factorPP   0.867197 0.1842721 0.1838681 22.14708  1.00000 2.525392e-06
## arr_factorST   1.422116 0.1869883 0.1861047 57.84174  1.00000 2.840770e-14
## frailty(Birth)       NA        NA        NA 51.01741 14.50753 5.870201e-06
```

```
aging_winterCens_male_arr <- coxph(
  Surv(Age.at.death, censrec) ~ arr_factor + arr:isogen + frailty(Birth),
  subset(aging_winterCens, sex=="male")
)
summary(aging_winterCens_male_arr)$coefficients[1:6,]
```

```
##                      coef  se(coef)       se2      Chisq       DF           p
## arr_factorAR    0.1718612 0.2369666 0.2350797  0.5259948  1.00000 0.468295617
## arr_factorCH    0.2284409 0.2380385 0.2352210  0.9209873  1.00000 0.337215866
## arr_factorCU   -0.1419979 0.2471495 0.2447829  0.3300991  1.00000 0.565600745
## arr_factorPP   -0.4570851 0.2346166 0.2327949  3.7955656  1.00000 0.051388507
## arr_factorST    0.0319134 0.2289338 0.2266372  0.0194324  1.00000 0.889133963
## frailty(Birth)         NA        NA        NA 30.9784877 10.90341 0.001047039
```

CH appears to have the lowest hazard ratio in males, so we will repeat the analysis using CH as the reference in males.

```
aging_winterCens$CH_factor <- factor(
  aging_winterCens$arr,
  levels = c("CH", "TL", "AR", "CU", "PP", "ST"))
aging_winterCens_male_CH <- coxph(
  Surv(Age.at.death, censrec) ~ CH_factor + arr:isogen + frailty(Birth),
  subset(aging_winterCens, sex=="male")
)
summary(aging_winterCens_male_CH)$coefficients[1:6,]
```

```
##                       coef  se(coef)       se2       Chisq       DF           p
## CH_factorTL    -0.22844094 0.2380385 0.2352210  0.92098733  1.00000 0.337215866
## CH_factorAR    -0.05657972 0.2120463 0.2100054  0.07119677  1.00000 0.789602190
## CH_factorCU    -0.37043883 0.2242561 0.2212743  2.72862935  1.00000 0.098563842
## CH_factorPP    -0.68552601 0.2121658 0.2096744 10.43991497  1.00000 0.001233209
## CH_factorST    -0.19652755 0.2058169 0.2022578  0.91176940  1.00000 0.339645381
## frailty(Birth)          NA        NA        NA 30.97848774 10.90341 0.001047039
```

We can also extract the sex effects within arrangements by subsetting for arrangement and comparing males and females. First, we do this for the full data.

```
aging_winterCens_AR <- coxph(
  Surv(Age.at.death, censrec) ~ sex + isogen + frailty(Birth),
  subset(aging_winterCens, arr=="AR")
)
summary(aging_winterCens_AR)$coefficients[1,]
```

```
##         coef     se(coef)          se2        Chisq           DF            p
## 5.032329e-01 9.759075e-02 9.630661e-02 2.659014e+01 1.000000e+00 2.515236e-07
```

```
aging_winterCens_CH <- coxph(
  Surv(Age.at.death, censrec) ~ sex + isogen + frailty(Birth),
  subset(aging_winterCens, arr=="CH")
)
summary(aging_winterCens_CH)$coefficients[1,]
```

```
##      coef   se(coef)       se2      Chisq         DF          p
```

```
## 0.1279738 0.1125608 0.1108491 1.2926119 1.0000000 0.2555672
```

```r
aging_winterCens_CU <- coxph(
  # exclude isogen from model because only one strain
  Surv(Age.at.death, censrec) ~ sex + frailty(Birth),
  subset(aging_winterCens, arr=="CU")
)
summary(aging_winterCens_CU)$coefficients[1,]
```

```
##       coef   se(coef)        se2      Chisq         DF          p
## 0.37662682 0.21697590 0.21103195 3.01300269 1.00000000 0.08259915
```

```r
aging_winterCens_PP <- coxph(
  Surv(Age.at.death, censrec) ~ sex + isogen + frailty(Birth),
  subset(aging_winterCens, arr=="PP")
)
summary(aging_winterCens_PP)$coefficients[1,]
```

```
##       coef   se(coef)        se2      Chisq         DF          p
## 0.23355034 0.09914796 0.09875222 5.54872866 1.00000000 0.01849412
```

```r
aging_winterCens_ST <- coxph(
  Surv(Age.at.death, censrec) ~ sex + isogen + frailty(Birth),
  subset(aging_winterCens, arr=="ST")
)
summary(aging_winterCens_ST)$coefficients[1,]
```

```
##        coef    se(coef)         se2       Chisq          DF           p
## 0.324581799 0.104862778 0.103690024 9.580884501 1.000000000 0.001966137
```

```r
aging_winterCens_TL <- coxph(
  Surv(Age.at.death, censrec) ~ sex + isogen + frailty(Birth),
  subset(aging_winterCens, arr=="TL")
)
summary(aging_winterCens_TL)$coefficients[1,]
```

```
##       coef   se(coef)        se2      Chisq         DF          p
## 0.20768162 0.11878842 0.11726269 3.05666568 1.00000000 0.08040651
```

Next, we do this for the data excluding day 1 deaths.

```r
aging_winterCens_no1_AR <- coxph(
  Surv(Age.at.death, censrec) ~ sex + isogen + frailty(Birth),
  subset(aging_winterCens_no1, arr=="AR")
)
summary(aging_winterCens_no1_AR)$coefficients[1,]
```

```
##         coef     se(coef)          se2        Chisq           DF            p
## 5.213034e-01 9.919787e-02 9.761536e-02 2.761700e+01 1.000000e+00 1.478734e-07
```

```r
aging_winterCens_no1_CH <- coxph(
  Surv(Age.at.death, censrec) ~ sex + isogen + frailty(Birth),
  subset(aging_winterCens_no1, arr=="CH")
)
summary(aging_winterCens_no1_CH)$coefficients[1,]
```

```
##      coef  se(coef)       se2     Chisq        DF         p
## 0.1253223 0.1133944 0.1116236 1.2214445 1.0000000 0.2690773
```

```
aging_winterCens_no1_CU <- coxph(
  # exclude isogen from model because only one strain
  Surv(Age.at.death, censrec) ~ sex + frailty(Birth),
  subset(aging_winterCens_no1, arr=="CU")
)
summary(aging_winterCens_no1_CU)$coefficients[1,]
```

```
##       coef   se(coef)        se2      Chisq         DF          p
## 0.37662682 0.21697590 0.21103195 3.01300269 1.00000000 0.08259915
```

```
aging_winterCens_no1_PP <- coxph(
  Surv(Age.at.death, censrec) ~ sex + isogen + frailty(Birth),
  subset(aging_winterCens_no1, arr=="PP")
)
summary(aging_winterCens_no1_PP)$coefficients[1,]
```

```
##       coef   se(coef)        se2      Chisq         DF          p
## 0.24328624 0.09934659 0.09916879 5.99693260 1.00000000 0.01433077
```

```
aging_winterCens_no1_ST <- coxph(
  Surv(Age.at.death, censrec) ~ sex + isogen + frailty(Birth),
  subset(aging_winterCens_no1, arr=="ST")
)
summary(aging_winterCens_no1_ST)$coefficients[1,]
```

```
##        coef    se(coef)         se2       Chisq          DF           p
## 0.331448364 0.105416189 0.104247801 9.885921058 1.000000000 0.001665481
```

```
aging_winterCens_no1_TL <- coxph(
  Surv(Age.at.death, censrec) ~ sex + isogen + frailty(Birth),
  subset(aging_winterCens_no1, arr=="TL")
)
summary(aging_winterCens_no1_TL)$coefficients[1,]
```

```
##      coef  se(coef)       se2     Chisq        DF         p
## 0.1963443 0.1200906 0.1184100 2.6731195 1.0000000 0.1020558
```

##### Fit Gompertz models to data

The goal of this analysis is to test for sex differences in the rate and shape parameters of a Gompertz model. We will analyze data within each arrangement, comparing between males and females. Data from each arrangement will be analyzed to test for sex differences in the rate parameter first. To do that, we will define functions to perform the model comparisons.

```
# This function works for all arrangements, other than CU
flexsurv_model_comparisons <- function(agingDataArr){
    NULLmodel <- flexsurvreg(
      Surv(Age.at.death, censrec) ~ 1,
      data=agingDataArr,
      dist = "gompertz")

    batchEffect <- flexsurvreg(
      Surv(Age.at.death, censrec) ~ Birth,
      data=agingDataArr,
      dist = "gompertz")
```

```
    sexEffect <- flexsurvreg(
      Surv(Age.at.death, censrec) ~ sex,
      data=agingDataArr,
      dist = "gompertz")

    strainEffect <- flexsurvreg(
      Surv(Age.at.death, censrec) ~ isogen,
      data=agingDataArr,
      dist = "gompertz")

    sexbatch <- flexsurvreg(
      Surv(Age.at.death, censrec) ~ sex + Birth,
      data=agingDataArr,
      dist = "gompertz")

    sexstrain <- flexsurvreg(
      Surv(Age.at.death, censrec) ~ sex + isogen,
      data=agingDataArr,
      dist = "gompertz")

    strainbatch <- flexsurvreg(
      Surv(Age.at.death, censrec) ~ isogen + Birth,
      data=agingDataArr,
      dist = "gompertz")

    sexstrainbatch <- flexsurvreg(
      Surv(Age.at.death, censrec) ~ sex + isogen + Birth,
      data=agingDataArr,
      dist = "gompertz")

    return.df <- data.frame(
        factors = c("batch", "sex", "strain", "sexbatch", "sexstrain", "sexstrainbatch"),
        pval = c(
            pchisq(2 * (logLik(batchEffect) - logLik(NULLmodel)), df = 1, lower.tail = FALSE),
            pchisq(2 * (logLik(sexEffect) - logLik(NULLmodel)), df = 1, lower.tail = FALSE),
            pchisq(2 * (logLik(strainEffect) - logLik(NULLmodel)), df = 1, lower.tail = FALSE),
            pchisq(2 * (logLik(sexbatch) - logLik(batchEffect)), df = 1, lower.tail = FALSE),
            pchisq(2 * (logLik(sexstrain) - logLik(strainEffect)), df = 1, lower.tail = FALSE),
            pchisq(2 * (logLik(sexstrainbatch) - logLik(strainbatch)), df = 1, lower.tail = FALSE)
        ),
        significant = NA
    )

    return.df$significant <- ""
    return.df$significant[return.df$pval < 0.05] <- "*"
    return.df$significant[return.df$pval < 0.005] <- "**"
    return.df$significant[return.df$pval < 0.0005] <- "***"
    return.df$significant[return.df$pval < 0.00005] <- "****"
    return.df$significant[return.df$pval < 0.000005] <- "******"

    return.df
}
```

```r
# Need a special function for CU because only one strain represented
# so cannot test for isogen effects
flexsurv_model_comparisons_CU <- function(agingDataArr){
    NULLmodel <- flexsurvreg(
      Surv(Age.at.death, censrec) ~ 1,
      data=agingDataArr,
      dist = "gompertz")

    batchEffect <- flexsurvreg(
      Surv(Age.at.death, censrec) ~ Birth,
      data=agingDataArr,
      dist = "gompertz")

    sexEffect <- flexsurvreg(
      Surv(Age.at.death, censrec) ~ sex,
      data=agingDataArr,
      dist = "gompertz")

    sexbatch <- flexsurvreg(
      Surv(Age.at.death, censrec) ~ sex + Birth,
      data=agingDataArr,
      dist = "gompertz")

    return.df <- data.frame(
        factors = c("batch", "sex", "sexbatch"),
        pval = c(
            pchisq(2 * (logLik(batchEffect) - logLik(NULLmodel)), df = 1, lower.tail = FALSE),
            pchisq(2 * (logLik(sexEffect) - logLik(NULLmodel)), df = 1, lower.tail = FALSE),
            pchisq(2 * (logLik(sexbatch) - logLik(batchEffect)), df = 1, lower.tail = FALSE)
        ),
        significant = NA
    )

    return.df$significant <- ""
    return.df$significant[return.df$pval < 0.05] <- "*"
    return.df$significant[return.df$pval < 0.005] <- "**"
    return.df$significant[return.df$pval < 0.0005] <- "***"
    return.df$significant[return.df$pval < 0.00005] <- "****"
    return.df$significant[return.df$pval < 0.000005] <- "******"

    return.df
}

# Wrapper function to perform analysis on each arrangement
flexsurv_arrangement_sex <- function(agingData){
    rbind(
        cbind(  arr="AR",
                flexsurv_model_comparisons(subset(agingData, arr=="AR")) ),
        cbind(  arr="CH",
                flexsurv_model_comparisons(subset(agingData, arr=="CH")) ),
        cbind(  arr="PP",
                flexsurv_model_comparisons(subset(agingData, arr=="PP")) ),
        cbind(  arr="ST",
```

```
                flexsurv_model_comparisons(subset(agingData, arr=="ST")) ),
        cbind(  arr="TL",
                flexsurv_model_comparisons(subset(agingData, arr=="TL")) ),
        cbind(  arr="CU",
                flexsurv_model_comparisons_CU(subset(agingData, arr=="CU")) )
    )
}
```

Perform analysis on the full data using the functions defined above.

```
flexsurv_arrangement_sex_winterCens <- flexsurv_arrangement_sex(aging_winterCens)
flexsurv_arrangement_sex_winterCens
```

```
##    arr         factors         pval significant
## 1   AR           batch 2.866234e-10      ******
## 2   AR             sex 3.154854e-04         ***
## 3   AR          strain 1.575507e-18      ******
## 4   AR       sexbatch 8.412547e-04          **
## 5   AR       sexstrain 1.326771e-05        ****
## 6   AR sexstrainbatch 6.962350e-05         ***
## 7   CH           batch 1.865444e-07      ******
## 8   CH             sex 6.990905e-01
## 9   CH          strain 4.935314e-21      ******
## 10  CH        sexbatch 9.119765e-01
## 11  CH       sexstrain 3.365394e-01
## 12  CH sexstrainbatch 8.542932e-01
## 13  PP           batch 2.260172e-09      ******
## 14  PP             sex 5.098465e-02
## 15  PP          strain 2.295777e-08      ******
## 16  PP        sexbatch 2.015949e-02           *
## 17  PP       sexstrain 4.928504e-02           *
## 18  PP sexstrainbatch 2.254876e-02           *
## 19  ST           batch 4.342460e-16      ******
## 20  ST             sex 6.234575e-03           *
## 21  ST          strain 8.378194e-03           *
## 22  ST        sexbatch 2.360301e-03          **
## 23  ST       sexstrain 1.748630e-02           *
## 24  ST sexstrainbatch 1.181340e-02           *
## 25  TL           batch 8.196475e-07      ******
## 26  TL             sex 1.380316e-01
## 27  TL          strain 6.360254e-10      ******
## 28  TL        sexbatch 1.354221e-01
## 29  TL       sexstrain 6.514239e-02
## 30  TL sexstrainbatch 1.439627e-01
## 31  CU           batch 2.055161e-06      ******
## 32  CU             sex 7.145972e-02
## 33  CU        sexbatch 2.503730e-02           *
```

Perform analysis on data excluding 1 day old deaths.

```
flexsurv_arrangement_sex_winterCens_no1 <- flexsurv_arrangement_sex(aging_winterCens_no1)
flexsurv_arrangement_sex_winterCens_no1
```

```
##    arr         factors         pval significant
## 1   AR           batch 3.114380e-10      ******
## 2   AR             sex 2.003111e-04         ***
```

```
## 3   AR          strain 1.131268e-18      ******
## 4   AR        sexbatch 5.486524e-04          **
## 5   AR       sexstrain 8.516812e-06        ****
## 6   AR sexstrainbatch 4.702207e-05        ****
## 7   CH           batch 4.291549e-07      ******
## 8   CH             sex 7.420652e-01
## 9   CH          strain 5.325048e-21      ******
## 10  CH        sexbatch 9.788509e-01
## 11  CH       sexstrain 3.529826e-01
## 12  CH sexstrainbatch 8.957762e-01
## 13  PP           batch 2.904316e-09      ******
## 14  PP             sex 3.704927e-02           *
## 15  PP          strain 1.666903e-08      ******
## 16  PP        sexbatch 1.291704e-02           *
## 17  PP       sexstrain 3.534489e-02           *
## 18  PP sexstrainbatch 1.479638e-02           *
## 19  ST           batch 6.762156e-16      ******
## 20  ST             sex 5.012252e-03           *
## 21  ST          strain 5.659597e-03           *
## 22  ST        sexbatch 1.992222e-03          **
## 23  ST       sexstrain 1.497134e-02           *
## 24  ST sexstrainbatch 1.128865e-02           *
## 25  TL           batch 1.366797e-06      ******
## 26  TL             sex 1.607201e-01
## 27  TL          strain 4.690517e-10      ******
## 28  TL        sexbatch 1.609482e-01
## 29  TL       sexstrain 7.634847e-02
## 30  TL sexstrainbatch 1.750466e-01
## 31  CU           batch 2.055161e-06      ******
## 32  CU             sex 7.145972e-02
## 33  CU        sexbatch 2.503730e-02           *
```

There is a significant effect of sex on the rate parameter consistently observed for AR and ST, but not for other arrangements.

Next, compare models with different shape parameters between males and females. Include Birth and isogen in all models as rate parameters. First, define two functions to perform the analyses, excluding the CU arrangement because there is only one strain sampled with that genotype.

```
shape_model_comparisons <- function(agingDataArr){
    NULLmodel <- flexsurvreg(
      Surv(Age.at.death, censrec) ~ Birth + isogen,
      anc = list(shape = ~ 1),
      data = agingDataArr,
      dist = "gompertz")

    sexShape <- flexsurvreg(
      Surv(Age.at.death, censrec) ~ Birth + isogen,
      anc = list(shape = ~ sex),
      data = agingDataArr,
      dist = "gompertz")

    sexRate <- flexsurvreg(
      Surv(Age.at.death, censrec) ~ Birth + isogen + sex,
      anc = list(shape = ~ 1),
```

```
      data = agingDataArr,
      dist = "gompertz")

    sexShapeRate <- flexsurvreg(
      Surv(Age.at.death, censrec) ~ Birth + isogen + sex,
      anc = list(shape = ~ sex),
      data = agingDataArr,
      dist = "gompertz")

    return.df <- data.frame(
        factors = c("sexShapeNULL", "sexRateNULL", "sexRateShape", "sexShapeRate"),
        pval = c(
            # test if shape parameter improves fit
          pchisq(2 * (logLik(sexShape) - logLik(NULLmodel)), df = 1, lower.tail = FALSE),
          # test if rate parameter improves fit
            pchisq(2 * (logLik(sexRate) - logLik(NULLmodel)), df = 1, lower.tail = FALSE),
            # test if adding Rate to a model with Shape improves fit
            pchisq(2 * (logLik(sexShapeRate) - logLik(sexShape)), df = 1, lower.tail = FALSE),
            # test if adding Shape to a model with Rate improves fit
            pchisq(2 * (logLik(sexShapeRate) - logLik(sexRate)), df = 1, lower.tail = FALSE)
        ),
        significant = NA
    )

    return.df$significant <- ""
    return.df$significant[return.df$pval < 0.05] <- "*"
    return.df$significant[return.df$pval < 0.005] <- "**"
    return.df$significant[return.df$pval < 0.0005] <- "***"
    return.df$significant[return.df$pval < 0.00005] <- "****"
    return.df$significant[return.df$pval < 0.000005] <- "******"

    return.df
}
shape_arrangement_sex <- function(agingData){
    rbind(
        cbind(  arr="AR",
                shape_model_comparisons(subset(agingData, arr=="AR")) ),
        cbind(  arr="CH",
                shape_model_comparisons(subset(agingData, arr=="CH")) ),
        cbind(  arr="PP",
                shape_model_comparisons(subset(agingData, arr=="PP")) ),
        cbind(  arr="ST",
                shape_model_comparisons(subset(agingData, arr=="ST")) ),
        cbind(  arr="TL",
                shape_model_comparisons(subset(agingData, arr=="TL")) )
    )
}
```

Perform the analysis comparing models with an without the shape parameter on the full data set.

```
shape_arrangement_sex_winterCens <- shape_arrangement_sex(aging_winterCens)
shape_arrangement_sex_winterCens
```

```
##    arr      factors         pval significant
## 1   AR sexShapeNULL 0.0360581540           *
```

```
## 2   AR  sexRateNULL 0.0000696235         ***
## 3   AR sexRateShape 0.0005504437          **
## 4   AR sexShapeRate 0.4752194883
## 5   CH sexShapeNULL 0.4630631347
## 6   CH  sexRateNULL 0.8542932131
## 7   CH sexRateShape 0.3147986087
## 8   CH sexShapeRate 0.2183487245
## 9   PP sexShapeNULL 0.0081046210           *
## 10  PP  sexRateNULL 0.0225487572           *
## 11  PP sexRateShape 0.6356155032
## 12  PP sexShapeRate 0.1540386368
## 13  ST sexShapeNULL 0.5537828304
## 14  ST  sexRateNULL 0.0118134001           *
## 15  ST sexRateShape 0.0038653660          **
## 16  ST sexShapeRate 0.1246478107
## 17  TL sexShapeNULL 0.0363266201           *
## 18  TL  sexRateNULL 0.1439626506
## 19  TL sexRateShape 0.7214651885
## 20  TL sexShapeRate 0.1233930159
```

Perform the analysis on the data excluding the one day old deaths.

```
shape_arrangement_sex_winterCens_no1 <- shape_arrangement_sex(aging_winterCens_no1)
shape_arrangement_sex_winterCens_no1
```

```
##    arr      factors         pval significant
## 1   AR sexShapeNULL 3.970394e-02           *
## 2   AR  sexRateNULL 4.702207e-05        ****
## 3   AR sexRateShape 3.024475e-04         ***
## 4   AR sexShapeRate 3.957744e-01
## 5   CH sexShapeNULL 4.453177e-01
## 6   CH  sexRateNULL 8.957762e-01
## 7   CH sexRateShape 3.351171e-01
## 8   CH sexShapeRate 2.215345e-01
## 9   PP sexShapeNULL 7.196495e-03           *
## 10  PP  sexRateNULL 1.479638e-02           *
## 11  PP sexRateShape 5.090886e-01
## 12  PP sexShapeRate 1.898674e-01
## 13  ST sexShapeNULL 5.609927e-01
## 14  ST  sexRateNULL 1.128865e-02           *
## 15  ST sexRateShape 3.476156e-03          **
## 16  ST sexShapeRate 1.169506e-01
## 17  TL sexShapeNULL 3.679887e-02           *
## 18  TL  sexRateNULL 1.750466e-01
## 19  TL sexRateShape 5.934908e-01
## 20  TL sexShapeRate 9.394522e-02
```

Adding a shape effect of sex to the model (sexShapeRate) does not improve model fit. Therefore, there is an affect of sex on the rate parameter in some arrangements, but no effect of sex on the shape.

Lastly, we will compare arrangements within each sex. To do so, we will compare the fit of models with arrangement as a rate and/or shape parameter. We will define two functions to perform the analysis on data from females and males separately. Strain (isogen) is not included in these models because flexsurvreg cannot handle isogen as a nested or interacting factor in a model with arrangement as a predictor.

```
arrangement_comparisons <- function(agingDataArr){
    # start with model that has batch
  batchRate <- flexsurvreg(
    Surv(Age.at.death, censrec) ~ Birth,
    anc = list(shape = ~ 1),
    data=agingDataArr,
    dist = "gompertz")

  # add arrangement as rate parameter
  arrRate <- flexsurvreg(
    Surv(Age.at.death, censrec) ~ Birth + arr,
    anc = list(shape = ~ 1),
    data=agingDataArr,
    dist = "gompertz")

  # add arrangement as shape parameter
    arrShape <- flexsurvreg(
      Surv(Age.at.death, censrec) ~ Birth,
      anc = list(shape = ~ arr),
      data=agingDataArr,
      dist = "gompertz")

    # add arrangement as both rate and shape parameter
    arrRateShape <- flexsurvreg(
      Surv(Age.at.death, censrec) ~ Birth + arr,
      anc = list(shape = ~ arr),
      data=agingDataArr,
      dist = "gompertz")

    return.df <- data.frame(
        factors = c("arrRate", "arrShape", "arrRateShape", "arrShapeRate"),
        pval = c(
            # test for effect of arrangement on rate
          pchisq(2 * (logLik(arrRate) - logLik(batchRate)), df = 2, lower.tail = FALSE),
          # test for effect of arrangement on shape
            pchisq(2 * (logLik(arrShape) - logLik(batchRate)), df = 2, lower.tail = FALSE),
            # test for effect of arrangement on rate if shape is included in the model
            pchisq(2 * (logLik(arrRateShape) - logLik(arrShape)), df = 2, lower.tail = FALSE),
            # test for the effect of arrangement on shape if rate is included in the model
            pchisq(2 * (logLik(arrRateShape) - logLik(arrRate)), df = 1, lower.tail = FALSE)
        ),
        significant = NA
    )

    return.df$significant <- ""
    return.df$significant[return.df$pval < 0.05] <- "*"
    return.df$significant[return.df$pval < 0.005] <- "**"
    return.df$significant[return.df$pval < 0.0005] <- "***"
    return.df$significant[return.df$pval < 0.00005] <- "****"
    return.df$significant[return.df$pval < 0.000005] <- "******"

    return.df
}
```

```
# Wrapper function to analyze data from each sex
arr_models <- function(agingData){
    rbind(
        cbind(  sex="female",
                arrangement_comparisons(subset(agingData, sex=="female")) ),
        cbind(  sex="male",
                arrangement_comparisons(subset(agingData, sex=="male")) )
    )
}
```

```
arr_models_winterCens <- arr_models(aging_winterCens)
arr_models_winterCens
```

```
##      sex      factors         pval significant
## 1 female      arrRate 2.241487e-09      ******
## 2 female     arrShape 1.038131e-06      ******
## 3 female arrRateShape 1.142683e-06      ******
## 4 female arrShapeRate 1.026033e-04         ***
## 5   male      arrRate 3.728415e-10      ******
## 6   male     arrShape 9.283270e-05         ***
## 7   male arrRateShape 8.217931e-13      ******
## 8   male arrShapeRate 2.854101e-08      ******
```

```
arr_models_winterCens_no1 <- arr_models(aging_winterCens_no1)
arr_models_winterCens_no1
```

```
##      sex      factors         pval significant
## 1 female      arrRate 3.278252e-09      ******
## 2 female     arrShape 1.272528e-06      ******
## 3 female arrRateShape 2.482842e-06      ******
## 4 female arrShapeRate 1.938979e-04         ***
## 5   male      arrRate 1.809467e-10      ******
## 6   male     arrShape 9.040359e-05         ***
## 7   male arrRateShape 3.329968e-13      ******
## 8   male arrShapeRate 2.306055e-08      ******
```

Arrangement affects both the shape and rate parameters in both males and females.

We can visualize the effects by plotting the age-dependent mortality rate. The time dependent mortality rate in Gompertz model is represented as $\mu(t) = ae^{bt}$, where $a$ is the initial mortality rate (rate parameter) and $b$ is the rate of aging (shape paramter). Calling `flexsurvreg()` gives us the model coefficients if we run it on each subset of data (i.e., each sex-by-arrangement combination). We will do this by writing a function to perform the analysis on each arrangement.

```
gompertz.model.extract <- function(aging_sex){
  Arrs <- levels(factor(aging_sex$arr))
  return.df <- data.frame(
    arr = Arrs,
    shape = NA,
    rate = NA
  )
  for(i in 1:length(Arrs)){
    return.df[i, 2:3] <- flexsurvreg(
      Surv(Age.at.death, censrec) ~ Birth,
        data = subset(aging_sex, arr==Arrs[i]),
        dist = "gompertz")$res[1:2,1]
```

```
  }
  return.df
}

gompertz_model_fit <- rbind(
  cbind(
    gompertz.model.extract(subset(aging_winterCens, sex=="female")),
    sex="female"
  ),
  cbind(
    gompertz.model.extract(subset(aging_winterCens, sex=="male")),
    sex="male"
  )
)
gompertz_model_fit
```

```
##    arr       shape        rate    sex
## 1   AR 0.009758308 0.020121771 female
## 2   CH 0.015231649 0.040453447 female
## 3   CU 0.038311328 0.010444613 female
## 4   PP 0.004679225 0.027280258 female
## 5   ST 0.017554860 0.033192226 female
## 6   TL 0.017904007 0.008941951 female
## 7   AR 0.004125545 0.019314911   male
## 8   CH 0.005592349 0.030685167   male
## 9   CU 0.087803202 0.024363090   male
## 10  PP 0.017967081 0.022516295   male
## 11  ST 0.011172240 0.066079637   male
## 12  TL 0.027454879 0.009679397   male
```

Plot the mortality rate over time. To do so, write a function to plot the natural log of the gompertz mortality rate, and another function to make the graphs.

```
gompertz.mortality <- function(x, ShapeRate){
  log(ShapeRate[2] * exp(ShapeRate[1] * x))
}

gompertz.mortality.plot <- function(GMF){
  ggplot() + xlim(0, 50) + ylim(-5,0.5) +
    scale_color_manual(name="",
      values = c(
          "AR" = "blue",
          "CU" = "black",
          "ST"="darkgreen",
          "CH"="orange",
          "PP"="red",
          "TL"="purple")) +
  geom_function(aes(color=GMF[1,1]), fun=gompertz.mortality,
                args = list( unlist(GMF[1,2:3]))) +
  geom_function(aes(color=GMF[2,1]), fun=gompertz.mortality,
                args = list( unlist(GMF[2,2:3]))) +
  geom_function(aes(color=GMF[3,1]), fun=gompertz.mortality,
                args = list( unlist(GMF[3,2:3]))) +
  geom_function(aes(color=GMF[4,1]), fun=gompertz.mortality,
                args = list( unlist(GMF[4,2:3]))) +
```

```
  geom_function(aes(color=GMF[5,1]), fun=gompertz.mortality,
                args = list( unlist(GMF[5,2:3]))) +
  geom_function(aes(color=GMF[6,1]), fun=gompertz.mortality,
                args = list( unlist(GMF[6,2:3]))) +
  xlab("time (days)") + ylab("ln(mortality rate)") +
  theme_bw()
}
```

Plot first for females.

```
gompertz.mortality.plot(subset(gompertz_model_fit, sex=="female"))
```

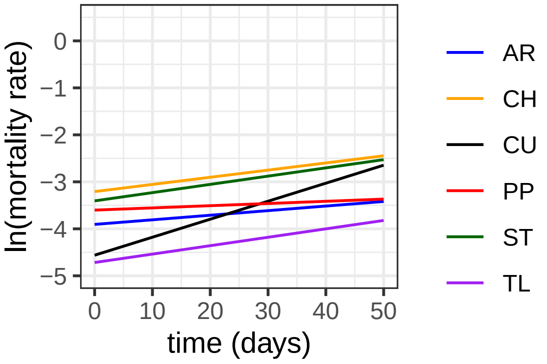


Plot next for males.

```
gompertz.mortality.plot(subset(gompertz_model_fit, sex=="male"))

## Warning: Removed 4 rows containing missing values or values outside the scale range
## (`geom_function()`).
```

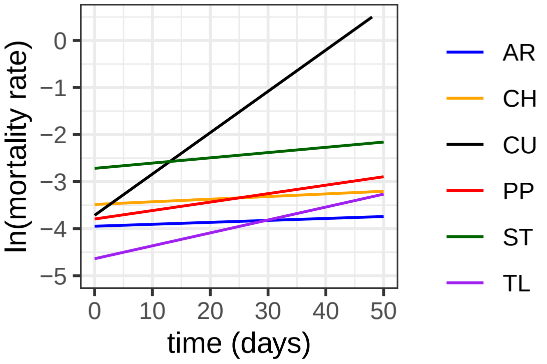


##### Export lifespan estimates for downstream analyses

We will use estimates of lifespan from this experiment in other analyses. To do so, we export estimates of lifespan for each strain. First, we will use the median value for each strain.

```
sex.median <- function(aging){
    strains <- levels(factor(aging$Strain))
    female.df <- data.frame(
        strain = strains,
        median = NA,
        obs = NA
    )
    male.df <- female.df
```

```
    for(i in 1:length(strains)){
        female.df$median[i] <- median(
          subset(aging, Strain==strains[i] & sex=="female")$Age.at.death
        )
        female.df$obs[i] <- length(
          subset(aging, Strain==strains[i] & sex=="female")[,1]
        )
    }
    female.df$sex <- "female"
    for(i in 1:length(strains)){
        male.df$median[i] <- median(subset(aging, Strain==strains[i] & sex=="male")$Age.at.death)
        male.df$obs[i] <- length(subset(aging, Strain==strains[i] & sex=="male")[,1])
    }
    male.df$sex <- "male"
    rbind(female.df, male.df)
}
sex_median_winterCens <- sex.median(aging_winterCens)
sex_median_winterCens_no1 <- sex.median(aging_winterCens_no1)

write.table(
  sex_median_winterCens, "sex_median_winterCens.tsv",
  quote=FALSE, sep="\t", row.names=FALSE)

write.table(
  sex_median_winterCens_no1, "sex_median_winterCens_no1.tsv",
  quote=FALSE, sep="\t", row.names=FALSE)
```

Second, we will use the intercept from a linear model for each strain. This allows us to calculate the mean value, while also modeling batch (emergence date) for each fly as a random effect. We will write a function to calculate the intercept, only including strains with measurements from at least 10 flies.

```
sex.intercept <- function(datas){
    strains <- levels(factor(datas$Strain))
    female.df <- data.frame(
        strain = strains,
        effect = NA,
        obs = NA
    )
    male.df <- female.df

    for(i in 1:length(strains)){
        # only consider if have >9 samples
      if(length(subset(datas, Strain==strains[i] & sex=="female")[,1]) > 9){
            if(length(levels(factor(
                subset(datas, sex=="female" & Strain==strains[i])$Birth))) > 1){
                female.df$effect[i] <- summary(lmer(Age.at.death ~ 1 + (1|Birth),
                                                    data=subset(datas,
                                                    Strain==strains[i] & sex=="female"))
                                                )$coefficients[1]
            }else{
                female.df$effect[i] <- summary(lm(Age.at.death ~ 1,
                                                    data=subset(datas,
                                                    Strain==strains[i] & sex=="female"))
                                                )$coefficients[1]
```

```
            }
        } else{
            female.df$effect[i] <- NA
        }
        female.df$obs[i] <- length(subset(datas,
                                          Strain==strains[i] & sex=="female")[,1])
    }
    female.df$sex <- "female"

    for(i in 1:length(strains)){
        # only consider if have >9 samples
      if(length(subset(datas, Strain==strains[i] & sex=="male")[,1]) > 9){
            if(length(levels(factor(
                subset(datas, sex=="male" & Strain==strains[i])$Birth))) > 1){
                male.df$effect[i] <- summary(lmer(Age.at.death ~ 1 + (1|Birth),
                                                  data=subset(datas,
                                                    Strain==strains[i] & sex=="male"))
                                              )$coefficients[1]
            } else{
                male.df$effect[i] <- summary(lm(Age.at.death ~ 1,
                                                 data=subset(datas,
                                                   Strain==strains[i] & sex=="male"))
                                              )$coefficients[1]
            }
        } else{
            male.df$effect[i] <- NA
        }
        male.df$obs[i] <- length(subset(datas, Strain==strains[i] & sex=="male")[,1])
    }
    male.df$sex <- "male"

    rbind(female.df, male.df)
}

sex_intercept_winterCens <- sex.intercept(aging_winterCens)
sex_intercept_winterCens_no1 <- sex.intercept(aging_winterCens_no1)

write.table(sex_intercept_winterCens, "sex_intercept_winterCens.tsv",
            quote=FALSE, sep="\t", row.names=FALSE)
write.table(sex_intercept_winterCens_no1, "sex_intercept_winterCens_no1.tsv",
            quote=FALSE, sep="\t", row.names=FALSE)
```

#### Make plots of the data and survival models

The analysis above demonstrated very similar results when we exclude 1 day old death and when we include all data. Because of these similarities, we will include all data in our plots.

We will first use autoplot to graph survivorship (Kaplan-Meier) curves. The graphing functions will ignore any variable other than the focal effects in order to produce clear graphs. Start with a plot comparing males and females.

```
autoplot(survfit(Surv(Age.at.death, censrec) ~ sex,
                 data=aging_winterCens),
                 ylab = "% surivival", xlab="time (days)") +
```

```
    scale_color_manual(values = c("female" = "red", "male" = "blue")) +
    annotate("text", x=2, y=0.40, label=paste("males"), hjust=0, color="blue") +
    annotate("text", x=23, y=0.50, label=paste("females"), hjust=0, color="red") +
    theme_bw() +
    theme(legend.position="none")
```

```
## Warning: `aes_string()` was deprecated in ggplot2 3.0.0.
## i Please use tidy evaluation idioms with `aes()`.
## i See also `vignette("ggplot2-in-packages")` for more information.
## i The deprecated feature was likely used in the ggfortify package.
##   Please report the issue at <https://github.com/sinhrks/ggfortify/issues>.
## This warning is displayed once every 8 hours.
## Call `lifecycle::last_lifecycle_warnings()` to see where this warning was
## generated.
```

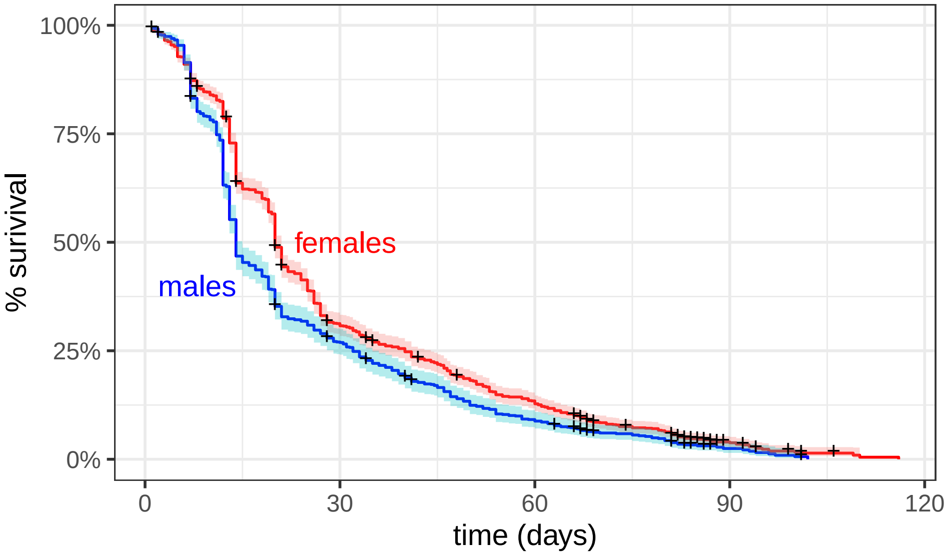


Create separate survivorship curves for males and females with each arrangement.

```
ggsurvplot_facet(survfit(
    Surv(Age.at.death, censrec) ~ sex + arr,
    data=aging_winterCens),
    aging_winterCens,
    facet.by = "arr",
    ncol=2,
    short.panel.labs=TRUE,
    xlab="time (days)",
    legend.title="") +
    scale_color_manual(values = c("female" = "magenta", "male" = "cornflowerblue"))
```

```
## Warning: Using `size` aesthetic for lines was deprecated in ggplot2 3.4.0.
## i Please use `linewidth` instead.
## i The deprecated feature was likely used in the ggpubr package.
##   Please report the issue at <https://github.com/kassambara/ggpubr/issues>.
## This warning is displayed once every 8 hours.
## Call `lifecycle::last_lifecycle_warnings()` to see where this warning was
## generated.
```

```
## Warning: ggtheme is not a valid theme.
```

```
## Please use `theme()` to construct themes.

## Ignoring unknown labels:
## * fill : ""
## * linetype : "1"
```

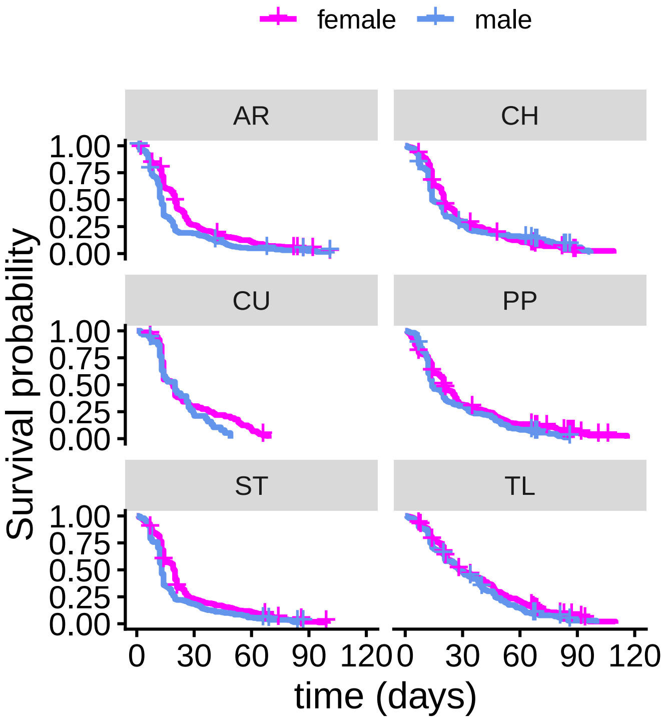


Create survival curves for each arrangement for males and females.

```
ggsurvplot_facet(survfit(Surv(Age.at.death, censrec) ~ sex + arr,
                         data=aging_winterCens), aging_winterCens,
    facet.by = "sex", ncol=1, short.panel.labs=TRUE, xlab="time (days)", legend.title="") +
    scale_color_manual(values = c(
      "AR" = "blue",
      "CU" = "black",
      "ST"="darkgreen",
      "CH"="orange",
      "PP"="red",
      "TL"="purple"))
```

```
## Warning: ggtheme is not a valid theme.
## Please use `theme()` to construct themes.

## Ignoring unknown labels:
## * fill : ""
## * linetype : "1"
```

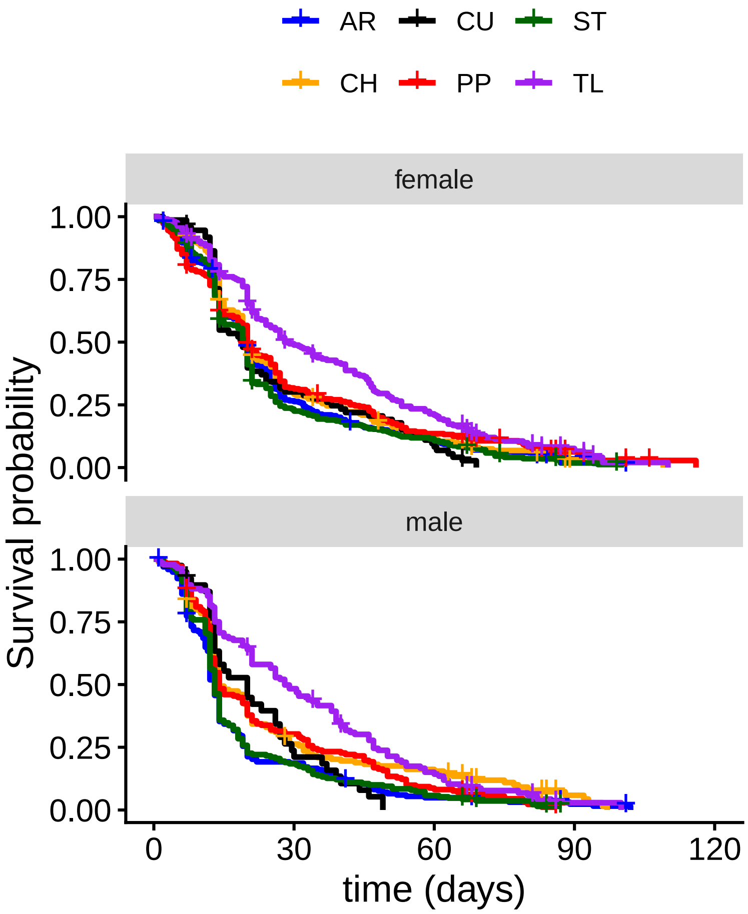


Use Gompertz model to calculate survival for males and females with each arrangement. This code chunk first defines a function to do the graphing, and then it calls that function to plot the model.

```
gompertz.arr.plot <- function(agingData){
    arrangements <- levels(factor(agingData$arr))
    colors <- c( "blue", "orange", "black", "red", "darkgreen", "purple")

        par(mfcol = c(6,2), mar = c(2, 4, 1, 0.5)) # mar = c(bottom, left, top, right)
        for(i in 1:length(arrangements)){
            if(i==1){ # print column heading if first graph
                plot(
                    flexsurvreg(Surv(Age.at.death, censrec) ~ 1,
                                data=subset(agingData, sex=="female" & arr==arrangements[i]),
                                dist = "gompertz"),
                     main="females", las=1, col=colors[i], xlim=c(0,100),
                     xlab="", ylab="Survival Probability"
                )
                text(x=90, y=0.9, labels=arrangements[i], col=colors[i])
            }else if(i==length(arrangements)){ # print X axis title if last graph
                plot(
                    flexsurvreg(Surv(Age.at.death, censrec) ~ 1,
                                data=subset(agingData, sex=="female" & arr==arrangements[i]),
                                dist = "gompertz"),
                    main="", las=1, col=colors[i], xlim=c(0,100),
                    xlab="time (days)", ylab="Survival Probability"
```

```
            )
            text(x=90, y=0.9, labels=arrangements[i], col=colors[i])
        }else{
            plot(
                flexsurvreg(Surv(Age.at.death, censrec) ~ 1,
                            data=subset(agingData, sex=="female" & arr==arrangements[i]),
                            dist = "gompertz"),
                 main="", las=1, col=colors[i], xlim=c(0,100),
                xlab="", ylab="Survival Probability"
            )
            text(x=90, y=0.9, labels=arrangements[i], col=colors[i])
        }
    }
    for(i in 1:length(arrangements)){
        if(i==1){ # print column heading if first graph
            plot(
                flexsurvreg(Surv(Age.at.death, censrec) ~ 1,
                            data=subset(agingData, sex=="male" & arr==arrangements[i]),
                            dist = "gompertz"),
                main="males", las=1, col=colors[i], xlim=c(0,100),
                xlab="", ylab=""
            )
            text(x=90, y=0.9, labels=arrangements[i], col=colors[i])
        }else if(i==length(arrangements)){ # print X axis title if last graph
            plot(
                flexsurvreg(Surv(Age.at.death, censrec) ~ 1,
                            data=subset(agingData, sex=="male" & arr==arrangements[i]),
                            dist = "gompertz"),
                main="", las=1, col=colors[i], xlim=c(0,100),
                xlab="time (days)", ylab=""
            )
            text(x=90, y=0.9, labels=arrangements[i], col=colors[i])
        }else{
            plot(
                flexsurvreg(Surv(Age.at.death, censrec) ~ 1,
                            data=subset(agingData, sex=="male" & arr==arrangements[i]),
                            dist = "gompertz"),
                main="", las=1, col=colors[i], xlim=c(0,100),
                xlab="", ylab=""
            )
            text(x=90, y=0.9, labels=arrangements[i], col=colors[i])
        }
    }
}

gompertz.arr.plot(aging_winterCens)
```

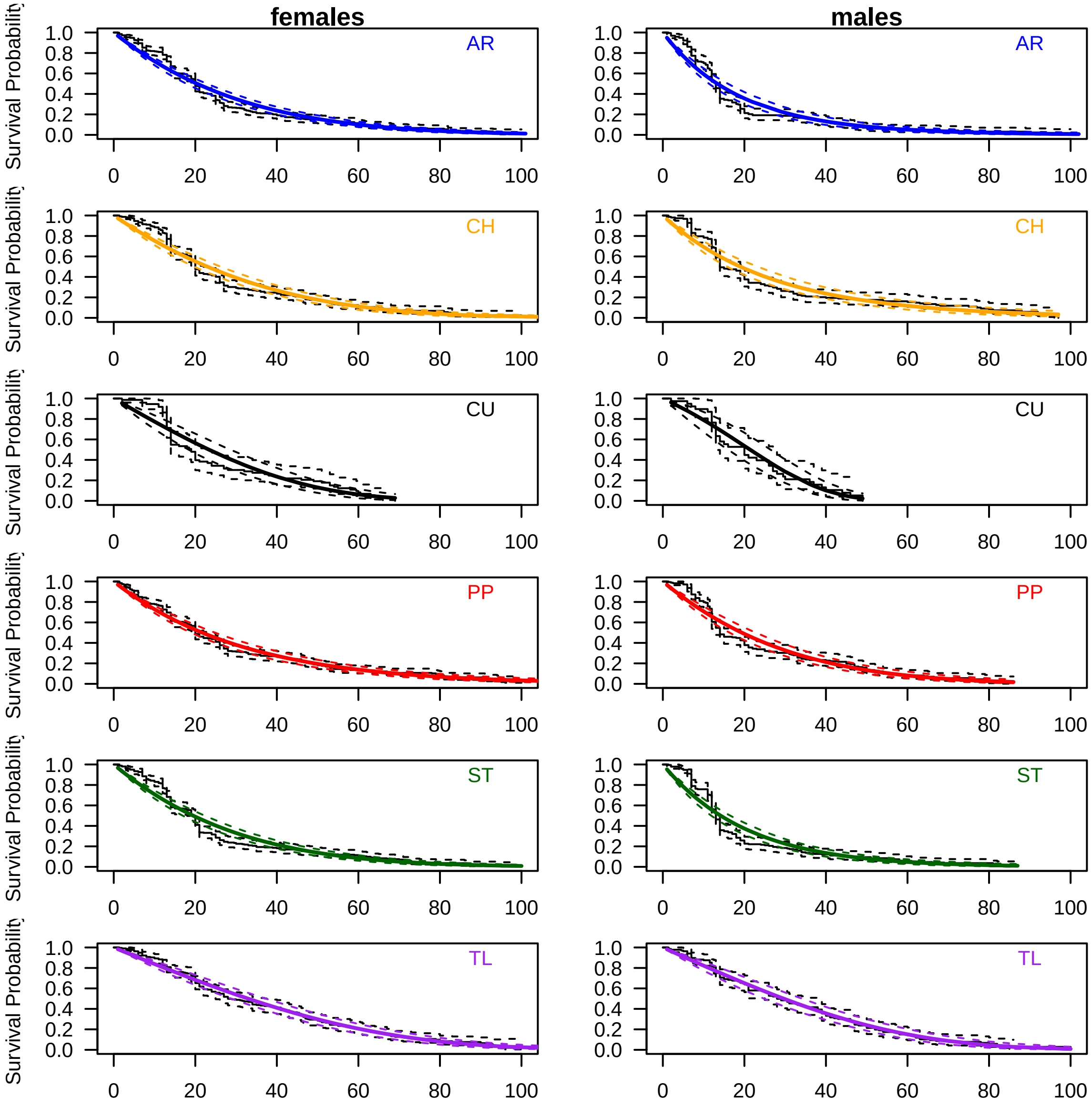

females
males
AR
CH
CU
PP
ST
TL
Survival Probability
1.0
0.8
0.6
0.4
0.2
0.0
0
20
40
60
80
100
