## Supplemental File S2 for "*Drosophila pseudoobscura* third chromosome inversion arrangements have temperature-dependent and sex-specific effects on life history traits"

### Supplementary File S2: Analyze lifespan data from *Drosophila pseudoobscura* at 18C, 22C, and 25C

2026-09-03

#### Overview

The analysis described below was performed on survival data from *Drosophila pseudoobscura* collected in 2025. Males and females carrying one of six different third chromosome inversions arrangements were sampled. The flies were sampled from 28 different strains, each of which is homozygous for one of six different chromosomal inversion arrangements. However, filtering of the data reduced the total number of strains analyzed to 27 (see below). Flies were collected and raised at either 18°C, 22°C, or 25°C.

Individual flies were collected upon emergence and stored in vials with standard *Drosophila* medium until they died. Each fly was transferred to a new vial every ~7 days until it died. The date of emergence from the pupal case (birth) and death were recorded for each fly. These dates were used to calculate the age at death (in days since emergence from the pupal case). Observations were predominately made on weekdays. If a fly died over an unobserved weekend or holiday, the midpoint of the unobserved period was recorded as the age at death. Flies that escaped were marked as censored data.

#### Install and Load Required Packages

```r
if (!require("tidyr")) install.packages("tidyr")
if (!require("ggplot2")) install.packages("ggplot2")
if (!require("ggfortify")) install.packages("ggfortify")
if (!require("survival")) install.packages("survival")
if (!require("survminer")) install.packages("survminer")
if (!require("flexsurv")) install.packages("flexsurv")
if (!require("lme4")) install.packages("lme4")
if (!require("coxme")) install.packages("coxme")
if (!require("tidychangepoint")) install.packages("tidychangepoint")
if (!require("cowplot")) install.packages("cowplot")

library(tidyr)
library(ggplot2)
library(ggfortify)
library(survival)
library(survminer)
library(flexsurv)
library(lme4)
library(coxme)
library(tidychangepoint) # to extract degrees of freedom from logLik object
library(cowplot)
```

#### Load, Prepare, and QC the Data

Load the survival data from file.

```r
aging_18 <- read.delim("Dpse_aging_18C_2025_08_11.txt")
aging_22 <- read.delim("Dpse_aging_22C_2025_07_01.txt")
aging_25 <- read.delim("Dpse_aging_25C_2025_07_01.txt")
```

Each row in the aging dataframes is a single fly, and the columns contain information about the fly. The columns are strain ID, sex of the fly, birth date, death date, age at death, any notes (related to censoring), if they fly is censored (0) or not (1).

First, confirm that all dates are formatted the same. To do so, check if any birth dates are singletons, which are probably errors that need to be corrected Construct a dataframe with the number of birth dates, and count the number of observations per birth date for each temperature.

```r
births_18 <- levels(factor(aging_18$Birth))
births_22 <- levels(factor(aging_22$Birth))
births_25 <- levels(factor(aging_25$Birth))

birthdates <- function(births, aging){
    Births.df <- data.frame(
        birth = births,
        count = NA
    )
    for(i in 1:length(births)){
        Births.df$count[i] <- length(subset(aging, Birth==births[i])[,1])
    }
    Births.df
}
Births_18.df <- birthdates(births_18, aging_18)
Births_22.df <- birthdates(births_22, aging_22)
Births_25.df <- birthdates(births_25, aging_25)
```

Next, make a plot of the birth dates per temperature. Birth dates will be treated as batches in the statistical analysis. Write a function to make the plot for each temperature.

```r
births.graph <- function(Births.df){
    Births.df %>% separate_wider_delim(
        birth,
        delim="-",
        names=c("day", "month", "year"),
        cols_remove = FALSE) -> Birth.graph
    Birth.graph$month[Birth.graph$month=="Feb"] <- "  Feb"
    Birth.graph$month[Birth.graph$month=="Mar"] <- " Mar"
    Birth.graph$month[Birth.graph$month=="Apr"] <- "Apr"

    Birth.graph$days <- as.numeric(Birth.graph$day)
    Birth.graph$day[Birth.graph$days < 10] <-
      paste(" ", Birth.graph$day[Birth.graph$days<10], sep="")
    Birth.graph$date <- paste(Birth.graph$month, Birth.graph$day)

    ggplot(Birth.graph, aes(x=date, y=count)) +
        geom_bar(stat="identity") +
        scale_x_discrete(guide = guide_axis(angle = 90), name=NULL) +
        scale_y_continuous(name="number of flies") +
        theme_bw()
}
```

Plot graphs for each temperature.

```r
births.graph(Births_18.df)
```

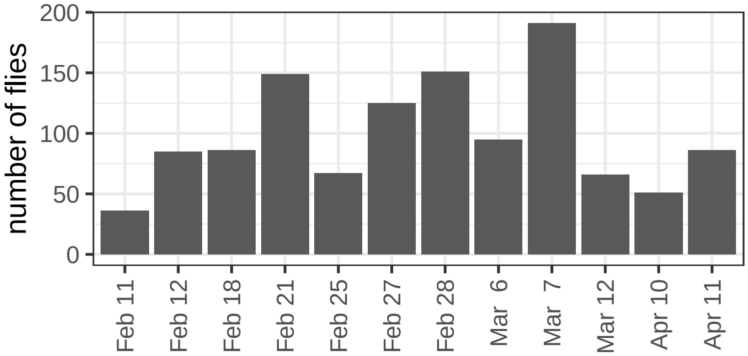


```r
births.graph(Births_22.df)
```

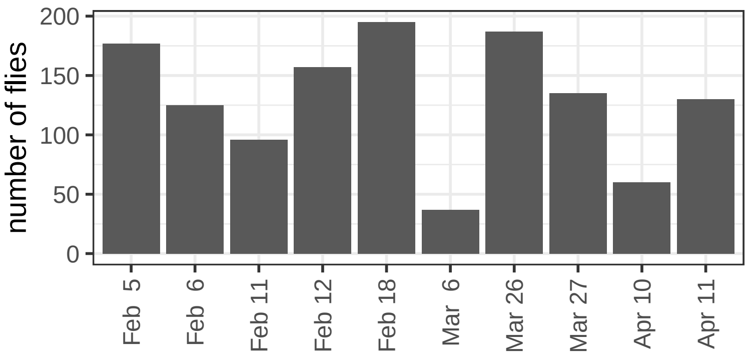


```r
births.graph(Births_25.df)
```

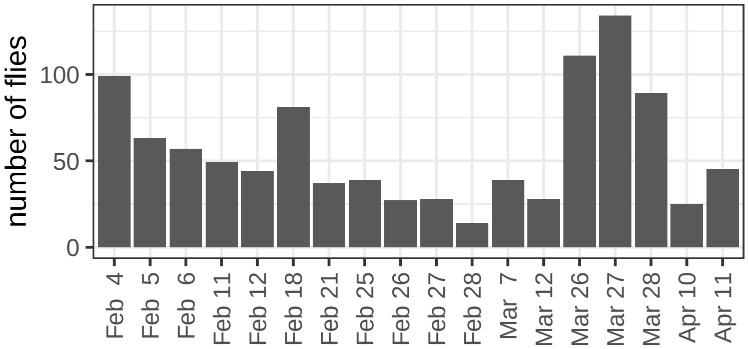


Assign strain information to each fly.

```r
strainIDs <- read.delim("Dpse-strains.tsv")

strain.assign <- function(strains, aging){
    aging$arr <- strains$arrangement[match(aging$Strain, strains$ID)]
    aging$isogen <- strains$isogen[match(aging$Strain, strains$ID)]
    aging$bal <- strains$balancer[match(aging$Strain, strains$ID)]

    # Omit data with non-existent strain ID
    aging <- subset(aging, !is.na(arr))
}
aging_18 <- strain.assign(strainIDs, aging_18)
aging_22 <- strain.assign(strainIDs, aging_22)
```

```
aging_25 <- strain.assign(strainIDs, aging_25)
```

Create a table with counts of strains assayed at each temperature in order to determine if any strains are under-represented in the data.

```
aging.strains <- function(aging, strains){
    aging_strains <- data.frame(
        strains = levels(factor(aging$Strain))
    )
    aging_strains$arr <- strains$arrangement[match(aging_strains$strains, strains$ID)]
    aging_strains$isogen <- strains$isogen[match(aging_strains$strains, strains$ID)]
    aging_strains$bal <- strains$balancer[match(aging_strains$strains, strains$ID)]
    for(i in 1:length(aging_strains$strains)){
        aging_strains$females[i] <- length(
          subset(aging, Strain==aging_strains$strains[i] & sex=="female")[,1]
          )
        aging_strains$males[i] <- length(
          subset(aging, Strain==aging_strains$strains[i] & sex=="male")[,1]
          )
    }
    aging_strains
}

aging_strains_18 <- aging.strains(aging_18, strainIDs)
aging_strains_22 <- aging.strains(aging_22, strainIDs)
aging_strains_25 <- aging.strains(aging_25, strainIDs)
```

```
aging_strains_18
```

```
##    strains arr     isogen bal females males
## 1        3  AR     DM1015   L       9    15
## 2        4  AR     DM1050   L      20    17
## 3        5  AR     DM1056   L      36    26
## 4        6  AR     DM1088   L       0     1
## 5        7  AR      KB635   L      27    12
## 6        8  AR      KB652   L      52    44
## 7       14  AR     MSH126   L      16    21
## 8       17  PP    BdA1137  10      54    33
## 9       18  PP     DM1020   B      21    10
## 10      19  PP     DM1038   B      25    15
## 11      20  PP     DM1049   B       9     7
## 12      22  PP     DM1065   L       7    10
## 13      24  PP     DM1085   B      40    18
## 14      26  ST      JR138   L      21    25
## 15      28  ST      JR209   L      34    20
## 16      31  ST       JR91   L      19    17
## 17      33  ST     MSH217   B      27    21
## 18      34  CH       JR32   B      14    16
## 19      35  CH        JR4   L      16    27
## 20      36  CH       JR20   L      31    26
## 21      38  CH      JR272   B      17    19
## 22      42  CH     MSH202   B      27    21
## 23      43  TL     MA1959          20    23
## 24      46  TL    SCI12-2   B      24    23
## 25      47  TL SPE123_2-3   B      18    15
```

```
## 26      48  TL SPE123_5-1   B      26    14
## 27      49  TL SPE123_6-3   B      24    12
## 28      53  CU SPE123_4-1   B      18    26
```

```
aging_strains_22
```

```
##    strains arr     isogen bal females males
## 1        3  AR     DM1015   L      29    21
## 2        4  AR     DM1050   L      21    18
## 3        5  AR     DM1056   L      40    35
## 4        7  AR      KB635   L      15     7
## 5        8  AR      KB652   L      39    23
## 6       14  AR     MSH126   L      37    21
## 7       17  PP    BdA1137  10      40    30
## 8       18  PP     DM1020   B      16    14
## 9       19  PP     DM1038   B      29    18
## 10      20  PP     DM1049   B      10     9
## 11      22  PP     DM1065   L       7     7
## 12      24  PP     DM1085   B      37    31
## 13      26  ST      JR138   L      38    21
## 14      28  ST      JR209   L      38    32
## 15      31  ST       JR91   L      17     9
## 16      33  ST     MSH217   B      49    28
## 17      34  CH       JR32   B      28     8
## 18      35  CH        JR4   L      16    12
## 19      36  CH       JR20   L      15    12
## 20      38  CH      JR272   B      26    22
## 21      42  CH     MSH202   B      37    43
## 22      43  TL     MA1959          23    31
## 23      46  TL    SCI12-2   B      28    18
## 24      47  TL SPE123_2-3   B      23    16
## 25      48  TL SPE123_5-1   B      19    20
## 26      49  TL SPE123_6-3   B      24     9
## 27      53  CU SPE123_4-1   B      46    34
```

```
aging_strains_25
```

```
##    strains arr     isogen bal females males
## 1        3  AR     DM1015   L      41    36
## 2        4  AR     DM1050   L      10     3
## 3        7  AR      KB635   L      11    12
## 4        8  AR      KB652   L      47    39
## 5       14  AR     MSH126   L      24    20
## 6       17  PP    BdA1137  10      50    84
## 7       18  PP     DM1020   B      12     6
## 8       19  PP     DM1038   B      33    14
## 9       20  PP     DM1049   B      17    11
## 10      22  PP     DM1065   L       6     2
## 11      24  PP     DM1085   B      16    11
## 12      26  ST      JR138   L      13    14
## 13      28  ST      JR209   L      60    54
## 14      31  ST       JR91   L      18     2
## 15      33  ST     MSH217   B      36    48
## 16      34  CH       JR32   B       9     8
## 17      35  CH        JR4   L       1     0
## 18      36  CH       JR20   L       1     1
```

```
## 19       38   CH       JR272   B       16     18
## 20       42   CH      MSH202   B       11      7
## 21       43   TL      MA1959           38     25
## 22       46   TL     SCI12-2   B        6      8
## 23       47   TL SPE123_2-3   B        8      4
## 24       48   TL SPE123_5-1   B       11      5
## 25       49   TL SPE123_6-3   B       32     19
## 26       53   CU SPE123_4-1   B        9     19
```

Strain 6 at 18°C only has one male and zero females. Remove strain 6 from the 18°C data.

```
aging_18 <- subset(aging_18, Strain!=6)
```

Remove strains 22, 35, and 36 from 25°C because they has fewer than 10 samples.

```
aging_25 <- subset(aging_25, Strain!=22 & Strain!=35 & Strain!=36)
```

Recalculate the strain count table with the low-count strains removed.

```
aging_strains_18 <- aging.strains(aging_18, strainIDs)
aging_strains_22 <- aging.strains(aging_22, strainIDs)
aging_strains_25 <- aging.strains(aging_25, strainIDs)
```

Write the data to file for analysis so the we do not need repeat the steps above.

```
write.table(aging_strains_18, "aging_strains_18.tsv",
            sep="\t", quote=FALSE, row.names=FALSE)
write.table(aging_strains_22, "aging_strains_22.tsv",
            sep="\t", quote=FALSE, row.names=FALSE)
write.table(aging_strains_25, "aging_strains_25.tsv",
            sep="\t", quote=FALSE, row.names=FALSE)
```

Create a second version of the data removing individuals that died one day after emergence by subsetting for flies that lived at least 2 days.

```
aging_18_no1 <- subset(aging_18, Age.at.death >= 2)
aging_22_no1 <- subset(aging_22, Age.at.death >= 2)
aging_25_no1 <- subset(aging_25, Age.at.death >= 2)
```

#### Cox proportional hazards analysis

Use the `coxph()` function in the `survival` package to model random effects as `frailty()`. The model will include the nested effect of isogen (within arrangement) as an interaction with arr. First, create functions to perform model comparisons.

```
# test for effect of arrangement
arr.effect.frailty <- function(agingData){
    anova(
        coxph(Surv(Age.at.death, censrec) ~ sex + arr +
                frailty(Birth), agingData),
        coxph(Surv(Age.at.death, censrec) ~ sex +
                frailty(Birth), agingData)
    )
}

# test for effect of sex
sex.effect.frailty <- function(agingData){
    anova(
```

```
        coxph(Surv(Age.at.death, censrec) ~ sex + arr +
                frailty(Birth), agingData),
        coxph(Surv(Age.at.death, censrec) ~ arr +
                frailty(Birth), agingData)
    )
}

# test for effect of interaction term
int.effect.frailty <- function(agingData){
    anova(
        coxph(Surv(Age.at.death, censrec) ~ sex + arr +
                frailty(Birth), agingData),
        coxph(Surv(Age.at.death, censrec) ~ sex + arr + sex*arr +
                frailty(Birth), agingData)
    )
}

# test for effect of isogen w/o interaction term
isogen.noint.effect.frailty <- function(agingData){
    anova(
        coxph(Surv(Age.at.death, censrec) ~ sex + arr +
                frailty(Birth), agingData),
        coxph(Surv(Age.at.death, censrec) ~ sex + arr + arr:isogen +
                frailty(Birth), agingData)
    )
}

# test for effect of isogen w/ interaction term
isogen.int.effect.frailty <- function(agingData){
    anova(
        coxph(Surv(Age.at.death, censrec) ~ sex + arr + sex*arr +
                frailty(Birth), agingData),
        coxph(Surv(Age.at.death, censrec) ~ sex + arr + sex*arr + arr:isogen +
                frailty(Birth), agingData)
    )
}

# test if interaction fits better w/isogen as random effect
int.isogen.effect.frailty <- function(agingData){
    anova(
        coxph(Surv(Age.at.death, censrec) ~ sex + arr + sex*arr + arr:isogen +
                frailty(Birth), agingData),
        coxph(Surv(Age.at.death, censrec) ~ sex + arr + arr:isogen +
                frailty(Birth), agingData)
    )
}
```

Create wrapper function to generate a data frame with the statistical test results.

```
coxph.frailty <- function(aging){
    DF <- data.frame(
        effect = c("arr", "sex", "int", "isogen-noint", "isogen-int", "int-isogen"),
        pval = c(
            arr.effect.frailty(aging)[2,4],
```

```
            sex.effect.frailty(aging)[2,4],
            int.effect.frailty(aging)[2,4],
            isogen.noint.effect.frailty(aging)[2,4],
            isogen.int.effect.frailty(aging)[2,4],
            int.isogen.effect.frailty(aging)[2,4]
        )
    )
    DF$sig <- NA
    DF$sig <- ""
    DF$sig[DF$pval<0.05] <- "*"
    DF$sig[DF$pval<0.005] <- "**"
    DF$sig[DF$pval<0.0005] <- "***"
    DF$sig[DF$pval<0.00005] <- "****"
    DF$sig[DF$pval<0.000005] <- "*****"
    DF
}
```

Perform the analysis on each temperature, and on both all the data and excluding 1 day old deaths.

```
coxph_frailty_18 <- coxph.frailty(aging_18)
coxph_frailty_22 <- coxph.frailty(aging_22)
coxph_frailty_25 <- coxph.frailty(aging_25)
coxph_frailty_18_no1 <- coxph.frailty(aging_18_no1)
coxph_frailty_22_no1 <- coxph.frailty(aging_22_no1)
coxph_frailty_25_no1 <- coxph.frailty(aging_25_no1)
```

Print outputs of each analysis for each temperature.

```
coxph_frailty_18
```

```
##         effect         pval   sig
## 1          arr 1.048113e-04   ***
## 2          sex 2.273614e-16 *****
## 3          int 5.772609e-01
## 4 isogen-noint 1.294569e-36 *****
## 5   isogen-int 3.425179e-37 *****
## 6   int-isogen 2.697711e-01
```

```
coxph_frailty_22
```

```
##         effect         pval   sig
## 1          arr 7.630497e-05   ***
## 2          sex 1.097174e-13 *****
## 3          int 6.700686e-01
## 4 isogen-noint 1.225026e-41 *****
## 5   isogen-int 2.714844e-42 *****
## 6   int-isogen 2.664117e-01
```

```
coxph_frailty_25
```

```
##         effect         pval   sig
## 1          arr 2.761260e-03    **
## 2          sex 1.614697e-06 *****
## 3          int 5.794525e-01
## 4 isogen-noint 2.696891e-33 *****
## 5   isogen-int 5.472231e-34 *****
## 6   int-isogen 2.031972e-01
```

At each temperature, the model that fits best has sex and arrangement effects with a strain effect (isogen) nested within arrangement. Adding an arrangement by sex interaction does not improve model fit. Similar results are observed with 1 day old deaths are excluded. Run this best fitting model for each temperature and each version of the data, and store the output.

```
aging_sex_arr_18 <- coxph(Surv(Age.at.death, censrec) ~ sex + arr + arr:isogen +
                        frailty(Birth), aging_18)
aging_sex_arr_22 <- coxph(Surv(Age.at.death, censrec) ~ sex + arr + arr:isogen +
                        frailty(Birth), aging_22)
aging_sex_arr_25 <- coxph(Surv(Age.at.death, censrec) ~ sex + arr + arr:isogen +
                        frailty(Birth), aging_25)
aging_sex_arr_18_no1 <- coxph(Surv(Age.at.death, censrec) ~ sex + arr + arr:isogen +
                            frailty(Birth), aging_18_no1)
aging_sex_arr_22_no1 <- coxph(Surv(Age.at.death, censrec) ~ sex + arr + arr:isogen +
                            frailty(Birth), aging_22_no1)
aging_sex_arr_25_no1 <- coxph(Surv(Age.at.death, censrec) ~ sex + arr + arr:isogen +
                            frailty(Birth), aging_25_no1)
```

In order to extract the effects of each arrangement on each sex, we need to run a model on data from each sex separately. First, set TL as reference because it is most often the longest lived.

```
aging_18$arr_factor <- factor(aging_18$arr,
                              levels = c("TL", "AR", "CH", "CU", "PP", "ST"))
aging_22$arr_factor <- factor(aging_22$arr,
                              levels = c("TL", "AR", "CH", "CU", "PP", "ST"))
aging_25$arr_factor <- factor(aging_25$arr,
                              levels = c("TL", "AR", "CH", "CU", "PP", "ST"))
aging_18_no1$arr_factor <- factor(aging_18_no1$arr,
                                levels = c("TL", "AR", "CH", "CU", "PP", "ST"))
aging_22_no1$arr_factor <- factor(aging_22_no1$arr,
                                levels = c("TL", "AR", "CH", "CU", "PP", "ST"))
aging_25_no1$arr_factor <- factor(aging_25_no1$arr,
                                levels = c("TL", "AR", "CH", "CU", "PP", "ST"))
```

This analysis must be performed using `coxme` in order to estimate arrangement effects because we need to treat isogen as random effect. We previously used the `coxph()` function, which only allows for fixed effects and frailty. Use `coxme()` to run a model with arrangement as a fixed effect (factored to make TL the reference), and the following random effects: isogen (strain) and Birth (batch).

```
aging_sex_arr_18_male <- coxme(Surv(Age.at.death, censrec) ~ arr_factor +
                               (1 | arr_factor/isogen) + (1|Birth),
                             subset(aging_18, sex=="male"))
aging_sex_arr_22_male <- coxme(Surv(Age.at.death, censrec) ~ arr_factor +
                               (1 | arr_factor/isogen) + (1|Birth),
                             subset(aging_22, sex=="male"))
aging_sex_arr_25_male <- coxme(Surv(Age.at.death, censrec) ~ arr_factor +
                               (1 | arr_factor/isogen) + (1|Birth),
                             subset(aging_25, sex=="male"))
aging_sex_arr_18_no1_male <- coxme(Surv(Age.at.death, censrec) ~  arr_factor +
                                   (1 | arr_factor/isogen) + (1|Birth),
                                 subset(aging_18_no1, sex=="male"))
aging_sex_arr_22_no1_male <- coxme(Surv(Age.at.death, censrec) ~  arr_factor +
                                   (1 | arr_factor/isogen) + (1|Birth),
                                 subset(aging_22_no1, sex=="male"))
aging_sex_arr_25_no1_male <- coxme(Surv(Age.at.death, censrec) ~  arr_factor +
                                   (1 | arr_factor/isogen) + (1|Birth),
```

```
                                    subset(aging_25_no1, sex=="male"))

aging_sex_arr_18_female <- coxme(Surv(Age.at.death, censrec) ~ arr_factor +
                                  (1 | arr_factor/isogen) + (1|Birth),
                                subset(aging_18, sex=="female"))
aging_sex_arr_22_female <- coxme(Surv(Age.at.death, censrec) ~ arr_factor +
                                  (1 | arr_factor/isogen) + (1|Birth),
                                subset(aging_22, sex=="female"))
aging_sex_arr_25_female <- coxme(Surv(Age.at.death, censrec) ~ arr_factor +
                                  (1 | arr_factor/isogen) + (1|Birth),
                                subset(aging_25, sex=="female"))
aging_sex_arr_18_no1_female <- coxme(Surv(Age.at.death, censrec) ~  arr_factor +
                                      (1 | arr_factor/isogen) + (1|Birth),
                                    subset(aging_18_no1, sex=="female"))
aging_sex_arr_22_no1_female <- coxme(Surv(Age.at.death, censrec) ~  arr_factor +
                                      (1 | arr_factor/isogen) + (1|Birth),
                                    subset(aging_22_no1, sex=="female"))
aging_sex_arr_25_no1_female <- coxme(Surv(Age.at.death, censrec) ~  arr_factor +
                                      (1 | arr_factor/isogen) + (1|Birth),
                                    subset(aging_25_no1, sex=="female"))
```

Create a single object with all of the statistical test results.

```
aging_sex_arr_coeff <- rbind(
    cbind(sex="male", temp="18C",
          data.frame(summary(aging_sex_arr_18_male)$coefficients[1:5,])),
    cbind(sex="male", temp="22C",
          data.frame(summary(aging_sex_arr_22_male)$coefficients[1:5,])),
    cbind(sex="male", temp="25C",
          data.frame(summary(aging_sex_arr_25_male)$coefficients[1:5,])),
    cbind(sex="female", temp="18C",
          data.frame(summary(aging_sex_arr_18_female)$coefficients[1:5,])),
    cbind(sex="female", temp="22C",
          data.frame(summary(aging_sex_arr_22_female)$coefficients[1:5,])),
    cbind(sex="female", temp="25C",
          data.frame(summary(aging_sex_arr_25_female)$coefficients[1:5,]))
)

aging_sex_arr_coeff_no1 <- rbind(
    cbind(sex="male", temp="18C",
          data.frame(summary(aging_sex_arr_18_no1_male)$coefficients[1:5,])),
    cbind(sex="male", temp="22C",
          data.frame(summary(aging_sex_arr_22_no1_male)$coefficients[1:5,])),
    cbind(sex="male", temp="25C",
          data.frame(summary(aging_sex_arr_25_no1_male)$coefficients[1:5,])),
    cbind(sex="female", temp="18C",
          data.frame(summary(aging_sex_arr_18_no1_female)$coefficients[1:5,])),
    cbind(sex="female", temp="22C",
          data.frame(summary(aging_sex_arr_22_no1_female)$coefficients[1:5,])),
    cbind(sex="female", temp="25C",
          data.frame(summary(aging_sex_arr_25_no1_female)$coefficients[1:5,]))
)
```

Print the output.

```
aging_sex_arr_coeff
```

```
##                    sex temp        coef exp.coef.  se.coef.     z          p
## arr_factorAR    male  18C  0.35872029 1.4314963 0.3603615  1.00 0.31951950
## arr_factorCH    male  18C  0.30936533 1.3625601 0.3742184  0.83 0.40840871
## arr_factorCU    male  18C  0.34774521 1.4158715 0.6328080  0.55 0.58264368
## arr_factorPP    male  18C  0.80118549 2.2281808 0.3697094  2.17 0.03022967
## arr_factorST    male  18C  0.52010598 1.6822059 0.3979511  1.31 0.19122647
## arr_factorAR1   male  22C  0.22557436 1.2530422 0.3743432  0.60 0.54678342
## arr_factorCH1   male  22C  0.56012015 1.7508829 0.3928081  1.43 0.15388608
## arr_factorCU1   male  22C  0.18174980 1.1993141 0.6479367  0.28 0.77908971
## arr_factorPP1   male  22C  0.55222841 1.7371197 0.3773705  1.46 0.14336922
## arr_factorST1   male  22C  0.19452722 1.2147365 0.4136872  0.47 0.63819227
## arr_factorAR2   male  25C -0.11398597 0.8922705 0.3616813 -0.32 0.75264333
## arr_factorCH2   male  25C  0.02322196 1.0234937 0.4184900  0.06 0.95574819
## arr_factorCU2   male  25C  0.27292695 1.3138043 0.5749826  0.47 0.63502225
## arr_factorPP2   male  25C -0.17923334 0.8359108 0.3594335 -0.50 0.61802239
## arr_factorST2   male  25C  0.03657671 1.0372539 0.3810820  0.10 0.92353549
## arr_factorAR3 female  18C  0.25496762 1.2904198 0.3238004  0.79 0.43103479
## arr_factorCH3 female  18C -0.03728966 0.9633970 0.3370545 -0.11 0.91190665
## arr_factorCU3 female  18C  0.06105728 1.0629598 0.5879873  0.10 0.91729540
## arr_factorPP3 female  18C  0.51377689 1.6715927 0.3263649  1.57 0.11543188
## arr_factorST3 female  18C  0.34715831 1.4150407 0.3542572  0.98 0.32710528
## arr_factorAR4 female  22C  0.46480539 1.5917044 0.3408228  1.36 0.17263861
## arr_factorCH4 female  22C  0.44232092 1.5563151 0.3577976  1.24 0.21637221
## arr_factorCU4 female  22C  0.30760843 1.3601683 0.5981556  0.51 0.60706913
## arr_factorPP4 female  22C  0.35742874 1.4296487 0.3479817  1.03 0.30435073
## arr_factorST4 female  22C  0.08610819 1.0899242 0.3758687  0.23 0.81879809
## arr_factorAR5 female  25C  0.19117400 1.2106701 0.3938361  0.49 0.62738188
## arr_factorCH5 female  25C  0.41053570 1.5076252 0.4663624  0.88 0.37870041
## arr_factorCU5 female  25C  0.49384791 1.6386093 0.7063051  0.70 0.48442761
## arr_factorPP5 female  25C  0.12185604 1.1295915 0.3960101  0.31 0.75830346
## arr_factorST5 female  25C  0.10654460 1.1124275 0.4111980  0.26 0.79555207
```

The only significant result is that PP significantly increases the hazard in males at 18°C. Test if significant PP stands up to multiple testing correction.

```
aging_sex_arr_coeff$padj <- p.adjust(aging_sex_arr_coeff$p, method="fdr")
```

Nothing is significant after adjusting for FDR. Similar results are observed when 1 day old deaths are excluded.

We can also extract the sex effects within arrangements by subsetting for arrangement and comparing males and females.

```
sex.coxph.frailty <- function(aging){
  arrangements <- levels(factor(aging$arr))
  return.df <- data.frame(
    arr = arrangements,
    coef=NA, se=NA, se2=NA, Chisq=NA, DF=NA, p=NA
  )
  for(i in 1:length(arrangements)){
    if(length(levels(factor(subset(aging, arr==arrangements[i])$isogen))) > 1){
       return.df[i,c(2:7)] <- summary(coxph(
          Surv(Age.at.death, censrec) ~ sex + isogen + frailty(Birth),
          subset(aging, arr==arrangements[i])
        ))$coefficients[1]
```

```r
    } else{
      return.df[i,c(2:7)] <- summary(coxph(
        Surv(Age.at.death, censrec) ~ sex + frailty(Birth),
        subset(aging, arr==arrangements[i])
       ))$coefficients[1]
    }
  }
  return.df
}
```

```r
sex.coxph.frailty(aging_18)
```

```
##   arr      coef        se       se2     Chisq        DF         p
## 1  AR 0.5531039 0.5531039 0.5531039 0.5531039 0.5531039 0.5531039
## 2  CH 0.8493722 0.8493722 0.8493722 0.8493722 0.8493722 0.8493722
## 3  CU 1.1289534 1.1289534 1.1289534 1.1289534 1.1289534 1.1289534
## 4  PP 0.6040457 0.6040457 0.6040457 0.6040457 0.6040457 0.6040457
## 5  ST 0.6490159 0.6490159 0.6490159 0.6490159 0.6490159 0.6490159
## 6  TL 0.4533357 0.4533357 0.4533357 0.4533357 0.4533357 0.4533357
```

```r
sex.coxph.frailty(aging_22)
```

```
##   arr      coef        se       se2     Chisq        DF         p
## 1  AR 0.2525086 0.2525086 0.2525086 0.2525086 0.2525086 0.2525086
## 2  CH 0.6420033 0.6420033 0.6420033 0.6420033 0.6420033 0.6420033
## 3  CU 0.2872802 0.2872802 0.2872802 0.2872802 0.2872802 0.2872802
## 4  PP 0.6272270 0.6272270 0.6272270 0.6272270 0.6272270 0.6272270
## 5  ST 0.6546943 0.6546943 0.6546943 0.6546943 0.6546943 0.6546943
## 6  TL 0.4452134 0.4452134 0.4452134 0.4452134 0.4452134 0.4452134
```

```r
sex.coxph.frailty(aging_25)
```

```
## Warning in coxpenal.fit(X, Y, istrat, offset, init = init, control, weights =
## weights, : Inner loop failed to coverge for iterations 2 3
```

```
##   arr      coef        se       se2     Chisq        DF         p
## 1  AR 0.4163490 0.4163490 0.4163490 0.4163490 0.4163490 0.4163490
## 2  CH 0.2618812 0.2618812 0.2618812 0.2618812 0.2618812 0.2618812
## 3  CU 0.4130043 0.4130043 0.4130043 0.4130043 0.4130043 0.4130043
## 4  PP 0.2319531 0.2319531 0.2319531 0.2319531 0.2319531 0.2319531
## 5  ST 0.8863643 0.8863643 0.8863643 0.8863643 0.8863643 0.8863643
## 6  TL 0.5675608 0.5675608 0.5675608 0.5675608 0.5675608 0.5675608
```

Using this approach, there is not a significant sex difference within temperatures and arrangements, but it may not be the best way to test for sex differences.

#### Fit Gompertz models

Next, use Gompertz models to test for sex differences in within each arrangement. The first goal is to test for sex effects on the rate and shape parameters in Gompertz model. The focus is therefore on comparisons that evaluate if sex improves the model fit.

Create a function to perform model comparison with different predictors. The function first fits models with different rate parameters (shape will be added later). Then, the function using a chi-square test to compare the log likelihood of nested models. Birth (batch), isogen (strain), and sex are included as rate parameters. The function is passed a subset of the data corresponding to a particular arrangement

```r
flexsurv_model_comparisons <- function(agingDataArr){
    NULLmodel <- flexsurvreg(Surv(Age.at.death, censrec) ~ 1,
                             data=agingDataArr, dist = "gompertz")
    batchEffect <- flexsurvreg(Surv(Age.at.death, censrec) ~ Birth,
                               data=agingDataArr, dist = "gompertz")
    sexEffect <- flexsurvreg(Surv(Age.at.death, censrec) ~ sex,
                             data=agingDataArr, dist = "gompertz")
    strainEffect <- flexsurvreg(Surv(Age.at.death, censrec) ~ isogen,
                                data=agingDataArr, dist = "gompertz")
    sexbatch <- flexsurvreg(Surv(Age.at.death, censrec) ~ sex + Birth,
                            data=agingDataArr, dist = "gompertz")
    sexstrain <- flexsurvreg(Surv(Age.at.death, censrec) ~ sex + isogen,
                             data=agingDataArr, dist = "gompertz")
    strainbatch <- flexsurvreg(Surv(Age.at.death, censrec) ~ isogen + Birth,
                               data=agingDataArr, dist = "gompertz")
    sexstrainbatch <- flexsurvreg(Surv(Age.at.death, censrec) ~ sex + isogen + Birth,
                                  data=agingDataArr, dist = "gompertz")

    return.df <- data.frame(
        factors = c("batch", "sex", "strain", "sexbatch", "sexstrain", "sexstrainbatch"),
        pval = c(
            pchisq(2 * (logLik(batchEffect) - logLik(NULLmodel)),
                   df = abs(deg_free(logLik(batchEffect) - deg_free(logLik(NULLmodel)))),
                   lower.tail = FALSE),
            pchisq(2 * (logLik(sexEffect) - logLik(NULLmodel)),
                   df = abs(deg_free(logLik(sexEffect) - deg_free(logLik(NULLmodel)))),
                   lower.tail = FALSE),
            pchisq(2 * (logLik(strainEffect) - logLik(NULLmodel)),
                   df = abs(deg_free(logLik(strainEffect) - deg_free(logLik(NULLmodel)))),
                   lower.tail = FALSE),
            pchisq(2 * (logLik(sexbatch) - logLik(batchEffect)),
                   df = abs(deg_free(logLik(sexbatch) - deg_free(logLik(batchEffect)))),
                   lower.tail = FALSE),
            pchisq(2 * (logLik(sexstrain) - logLik(strainEffect)),
                   df = abs(deg_free(logLik(sexstrain) - deg_free(logLik(strainEffect)))),
                   lower.tail = FALSE),
            pchisq(2 * (logLik(sexstrainbatch) - logLik(strainbatch)),
                   df = abs(deg_free(logLik(sexstrainbatch) - deg_free(logLik(strainbatch)))),
                   lower.tail = FALSE)
        ),
        significant = NA
    )

    return.df$significant <- ""
    return.df$significant[return.df$pval < 0.05] <- "*"
    return.df$significant[return.df$pval < 0.005] <- "**"
    return.df$significant[return.df$pval < 0.0005] <- "***"
    return.df$significant[return.df$pval < 0.00005] <- "****"
    return.df$significant[return.df$pval < 0.000005] <- "*****"

    return.df
}
```

Need a separate function for CU because there is only one strain (isogen for CU), which means we cannot test for isogen effects for CU.

```
flexsurv_model_comparisons_CU <- function(agingDataArr){
    NULLmodel <- flexsurvreg(Surv(Age.at.death, censrec) ~ 1,
                             data=agingDataArr, dist = "gompertz")
    batchEffect <- flexsurvreg(Surv(Age.at.death, censrec) ~ Birth,
                               data=agingDataArr, dist = "gompertz")
    sexEffect <- flexsurvreg(Surv(Age.at.death, censrec) ~ sex,
                             data=agingDataArr, dist = "gompertz")
    sexbatch <- flexsurvreg(Surv(Age.at.death, censrec) ~ sex + Birth,
                            data=agingDataArr, dist = "gompertz")

    return.df <- data.frame(
        factors = c("batch", "sex", "sexbatch"),
        pval = c(
            pchisq(2 * (logLik(batchEffect) - logLik(NULLmodel)),
                   df = abs(deg_free(logLik(batchEffect) - deg_free(logLik(NULLmodel)))),
                   lower.tail = FALSE),
            pchisq(2 * (logLik(sexEffect) - logLik(NULLmodel)),
                   df = abs(deg_free(logLik(sexEffect) - deg_free(logLik(NULLmodel)))),
                   lower.tail = FALSE),
            pchisq(2 * (logLik(sexbatch) - logLik(batchEffect)),
                   df = abs(deg_free(logLik(sexbatch) - deg_free(logLik(batchEffect)))),
                   lower.tail = FALSE)
        ),
        significant = NA
    )

    return.df$significant <- ""
    return.df$significant[return.df$pval < 0.05] <- "*"
    return.df$significant[return.df$pval < 0.005] <- "**"
    return.df$significant[return.df$pval < 0.0005] <- "***"
    return.df$significant[return.df$pval < 0.00005] <- "****"
    return.df$significant[return.df$pval < 0.000005] <- "******"

    return.df
}
```

Lastly, here is a wrapper function to call the functions above for data from each arrangement.

```
flexsurv_arrangement_sex <- function(agingData){
    rbind(
        cbind(  arr="AR",
                flexsurv_model_comparisons(subset(agingData, arr=="AR")) ),
        cbind(  arr="CH",
                flexsurv_model_comparisons(subset(agingData, arr=="CH")) ),
        cbind(  arr="PP",
                flexsurv_model_comparisons(subset(agingData, arr=="PP")) ),
        cbind(  arr="ST",
                flexsurv_model_comparisons(subset(agingData, arr=="ST")) ),
        cbind(  arr="TL",
                flexsurv_model_comparisons(subset(agingData, arr=="TL")) ),
        cbind(  arr="CU",
                flexsurv_model_comparisons_CU(subset(agingData, arr=="CU")) )
    )
```

```
}
```

Call the wrapper function for data from each temperature.

```
fs_arr_sex_18 <- flexsurv_arrangement_sex(aging_18)
fs_arr_sex_22 <- flexsurv_arrangement_sex(aging_22)
fs_arr_sex_25 <- flexsurv_arrangement_sex(aging_25)

fs_arr_sex_18_no1 <- flexsurv_arrangement_sex(aging_18_no1)
fs_arr_sex_22_no1 <- flexsurv_arrangement_sex(aging_22_no1)
fs_arr_sex_25_no1 <- flexsurv_arrangement_sex(aging_25_no1)
```

Create a single data frame that summarizes the statistical test results for each temperature. Make a separate object for the entire data set and the data excluding death after one day.

```
fs_arr_sex <- cbind(
    fs_arr_sex_18[,1:2],
    pval_18 = fs_arr_sex_18$pval,
    pval_22 = fs_arr_sex_22$pval,
    pval_25 = fs_arr_sex_25$pval,
    sig_18 = fs_arr_sex_18$significant,
    sig_22 = fs_arr_sex_22$significant,
    sig_25 = fs_arr_sex_25$significant
    )

fs_arr_sex_no1 <- cbind(
    fs_arr_sex_18_no1[,1:2],
    pval_18 = fs_arr_sex_18_no1$pval,
    pval_22 = fs_arr_sex_22_no1$pval,
    pval_25 = fs_arr_sex_25_no1$pval,
    sig_18 = fs_arr_sex_18_no1$significant,
    sig_22 = fs_arr_sex_22_no1$significant,
    sig_25 = fs_arr_sex_25_no1$significant
    )
```

Print the statistical test results.

```
fs_arr_sex
```

```
##    arr        factors      pval_18      pval_22      pval_25 sig_18 sig_22
## 1   AR          batch 3.792636e-01 2.824142e-03 1.552141e-07            **
## 2   AR            sex 5.913371e-04 1.930397e-01 3.730429e-01     **
## 3   AR         strain 3.142091e-12 1.164331e-11 9.737185e-15 ****** ******
## 4   AR       sexbatch 3.980583e-01 9.885262e-01 1.000000e+00
## 5   AR      sexstrain 1.420638e-02 9.420511e-01 7.603209e-01      *
## 6   AR sexstrainbatch 8.875064e-01 9.999330e-01 9.999841e-01
## 7   CH          batch 3.070113e-03 8.044632e-02 7.708400e-03     **
## 8   CH            sex 9.072445e-03 2.120251e-02 9.979968e-01      *      *
## 9   CH         strain 2.816175e-02 2.330268e-11 2.866934e-01      * ******
## 10  CH       sexbatch 1.182125e-01 3.873442e-01 1.000000e+00
## 11  CH      sexstrain 3.180800e-03 4.430331e-01 9.995685e-01     **
## 12  CH sexstrainbatch 5.687731e-02 9.691953e-01 1.000000e+00
## 13  PP          batch 1.527470e-03 4.140879e-01 6.582813e-07     **
## 14  PP            sex 4.020262e-03 3.316776e-02 5.100393e-01     **      *
## 15  PP         strain 7.916229e-06 8.435368e-07 1.101010e-02   **** ******
## 16  PP       sexbatch 8.068148e-01 8.453166e-01 1.000000e+00
```

```
## 17  PP      sexstrain 1.522953e-02 6.073530e-02 9.856487e-01      *
## 18  PP sexstrainbatch 9.361280e-01 7.643709e-01 1.000000e+00
## 19  ST          batch 6.680801e-01 3.192002e-08 3.565691e-05        ******
## 20  ST            sex 1.028928e-02 7.074599e-02 1.703532e-01      *
## 21  ST         strain 1.318448e-10 1.326004e-05 1.179052e-06 ******   ****
## 22  ST       sexbatch 5.755970e-01 1.867263e-01 9.660816e-01
## 23  ST      sexstrain 4.862961e-02 8.496261e-02 2.265835e-01      *
## 24  ST sexstrainbatch 4.125961e-01 3.034112e-01 7.141946e-01
## 25  TL          batch 3.821246e-01 1.088678e-08 7.314419e-05        ******
## 26  TL            sex 9.200339e-02 2.084429e-01 1.455978e-01
## 27  TL         strain 3.111155e-04 4.502272e-04 1.460792e-09    ***    ***
## 28  TL       sexbatch 7.931764e-01 9.516971e-01 9.997854e-01
## 29  TL      sexstrain 4.332041e-01 6.992015e-01 4.530910e-01
## 30  TL sexstrainbatch 8.405366e-01 9.627485e-01 9.998913e-01
## 31  CU          batch 1.779376e-02 2.566775e-02 1.685699e-02      *      *
## 32  CU            sex 7.142787e-03 7.751829e-01 6.336921e-01      *
## 33  CU       sexbatch 4.696740e-01 1.000000e+00 9.990458e-01
##    sig_25
## 1  ******
## 2
## 3  ******
## 4
## 5
## 6
## 7       *
## 8
## 9
## 10
## 11
## 12
## 13 ******
## 14
## 15      *
## 16
## 17
## 18
## 19   ****
## 20
## 21 ******
## 22
## 23
## 24
## 25    ***
## 26
## 27 ******
## 28
## 29
## 30
## 31      *
## 32
## 33
```

Including strain (isogen) and batch (birth) as a rate parameter often improves model fit. Including a sex effect significantly improves the fit of the model for most arrangements, but only at 18°C, demonstrating that sex affects Gompertz rate parameter at the lowest temperature. However, the effect of sex is not as consistent at 22°C or 25°C. Similar results are observed when 1 day old deaths are excluded.

Next, add a shape parameter to the Gompertz models, and test if adding sex as a shape effect significantly improves model fit. In all cases, include Birth and isogen in all models as rate parameters because they affect rate as demonstrated above. Do not analyze CU because cannot perform analysis with isogen as rate parameter with only on CU strain sampled.

```
shape_model_comparisons <- function(agingDataArr){
    NULLmodel <- flexsurvreg(Surv(Age.at.death, censrec) ~ Birth + isogen,
                  anc = list(shape = ~ 1), data = agingDataArr, dist = "gompertz")
    sexShape <- flexsurvreg(Surv(Age.at.death, censrec) ~ Birth + isogen,
                  anc = list(shape = ~ sex), data = agingDataArr, dist = "gompertz")
    sexRate <- flexsurvreg(Surv(Age.at.death, censrec) ~ Birth + isogen + sex,
                  anc = list(shape = ~ 1), data = agingDataArr, dist = "gompertz")
    sexShapeRate <- flexsurvreg(Surv(Age.at.death, censrec) ~ Birth + isogen + sex,
                  anc = list(shape = ~ sex), data = agingDataArr, dist = "gompertz")

    return.df <- data.frame(
        factors = c("sexShapeNULL", "sexRateNULL", "sexRateShape", "sexShapeRate"),
        pval = c(

            pchisq(2 * (logLik(sexShape) - logLik(NULLmodel)),
                   df = abs(deg_free(logLik(sexShape) - deg_free(logLik(NULLmodel)))),
                   lower.tail = FALSE),
            pchisq(2 * (logLik(sexRate) - logLik(NULLmodel)),
                   df = abs(deg_free(logLik(sexRate) - deg_free(logLik(NULLmodel)))),
                   lower.tail = FALSE),
            pchisq(2 * (logLik(sexShapeRate) - logLik(sexShape)),
                   df = abs(deg_free(logLik(sexShapeRate) - deg_free(logLik(sexShape)))),
                   lower.tail = FALSE), # test if adding Rate to a model with Shape improves fit
            pchisq(2 * (logLik(sexShapeRate) - logLik(sexRate)),
                   df = abs(deg_free(logLik(sexShapeRate) - deg_free(logLik(sexRate)))),
                   lower.tail = FALSE) # test if adding Shape to a model with Rate improves fit
        ),
        significant = NA
    )

    return.df$significant <- ""
    return.df$significant[return.df$pval < 0.05] <- "*"
    return.df$significant[return.df$pval < 0.005] <- "**"
    return.df$significant[return.df$pval < 0.0005] <- "***"
    return.df$significant[return.df$pval < 0.00005] <- "****"
    return.df$significant[return.df$pval < 0.000005] <- "******"

    return.df
}
```

Wrapper function to run analysis on data from each arrangement.

```
shape_arrangement_sex <- function(agingData){
    rbind(
        cbind(  arrangement="AR",
                shape_model_comparisons(subset(agingData, arr=="AR")) ),
        cbind(  arrangement="CH",
                shape_model_comparisons(subset(agingData, arr=="CH")) ),
```

```r
        cbind(  arrangement="PP",
                shape_model_comparisons(subset(agingData, arr=="PP")) ),
        cbind(  arrangement="ST",
                shape_model_comparisons(subset(agingData, arr=="ST")) ),
        cbind(  arrangement="TL",
                shape_model_comparisons(subset(agingData, arr=="TL")) )
    )
}
```

Call wrapper function for data from each temperature.

```r
shape_arr_sex_18 <- shape_arrangement_sex(aging_18)
shape_arr_sex_22 <- shape_arrangement_sex(aging_22)
shape_arr_sex_25 <- shape_arrangement_sex(aging_25)

shape_arr_sex_18_no1 <- shape_arrangement_sex(aging_18_no1)
shape_arr_sex_22_no1 <- shape_arrangement_sex(aging_22_no1)
shape_arr_sex_25_no1 <- shape_arrangement_sex(aging_25_no1)
```

Generate a single data frame with statistical test results for complete dataset and data excluding 1 day old deaths.

```r
shape_arr_sex <- cbind(
    shape_arr_sex_18[,1:2],
    pval_18 = shape_arr_sex_18$pval,
    pval_22 = shape_arr_sex_22$pval,
    pval_25 = shape_arr_sex_25$pval,
    sig_18 = shape_arr_sex_18$significant,
    sig_22 = shape_arr_sex_22$significant,
    sig_25 = shape_arr_sex_25$significant
)

shape_arr_sex_no1 <- cbind(
    shape_arr_sex_18_no1[,1:2],
    pval_18 = shape_arr_sex_18_no1$pval,
    pval_22 = shape_arr_sex_22_no1$pval,
    pval_25 = shape_arr_sex_25_no1$pval,
    sig_18 = shape_arr_sex_18_no1$significant,
    sig_22 = shape_arr_sex_22_no1$significant,
    sig_25 = shape_arr_sex_25_no1$significant
)

shape_arr_sex
```

```
##    arrangement      factors    pval_18   pval_22   pval_25 sig_18 sig_22 sig_25
## 1           AR sexShapeNULL 0.99829765 0.9998564 1.0000000
## 2           AR  sexRateNULL 0.88750643 0.9999330 0.9999841
## 3           AR sexRateShape 0.99657467 1.0000000 0.9999992
## 4           AR sexShapeRate 1.00000000 1.0000000 1.0000000
## 5           CH sexShapeNULL 0.76732628 1.0000000 1.0000000
## 6           CH  sexRateNULL 0.05687731 0.9691953 1.0000000
## 7           CH sexRateShape 0.50710651 0.6554142 1.0000000
## 8           CH sexShapeRate 0.99998851 0.9839496 1.0000000
## 9           PP sexShapeNULL 0.98447844 0.9467875 0.9999999
## 10          PP  sexRateNULL 0.93612800 0.7643709 1.0000000
```

```
## 11          PP sexRateShape 0.99999929 0.9997777 1.0000000
## 12          PP sexShapeRate 1.00000000 1.0000000 1.0000000
## 13          ST sexShapeNULL 0.99002363 0.9080470 0.9691727
## 14          ST  sexRateNULL 0.41259614 0.3034112 0.7141946
## 15          ST sexRateShape 0.62714066 0.9128108 0.9987755
## 16          ST sexShapeRate 0.99964819 1.0000000 1.0000000
## 17          TL sexShapeNULL 0.90166979 0.9993280 0.9999992
## 18          TL  sexRateNULL 0.84053659 0.9627485 0.9998913
## 19          TL sexRateShape 0.99999996 0.9992359 1.0000000
## 20          TL sexShapeRate 1.00000000 1.0000000 1.0000000
```

Adding a shape parameter does not significantly improve model fit at any temperature. We therefore conclude that there is not a significant effect of sex on the shape of the survival curve.

We will next compare arrangements within each sex. The model will include batch (Birth) as a predictor, but not isogen (strain) because flexsurvreg cannot handle isogen as nested or interacting factor in model with arrangement. Define a function to perform the model comparisons.

```
arrangement.comparisons <- function(agingDataArr){
    batchRate <- flexsurvreg(Surv(Age.at.death, censrec) ~ Birth,
                             anc = list(shape = ~ 1),
                             data=agingDataArr, dist = "gompertz")
    arrRate <- flexsurvreg(Surv(Age.at.death, censrec) ~ Birth + arr,
                           anc = list(shape = ~ 1),
                           data=agingDataArr, dist = "gompertz")
    arrShape <- flexsurvreg(Surv(Age.at.death, censrec) ~ Birth,
                            anc = list(shape = ~ arr),
                            data=agingDataArr, dist = "gompertz")
    arrRateShape <- flexsurvreg(Surv(Age.at.death, censrec) ~ Birth + arr,
                                anc = list(shape = ~ arr),
                                data=agingDataArr, dist = "gompertz")

    return.df <- data.frame(
        factors = c("arrRate", "arrShape", "arrRateShape", "arrShapeRate"),
        pval = c(
            pchisq(2 * (logLik(arrRate) - logLik(batchRate)),
                   df = abs(deg_free(logLik(arrRate) - deg_free(logLik(batchRate)))),
                   lower.tail = FALSE),
            pchisq(2 * (logLik(arrShape) - logLik(batchRate)),
                   df = abs(deg_free(logLik(arrShape) - deg_free(logLik(batchRate)))),
                   lower.tail = FALSE),
            pchisq(2 * (logLik(arrRateShape) - logLik(arrShape)),
                   df = abs(deg_free(logLik(arrRateShape) - deg_free(logLik(arrShape)))),
                   lower.tail = FALSE),
            pchisq(2 * (logLik(arrRateShape) - logLik(arrRate)),
                   df = abs(deg_free(logLik(arrRateShape) - deg_free(logLik(arrRate)))),
                   lower.tail = FALSE)
        ),
        significant = NA
    )

    return.df$significant <- ""
    return.df$significant[return.df$pval < 0.05] <- "*"
    return.df$significant[return.df$pval < 0.005] <- "**"
    return.df$significant[return.df$pval < 0.0005] <- "***"
```

```
    return.df$significant[return.df$pval < 0.00005] <- "****"
    return.df$significant[return.df$pval < 0.000005] <- "******"

    return.df
}
```

Use a wrapper function to perform analysis on male and female data separately for each temperature. First, analyze the complete data set.

```
arr.models <- function(agingData){
    rbind(
        cbind(  sex="female",
                arrangement.comparisons(subset(agingData, sex=="female")) ),
        cbind(  sex="male",
                arrangement.comparisons(subset(agingData, sex=="male")) )
    )
}

arr_models_18 <- arr.models(aging_18)
arr_models_22 <- arr.models(aging_22)
arr_models_25 <- arr.models(aging_25)

arr_models <- cbind(
    arr_models_18[,1:2],
    pval_18 = arr_models_18$pval,
    pval_22 = arr_models_22$pval,
    pval_25 = arr_models_25$pval,
    sig_18 = arr_models_18$significant,
    sig_22 = arr_models_22$significant,
    sig_25 = arr_models_25$significant
)

arr_models
```

```
##      sex      factors    pval_18   pval_22   pval_25 sig_18 sig_22 sig_25
## 1 female      arrRate 0.63700468 0.5498012 0.9216044
## 2 female     arrShape 0.71448121 0.1654886 0.8348673
## 3 female arrRateShape 0.76855438 0.9272887 0.9999975
## 4 female arrShapeRate 0.82583111 0.5774295 0.9999133
## 5   male      arrRate 0.64624459 0.6941492 0.9999968
## 6   male     arrShape 0.90415154 0.9999803 0.9775719
## 7   male arrRateShape 0.01923095 0.8080563 1.0000000      *
## 8   male arrShapeRate 0.05622436 0.9999093 0.9998603
```

Next, analyze data excluding 1 day old deaths.

```
arr_models_18_no1 <- arr.models(aging_18_no1)
arr_models_22_no1 <- arr.models(aging_22_no1)
arr_models_25_no1 <- arr.models(aging_25_no1)

arr_models_no1 <- cbind(
    arr_models_18_no1[,1:2],
    pval_18 = arr_models_18_no1$pval,
    pval_22 = arr_models_22_no1$pval,
    pval_25 = arr_models_25_no1$pval,
```

```r
    sig_18 = arr_models_18_no1$significant,
    sig_22 = arr_models_22_no1$significant,
    sig_25 = arr_models_25_no1$significant
)

arr_models_no1
```

```
##      sex      factors    pval_18   pval_22   pval_25 sig_18 sig_22 sig_25
## 1 female      arrRate 0.68269190 0.5832379 0.9622938
## 2 female     arrShape 0.70175024 0.1611662 0.8310373
## 3 female arrRateShape 0.83932453 0.9254569 0.9999997
## 4 female arrShapeRate 0.85184079 0.5370851 0.9997912
## 5   male      arrRate 0.70012951 0.6775049 0.9999875
## 6   male     arrShape 0.90203990 0.9999788 0.9725611
## 7   male arrRateShape 0.02588404 0.7815487 1.0000000      *
## 8   male arrShapeRate 0.06077846 0.9998524 0.9999029
```

Including arrangement as a rate parameter improves model fit for males at 18°C, but not in any other sex or temperature combinations. Including arrangement as a shape parameter does not improve model fit in these data.

##### Compare across temperatures

To perform statistical analysis across temperatures, first create a data frames that include flies raised at all three temperatures.

```r
aging_temps <- rbind(
    cbind(aging_18, temp="18C"),
    cbind(aging_22, temp="22C"),
    cbind(aging_25, temp="25C")
)

aging_temps_no1 <- rbind(
    cbind(aging_18_no1, temp="18C"),
    cbind(aging_22_no1, temp="22C"),
    cbind(aging_25_no1, temp="25C")
)
```

Define functions to perform comparisons between nested Cox proportional hazard models.

```r
arr.effect.temp <- function(agingData){ # test for effect of sex
    anova(
        coxph(Surv(Age.at.death, censrec) ~ arr + temp +
                frailty(Birth), agingData),
        coxph(Surv(Age.at.death, censrec) ~ temp +
                frailty(Birth), agingData)
    )
}

arrisogen.effect.temp <- function(agingData){ # test for effect of sex
    anova(
        coxph(Surv(Age.at.death, censrec) ~ arr + temp + arr:isogen +
                frailty(Birth), agingData),
        coxph(Surv(Age.at.death, censrec) ~ arr + temp +
                frailty(Birth), agingData)
    )
```

```
}

sex.effect.temp <- function(agingData){ # test for effect of arrangement
    anova(
        coxph(Surv(Age.at.death, censrec) ~ sex + temp +
                frailty(Birth), agingData),
        coxph(Surv(Age.at.death, censrec) ~ temp +
                frailty(Birth), agingData)
    )
}

temp.effect.sexarrisogen <- function(agingData){ # test for effect of sex
    anova(
        coxph(Surv(Age.at.death, censrec) ~ arr + sex + arr:isogen + temp +
                frailty(Birth), agingData),
        coxph(Surv(Age.at.death, censrec) ~ arr + sex + arr:isogen +
                frailty(Birth), agingData)
    )
}

sexarrisogen.effect.temp <- function(agingData){ # test for effect of sex
    anova(
        coxph(Surv(Age.at.death, censrec) ~ sex + arr + temp + arr:isogen +
                frailty(Birth), agingData),
        coxph(Surv(Age.at.death, censrec) ~ arr + temp + arr:isogen +
                frailty(Birth), agingData)
    )
}

arrtempint.effect.temp <- function(agingData){ # test for effect of sex
    anova(
        coxph(Surv(Age.at.death, censrec) ~ arr + temp + arr*temp +
                frailty(Birth), agingData),
        coxph(Surv(Age.at.death, censrec) ~ arr + temp +
                frailty(Birth), agingData)
    )
}

arrtempintisogen.effect.temp <- function(agingData){ # test for effect of sex
    anova(
        coxph(Surv(Age.at.death, censrec) ~ arr + temp + arr*temp + arr:isogen +
                frailty(Birth), agingData),
        coxph(Surv(Age.at.death, censrec) ~ arr + temp + + arr:isogen +
                frailty(Birth), agingData)
    )
}

sextempint.effect.temp <- function(agingData){ # test for effect of arrangement
    anova(
        coxph(Surv(Age.at.death, censrec) ~ sex + temp + sex*temp +
                frailty(Birth), agingData),
        coxph(Surv(Age.at.death, censrec) ~ sex + temp +
                frailty(Birth), agingData)
```

```
    )
}

sexarrint.effect.temp <- function(agingData){ # test for effect of interaction term
    anova(
        coxph(Surv(Age.at.death, censrec) ~ sex + arr + temp +
                frailty(Birth), agingData),
        coxph(Surv(Age.at.death, censrec) ~ sex + arr + sex*arr + temp +
                frailty(Birth), agingData)
    )
}

sexarrintisogen.effect.temp <- function(agingData){ # test for effect of interaction term
    anova(
        coxph(Surv(Age.at.death, censrec) ~ sex + arr + temp + arr:isogen +
                frailty(Birth), agingData),
        coxph(Surv(Age.at.death, censrec) ~ sex + arr + sex*arr + temp + arr:isogen +
                frailty(Birth), agingData)
    )
}
```

Define a wrapper function to call each model comparison, and then run the model comparison on the data.

```
coxph.temp <- function(aging){
    DF <- data.frame(
        effect = c("arr", "isogen", "sex", "sex (w/isogen)", "temp",
                "arr-temp-int", "arr-temp-int (w/ isogen)",
                "sex-temp-int", "sex-arr-int", "sex-arr-int (w/isogen)"),
        pval = c(
            arr.effect.temp(aging)[2,4],
            arrisogen.effect.temp(aging)[2,4],
            sex.effect.temp(aging)[2,4],
            temp.effect.sexarrisogen(aging)[2,4],
            sexarrisogen.effect.temp(aging)[2,4],
            arrtempint.effect.temp(aging)[2,4],
            arrtempintisogen.effect.temp(aging)[2,4],
            sextempint.effect.temp(aging)[2,4],
            sexarrint.effect.temp(aging)[2,4],
            sexarrintisogen.effect.temp(aging)[2,4]
        )
    )
    DF$sig <- NA
    DF$sig <- ""
    DF$sig[DF$pval<0.05] <- "*"
    DF$sig[DF$pval<0.005] <- "**"
    DF$sig[DF$pval<0.0005] <- "***"
    DF$sig[DF$pval<0.00005] <- "****"
    DF$sig[DF$pval<0.000005] <- "*****"
    DF
}

coxph_temps <- coxph.temp(aging_temps)
coxph_temps_no1 <- coxph.temp(aging_temps_no1)
```

Print the results of the analysis of the complete data.

```
coxph_temps
```

```
##                       effect          pval   sig
## 1                        arr  5.617012e-10 *****
## 2                     isogen  8.855853e-86 *****
## 3                        sex  3.095595e-27 *****
## 4             sex (w/isogen) 8.901585e-182 *****
## 5                       temp  2.293052e-20 *****
## 6               arr-temp-int  2.447073e-04   ***
## 7  arr-temp-int (w/ isogen)  8.638283e-07 *****
## 8               sex-temp-int  7.947923e-01
## 9                sex-arr-int  4.206939e-01
## 10   sex-arr-int (w/isogen)  9.999891e-01
```

Model fit is significantly improved when including arrangement, sex, temperature and interaction between arrangement and temperature. Similar results are observed excluding 1 day old deaths.

Run the model that fits best to extract key model components. First, run the model on all of the data.

```
# run model
aging_sex_arr_temp <- coxph(Surv(Age.at.death, censrec) ~
                              sex + arr + temp + arr*temp + arr:isogen +
                              frailty(Birth), aging_temps)

# extract coefficients from model
aging_sex_arr_temp_coeff <- data.frame(summary(aging_sex_arr_temp)$coefficients)
aging_sex_arr_temp_coeff$hazard <- exp(aging_sex_arr_temp_coeff$coef)
subset(aging_sex_arr_temp_coeff, p<0.05)
```

```
##                            coef   se.coef.        se2       Chisq       DF
## sexmale               0.4477244 0.03599392 0.03598776  154.725993  1.00000
## arrCH                 7.2864903 0.15331728 0.15327873 2258.679426  1.00000
## arrCU                 7.0700645 0.19105846 0.19100039 1369.349110  1.00000
## arrPP                 7.1928134 0.13813418 0.13808749 2711.410543  1.00000
## arrST                 5.4475191 0.14899952 0.14895106 1336.680936  1.00000
## temp22C               0.8388660 0.09669082 0.09650325   75.268741  1.00000
## temp25C               1.9172401 0.10839953 0.10818009  312.822681  1.00000
## frailty(Birth)               NA         NA         NA  210.170466 18.58978
## arrCH:temp22C         0.5818853 0.13799636 0.13794690   17.780317  1.00000
## arrCH:temp25C         0.8066706 0.18632884 0.18618084   18.742705  1.00000
## arrPP:temp25C        -0.2953072 0.13847978 0.13844407    4.547527  1.00000
## arrAR:isogenDM1015    7.3698781 0.12534376 0.12529409 3457.125816  1.00000
## arrPP:isogenDM1020    0.7436113 0.13947408 0.13929325   28.425291  1.00000
## arrPP:isogenDM1038    0.2378456 0.10884261 0.10881967    4.775208  1.00000
## arrPP:isogenDM1049    1.1125611 0.14478513 0.14468954   59.047262  1.00000
## arrAR:isogenDM1050    7.2952161 0.14666110 0.14662142 2474.266377  1.00000
## arrAR:isogenDM1056    7.7054120 0.12678196 0.12676409 3693.829294  1.00000
## arrST:isogenJR138     1.9049259 0.11647525 0.11643453  267.478485  1.00000
## arrST:isogenJR209     1.1068273 0.09948077 0.09939619  123.788835  1.00000
## arrCH:isogenJR272    -0.8228138 0.12885155 0.12883270   40.777803  1.00000
## arrCH:isogenJR32     -0.6128557 0.14111652 0.14110217   18.860828  1.00000
## arrST:isogenJR91      2.7198026 0.13570570 0.13567046  401.678480  1.00000
## arrAR:isogenKB635     7.6905795 0.14615418 0.14612588 2768.825791  1.00000
## arrAR:isogenKB652     6.3945462 0.11331362 0.11328139 3184.602353  1.00000
##                                 p       hazard
```

```
## sexmale             1.607211e-35    1.5647474
## arrCH               0.000000e+00 1460.4360176
## arrCU              9.615795e-300 1176.2238427
## arrPP               0.000000e+00 1329.8393276
## arrST              1.207882e-292  232.1814227
## temp22C             4.108124e-18    2.3137416
## temp25C             5.300679e-70    6.8021591
## frailty(Birth)      1.923083e-34           NA
## arrCH:temp22C       2.479349e-05    1.7894088
## arrCH:temp25C       1.495945e-05    2.2404362
## arrPP:temp25C       3.296634e-02    0.7443029
## arrAR:isogenDM1015  0.000000e+00 1587.4403194
## arrPP:isogenDM1020  9.738478e-08    2.1035182
## arrPP:isogenDM1038  2.887237e-02    1.2685133
## arrPP:isogenDM1049  1.539300e-14    3.0421395
## arrAR:isogenDM1050  0.000000e+00 1473.2352583
## arrAR:isogenDM1056  0.000000e+00 2220.3320041
## arrST:isogenJR138   4.022276e-60    6.7189098
## arrST:isogenJR209   9.370338e-29    3.0247465
## arrCH:isogenJR272   1.705594e-10    0.4391941
## arrCH:isogenJR32    1.406106e-05    0.5418014
## arrST:isogenJR91    2.374393e-89   15.1773259
## arrAR:isogenKB635   0.000000e+00 2187.6420387
## arrAR:isogenKB652   0.000000e+00  598.5716372
```

Next, run the model and extract components for data excluding one day old deaths.

```r
aging_sex_arr_temp_no1 <- coxph(Surv(Age.at.death, censrec) ~
                                  sex + arr + temp + arr*temp + arr:isogen +
                                  frailty(Birth), aging_temps_no1)
aging_sex_arr_temp_no1_coeff <- data.frame(summary(aging_sex_arr_temp_no1)$coefficients)
subset(aging_sex_arr_temp_no1_coeff, p<0.05)
```

```
##                           coef   se.coef.        se2      Chisq      DF
## sexmale              0.4491105 0.03613972 0.03613342 154.431840  1.0000
## temp22C              0.8560148 0.09710355 0.09691133  77.712758  1.0000
## temp25C              1.9194597 0.10910592 0.10887929 309.500477  1.0000
## frailty(Birth)              NA         NA         NA 209.199580 18.5828
## arrCH:temp22C        0.5737571 0.13842709 0.13837702  17.179640  1.0000
## arrCH:temp25C        0.8147593 0.18749172 0.18733799  18.884020  1.0000
## arrPP:temp25C       -0.2930652 0.13943788 0.13940086   4.417401  1.0000
## arrPP:isogenDM1020   0.7506895 0.13966150 0.13947681  28.891310  1.0000
## arrPP:isogenDM1049   1.1124801 0.14577003 0.14567239  58.243573  1.0000
## arrST:isogenJR138    0.2469554 0.11724745 0.11720598   4.436395  1.0000
## arrST:isogenJR209   -0.5345272 0.09975699 0.09966957  28.711309  1.0000
## arrCH:isogenJR272   -0.8196453 0.12907439 0.12905458  40.324753  1.0000
## arrCH:isogenJR32    -0.6211977 0.14189041 0.14187551  19.166979  1.0000
## arrST:isogenJR91     1.0747256 0.13657688 0.13654054  61.921415  1.0000
## arrTL:isogenMA1959  -1.0299102 0.12740679 0.12739519  65.345182  1.0000
## arrTL:isogenSCI12-2 -0.9046275 0.14093194 0.14091795  41.202227  1.0000
## arrTL:isogenSPE123_2-3 -0.4298884 0.14980171 0.14978101   8.235271  1.0000
## arrTL:isogenSPE123_5-1 -0.7080110 0.14331543 0.14329465  24.405862  1.0000
##                                   p
## sexmale                1.863612e-35
## temp22C                1.191691e-18
```

```
## temp25C                2.805709e-69
## frailty(Birth)         2.982266e-34
## arrCH:temp22C          3.400616e-05
## arrCH:temp25C          1.389115e-05
## arrPP:temp25C          3.557418e-02
## arrPP:isogenDM1020     7.655560e-08
## arrPP:isogenDM1049     2.315923e-14
## arrST:isogenJR138      3.518046e-02
## arrST:isogenJR209      8.401169e-08
## arrCH:isogenJR272      2.150667e-10
## arrCH:isogenJR32       1.197674e-05
## arrST:isogenJR91       3.574410e-15
## arrTL:isogenMA1959     6.286341e-16
## arrTL:isogenSCI12-2    1.372660e-10
## arrTL:isogenSPE123_2-3 4.108405e-03
## arrTL:isogenSPE123_5-1 7.803089e-07
```

In order to directly measure arrangement effects within each sex, need to analyze data from males and females separately. When doing so, need to use `coxme()` in order to estimate arrangement effects so that we can model isogen (strain) as a random effect. To do this, we will use TL as a baseline (reference) for arrangements because it has the longest lifespan (smallest hazard) on average (see above). Here is the analysis of the entire data.

```
aging_sex_arr_temp_male <- coxme(Surv(Age.at.death, censrec) ~
                                   arr_factor + temp + arr_factor*temp +
                                   (1 | arr_factor/isogen) + (1|Birth),
                                 subset(aging_temps, sex=="male"))
aging_sex_arr_temp_female <- coxme(Surv(Age.at.death, censrec) ~
                                     arr_factor + temp + arr_factor*temp +
                                     (1 | arr_factor/isogen) + (1|Birth),
                                   subset(aging_temps, sex=="female"))

aging_sex_arr_temp_coefficients <- rbind(
    cbind(sex="male", data.frame(summary(aging_sex_arr_temp_male)$coefficients)),
    cbind(sex="female", data.frame(summary(aging_sex_arr_temp_female)$coefficients))
)

subset(aging_sex_arr_temp_coefficients, p<0.05)
```

```
##                          sex       coef exp.coef.  se.coef.     z            p
## arr_factorPP            male  0.6194080  1.857828 0.3032017  2.04 4.106327e-02
## temp22C                 male  0.9478848  2.580246 0.1674354  5.66 1.503202e-08
## temp25C                 male  1.9687820  7.161948 0.2015654  9.77 1.552937e-22
## arr_factorPP1         female  0.6915974  1.996903 0.3079233  2.25 2.470364e-02
## temp22C1              female  1.1291332  3.092974 0.1484442  7.61 2.817319e-14
## temp25C1              female  2.0562596  7.816677 0.1741231 11.81 3.497433e-32
## arr_factorCH:temp22C1 female  0.4720457  1.603271 0.2007465  2.35 1.870032e-02
## arr_factorPP:temp22C1 female -0.4266805  0.652672 0.1862846 -2.29 2.199373e-02
## arr_factorCH:temp25C1 female  0.9285843  2.530924 0.2702834  3.44 5.912547e-04
```

Next, we will analyze the data excluding 1 day old deaths.

```
aging_sex_arr_temp_male_no1 <- coxme(Surv(Age.at.death, censrec) ~
                                       arr_factor + temp + arr_factor*temp +
                                       (1 | arr_factor/isogen) + (1|Birth),
                                     subset(aging_temps_no1, sex=="male"))
```

```r
aging_sex_arr_temp_female_no1 <- coxme(Surv(Age.at.death, censrec) ~
                                        arr_factor + temp + arr_factor*temp +
                                        (1 | arr_factor/isogen) + (1|Birth),
                                      subset(aging_temps_no1, sex=="female"))
aging_sex_arr_temp_no1_coefficients <- rbind(
    cbind(sex="male", data.frame(summary(aging_sex_arr_temp_male_no1)$coefficients)),
    cbind(sex="female", data.frame(summary(aging_sex_arr_temp_female_no1)$coefficients))
)

subset(aging_sex_arr_temp_no1_coefficients, p<0.05)
```

```
##                             sex       coef exp.coef.  se.coef.     z            p
## arr_factorPP               male  0.6127300 1.8454627 0.3043299  2.01 4.407523e-02
## temp22C                    male  0.9537003 2.5952953 0.1676750  5.69 1.286932e-08
## temp25C                    male  1.9473161 7.0098485 0.2035933  9.56 1.124870e-21
## arr_factorPP1            female  0.6758505 1.9657041 0.3081069  2.19 2.826715e-02
## temp22C1                 female  1.1314208 3.1000578 0.1486472  7.61 2.710381e-14
## temp25C1                 female  2.0640940 7.8781571 0.1745273 11.83 2.838550e-32
## arr_factorCH:temp22C1 female  0.4759955 1.6096158 0.2012601  2.37 1.802634e-02
## arr_factorPP:temp22C1 female -0.4117399 0.6624966 0.1866345 -2.21 2.737497e-02
## arr_factorCH:temp25C1 female  0.9189440 2.5066420 0.2722561  3.38 7.373717e-04
```

Repeat with PP as a reference because it has a significant effect on survival.

```r
aging_temps_no1$arr_PP <- factor(aging_temps_no1$arr,
  levels = c("PP", "AR", "CH", "CU", "ST", "TL"))

aging_sex_arr_temp_male_PP_no1 <- coxme(Surv(Age.at.death, censrec) ~
                                   arr_PP + temp + arr_PP*temp +
                                   (1 | arr_PP/isogen) + (1|Birth),
                                 subset(aging_temps_no1, sex=="male"))
aging_sex_arr_temp_female_PP_no1 <- coxme(Surv(Age.at.death, censrec) ~
                                     arr_PP + temp + arr_PP*temp +
                                     (1 | arr_PP/isogen) + (1|Birth),
                                   subset(aging_temps_no1, sex=="female"))

aging_sex_arr_temp_coefficients_PP_no1 <- rbind(
    cbind(sex="male", data.frame(summary(aging_sex_arr_temp_male_PP_no1)$coefficients)),
    cbind(sex="female", data.frame(summary(aging_sex_arr_temp_female_PP_no1)$coefficients))
)

subset(aging_sex_arr_temp_coefficients_PP_no1, p<0.05)
```

```
##                       sex       coef exp.coef.  se.coef.     z            p
## arr_PPTL             male -0.6127300 0.5418695 0.3043299 -2.01 4.407523e-02
## temp22C              male  0.7002549 2.0142661 0.1576677  4.44 8.940328e-06
## temp25C              male  1.5812372 4.8609660 0.1568812 10.08 6.827915e-24
## arr_PPCH:temp22C     male  0.4609780 1.5856240 0.2168350  2.13 3.350832e-02
## arr_PPCH:temp25C     male  0.9170300 2.5018489 0.2678647  3.42 6.182439e-04
## arr_PPCU:temp25C     male  0.8256011 2.2832529 0.3498890  2.36 1.829423e-02
## arr_PPCH1          female -0.6502724 0.5219036 0.3099581 -2.10 3.591079e-02
## arr_PPTL1          female -0.6758505 0.5087236 0.3081069 -2.19 2.826715e-02
## temp22C1           female  0.7196809 2.0537778 0.1285323  5.60 2.153180e-08
## temp25C1           female  1.8033371 6.0698693 0.1380983 13.06 5.694331e-39
## arr_PPAR:temp22C1 female  0.3489374 1.4175604 0.1706790  2.04 4.091328e-02
```

```
## arr_PPCH:temp22C1 female  0.8877354 2.4296212 0.1935660  4.59 4.513530e-06
## arr_PPCU:temp22C1 female  0.6259515 1.8700245 0.3138772  1.99 4.612404e-02
## arr_PPTL:temp22C1 female  0.4117399 1.5094417 0.1866345  2.21 2.737497e-02
## arr_PPAR:temp25C1 female  0.4127839 1.5110185 0.1874606  2.20 2.766691e-02
## arr_PPCH:temp25C1 female  1.1797010 3.2534011 0.2550856  4.62 3.750944e-06
```

Now with 1 day deaths excluded.

```r
aging_temps$arr_PP <- factor(aging_temps$arr,
  levels = c("PP", "AR", "CH", "CU", "ST", "TL"))

aging_sex_arr_temp_male_PP <- coxme(Surv(Age.at.death, censrec) ~
                                  arr_PP + temp + arr_PP*temp +
                                  (1 | arr_PP/isogen) + (1|Birth),
                                subset(aging_temps, sex=="male"))
aging_sex_arr_temp_female_PP <- coxme(Surv(Age.at.death, censrec) ~
                                    arr_PP + temp + arr_PP*temp +
                                    (1 | arr_PP/isogen) + (1|Birth),
                                  subset(aging_temps, sex=="female"))

aging_sex_arr_temp_coefficients_PP <- rbind(
    cbind(sex="male", data.frame(summary(aging_sex_arr_temp_male_PP)$coefficients)),
    cbind(sex="female", data.frame(summary(aging_sex_arr_temp_female_PP)$coefficients))
)

subset(aging_sex_arr_temp_coefficients_PP, p<0.05)
```

```
##                      sex       coef exp.coef.  se.coef.     z            p
## arr_PPTL            male -0.6194080 0.5382630 0.3032017 -2.04 4.106327e-02
## temp22C             male  0.6835632 1.9809236 0.1570746  4.35 1.350017e-05
## temp25C             male  1.5742693 4.8272133 0.1559222 10.10 5.724418e-24
## arr_PPCH:temp22C    male  0.4705797 1.6009220 0.2163259  2.18 2.960555e-02
## arr_PPCH:temp25C    male  0.8999922 2.4595839 0.2669746  3.37 7.487463e-04
## arr_PPCU:temp25C    male  0.7990352 2.2233949 0.3492103  2.29 2.213052e-02
## arr_PPCH1         female -0.6569647 0.5184225 0.3096487 -2.12 3.386757e-02
## arr_PPTL1         female -0.6915974 0.5007755 0.3079233 -2.25 2.470364e-02
## temp22C1          female  0.7024527 2.0186980 0.1280865  5.48 4.153343e-08
## temp25C1          female  1.7978020 6.0363650 0.1371646 13.11 3.006534e-39
## arr_PPAR:temp22C1 female  0.3553526 1.4266836 0.1701690  2.09 3.677675e-02
## arr_PPCH:temp22C1 female  0.8987262 2.4564722 0.1928071  4.66 3.142631e-06
## arr_PPCU:temp22C1 female  0.6369629 1.8907299 0.3136844  2.03 4.229711e-02
## arr_PPTL:temp22C1 female  0.4266805 1.5321631 0.1862846  2.29 2.199373e-02
## arr_PPAR:temp25C1 female  0.4180866 1.5190522 0.1862109  2.25 2.475329e-02
## arr_PPCH:temp25C1 female  1.1870418 3.2773718 0.2526050  4.70 2.611803e-06
```

Key results:

- Hazard increases at 22° and 25° in both males and females, meaning lifespan is shorter with warmer temperatures.
- PP has increased hazard (shorter lifespan) in both males and females.
- Significant positive interaction between CH and temperature for females, meaning the effect of temperature on lifespan in females is greater for CH than other arrangements.
- Significant negative interaction between PP and 22° in females, which means females carrying the PP arrangement are less affected by the increase from 18° to 22°.

Lastly, we will use Gompertz model to test for the effect of temperature on both the rate and shape of the survival curve. We will perform this analysis within each sex and arrangement combination by contrasting across temperatures. To do this, we define functions to perform the analyses, and then call those functions on the complete data and the data excluding 1 day old deaths.

```
temp_shape_rate_comparisons <- function(agingDataArr){
    NULLmodel <- flexsurvreg(Surv(Age.at.death, censrec) ~ Birth + isogen,
                             anc = list(shape = ~ 1), data = agingDataArr,
                             dist = "gompertz")
    tempShape <- flexsurvreg(Surv(Age.at.death, censrec) ~ Birth + isogen,
                             anc = list(shape = ~ temp), data = agingDataArr,
                             dist = "gompertz")
    tempRate <- flexsurvreg(Surv(Age.at.death, censrec) ~ Birth + isogen + temp,
                            anc = list(shape = ~ 1), data = agingDataArr,
                            dist = "gompertz")
    tempShapeRate <- flexsurvreg(Surv(Age.at.death, censrec) ~ Birth + isogen + temp,
                                 anc = list(shape = ~ temp), data = agingDataArr,
                                 dist = "gompertz")

    return.df <- data.frame(
        factors = c("tempShapeNULL", "tempRateNULL", "tempRateShape", "tempShapeRate"),
        pval = c(
            pchisq(2 * (logLik(tempShape) - logLik(NULLmodel)),
                   df = 1, lower.tail = FALSE),
            pchisq(2 * (logLik(tempRate) - logLik(NULLmodel)),
                   df = 1, lower.tail = FALSE),
             # test if adding Rate to a model with Shape improves fit
            pchisq(2 * (logLik(tempShapeRate) - logLik(tempShape)),
                   df = 1, lower.tail = FALSE),
            # test if adding Shape to a model with Rate improves fit
            pchisq(2 * (logLik(tempShapeRate) - logLik(tempRate)),
                   df = 1, lower.tail = FALSE)
        ),
        significant = NA
    )

    return.df$significant <- ""
    return.df$significant[return.df$pval < 0.05] <- "*"
    return.df$significant[return.df$pval < 0.005] <- "**"
    return.df$significant[return.df$pval < 0.0005] <- "***"
    return.df$significant[return.df$pval < 0.00005] <- "****"
    return.df$significant[return.df$pval < 0.000005] <- "******"

    return.df
}
temp_shape_rate <- function(agingData){
    rbind(
        cbind(  arrangement="AR", sex="female",
            temp_shape_rate_comparisons(subset(agingData, arr=="AR" & sex=="female"))),
        cbind(  arrangement="CH", sex="female",
            temp_shape_rate_comparisons(subset(agingData, arr=="CH" & sex=="female"))),
        cbind(  arrangement="PP", sex="female",
            temp_shape_rate_comparisons(subset(agingData, arr=="PP" & sex=="female"))),
        cbind(  arrangement="ST", sex="female",
            temp_shape_rate_comparisons(subset(agingData, arr=="ST" & sex=="female"))),
        cbind(  arrangement="TL", sex="female",
```

```
            temp_shape_rate_comparisons(subset(agingData, arr=="TL" & sex=="female"))),
        cbind(  arrangement="AR", sex="male",
            temp_shape_rate_comparisons(subset(agingData, arr=="AR" & sex=="male"))),
        cbind(  arrangement="CH", sex="male",
            temp_shape_rate_comparisons(subset(agingData, arr=="CH" & sex=="male"))),
        cbind(  arrangement="PP", sex="male",
            temp_shape_rate_comparisons(subset(agingData, arr=="PP" & sex=="male"))),
        cbind(  arrangement="ST", sex="male",
            temp_shape_rate_comparisons(subset(agingData, arr=="ST" & sex=="male"))),
        cbind(  arrangement="TL", sex="male",
            temp_shape_rate_comparisons(subset(agingData, arr=="TL" & sex=="male")))
    )
}

temps_shape_rate <- temp_shape_rate(aging_temps)
temps_shape_rate_no1 <- temp_shape_rate(aging_temps_no1)
```

Taking subsets of the data allows us to examine how specific factors affect model fit. Doing this first for the effect of the rate parameter reveals that adding temperature as a rate parameter improves model fit for all arrangements in both males and females.

```
subset(temps_shape_rate, factors=="tempRateShape")
```

```
##    arrangement    sex       factors         pval significant
## 3           AR female tempRateShape 7.370260e-20      ******
## 7           CH female tempRateShape 4.053913e-11      ******
## 11          PP female tempRateShape 8.486493e-22      ******
## 15          ST female tempRateShape 2.041148e-24      ******
## 19          TL female tempRateShape 1.434645e-26      ******
## 23          AR   male tempRateShape 7.820247e-15      ******
## 27          CH   male tempRateShape 1.099971e-06      ******
## 31          PP   male tempRateShape 1.622563e-12      ******
## 35          ST   male tempRateShape 6.326862e-10      ******
## 39          TL   male tempRateShape 5.340940e-11      ******
```

A similar approach can assess if adding temperature as a shape parameter affect model fit.

```
subset(temps_shape_rate, factors=="tempShapeRate")
```

```
##    arrangement    sex       factors         pval significant
## 4           AR female tempShapeRate 6.394103e-04          **
## 8           CH female tempShapeRate 2.993109e-01
## 12          PP female tempShapeRate 8.861751e-06        ****
## 16          ST female tempShapeRate 5.612851e-04          **
## 20          TL female tempShapeRate 3.861638e-03          **
## 24          AR   male tempShapeRate 1.244698e-01
## 28          CH   male tempShapeRate 1.188539e-01
## 32          PP   male tempShapeRate 9.355920e-03           *
## 36          ST   male tempShapeRate 8.506845e-01
## 40          TL   male tempShapeRate 2.537435e-03          **
```

This reveals adding temperature as a shape parameter only improves model fit for some arrangements in some sexes. Interpreting these results, we can see, for example, that temperature affects both the shape and rate of the survival curve in females carrying the AR or ST arrangements, but temperature only affects the rate parameter in AR and ST males. In contrast, temperature affects both the shape and rate parameters in PP and TL flies of both sexes. In both males and females carrying the CH arrangement, temperature only affects the rate parameter and not shape.

What is the effect of temperature on the rate and shape parameters? Run models where the is a significant effect of temperature on the shape parameter. Write a function to analyze model, and then call the model for each data subset with a significant shape effect.

```
gompertz.shape <- function(agingData, ARR, SEX){
  flexsurvreg(Surv(Age.at.death, censrec) ~ Birth + isogen + temp,
              anc = list(shape = ~ temp),
              data = subset(agingData, arr==ARR & sex==SEX),
              dist = "gompertz")
}
gompertz_AR_female <- gompertz.shape(aging_temps, "AR", "female")
gompertz_ST_female <- gompertz.shape(aging_temps, "ST", "female")
gompertz_PP_female <- gompertz.shape(aging_temps, "PP", "female")
gompertz_PP_male   <- gompertz.shape(aging_temps, "PP", "male")
gompertz_TL_female <- gompertz.shape(aging_temps, "TL", "female")
gompertz_TL_male   <- gompertz.shape(aging_temps, "TL", "male")
```

View the temperature effects in in the model outputs. Write a function to extract that from the model outputs

```
temp.effects <- function(gomp_arr_sex){
 as.data.frame(gomp_arr_sex$res)[
    grepl(paste(c("temp","shape","rate"), collapse = "|") ,
    rownames(as.data.frame(gomp_arr_sex$res))),
  ]
}
```

```
temp.effects(gompertz_AR_female)
```

```
##                         est          L95%          U95%          se
## shape           0.034403827  0.029127906  0.039679747 0.002691846
## rate            0.004411752  0.002318477  0.008394976 0.001448165
## temp22C         1.635094895  1.175969829  2.094219961 0.234251787
## temp25C         2.251149080  1.753102766  2.749195393 0.254109931
## shape(temp22C) -0.013640026 -0.021618072 -0.005661981 0.004070506
## shape(temp25C) -0.004168444 -0.016131177  0.007794290 0.006103548
```

The shape and rate parameters for AR females are both positive. The effect of increasing temperature on the rate parameter is positive, meaning that the initial mortality rate increases as temperature increases. The effect of increasing temperature on the shape parameter is negative, which means the rate of aging is slower at higher temperatures.

```
temp.effects(gompertz_ST_female)
```

```
##                         est          L95%          U95%          se
## shape           0.036513303  0.029089217  0.043937389 0.003787869
## rate            0.004971228  0.002380529  0.010381352 0.001867654
## temp22C         1.576312198  0.943137635  2.209486760 0.323054182
## temp25C         3.027036915  2.410424069  3.643649762 0.314604172
## shape(temp22C) -0.011434707 -0.021306279 -0.001563136 0.005036609
## shape(temp25C) -0.021234244 -0.034107613 -0.008360874 0.006568166
```

The shape and rate parameters for ST females are both positive. The effect of increasing temperature on the rate parameter is positive, meaning that the initial mortality rate increases as temperature increases. The effect of increasing temperature on the shape parameter is negative, which means the rate of aging is slower at higher temperatures.

```r
temp.effects(gompertz_PP_female)
```

```
##                          est          L95%          U95%          se
## shape            0.037872893  0.031712877  0.044032908 0.003142923
## rate             0.004521578  0.002483208  0.008233168 0.001382590
## temp22C          1.137568792  0.630005773  1.645131811 0.258965483
## temp25C          2.417732119  1.920405515  2.915058722 0.253742726
## shape(temp22C) -0.015151162 -0.025030491 -0.005271834 0.005040566
## shape(temp25C) -0.024814196 -0.037090777 -0.012537616 0.006263676
```

The shape and rate parameters for PP females are both positive. The effect of increasing temperature on the rate parameter is positive, meaning that the initial mortality rate increases as temperature increases. The effect of increasing temperature on the shape parameter is negative, which means the rate of aging is slower at higher temperatures.

```r
temp.effects(gompertz_PP_male)
```

```
##                          est          L95%          U95%          se
## shape            0.041086210  0.032041551  0.050130869 0.004614707
## rate             0.006760992  0.002486286  0.018385260 0.003450859
## temp22C          0.978544246  0.408646297  1.548442194 0.290769602
## temp25C          2.002714082  1.424431753  2.580996410 0.295047426
## shape(temp22C) -0.015106806 -0.028489875 -0.001723738 0.006828222
## shape(temp25C) -0.017767505 -0.034805457 -0.000729552 0.008692993
```

The shape and rate parameters for PP males are both positive. The effect of increasing temperature on the rate parameter is positive, meaning that the initial mortality rate increases as temperature increases. The effect of increasing temperature on the shape parameter is negative, which means the rate of aging is slower at higher temperatures.

```r
temp.effects(gompertz_TL_female)
```

```
##                          est          L95%          U95%           se
## shape            0.046892264  0.0390907505  0.054693777 0.0039804370
## rate             0.001541628  0.0007681467  0.003093961 0.0005479277
## temp22C          2.190421995  1.5582986682  2.822545321 0.3225178277
## temp25C          3.299088428  2.6502373697  3.947939487 0.3310525416
## shape(temp22C) -0.014352986 -0.0259213478 -0.002784623 0.0059023341
## shape(temp25C) -0.017631516 -0.0330110740 -0.002251957 0.0078468576
```

The shape and rate parameters for TL females are both positive. The effect of increasing temperature on the rate parameter is positive, meaning that the initial mortality rate increases as temperature increases. The effect of increasing temperature on the shape parameter is negative, which means the rate of aging is slower at higher temperatures.

```r
temp.effects(gompertz_TL_male)
```

```
##                          est          L95%          U95%           se
## shape            0.048484259  0.0385470101  0.058421507 0.0050701179
## rate             0.001864376  0.0007517471  0.004623757 0.0008639841
## temp22C          1.789853373  1.0599630840  2.519743662 0.3723998474
## temp25C          2.611769007  1.8267003670  3.396837647 0.4005525847
## shape(temp22C) -0.020833753 -0.0357054800 -0.005962026 0.0075877552
## shape(temp25C) -0.023045619 -0.0473993840  0.001308145 0.0124256184
```

The shape and rate parameters for TL males are both positive. The effect of increasing temperature on the rate parameter is positive, meaning that the initial mortality rate increases as temperature increases. The effect of increasing temperature on the shape parameter is negative, which means the rate of aging is slower at higher temperatures.

#### Export average lifespan for each strain

We will export the average lifespan for males and feamles from each strain so that we can use those data in analyses comparing lifespan with other life history traits. First, we will use the median value for each strain. To do this, we will write a function that calculates median lifespan for each sex from all strains. Then we will call that function for the data from each temperature. Lastly, we will export the median values to file.

```
sex.median <- function(aging){
    strains <- levels(factor(aging$Strain))
    female.df <- data.frame(
        strain = strains,
        median = NA,
        obs = NA
    )
    male.df <- female.df
    for(i in 1:length(strains)){
        female.df$median[i] <- median(subset(aging,
                                             Strain==strains[i] &
                                               sex=="female")$Age.at.death)
        female.df$obs[i] <- length(subset(aging,
                                          Strain==strains[i] &
                                            sex=="female")[,1])
    }
    female.df$sex <- "female"
    for(i in 1:length(strains)){
        male.df$median[i] <- median(subset(aging,
                                           Strain==strains[i] &
                                             sex=="male")$Age.at.death)
        male.df$obs[i] <- length(subset(aging,
                                        Strain==strains[i] &
                                          sex=="male")[,1])
    }
    male.df$sex <- "male"
    rbind(female.df, male.df)
}
# exclude strain 6 from 18C because not enough data
sex_median_18 <- sex.median(subset(aging_18, Strain!=6))
sex_median_22 <- sex.median(aging_22)
sex_median_25 <- sex.median(aging_25)

write.table(sex_median_18, "sex_median_18.tsv",
            quote=FALSE, sep="\t", row.names=FALSE)
write.table(sex_median_22, "sex_median_22.tsv",
            quote=FALSE, sep="\t", row.names=FALSE)
write.table(sex_median_25, "sex_median_25.tsv",
            quote=FALSE, sep="\t", row.names=FALSE)
```

Second, we will use the intercept from a linear model for each strain. This allows us to calculate the mean value, while also modeling batch (emergence date) for each fly as a random effect. We will write a function to calculate the intercept, only including strains with measurements from at least 10 flies.

```r
sex.intercept <- function(datas){
    strains <- levels(factor(datas$Strain))
    female.df <- data.frame(
        strain = strains,
        effect = NA,
        obs = NA
    )
    male.df <- female.df

    for(i in 1:length(strains)){
        # only consider if have >9 samples
      if(length(subset(datas, Strain==strains[i] & sex=="female")[,1]) > 9){
            if(length(levels(factor(subset(datas, sex=="female" &
                                               Strain==strains[i])$Birth))) > 1){
                female.df$effect[i] <- summary(lmer(Age.at.death ~ 1 + (1|Birth),
                                                     data=subset(datas,
                                                                 Strain==strains[i] &
                                                                   sex=="female"))
                                                 )$coefficients[1]
            }else{
                female.df$effect[i] <- summary(lm(Age.at.death ~ 1,
                                                   data=subset(datas,
                                                               Strain==strains[i] &
                                                                 sex=="female"))
                                                 )$coefficients[1]
            }
        } else{
            female.df$effect[i] <- NA
        }
        female.df$obs[i] <- length(subset(datas,
                                          Strain==strains[i] & sex=="female")[,1])
    }
    female.df$sex <- "female"

    for(i in 1:length(strains)){
        # only consider if have >9 samples
      if(length(subset(datas, Strain==strains[i] & sex=="male")[,1]) > 9){
            if(length(levels(factor(subset(datas, sex=="male" &
                                               Strain==strains[i])$Birth))) > 1){
                male.df$effect[i] <- summary(lmer(Age.at.death ~ 1 + (1|Birth),
                                                   data=subset(datas,
                                                               Strain==strains[i] &
                                                                 sex=="male"))
                                               )$coefficients[1]
            } else{
                male.df$effect[i] <- summary(lm(Age.at.death ~ 1,
                                                 data=subset(datas,
                                                             Strain==strains[i] &
                                                               sex=="male"))
                                               )$coefficients[1]
            }
        } else{
            male.df$effect[i] <- NA
```

```
        }
        male.df$obs[i] <- length(subset(datas,
                                        Strain==strains[i] & sex=="male")[,1])
    }
    male.df$sex <- "male"

    rbind(female.df, male.df)
}
sex_intercept_18 <- sex.intercept(aging_18)
sex_intercept_22 <- sex.intercept(aging_22)
sex_intercept_25 <- sex.intercept(aging_25)

write.table(sex_intercept_18, "sex_intercept_18.tsv",
            quote=FALSE, sep="\t", row.names=FALSE)
write.table(sex_intercept_22, "sex_intercept_22.tsv",
            quote=FALSE, sep="\t", row.names=FALSE)
write.table(sex_intercept_25, "sex_intercept_25.tsv",
            quote=FALSE, sep="\t", row.names=FALSE)
```

#### Compare 22C lifespans across strains from the two trials

Two separate trials were performed measuring lifespan at 22°C. We will compare the results of those two trials to test if the lifespan measurements are reproducible. Our specific focus is on the average lifespan measured for each strain. We will use two different measures of average lifespan for each strain: median and mean. To do the comparison, we will first load the data from the other experiment.

```
sex_aging_winterCens <- read.delim("sex_median_winterCens.tsv")
sex_agingInt_winterCens <- read.delim("sex_intercept_winterCens.tsv")
```

Create a function to compare the two trials at 22°C. The function generates a single dataframe for data from the two trials, and it also creates a plot showing the relationships between average lifespan for the strains between the two trials.

```
lifespan.corr <- function(aging1, aging2, strains){
    arrcols <- c( "AR"="blue", "CH"="orange", "CU"="black",
                  "PP"="red", "ST"="darkgreen", "TL"="purple")
    female.df <- data.frame(
        strain = intersect(subset(aging1, sex=="female")$strain,
                           subset(aging2, sex=="female")$strain),
        sex = "female",
        aging1 = NA,
        aging2 = NA
    )
    female.df$aging1 <- subset(aging1, sex=="female")[
      match(female.df$strain, subset(aging1, sex=="female")$strain), 2]
    female.df$aging2 <- subset(aging2, sex=="female")[
      match(female.df$strain, subset(aging2, sex=="female")$strain), 2]
    female.df$arr <- strains$arrangement[
      match(female.df$strain, strains$ID)]

    male.df <- data.frame(
        strain = intersect(
          subset(aging1, sex=="male")$strain,
          subset(aging2, sex=="male")$strain),
```

```
        sex = "male",
        aging1 = NA,
        aging2 = NA
    )
    male.df$aging1 <- subset(aging1, sex=="male")[
      match(male.df$strain, subset(aging1, sex=="male")$strain), 2]
    male.df$aging2 <- subset(aging2, sex=="male")[
      match(male.df$strain, subset(aging2, sex=="male")$strain), 2]
    male.df$arr <- strains$arrangement[
      match(male.df$strain, strains$ID)]

    print(
        ggplot(na.omit(rbind(female.df, male.df)),
               aes(x=aging1, y=aging2, color=arr)) +
        geom_point() +
        scale_color_manual(values=arrcols, name="") +
        facet_wrap(~sex, ncol=2) +
        scale_x_continuous("avg lifespan (days), trial 1") +
        scale_y_continuous("avg lifespan (days), trial 2") +
        theme_bw()
    )
    na.omit(rbind(female.df, male.df))
}
```

Call the function for the median lifespan.

```
sex_median_corr_22 <- lifespan.corr(sex_aging_winterCens,
                                    sex_median_22, strainIDs)
```

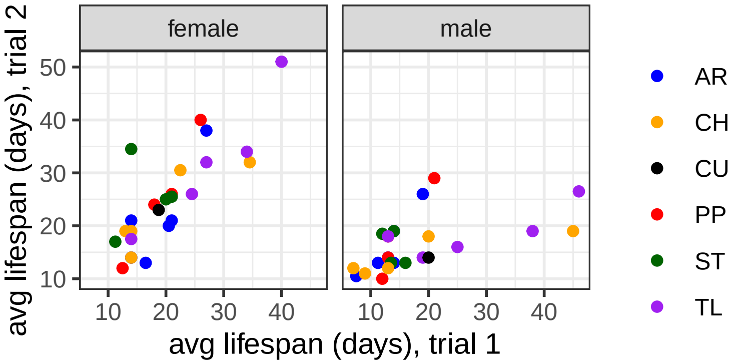


```
sex_intercept_corr_22 <- lifespan.corr(sex_agingInt_winterCens,
                                       sex_intercept_22, strainIDs)
```

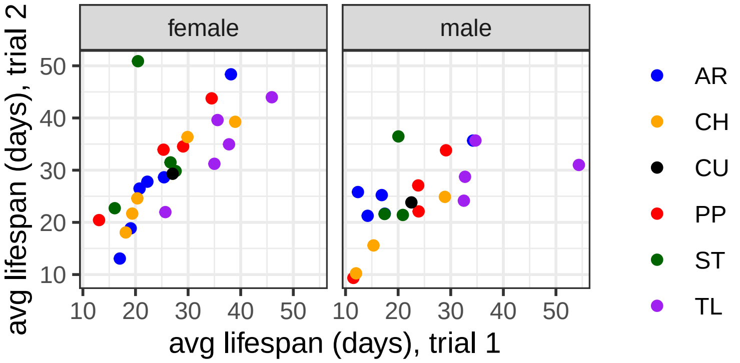

Make a version of the figure for publication.

```
arrcols <- c( "AR"="blue", "CH"="orange", "CU"="black",
                   "PP"="red", "ST"="darkgreen", "TL"="purple")
ggplot(sex_intercept_corr_22, aes(x=aging1, y=aging2, color=arr)) +
  geom_point() +
  scale_color_manual(values=arrcols, name="") +
  facet_wrap(~sex, nrow=2) +
  scale_x_continuous("avg lifespan (days), trial 1") +
  scale_y_continuous("avg lifespan (days), trial 2") +
  theme_bw()
```

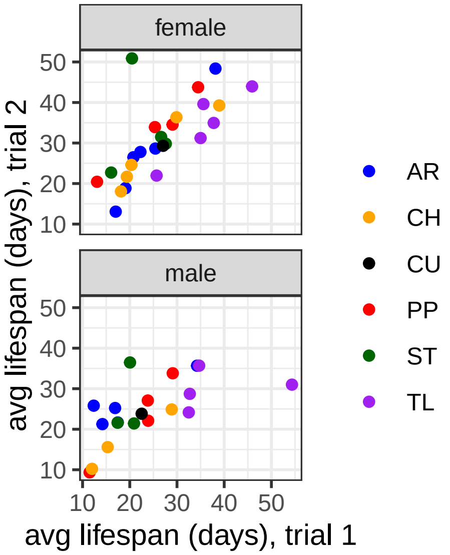


In the graphs above, each point represents an average lifespan measurement for a strain. We have colored the points (strains) by the arrangement they carry. Both graphs show that lifespan measurements across the two trials are similar between strains.

To test if the average lifespan across trials is significantly correlated, we construct a linear model with average lifespan in trial 2 as the response and the following predictors: average lifespan in trial 1, sex of the flies, and arrangement the strain carries. This allows us to consider arrangement and sex effects as co-variates in the analysis of strain-level correlations between trials. If there is a significant effect of trial 1 average lifespan on trial 2 lifespan, then the trials are significantly correlated.

We first run the model on the median lifespans.

```
summary(lm(aging2 ~ aging1 + sex + arr, data= sex_median_corr_22))
```

```
##
## Call:
## lm(formula = aging2 ~ aging1 + sex + arr, data = sex_median_corr_22)
##
## Residuals:
##      Min       1Q   Median       3Q      Max
## -10.7208  -3.9323  -0.2388   2.6352  15.2178
##
## Coefficients:
##             Estimate Std. Error t value Pr(>|t|)
## (Intercept) 13.21699    2.73565   4.831 1.84e-05 ***
## aging1       0.56238    0.11171   5.034 9.53e-06 ***
## sexmale     -7.32343    1.73050  -4.232 0.000123 ***
```

```
## arrCH       -1.70300    2.60774  -0.653 0.517278
## arrCU       -1.95142    4.62270  -0.422 0.675077
## arrPP        2.47409    2.75593   0.898 0.374446
## arrST        2.57348    2.75458   0.934 0.355514
## arrTL        0.06992    2.91362   0.024 0.980969
## ---
## Signif. codes:  0 '***' 0.001 '**' 0.01 '*' 0.05 '.' 0.1 ' ' 1
##
## Residual standard error: 6.032 on 42 degrees of freedom
## Multiple R-squared:  0.5924, Adjusted R-squared:  0.5244
## F-statistic: 8.719 on 7 and 42 DF,  p-value: 1.418e-06
```

Next, we run the analysis on the mean lifespans.

```
summary(lm(aging2 ~ aging1 + sex + arr, data= sex_intercept_corr_22))
```

```
##
## Call:
## lm(formula = aging2 ~ aging1 + sex + arr, data = sex_intercept_corr_22)
##
## Residuals:
##      Min       1Q   Median       3Q      Max
## -13.7079  -2.8410   0.1616   2.6537  19.3724
##
## Coefficients:
##             Estimate Std. Error t value Pr(>|t|)
## (Intercept)  10.2242     3.6744   2.783  0.00844 **
## aging1        0.8360     0.1320   6.335 2.21e-07 ***
## sexmale      -3.6985     1.9205  -1.926  0.06184 .
## arrCH        -4.1190     2.9035  -1.419  0.16437
## arrCU        -2.5299     4.8126  -0.526  0.60225
## arrPP        -0.1279     2.9137  -0.044  0.96523
## arrST         4.1758     3.0149   1.385  0.17434
## arrTL        -7.2621     3.4677  -2.094  0.04315 *
## ---
## Signif. codes:  0 '***' 0.001 '**' 0.01 '*' 0.05 '.' 0.1 ' ' 1
##
## Residual standard error: 6.234 on 37 degrees of freedom
## Multiple R-squared:  0.6234, Adjusted R-squared:  0.5521
## F-statistic: 8.749 on 7 and 37 DF,  p-value: 2.624e-06
```

In both cases, there is a significant correlation between the average lifespan measured in trials 1 and 2 ($p < 10^{-6}$). There is also a significant effect of sex, demonstrating that sex differences in lifespan at 22°C are shared across trials. We therefore conclude that the strain and sex differences in lifespan that we detect are reproducible.

#### Make plots of the data and survival models

The analysis above revealed very similar results when we exclude 1 day old death and when we include all data. Because of these similarities, we will include all data in our plots.

We will first use autoplot to graph survivorship (Kaplan-Meier) curves. The graphing functions will ignore any variable other than the focal effects in order to produce clear graphs. Start with a plot comparing males and females at each temperature. To do so, write a function that makes the plots for each temperature, and then call that function using data from each temperature.

```r
aging.sex.temp.plot <- function(aging){
    autoplot(survfit(Surv(Age.at.death, censrec) ~ sex, data=aging),
             ylab = "% surivival", xlab="time (days)") +
        scale_color_manual(values = c("female" = "red", "male" = "blue")) +
        scale_x_continuous(limits=c(0,140)) +
        theme_bw() +
        theme(legend.position="none")
}

plot_grid(
    ggdraw(aging.sex.temp.plot(aging_18)) +
        draw_label(label="female", x=0.3, y=0.5, colour="red") +
        draw_label(label="male", x=0.3, y=0.4, colour="blue"),
    aging.sex.temp.plot(aging_22),
    aging.sex.temp.plot(aging_25),
    labels = c('18C', '22C', '25C'),
    label_x=0.8, label_y=0.9, label_size = 12, nrow=3
)
```

```
## Warning: `aes_string()` was deprecated in ggplot2 3.0.0.
## i Please use tidy evaluation idioms with `aes()`.
## i See also `vignette("ggplot2-in-packages")` for more information.
## i The deprecated feature was likely used in the ggfortify package.
##   Please report the issue at <https://github.com/sinhrks/ggfortify/issues>.
## This warning is displayed once every 8 hours.
## Call `lifecycle::last_lifecycle_warnings()` to see where this warning was
## generated.
```

```
## Warning: Removed 1 row containing missing values or values outside the scale range
## (`geom_step()`).
```

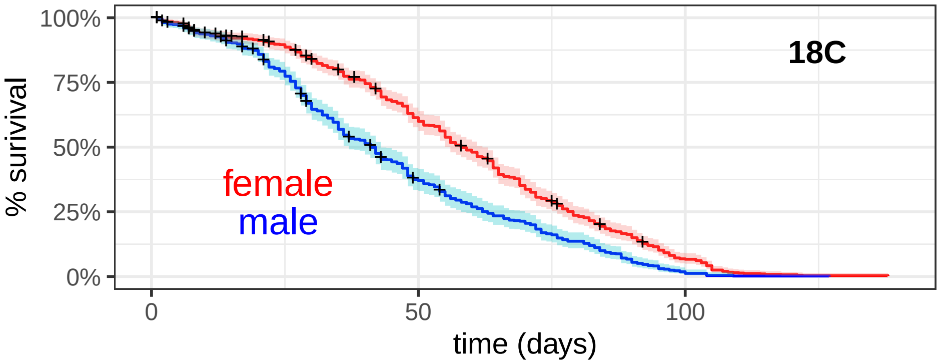


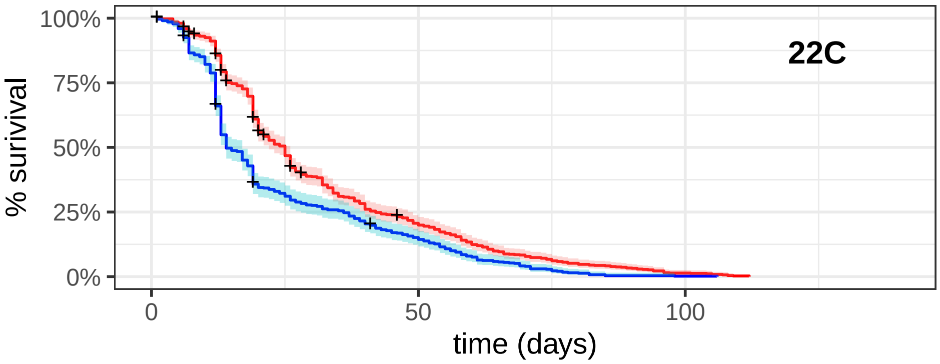


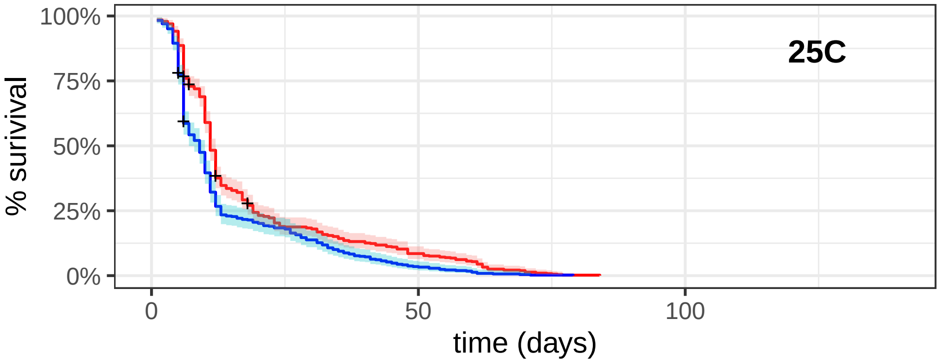


Next, we will create separate survivorship curves for males and females with each arrangement. This will allow us to compare males vs females within arrangements at each temperature.

```
ggsurvplot_facet(survfit(Surv(Age.at.death, censrec) ~ sex + arr,
                         data=aging_temps), aging_temps,
                 facet.by = c("arr","temp"), short.panel.labs=TRUE,
                 xlab="lifespan (days)", legend.title="") +
    scale_color_manual(values =
                         c("female" = "magenta", "male" = "cornflowerblue")) +
  scale_y_continuous(breaks=c(0,0.5,1))
```

```
## Warning: Using `size` aesthetic for lines was deprecated in ggplot2 3.4.0.
## i Please use `linewidth` instead.
## i The deprecated feature was likely used in the ggpubr package.
##   Please report the issue at <https://github.com/kassambara/ggpubr/issues>.
## This warning is displayed once every 8 hours.
## Call `lifecycle::last_lifecycle_warnings()` to see where this warning was
## generated.
```

```
## Warning: ggtheme is not a valid theme.
## Please use `theme()` to construct themes.
```

```
## Scale for y is already present.
## Adding another scale for y, which will replace the existing scale.
## Ignoring unknown labels:
## * fill : ""
## * linetype : "1"
```

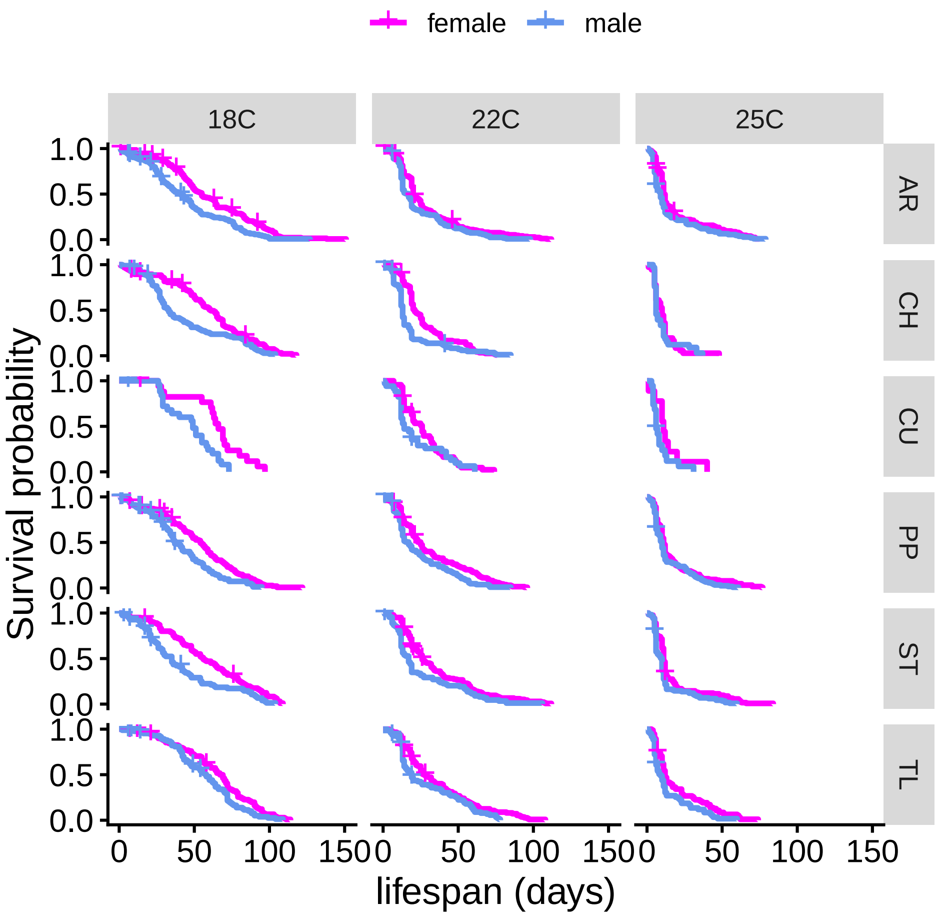


The next graphs compare arrangements within each sex and temperature.

```
ggsurvplot_facet(survfit(Surv(Age.at.death, censrec) ~ sex + arr,
                         data=aging_temps), aging_temps,
                 facet.by = c("sex", "temp"), short.panel.labs=TRUE,
                 xlab="lifespan (days)", legend.title="") +
    scale_color_manual(values = c("AR" = "blue", "CU" = "black", "ST"="darkgreen",
                                  "CH"="orange", "PP"="red", "TL"="purple"))
```

```
## Warning: ggtheme is not a valid theme.
## Please use `theme()` to construct themes.
```

```
## Ignoring unknown labels:
## * fill : ""
## * linetype : "1"
```

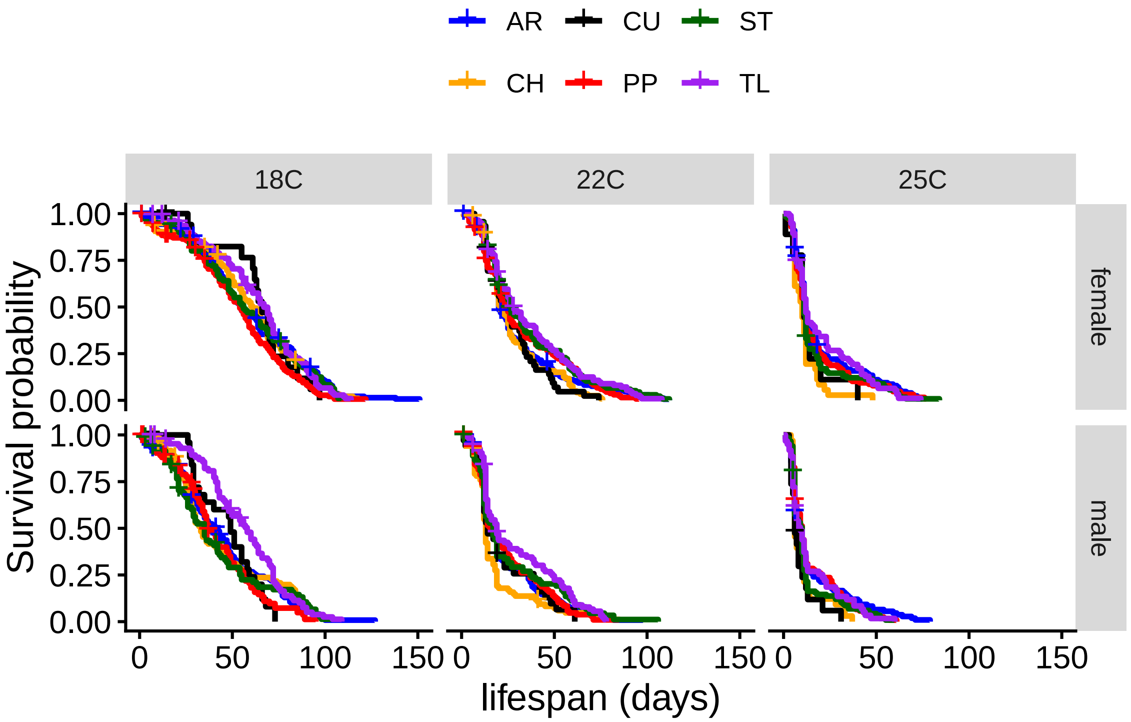


Lastly, we will fit a Gompertz model to each sex, strain, and temperature combination, and then plot the models. We will first generate separate figures for data from each temperature. To do so, we will first define a function to make the plots.

```
gompertz.arr.plot <- function(agingData, xlimit){
    arrangements <- levels(factor(agingData$arr))
    colors <- c( "blue", "orange", "black", "red", "darkgreen", "purple")

        par(mfcol = c(6,2), mar = c(2, 4, 1, 0.5)) # mar = c(bottom,left,top,right)
        for(i in 1:length(arrangements)){
            if(i==1){ # print column heading if first graph
                plot(
                    flexsurvreg(Surv(Age.at.death, censrec) ~ 1,
                             data=subset(agingData, sex=="female" & arr==arrangements[i]),
                             dist = "gompertz"),
                    main="females", las=1, xlab="", ylab="Survival Probability",
                    col=colors[i], xlim=xlimit
                )
                text(x=0.9*xlimit[2], y=0.9, labels=arrangements[i], col=colors[i])
            }else if(i==length(arrangements)){ # print X axis title if last graph
                plot(
                    flexsurvreg(Surv(Age.at.death, censrec) ~ 1,
                             data=subset(agingData, sex=="female" & arr==arrangements[i]),
                             dist = "gompertz"),
                    main="", las=1, xlab="time (days)", ylab="Survival Probability",
                    col=colors[i], xlim=xlimit
                )
                text(x=0.9*xlimit[2], y=0.9, labels=arrangements[i], col=colors[i])
            }else{
                plot(
                    flexsurvreg(Surv(Age.at.death, censrec) ~ 1,
```

```
                                data=subset(agingData, sex=="female" & arr==arrangements[i]),
                                dist = "gompertz"),
                      main="", las=1, xlab="", ylab="Survival Probability",
                      col=colors[i], xlim=xlimit
                )
                text(x=0.9*xlimit[2], y=0.9, labels=arrangements[i], col=colors[i])
            }
        }
        for(i in 1:length(arrangements)){
            if(i==1){ # print column heading if first graph
                plot(
                    flexsurvreg(Surv(Age.at.death, censrec) ~ 1,
                                data=subset(agingData, sex=="male" & arr==arrangements[i]),
                                dist = "gompertz"),
                    main="males", las=1, xlab="", ylab="", col=colors[i], xlim=xlimit
                )
                text(x=0.9*xlimit[2], y=0.9, labels=arrangements[i], col=colors[i])
            }else if(i==length(arrangements)){ # print X axis title if last graph
                plot(
                    flexsurvreg(Surv(Age.at.death, censrec) ~ 1,
                                data=subset(agingData, sex=="male" & arr==arrangements[i]),
                                dist = "gompertz"),
                    main="", las=1, xlab="time (days)", ylab="", col=colors[i], xlim=xlimit
                )
                text(x=0.9*xlimit[2], y=0.9, labels=arrangements[i], col=colors[i])
            }else{
                plot(
                    flexsurvreg(Surv(Age.at.death, censrec) ~ 1,
                                data=subset(agingData, sex=="male" & arr==arrangements[i]),
                                dist = "gompertz"),
                    main="", las=1, xlab="", ylab="", col=colors[i], xlim=xlimit
                )
                text(x=0.9*xlimit[2], y=0.9, labels=arrangements[i], col=colors[i])
            }
        }
}
```

Next, we call the function with data from each temperature.

```
gompertz.arr.plot(aging_25, c(0,80))
```

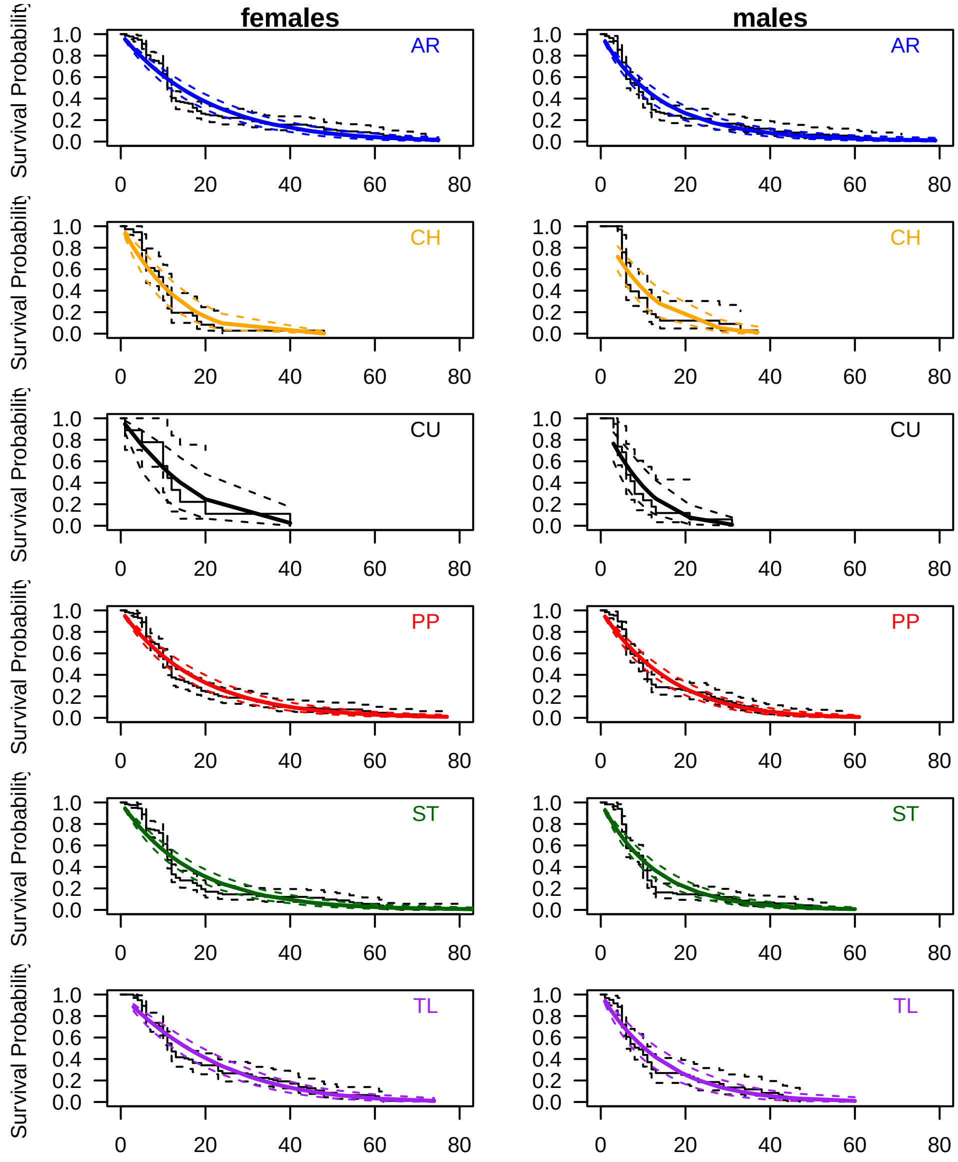


```r
gompertz.arr.plot(aging_22, c(0,100))
```

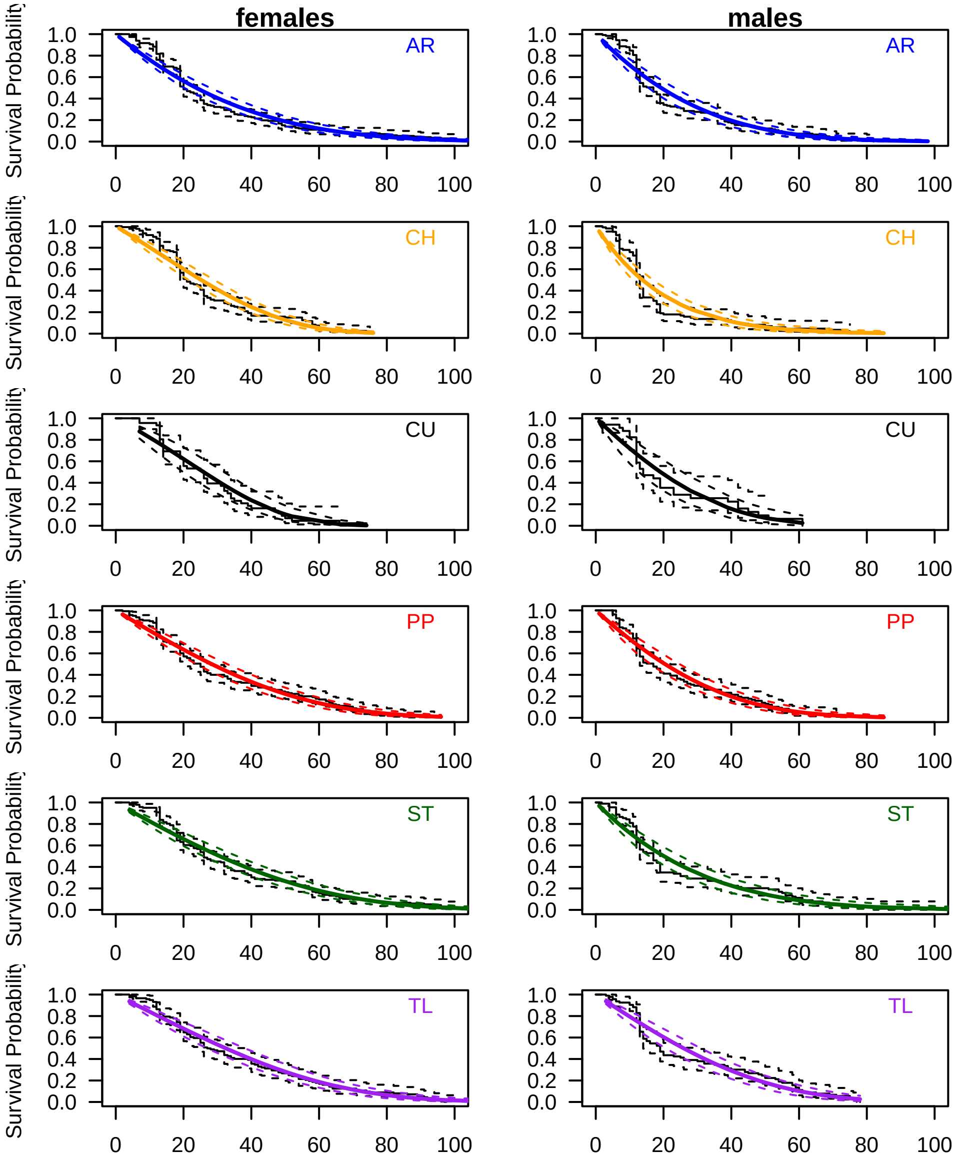


```
gompertz.arr.plot(aging_18, c(0,120))
```

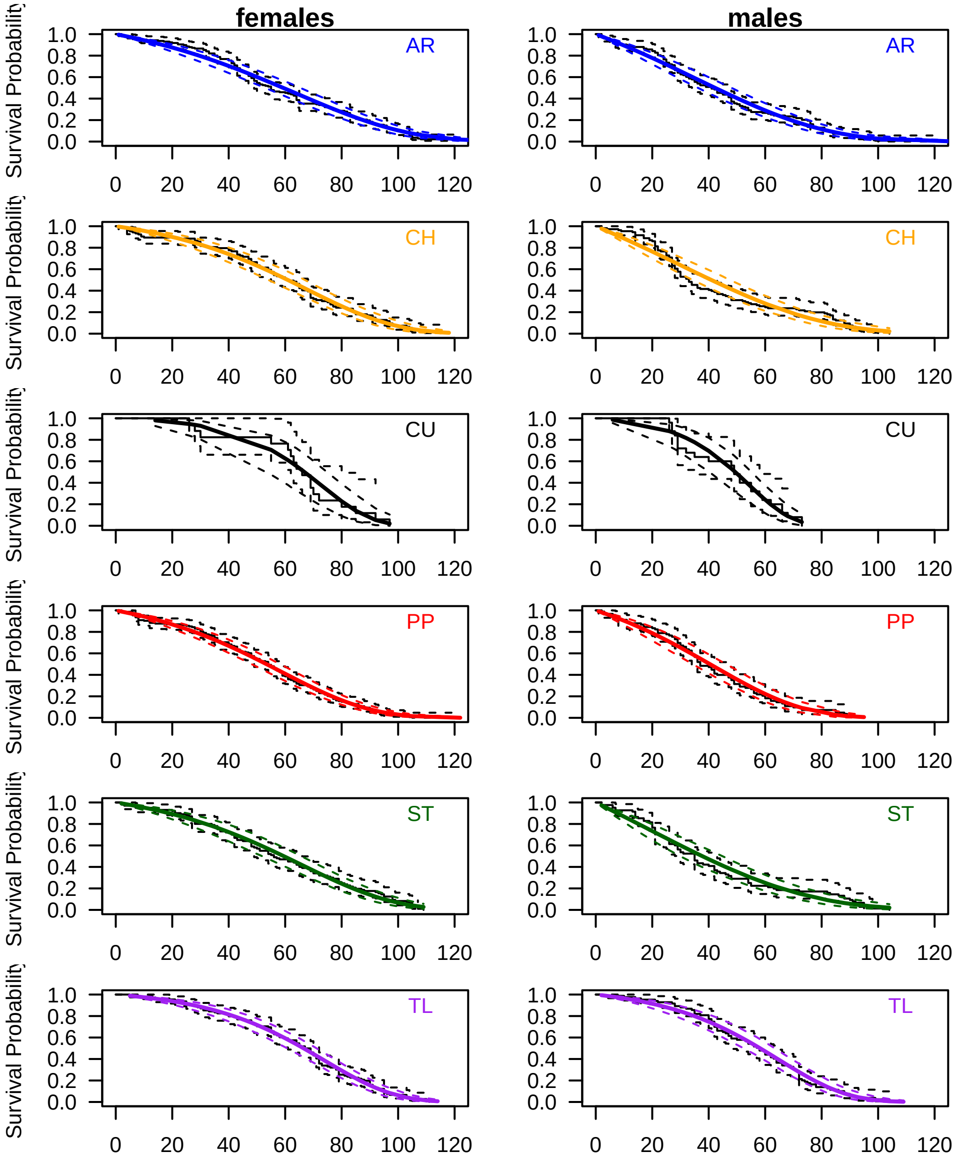


Finally, we will plot the Gompertz models for males and females separately, faceting by arrangement and temperature. Start by defining a function to make the graphs.

```
gompertz.arr.plot.temp <- function(agingData, xlimit){
    arrangements <- levels(factor(agingData$arr))
    colors <- c( "blue", "orange", "black", "red", "darkgreen", "purple")

        par(mfcol = c(6,3), mar = c(2, 4, 1, 0.5)) # mar = c(bottom, left, top, right)
        for(i in 1:length(arrangements)){
            if(i==1){ # print column heading if first graph
                plot(
                    flexsurvreg(Surv(Age.at.death, censrec) ~ 1,
                                data=subset(agingData, temp=="18C" & arr==arrangements[i]),
                                dist = "gompertz"),
                    main="18C", las=1, xlab="", ylab="Survival Probability",
                    col=colors[i], xlim=xlimit
                )
                text(x=0.9*xlimit[2], y=0.9, labels=arrangements[i], col=colors[i])
```

```
        }else if(i==length(arrangements)){ # print X axis title if last graph
            plot(
                flexsurvreg(Surv(Age.at.death, censrec) ~ 1,
                            data=subset(agingData, temp=="18C" & arr==arrangements[i]),
                            dist = "gompertz"),
                main="", las=1, xlab="time (days)", ylab="Survival Probability",
                col=colors[i], xlim=xlimit
            )
            text(x=0.9*xlimit[2], y=0.9, labels=arrangements[i], col=colors[i])
        }else{
            plot(
                flexsurvreg(Surv(Age.at.death, censrec) ~ 1,
                            data=subset(agingData, temp=="18C" & arr==arrangements[i]),
                            dist = "gompertz"),
                main="", las=1, xlab="", ylab="Survival Probability",
                col=colors[i], xlim=xlimit
            )
            text(x=0.9*xlimit[2], y=0.9, labels=arrangements[i], col=colors[i])
        }
    }
    for(i in 1:length(arrangements)){
        if(i==1){ # print column heading if first graph
            plot(
                flexsurvreg(Surv(Age.at.death, censrec) ~ 1,
                            data=subset(agingData, temp=="22C" & arr==arrangements[i]),
                            dist = "gompertz"),
                main="22C", las=1, xlab="", ylab="", col=colors[i], xlim=xlimit
            )
            text(x=0.9*xlimit[2], y=0.9, labels=arrangements[i], col=colors[i])
        }else if(i==length(arrangements)){ # print X axis title if last graph
            plot(
                flexsurvreg(Surv(Age.at.death, censrec) ~ 1,
                            data=subset(agingData, temp=="22C" & arr==arrangements[i]),
                            dist = "gompertz"),
                main="", las=1, xlab="time (days)", ylab="", col=colors[i], xlim=xlimit
            )
            text(x=0.9*xlimit[2], y=0.9, labels=arrangements[i], col=colors[i])
        }else{
            plot(
                flexsurvreg(Surv(Age.at.death, censrec) ~ 1,
                            data=subset(agingData, temp=="22C" & arr==arrangements[i]),
                            dist = "gompertz"),
                main="", las=1, xlab="", ylab="", col=colors[i], xlim=xlimit
            )
            text(x=0.9*xlimit[2], y=0.9, labels=arrangements[i], col=colors[i])
        }
    }
    for(i in 1:length(arrangements)){
        if(i==1){ # print column heading if first graph
            plot(
                flexsurvreg(Surv(Age.at.death, censrec) ~ 1,
                            data=subset(agingData, temp=="25C" & arr==arrangements[i]),
                            dist = "gompertz"),
```

```
                    main="25C", las=1, xlab="", ylab="", col=colors[i], xlim=xlimit
                )
                text(x=0.9*xlimit[2], y=0.9, labels=arrangements[i], col=colors[i])
            }else if(i==length(arrangements)){ # print X axis title if last graph
                plot(
                    flexsurvreg(Surv(Age.at.death, censrec) ~ 1,
                                data=subset(agingData, temp=="25C" & arr==arrangements[i]),
                                dist = "gompertz"),
                    main="", las=1, xlab="time (days)", ylab="", col=colors[i], xlim=xlimit
                )
                text(x=0.9*xlimit[2], y=0.9, labels=arrangements[i], col=colors[i])
            }else{
                plot(
                    flexsurvreg(Surv(Age.at.death, censrec) ~ 1,
                                data=subset(agingData, temp=="25C" & arr==arrangements[i]),
                                dist = "gompertz"),
                    main="", las=1, xlab="", ylab="", col=colors[i], xlim=xlimit
                )
                text(x=0.9*xlimit[2], y=0.9, labels=arrangements[i], col=colors[i])
            }
        }
}
```

Call the function on data from females.

```
gompertz.arr.plot.temp(subset(aging_temps, sex=="female"), c(0,100))
```

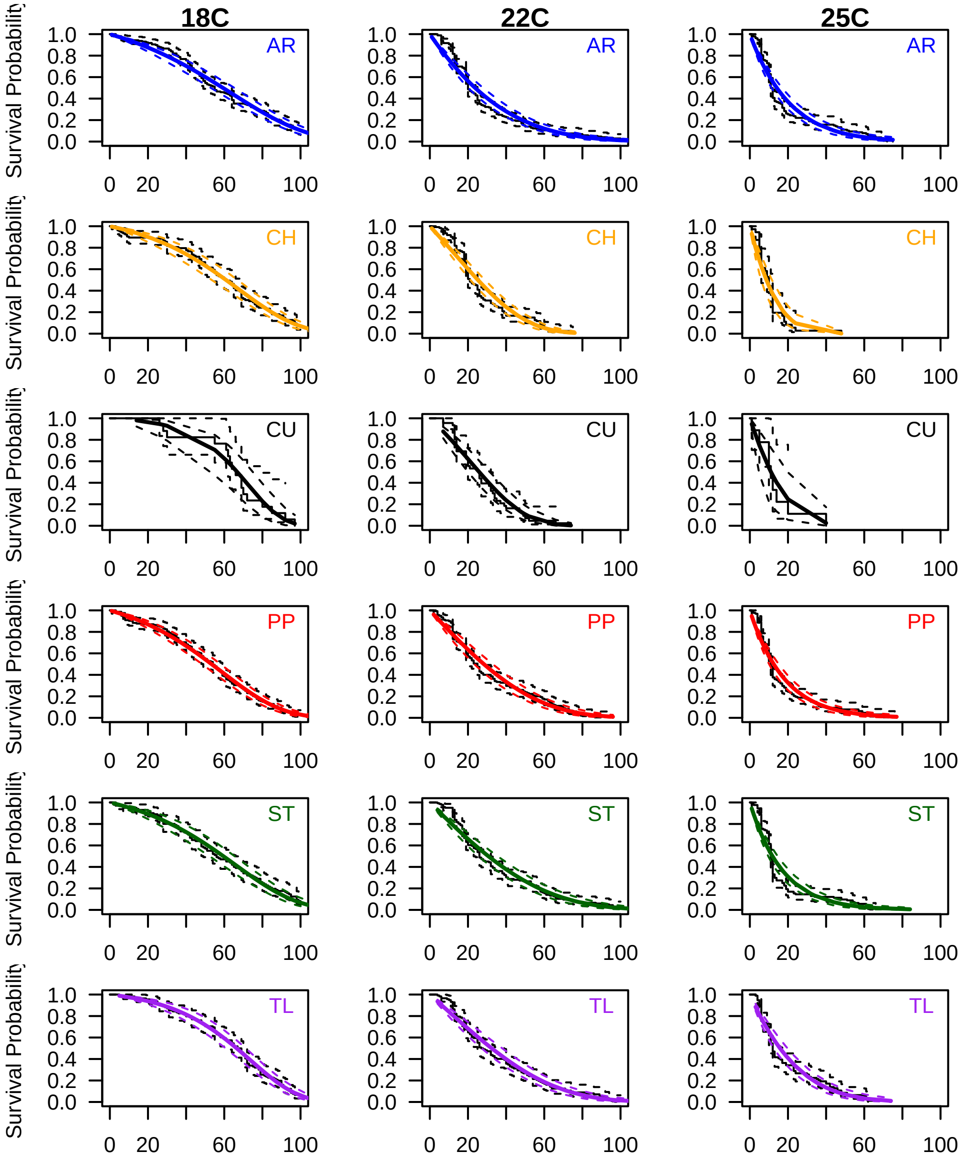


Call the function on data from males.

```
gompertz.arr.plot.temp(subset(aging_temps, sex=="male"), c(0,100))
```

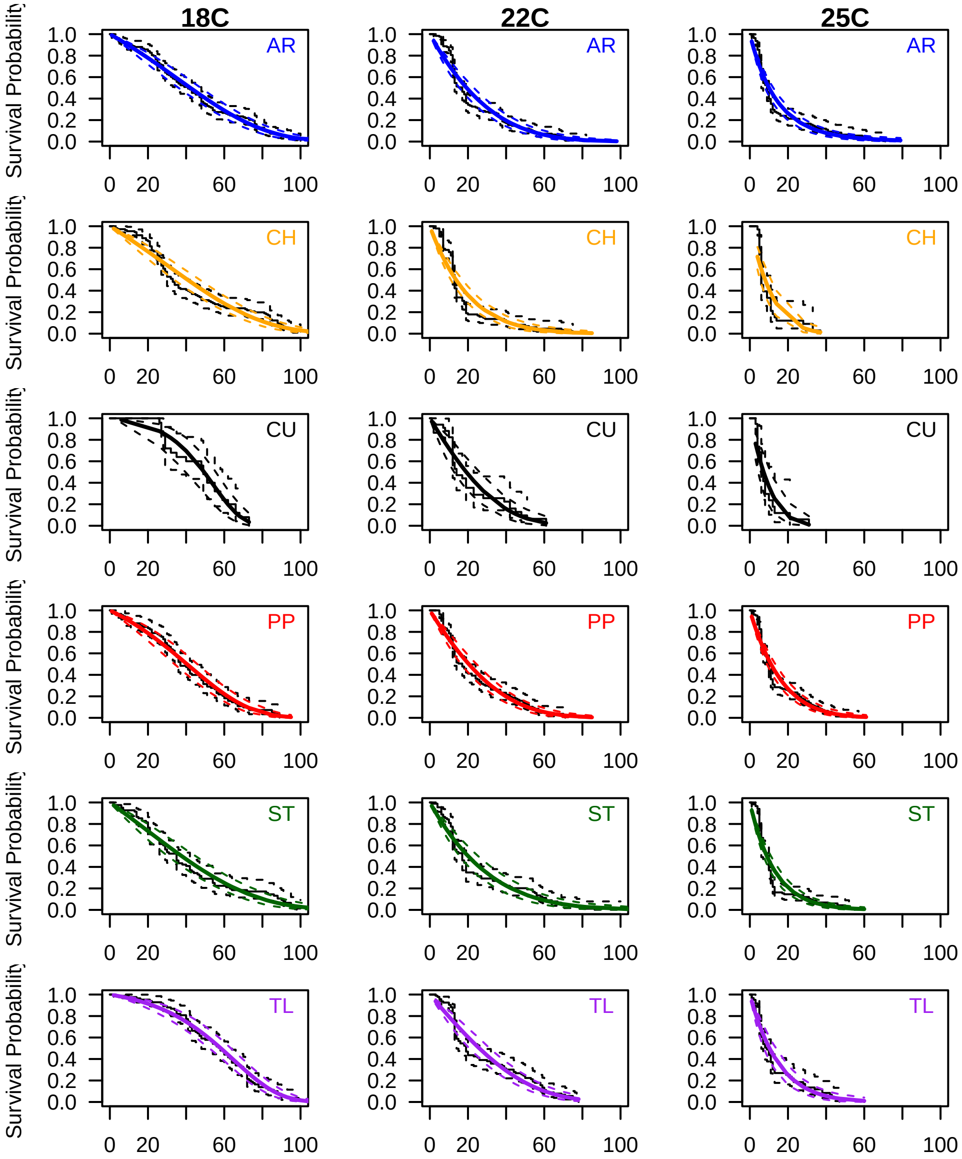


An alternative way to visualize the Gompertz model is to plotting the log of the age-dependent mortality rate over time.
The time dependent mortality rate in Gompertz model is represented as $\mu(t) = ae^{bt}$, where $a$ is the initial mortality rate (rate parameter) and $b$ is the rate of aging (shape paramter). Calling `flexsurvreg()` gives us the model coefficients if we run it on each subset of data (i.e., each sex-by-arrangement combination). We will do this by writing a function to perform the analysis on each arrangement.

```
gompertz.model.extract <- function(aging_sex){
  Arrs <- levels(factor(aging_sex$arr))
  return.df <- data.frame(
    arr = Arrs,
    shape = NA,
    rate = NA
  )
  for(i in 1:length(Arrs)){
    return.df[i, 2:3] <- flexsurvreg(
      Surv(Age.at.death, censrec) ~ Birth,
        data = subset(aging_sex, arr==Arrs[i]),
```

```r
        dist = "gompertz")$res[1:2,1]
  }
  return.df
}

gompertz_model_fit_sextemp <- rbind(
  cbind(
    gompertz.model.extract(subset(aging_temps, sex=="female" & temp=="18C")),
    sex="female", temp="18C"
  ),
  cbind(
    gompertz.model.extract(subset(aging_temps, sex=="male" & temp=="18C")),
    sex="male", temp="18C"
  ),
  cbind(
    gompertz.model.extract(subset(aging_temps, sex=="female" & temp=="22C")),
    sex="female", temp="22C"
  ),
  cbind(
    gompertz.model.extract(subset(aging_temps, sex=="male" & temp=="22C")),
    sex="male", temp="22C"
  ),
  cbind(
    gompertz.model.extract(subset(aging_temps, sex=="female" & temp=="25C")),
    sex="female", temp="25C"
  ),
  cbind(
    gompertz.model.extract(subset(aging_temps, sex=="male" & temp=="25C")),
    sex="male", temp="25C"
  )
)
gompertz_model_fit_sextemp
```

```
##    arr      shape         rate    sex temp
## 1   AR 0.02771970 0.0082785922 female  18C
## 2   CH 0.03777233 0.0028655751 female  18C
## 3   CU 0.07665723 0.0004101269 female  18C
## 4   PP 0.03589746 0.0043127712 female  18C
## 5   ST 0.03234556 0.0055713676 female  18C
## 6   TL 0.04558694 0.0026086905 female  18C
## 7   AR 0.02545997 0.0076519249   male  18C
## 8   CH 0.02919926 0.0378545283   male  18C
## 9   CU 0.09292013 0.0005637864   male  18C
## 10  PP 0.03744577 0.0065762500   male  18C
## 11  ST 0.02051627 0.0106120878   male  18C
## 12  TL 0.04803768 0.0039755824   male  18C
## 13  AR 0.01389868 0.0243178259 female  22C
## 14  CH 0.03338845 0.0108293067 female  22C
## 15  CU 0.05986695 0.0029720543 female  22C
## 16  PP 0.02051446 0.0145777639 female  22C
## 17  ST 0.02426794 0.0154786788 female  22C
## 18  TL 0.03283762 0.0092455006 female  22C
## 19  AR 0.02221625 0.0223074766   male  22C
## 20  CH 0.01974567 0.0726068141   male  22C
```

```
## 21   CU 0.04281504 0.0145976750   male  22C
## 22   PP 0.01936413 0.0621338208   male  22C
## 23   ST 0.02840290 0.0255748080   male  22C
## 24   TL 0.03336668 0.0138151278   male  22C
## 25   AR 0.02892804 0.0618341206 female  25C
## 26   CH 0.14618246 0.0536031903 female  25C
## 27   CU 0.07916755 0.0569984531 female  25C
## 28   PP 0.02000751 0.0287241337 female  25C
## 29   ST 0.01536296 0.0965560599 female  25C
## 30   TL 0.03439788 0.0775661126 female  25C
## 31   AR 0.02520965 0.1238766363   male  25C
## 32   CH 0.08091717 0.0974631814   male  25C
## 33   CU 0.48612172 0.0018945471   male  25C
## 34   PP 0.03142126 0.0221624910   male  25C
## 35   ST 0.02854680 0.0284226943   male  25C
## 36   TL 0.03002329 0.0301833069   male  25C
```

Plot the mortality rate over time. To do so, write a function to plot the natural log of the gompertz mortality rate, and another function to make the graphs for each sex by temperature combination.

```
gompertz.mortality <- function(x, ShapeRate){
  log(ShapeRate[2] * exp(ShapeRate[1] * x))
}


gompertz.mortality.plot.sextemp <- function(GMF){
  ggplot() + xlim(0, 50) + ylim(-8,2) +
    scale_color_manual(name="", guide="none",
      values = c(
          "AR" = "blue",
          "CU" = "black",
          "ST"="darkgreen",
          "CH"="orange",
          "PP"="red",
          "TL"="purple")) +
  geom_function(aes(color=GMF[1,1]), fun=gompertz.mortality,
              args = list( unlist(GMF[1,2:3]))) +
  geom_function(aes(color=GMF[2,1]), fun=gompertz.mortality,
              args = list( unlist(GMF[2,2:3]))) +
  geom_function(aes(color=GMF[3,1]), fun=gompertz.mortality,
              args = list( unlist(GMF[3,2:3]))) +
  geom_function(aes(color=GMF[4,1]), fun=gompertz.mortality,
              args = list( unlist(GMF[4,2:3]))) +
  geom_function(aes(color=GMF[5,1]), fun=gompertz.mortality,
              args = list( unlist(GMF[5,2:3]))) +
  geom_function(aes(color=GMF[6,1]), fun=gompertz.mortality,
              args = list( unlist(GMF[6,2:3]))) +
  xlab("time (days)") + ylab("ln(mortality rate)") + theme_bw()
}
```

Plot for each sex and temperature combination.

```
plot_grid(
  gompertz.mortality.plot.sextemp(
    subset(gompertz_model_fit_sextemp, sex=="female" & temp=="18C")),
  gompertz.mortality.plot.sextemp(
```

```
    subset(gompertz_model_fit_sextemp, sex=="female" & temp=="22C")),
  gompertz.mortality.plot.sextemp(
    subset(gompertz_model_fit_sextemp, sex=="female" & temp=="25C")),
  gompertz.mortality.plot.sextemp(
    subset(gompertz_model_fit_sextemp, sex=="male" & temp=="18C")),
  gompertz.mortality.plot.sextemp(
    subset(gompertz_model_fit_sextemp, sex=="male" & temp=="22C")),
  gompertz.mortality.plot.sextemp(
    subset(gompertz_model_fit_sextemp, sex=="male" & temp=="25C")),
  labels=c("18\u00B0C", "22\u00B0C", "25\u00B0C", "18\u00B0C", "22\u00B0C", "25\u00B0C"),
  nrow=2, label_size = 10, label_x=0.25, label_y=0.95
)
```

```
## Warning: Removed 33 rows containing missing values or values outside the scale range
## (`geom_function()`).
```

```
## Warning: Removed 66 rows containing missing values or values outside the scale range
## (`geom_function()`).
```

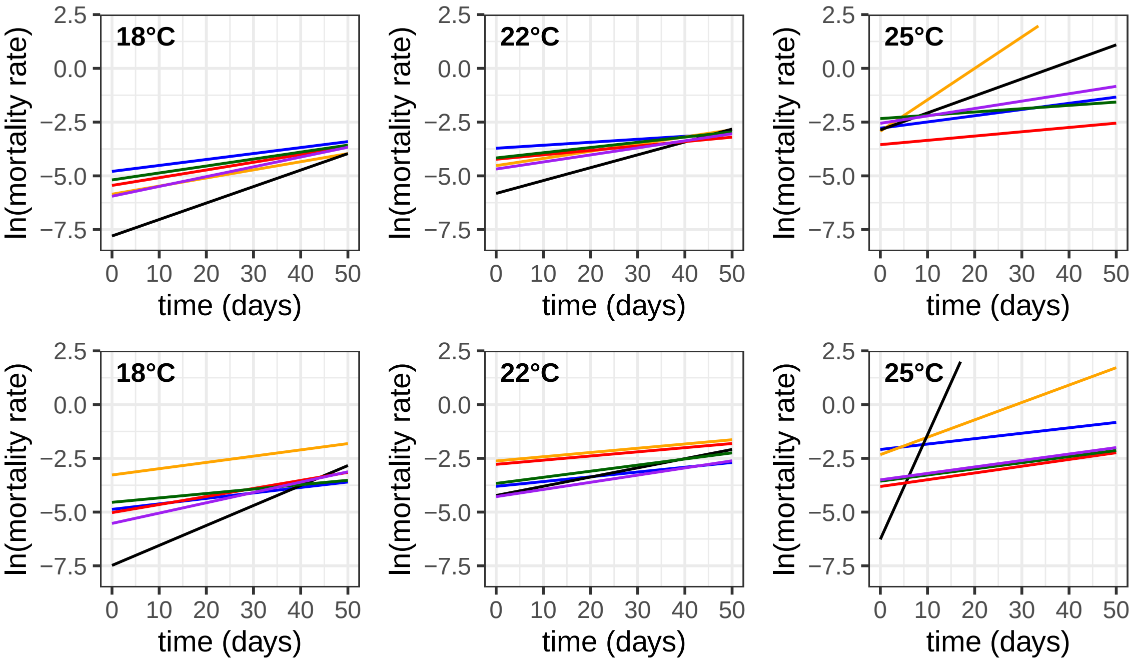
