## Supplemental File S3 for "*Drosophila pseudoobscura* third chromosome inversion arrangements have temperature-dependent and sex-specific effects on life history traits"

### Supplementary File S3: Analyze development rate data from *Drosophila pseudoobscura* at three different temperatures

2026-09-03

#### Overview

The analysis described below was performed on developmental time measurements from *Drosophila pseudoobscura* collected in 2025.
Males and females carrying one of six different third chromosome inversions arrangements were sampled. The flies were sampled from 28 different strains, each of which is homozygous for one of six different chromosomal inversion arrangements.

Five female flies from a given strain were place into a vial with standard *Drosophila* medium, and allowed to lay eggs for 24 hrs. Emerging flies were collected daily, and the sex at emergence from pupation was recorded. Egg-to-adult development time was recorded as the difference in days from when the female (i.e., mother) was placed in the vial and the date when the fly emerged. Flies were raised at 18°C, 22°C, or 25°C throughout the experiment (both parents and offspring).

#### Install and Load Required Packages

```
if (!require("lme4")) install.packages("lme4")
if (!require("nlme")) install.packages("nlme")
if (!require("ggplot2")) install.packages("ggplot2")
if (!require("ggridges")) install.packages("ggridges")
if (!require("tidyr")) install.packages("tidyr")
if (!require("dplyr")) install.packages("dplyr")
if (!require("cowplot")) install.packages("cowplot")

library(lme4)
library(nlme)
library(ggplot2)
library(ggridges)
library(tidyr)
library(dplyr)
library(cowplot)
```

#### Load, Prepare, and QC the Data

Load the data on the egg-to-adult development time from each temperature.

```
dev_18 <- read.delim("Dpse-devrate-18c_2025_08.txt")
dev_22 <- read.delim("Dpse-devrate-22c_2025_08.txt")
dev_25 <- read.delim("Dpse-devrate-25c_2025_08.txt")
```

Confirm that all dates are formatted the same. First, create objects that have the egg laying dates and emergence (birth) dates for the flies.

```
laid_18 <- levels(factor(dev_18$Date.Laid))
laid_22 <- levels(factor(dev_22$Date.Laid))
laid_25 <- levels(factor(dev_25$Date.Laid))

births_18 <- levels(factor(dev_18$Date.of.Emergence))
births_22 <- levels(factor(dev_22$Date.of.Emergence))
births_25 <- levels(factor(dev_25$Date.of.Emergence))
```

Next, count the number of flies with each egg laying and emergence date at each temperature. Check if any dates are singletons, which may be errors that need to be corrected.

```
# look at number of observations per birth date.
datecounter <- function(dates, counts){
    counts.df <- data.frame(
      date = counts
    )
  Dates.df <- data.frame(
        date = dates,
        count = NA
    )
    for(i in 1:length(dates)){
        Dates.df$count[i] <- length(subset(counts.df, date==dates[i])[,1])
    }
    Dates.df
}

laid_18.df <- datecounter(laid_18, dev_18$Date.Laid)
laid_22.df <- datecounter(laid_22, dev_22$Date.Laid)
laid_25.df <- datecounter(laid_25, dev_25$Date.Laid)

Births_18.df <- datecounter(births_18, dev_18$Date.of.Emergence)
Births_22.df <- datecounter(births_22, dev_22$Date.of.Emergence)
Births_25.df <- datecounter(births_25, dev_25$Date.of.Emergence)
```

The egg laying data from 18°C and 25°C look fine. However, there are only 2 samples from the 28-May egg laying date in the 22°C flies.

```
laid_22.df
```

```
##           date count
## 1   1-Jul-2025   431
## 2  10-Jun-2025   209
## 3  16-Jun-2025   101
## 4  17-Jun-2025   129
## 5   2-Jun-2025   313
## 6  23-Jun-2025   302
## 7  24-Jun-2025   416
## 8  28-May-2025     2
## 9   3-Jun-2025   418
## 10 30-Jun-2025   517
## 11  9-Jun-2025   194
```

We will remove the 28-May samples because we cannot effectively model batch effects with so few samples.

```
dev_22 <- subset(dev_22, Date.Laid != "28-May-2025")
laid_22 <- levels(factor(dev_22$Date.Laid))
```

```
laid_22.df <- datecounter(laid_22, dev_22$Date.Laid)
Births_22.df <- datecounter(births_22, dev_22$Date.of.Emergence)
laid_22.df
```

```
##           date count
## 1   1-Jul-2025   431
## 2  10-Jun-2025   209
## 3  16-Jun-2025   101
## 4  17-Jun-2025   129
## 5   2-Jun-2025   313
## 6  23-Jun-2025   302
## 7  24-Jun-2025   416
## 8   3-Jun-2025   418
## 9  30-Jun-2025   517
## 10  9-Jun-2025   194
```

We will next load an object that contains information about the strains. Use that data frame to assign strain information to the samples in the data file. Also create a column `batch` that indicates the vial in which the flies are collected. The vial has a strain number, code, and date when it was set up. Combining those three pieces of information gives a unique batch (vial ID) for each vial.

```
# load strain information
strainIDs <- read.delim("Dpse-strains.tsv")

# assign strain names
strain.assign <- function(strains, datas){
    datas$arr <- strains$arrangement[match(datas$Strain, strains$ID)]
    datas$isogen <- strains$isogen[match(datas$Strain, strains$ID)]
    datas$bal <- strains$balancer[match(datas$Strain, strains$ID)]

    # add batch (strain + date + vial code) to each dataframe
    datas$batch <- paste(datas$Strain, datas$Date.Laid, datas$Vial.Code, sep="_")

    # Omit data with non-existent strain ID
    datas <- subset(datas, !is.na(arr))
}
dev_18 <- strain.assign(strainIDs, dev_18)
dev_22 <- strain.assign(strainIDs, dev_22)
dev_25 <- strain.assign(strainIDs, dev_25)
```

Create a table with all strains assayed for each temperature.

```
aging.strains <- function(aging, strains){
    aging_strains <- data.frame(
        strains = levels(factor(aging$Strain))
    )
    aging_strains$arr <- strains$arrangement[match(aging_strains$strains, strains$ID)]
    aging_strains$isogen <- strains$isogen[match(aging_strains$strains, strains$ID)]
    aging_strains$bal <- strains$balancer[match(aging_strains$strains, strains$ID)]
    for(i in 1:length(aging_strains$strains)){
        aging_strains$females[i] <- length(subset(aging,
                                                  Strain==aging_strains$strains[i] &
                                                    Sex=="female")[,1])
        aging_strains$males[i] <- length(subset(aging,
                                                Strain==aging_strains$strains[i] &
```

```
                                                         Sex=="male")[,1])
    }
    aging_strains
}

dev_strains_18 <- aging.strains(dev_18, strainIDs)
dev_strains_22 <- aging.strains(dev_22, strainIDs)
dev_strains_25 <- aging.strains(dev_25, strainIDs)
```

Check if any strains are under-sampled at a temperature. Remove under-sampled strains from each temperature. Set a threshold of 10 flies for a strain to be included. Start by checking 18°C flies.

```
dev_strains_18[order(dev_strains_18$females + dev_strains_18$males),]
```

```
##    strains arr     isogen bal females males
## 1        3  AR     DM1015   L      12    17
## 18      35  CH        JR4   L      20    13
## 11      22  PP     DM1065   L      22    15
## 2        4  AR     DM1050   L      14    24
## 3        5  AR     DM1056   L      17    22
## 21      42  CH     MSH202   B      21    21
## 12      24  PP     DM1085   B      32    20
## 19      36  CH       JR20   L      25    30
## 24      47  TL SPE123_2-3   B      29    29
## 25      48  TL SPE123_5-1   B      33    34
## 26      49  TL SPE123_6-3   B      36    31
## 6       14  AR     MSH126   L      29    40
## 8       18  PP     DM1020   B      30    48
## 20      38  CH      JR272   B      41    37
## 16      33  ST     MSH217   B      50    37
## 17      34  CH       JR32   B      50    59
## 4        7  AR      KB635   L      63    51
## 15      31  ST       JR91   L      58    62
## 27      53  CU SPE123_4-1   B      65    65
## 23      46  TL    SCI12-2   B      64    69
## 10      20  PP     DM1049   B      65    69
## 7       17  PP    BdA1137  10      72    74
## 9       19  PP     DM1038   B      77    85
## 14      28  ST      JR209   L     114    84
## 13      26  ST      JR138   L      96   105
## 22      43  TL     MA1959         111   121
## 5        8  AR      KB652   L     140   151
```

All strains at 18°C have at least 29 flies.

Next, check the 22°C samples.

```
dev_strains_22[order(dev_strains_22$females + dev_strains_22$males),]
```

```
##    strains arr     isogen bal females males
## 9       22  PP     DM1065   L       3     3
## 16      35  CH        JR4   L      12    16
## 17      36  CH       JR20   L      12    23
## 18      38  CH      JR272   B      27    25
## 23      48  TL SPE123_5-1   B      32    21
## 13      31  ST       JR91   L      31    27
```

```
## 4          7  AR      KB635   L      25    34
## 22        47  TL SPE123_2-3   B      32    32
## 8         19  PP     DM1038   B      47    24
## 1          3  AR     DM1015   L      37    36
## 10        24  PP     DM1085   B      41    35
## 3          5  AR     DM1056   L      49    36
## 15        34  CH       JR32   B      44    41
## 24        49  TL SPE123_6-3   B      52    51
## 11        26  ST      JR138   L      47    60
## 21        46  TL    SCI12-2   B      74    56
## 19        42  CH     MSH202   B      74    62
## 2          4  AR     DM1050   L      86    84
## 5          8  AR      KB652   L     106    68
## 6         14  AR     MSH126   L      86    91
## 20        43  TL     MA1959          96   109
## 25        53  CU SPE123_4-1   B     103   108
## 12        28  ST      JR209   L     114   101
## 14        33  ST     MSH217   B     120   139
## 7         17  PP    BdA1137  10     196   200
```

Remove strain 22 at 22°C because <10 flies.

```
dev_22 <- subset(dev_22, Strain!=22)
dev_strains_22 <- aging.strains(dev_22, strainIDs)
dev_strains_22[order(dev_strains_22$females + dev_strains_22$males),]
```

```
##    strains arr     isogen bal females males
## 15      35  CH        JR4   L      12    16
## 16      36  CH       JR20   L      12    23
## 17      38  CH      JR272   B      27    25
## 22      48  TL SPE123_5-1   B      32    21
## 12      31  ST       JR91   L      31    27
## 4        7  AR      KB635   L      25    34
## 21      47  TL SPE123_2-3   B      32    32
## 8       19  PP     DM1038   B      47    24
## 1        3  AR     DM1015   L      37    36
## 9       24  PP     DM1085   B      41    35
## 3        5  AR     DM1056   L      49    36
## 14      34  CH       JR32   B      44    41
## 23      49  TL SPE123_6-3   B      52    51
## 10      26  ST      JR138   L      47    60
## 20      46  TL    SCI12-2   B      74    56
## 18      42  CH     MSH202   B      74    62
## 2        4  AR     DM1050   L      86    84
## 5        8  AR      KB652   L     106    68
## 6       14  AR     MSH126   L      86    91
## 19      43  TL     MA1959          96   109
## 24      53  CU SPE123_4-1   B     103   108
## 11      28  ST      JR209   L     114   101
## 13      33  ST     MSH217   B     120   139
## 7       17  PP    BdA1137  10     196   200
```

Lastly, check the 25°C samples.

```
dev_strains_25[order(dev_strains_25$females + dev_strains_25$males),]
```

```
##    strains arr      isogen bal  females males
## 20      46  TL     SCI12-2   B        0     1
## 8       20  PP      DM1049   B        2     1
## 15      35  CH         JR4   L        1     2
## 22      48  TL SPE123_5-1    B        3     4
## 5       14  AR      MSH126   L        4     4
## 16      36  CH        JR20   L        7     3
## 9       24  PP      DM1085   B        5     6
## 14      34  CH        JR32   B        5     6
## 13      33  ST      MSH217   B        4     9
## 17      38  CH       JR272   B        6     8
## 24      53  CU SPE123_4-1    B       12     4
## 7       19  PP      DM1038   B       12     8
## 10      26  ST       JR138   L       11    10
## 18      42  CH      MSH202   B       13    10
## 21      47  TL SPE123_2-3    B       14    10
## 3        7  AR       KB635   L       15    16
## 23      49  TL SPE123_6-3    B       18    15
## 2        4  AR      DM1050   L       20    15
## 4        8  AR       KB652   L       22    17
## 1        3  AR      DM1015   L       34    32
## 12      31  ST        JR91   L       40    36
## 11      28  ST       JR209   L       55    64
## 6       17  PP     BdA1137  10       61    77
## 19      43  TL      MA1959          116   113
```

Remove strains 14, 20, 35, 46, and 48 at 25 °C because <10 flies.

```r
dev_25 <- subset(dev_25,
                 Strain!=14 & Strain!=20 & Strain!=35 & Strain!=46 & Strain!=48)
dev_strains_25 <- aging.strains(dev_25, strainIDs)
dev_strains_25[order(dev_strains_25$females + dev_strains_25$males),]
```

```
##    strains arr      isogen bal  females males
## 13      36  CH        JR20   L        7     3
## 7       24  PP      DM1085   B        5     6
## 12      34  CH        JR32   B        5     6
## 11      33  ST      MSH217   B        4     9
## 14      38  CH       JR272   B        6     8
## 19      53  CU SPE123_4-1    B       12     4
## 6       19  PP      DM1038   B       12     8
## 8       26  ST       JR138   L       11    10
## 15      42  CH      MSH202   B       13    10
## 17      47  TL SPE123_2-3    B       14    10
## 3        7  AR       KB635   L       15    16
## 18      49  TL SPE123_6-3    B       18    15
## 2        4  AR      DM1050   L       20    15
## 4        8  AR       KB652   L       22    17
## 1        3  AR      DM1015   L       34    32
## 10      31  ST        JR91   L       40    36
## 9       28  ST       JR209   L       55    64
## 5       17  PP     BdA1137  10       61    77
## 16      43  TL      MA1959          116   113
```

Write data to file after removing the strains with <10 flies.

```r
write.table(dev_strains_18, "dev_strains_18.tsv",
            sep="\t", quote=FALSE, row.names=FALSE)
write.table(dev_strains_22, "dev_strains_22.tsv",
            sep="\t", quote=FALSE, row.names=FALSE)
write.table(dev_strains_25, "dev_strains_25.tsv",
            sep="\t", quote=FALSE, row.names=FALSE)
```

### Perform statistical tests to identify what affects development time at each temperature

First, test for the effect of arrangement and sex on development time within each temperature. Use a mixed model with include vial (batch) and strain (nested in arrangement) as random effects. We will use `lmer()` from the `lme4` package to perform comparisons between nested models. If a factor has a significant effect on model fit, then it is interpreted to affect development time. Define functions to perform model comparisons.

```r
# test for interaction effect
sex.arr.int.lmer <- function(devdata){
    return.anova <- anova(
        lmer(time ~ Sex + arr + Sex*arr + (1|arr:isogen) + (1|batch), data=devdata),
        lmer(time ~ Sex + arr + (1|arr:isogen) + (1|batch), data=devdata)
    )
    rownames(return.anova) <- c("model1", "model2")
    return.anova
}

# test for a sex effect
sex.arr.lmer <- function(devdata){
    return.anova <- anova(
        lmer(time ~ Sex + arr + (1|arr:isogen) + (1|batch), data=devdata),
        lmer(time ~ arr + (1|arr:isogen) + (1|batch), data=devdata)
    )
    rownames(return.anova) <- c("model1", "model2")
    return.anova
}

# test for arrangement effect
arr.sex.lmer <- function(devdata){
    return.anova <- anova(
        lmer(time ~ Sex + arr + (1|arr:isogen) + (1|batch), data=devdata),
        lmer(time ~ Sex + (1|arr:isogen) + (1|batch), data=devdata)
    )
    rownames(return.anova) <- c("model1", "model2")
    return.anova
}
```

Test the 18°C flies first by comparing models with and without an interaction term.

```r
sex.arr.int.lmer(dev_18)
```

```
## refitting model(s) with ML (instead of REML)

## Data: devdata
## Models:
## lmer(time ~ Sex + arr + (1 | arr:isogen) + (1 | batch), data = devdata): time ~ Sex + arr + (1 | arr
```

```
## lmer(time ~ Sex + arr + Sex * arr + (1 | arr:isogen) + (1 | batch), data = devdata): time ~ Sex + ar
##        npar    AIC    BIC  logLik -2*log(L)  Chisq Df Pr(>Chisq)
## model1   10 9694.7 9754.1 -4837.4    9674.7
## model2   15 9696.9 9785.9 -4833.4    9666.9 7.8343  5     0.1656
```

The interaction term does not improve model fit, and so we conclude that there is not a significant interaction effect at 18°C. We will next test for the sex effect at 18°C.

```
sex.arr.lmer(dev_18)
```

```
## refitting model(s) with ML (instead of REML)

## Data: devdata
## Models:
## lmer(time ~ arr + (1 | arr:isogen) + (1 | batch), data = devdata): time ~ arr + (1 | arr:isogen) + (
## lmer(time ~ Sex + arr + (1 | arr:isogen) + (1 | batch), data = devdata): time ~ Sex + arr + (1 | arr
##        npar    AIC    BIC  logLik -2*log(L)  Chisq Df Pr(>Chisq)
## model1    9 9833.7 9887.1 -4907.8    9815.7
## model2   10 9694.7 9754.1 -4837.4    9674.7 140.94  1  < 2.2e-16 ***
## ---
## Signif. codes:  0 '***' 0.001 '**' 0.01 '*' 0.05 '.' 0.1 ' ' 1
```

Including sex in the model does improve model fit, and so we conclude that sex affects development time at 18°C. Lastly, we will test for an arrangement effect at 18°C.

```
arr.sex.lmer(dev_18)
```

```
## refitting model(s) with ML (instead of REML)

## Data: devdata
## Models:
## lmer(time ~ Sex + (1 | arr:isogen) + (1 | batch), data = devdata): time ~ Sex + (1 | arr:isogen) + (
## lmer(time ~ Sex + arr + (1 | arr:isogen) + (1 | batch), data = devdata): time ~ Sex + arr + (1 | arr
##        npar    AIC    BIC  logLik -2*log(L)  Chisq Df Pr(>Chisq)
## model1    5 9692.0 9721.7 -4841.0    9682.0
## model2   10 9694.7 9754.1 -4837.4    9674.7 7.2621  5     0.2019
```

Arrangement does not improve model fit at 18°C, which means arrangement does not affect developmental rate at 18°C.

We can perform the same three tests for flies at 22°C.

```
sex.arr.int.lmer(dev_22)
```

```
## refitting model(s) with ML (instead of REML)

## Data: devdata
## Models:
## lmer(time ~ Sex + arr + (1 | arr:isogen) + (1 | batch), data = devdata): time ~ Sex + arr + (1 | arr
## lmer(time ~ Sex + arr + Sex * arr + (1 | arr:isogen) + (1 | batch), data = devdata): time ~ Sex + ar
##        npar    AIC    BIC  logLik -2*log(L)  Chisq Df Pr(>Chisq)
## model1   10 9081.3 9141.4 -4530.6    9061.3
## model2   15 9089.5 9179.7 -4529.8    9059.5 1.7891  5     0.8775
```

There is not a significant interaction effect at 22°C.

```
sex.arr.lmer(dev_22)
```

```
## refitting model(s) with ML (instead of REML)

## Data: devdata
```

```
## Models:
## lmer(time ~ arr + (1 | arr:isogen) + (1 | batch), data = devdata): time ~ arr + (1 | arr:isogen) + (
## lmer(time ~ Sex + arr + (1 | arr:isogen) + (1 | batch), data = devdata): time ~ Sex + arr + (1 | arr
##        npar    AIC    BIC  logLik -2*log(L)  Chisq Df Pr(>Chisq)
## model1    9 9241.6 9295.8 -4611.8    9223.6
## model2   10 9081.3 9141.4 -4530.6    9061.3 162.35  1  < 2.2e-16 ***
## ---
## Signif. codes:  0 '***' 0.001 '**' 0.01 '*' 0.05 '.' 0.1 ' ' 1
```

There is a significant sex effect at 22°C.

```r
arr.sex.lmer(dev_22)
```

```
## refitting model(s) with ML (instead of REML)

## Data: devdata
## Models:
## lmer(time ~ Sex + (1 | arr:isogen) + (1 | batch), data = devdata): time ~ Sex + (1 | arr:isogen) + (
## lmer(time ~ Sex + arr + (1 | arr:isogen) + (1 | batch), data = devdata): time ~ Sex + arr + (1 | arr
##        npar    AIC    BIC  logLik -2*log(L)  Chisq Df Pr(>Chisq)
## model1    5 9072.2 9102.2 -4531.1    9062.2
## model2   10 9081.3 9141.4 -4530.6    9061.3 0.8631  5     0.9728
```

And there is not a significant arrangement effect at 22°C. Therefore, similar results at 18°C and 22°C, where only sex affects development time.

And we can perform the three tests for flies at 25°C.

```r
sex.arr.int.lmer(dev_25)
```

```
## refitting model(s) with ML (instead of REML)

## Data: devdata
## Models:
## lmer(time ~ Sex + arr + (1 | arr:isogen) + (1 | batch), data = devdata): time ~ Sex + arr + (1 | arr
## lmer(time ~ Sex + arr + Sex * arr + (1 | arr:isogen) + (1 | batch), data = devdata): time ~ Sex + ar
##        npar    AIC    BIC  logLik -2*log(L)  Chisq Df Pr(>Chisq)
## model1   10 2544.5 2592.8 -1262.2    2524.5
## model2   15 2539.0 2611.5 -1254.5    2509.0 15.491  5   0.008458 **
## ---
## Signif. codes:  0 '***' 0.001 '**' 0.01 '*' 0.05 '.' 0.1 ' ' 1
```

Unlike the other two temperatures, there is a significant interaction effect at 25°C.

```r
sex.arr.lmer(dev_25)
```

```
## refitting model(s) with ML (instead of REML)

## Data: devdata
## Models:
## lmer(time ~ arr + (1 | arr:isogen) + (1 | batch), data = devdata): time ~ arr + (1 | arr:isogen) + (
## lmer(time ~ Sex + arr + (1 | arr:isogen) + (1 | batch), data = devdata): time ~ Sex + arr + (1 | arr
##        npar    AIC    BIC  logLik -2*log(L)  Chisq Df Pr(>Chisq)
## model1    9 2595.0 2638.5 -1288.5    2577.0
## model2   10 2544.5 2592.8 -1262.2    2524.5 52.556  1  4.181e-13 ***
## ---
## Signif. codes:  0 '***' 0.001 '**' 0.01 '*' 0.05 '.' 0.1 ' ' 1
```

And there is also a sex effect at 25°C.

```r
arr.sex.lmer(dev_25)
```

```
## Warning in checkConv(attr(opt, "derivs"), opt$par, ctrl = control$checkConv, :
## Model failed to converge with max|grad| = 0.00225529 (tol = 0.002, component 1)

## refitting model(s) with ML (instead of REML)

## Data: devdata
## Models:
## lmer(time ~ Sex + (1 | arr:isogen) + (1 | batch), data = devdata): time ~ Sex + (1 | arr:isogen) + (
## lmer(time ~ Sex + arr + (1 | arr:isogen) + (1 | batch), data = devdata): time ~ Sex + arr + (1 | arr
##        npar    AIC    BIC  logLik -2*log(L)  Chisq Df Pr(>Chisq)
## model1    5 2543.0 2567.2 -1266.5    2533.0
## model2   10 2544.5 2592.8 -1262.2    2524.5 8.4941  5      0.131
```

But there is no arrangement effect at 25°C.

We will next use the `lme()` function from the `nlme` package to estimate the effects of the predictors that significantly improve the model fit. As above, we will define functions to perform the ANOVA.

```r
# The full model with an interaction term
sex.arr.int.lme <- function(devdata){
    anova(
        lme(fixed = time ~ Sex + arr + Sex*arr,
            data = devdata,
            random = list(~1|isogen, ~1|batch)
        )
    )
}

# A model with both Sex and arrangement effects
sex.arr.lme <- function(devdata){
    anova(
        lme(fixed = time ~ Sex + arr,
            data = devdata,
            random = list(~1|isogen, ~1|batch)
        )
    )
}
```

We will call the model without an interaction term on the data from 18°C and 22°C.

```r
sex.arr.lme(dev_18) # sex effect
```

```
##             numDF denDF   F-value p-value
## (Intercept)     1  2385 21606.346  <.0001
## Sex             1  2385   145.129  <.0001
## arr             5    21     1.234  0.3285
```

This confirms that sex significantly affects development time at 18°C. To measure the effect of sex, we can call `lme()` without wrapping it in the `anova()` function.

```r
lme(fixed = time ~ Sex + arr,
    data = dev_18,
    random = list(~1|isogen, ~1|batch)
)
```

```
## Linear mixed-effects model fit by REML
##   Data: dev_18
```

```
##   Log-restricted-likelihood: -4840.229
##   Fixed: time ~ Sex + arr
## (Intercept)     Sexmale       arrCH       arrCU       arrPP       arrST
##  21.1410623   0.5603029   0.9700284   1.0410907   0.1816686   0.3506533
##       arrTL
##   0.7589593
##
## Random effects:
##  Formula: ~1 | isogen
##         (Intercept)
## StdDev:   0.5867596
##
##  Formula: ~1 | batch %in% isogen
##         (Intercept) Residual
## StdDev:    1.715691 1.136982
##
## Number of Observations: 2799
## Number of Groups:
##            isogen batch %in% isogen
##                27               413
```

Being male has a positive effect on egg-to-adult development time, which means males develop slower than females. The average effect is approximately half a day.

```r
sex.arr.lme(dev_22) # sex effect
```

```
##             numDF denDF   F-value p-value
## (Intercept)     1  2605 28757.144  <.0001
## Sex             1  2605   166.856  <.0001
## arr             5    18     0.124  0.9851
```

This confirms a significant sex effect at 22°C. To measure the effect, we call `lme()` without wrapping it in `anova()`.

```r
lme(fixed = time ~ Sex + arr,
    data = dev_22,
    random = list(~1|isogen, ~1|batch)
)
```

```
## Linear mixed-effects model fit by REML
##   Data: dev_22
##   Log-restricted-likelihood: -4536.02
##   Fixed: time ~ Sex + arr
## (Intercept)     Sexmale       arrCH       arrCU       arrPP       arrST
## 16.97049784  0.47423528  0.09054609 -0.25617367  0.04510645  0.07543112
##       arrTL
## -0.05506712
##
## Random effects:
##  Formula: ~1 | isogen
##         (Intercept)
## StdDev:   0.3998763
##
##  Formula: ~1 | batch %in% isogen
##         (Intercept)  Residual
## StdDev:    1.043637 0.9458031
```

```
##
## Number of Observations: 3022
## Number of Groups:
##            isogen batch %in% isogen
##                24              416
```

Similar to what we observe at 18°C, males at 22°C take approximately half a day to longer than females to go from egg to adult.

And we call the function with the interaction term for the data from 25°C.

```r
sex.arr.int.lme(dev_25) # Sex and Sex-by-Arr interaction
```

```
##             numDF denDF  F-value p-value
## (Intercept)     1   753 9467.411  <.0001
## Sex             1   753   55.207  <.0001
## arr             5    13    1.442  0.2739
## Sex:arr         5   753    3.127  0.0084
```

This confirms the sex effect and the sex-by-arrangement interaction. We can first measure the sex effect using a model without the interaction term.

```r
lme(fixed = time ~ Sex + arr,
    data = dev_25,
    random = list(~1|isogen, ~1|batch)
)
```

```
## Linear mixed-effects model fit by REML
##   Data: dev_25
##   Log-restricted-likelihood: -1264.955
##   Fixed: time ~ Sex + arr
## (Intercept)     Sexmale      arrCH      arrCU      arrPP      arrST
##  14.7149573   0.4150433  0.5461866 -0.7673863  0.1558050 -0.3243702
##       arrTL
##  -0.6023297
##
## Random effects:
##  Formula: ~1 | isogen
##         (Intercept)
## StdDev:   0.5409164
##
##  Formula: ~1 | batch %in% isogen
##         (Intercept) Residual
## StdDev:   0.8637175 0.803471
##
## Number of Observations: 929
## Number of Groups:
##            isogen batch %in% isogen
##                19              170
```

At 25°C, males take 0.415 days longer to complete egg-to-adult development relative to females. By adding the sex-by-arrangement interaction to the model, we can identify which arrangements have the smallest and largest sex differences.

```r
lme(fixed = time ~ Sex + arr + Sex:arr,
    data = dev_25,
    random = list(~1|isogen, ~1|batch)
)
```

```
## Linear mixed-effects model fit by REML
##   Data: dev_25
##   Log-restricted-likelihood: -1260.435
##   Fixed: time ~ Sex + arr + Sex:arr
##   (Intercept)       Sexmale         arrCH         arrCU         arrPP
##   14.77845724    0.27754546    0.60710986   -1.05684155    0.14831794
##         arrST         arrTL Sexmale:arrCH Sexmale:arrCU Sexmale:arrPP
##   -0.53011735   -0.64346399   -0.14523788    1.37369789    0.02084983
## Sexmale:arrST Sexmale:arrTL
##    0.40827713    0.07850184
##
## Random effects:
##  Formula: ~1 | isogen
##         (Intercept)
## StdDev:   0.5431382
##
##  Formula: ~1 | batch %in% isogen
##         (Intercept)  Residual
## StdDev:   0.8738578 0.7967854
##
## Number of Observations: 929
## Number of Groups:
##            isogen batch %in% isogen
##                19               170
```

It can be clearer to see the effects if analyze data from males and females separately.

```
lme(fixed = time ~ arr,
    data = subset(dev_25, Sex=="female"),
    random = list(~1|isogen, ~1|batch)
)
```

```
## Linear mixed-effects model fit by REML
##   Data: subset(dev_25, Sex == "female")
##   Log-restricted-likelihood: -694.3155
##   Fixed: time ~ arr
## (Intercept)       arrCH       arrCU       arrPP       arrST       arrTL
## 14.69816741  0.70542841 -0.97716843  0.06112326 -0.42787664 -0.54823116
##
## Random effects:
##  Formula: ~1 | isogen
##         (Intercept)
## StdDev:    0.503124
##
##  Formula: ~1 | batch %in% isogen
##         (Intercept)  Residual
## StdDev:   0.9600748 0.8389325
##
## Number of Observations: 470
## Number of Groups:
##            isogen batch %in% isogen
##                19               147
```

```
lme(fixed = time ~ arr,
    data = subset(dev_25, Sex=="male"),
    random = list(~1|isogen, ~1|batch)
```

```
)
```

```
## Linear mixed-effects model fit by REML
##   Data: subset(dev_25, Sex == "male")
##   Log-restricted-likelihood: -590.0381
##   Fixed: time ~ arr
## (Intercept)       arrCH       arrCU       arrPP       arrST       arrTL
##  15.1233727   0.3972763  -0.1233727   0.3433824  -0.2689811  -0.5152958
##
## Random effects:
##  Formula: ~1 | isogen
##         (Intercept)
## StdDev:   0.6433525
##
##  Formula: ~1 | batch %in% isogen
##         (Intercept)  Residual
## StdDev:   0.7302888 0.7098037
##
## Number of Observations: 459
## Number of Groups:
##            isogen batch %in% isogen
##                19               129
```

The reference strain in AR, in both cases. Note that the sign of the effect of each arrangement is the same in both sexes, indicating that each arrangement has the same direction of effect relative to AR. However, we can see if the rank order changes between sexes. To do so, we will calculate the average lifespan of each arrangement within each sex by adding their effects to the within sex intercept. We will write a function to do this and make a dataframe of the estimates, and then sort by average female lifespan.

```
dev.effect <- function(devtime){
  FEs <- lme(fixed = time ~ arr,
      data = devtime,
      random = list(~1|isogen, ~1|batch)
  )$coefficients$fixed

  data.frame(
    arr = c("AR", "CH", "CU", "PP", "ST", "TL"),
    effect = c(
     FEs[1],
     FEs[1] + FEs[2],
     FEs[1] + FEs[3],
     FEs[1] + FEs[4],
     FEs[1] + FEs[5],
     FEs[1] + FEs[6]
    )
  )
}
dev_25_arr <- cbind(
  dev.effect(subset(dev_25, Sex == "female")),
  male = dev.effect(subset(dev_25, Sex == "male"))[,2]
)
colnames(dev_25_arr)[2] <- "female"
dev_25_arr$sexdiff <- dev_25_arr$female - dev_25_arr$male
dev_25_arr[order(dev_25_arr$female),]
```

```
##   arr   female     male    sexdiff
## 3  CU 13.72100 15.00000 -1.2790010
## 6  TL 14.14994 14.60808 -0.4581406
## 5  ST 14.27029 14.85439 -0.5841009
## 1  AR 14.69817 15.12337 -0.4252053
## 4  PP 14.75929 15.46676 -0.7074645
## 2  CH 15.40360 15.52065 -0.1170532
```

In all cases, females develop faster than males. However, the sex difference in development time is smallest in flies with the CH arrangement (0.12 days). In contrast, the sex difference is especially pronounced in flies with the CU arrangement, where males take 1.28 days more than females to complete egg-to-adult development.

##### Test how temperature affects development time, with respect to arragement and sex

We will next analyze data from all 3 temperatures in a single model. This will allow us to evaluate how temperature affects the relationships between arrangement, sex, and development time. First, create a single data frame with results from all three temperatures. In addition to binding the data from each temperature, we will add a new batch label that includes temperature.

```
dev_temp <- rbind(
    cbind(temp = "18C", dev_18),
    cbind(temp = "22C", dev_22),
    cbind(temp = "25C", dev_25)
)
dev_temp$batch_temp <- paste(dev_temp$batch, dev_temp$temp)
```

Now, use `lmer()` from the `lme4` package to test for the effects of arrangement, sex, and temperature on development time. As above, we will use model comparisons to evaluate the effects of each factor. First, test if temperature affects development time in a model with sex and arrangement as fixed effects and strain (nested withing arrangement) and batch as random effects.

```
{ temp.anova <- anova (
    lmer(time ~ Sex + arr + temp + (1|arr:isogen) + (1|batch_temp),
         data=dev_temp),
    lmer(time ~ Sex + arr + (1|arr:isogen) + (1|batch_temp),
         data=dev_temp)
  )
    rownames(temp.anova) <- c("model1", "model2")
    temp.anova
}
```

```
## refitting model(s) with ML (instead of REML)

## Data: dev_temp
## Models:
## lmer(time ~ Sex + arr + (1 | arr:isogen) + (1 | batch_temp), data = dev_temp): time ~ Sex + arr + (1
## lmer(time ~ Sex + arr + temp + (1 | arr:isogen) + (1 | batch_temp), data = dev_temp): time ~ Sex + a
##        npar   AIC   BIC logLik -2*log(L)  Chisq Df Pr(>Chisq)
## model1   10 22992 23060 -11486     22972
## model2   12 21592 21673 -10784     21568 1404.6  2  < 2.2e-16 ***
## ---
## Signif. codes:  0 '***' 0.001 '**' 0.01 '*' 0.05 '.' 0.1 ' ' 1
```

Including temperature in the model significantly improves fit. Therefore, temperature affects development time. Let us next do a comparison between models with and without sex as a factor.

```
{ temp.anova <- anova (
    lmer(time ~ Sex + arr + temp + (1|arr:isogen) + (1|batch_temp),
         data=dev_temp),
    lmer(time ~ arr + temp + (1|arr:isogen) + (1|batch_temp),
         data=dev_temp)
  )
    rownames(temp.anova) <- c("model1", "model2")
    temp.anova
}
```

```
## Warning in checkConv(attr(opt, "derivs"), opt$par, ctrl = control$checkConv, :
## Model failed to converge with max|grad| = 0.0139975 (tol = 0.002, component 1)

## refitting model(s) with ML (instead of REML)

## Data: dev_temp
## Models:
## lmer(time ~ arr + temp + (1 | arr:isogen) + (1 | batch_temp), data = dev_temp): time ~ arr + temp +
## lmer(time ~ Sex + arr + temp + (1 | arr:isogen) + (1 | batch_temp), data = dev_temp): time ~ Sex + a
##        npar   AIC   BIC logLik -2*log(L)  Chisq Df Pr(>Chisq)
## model1   11 21936 22011 -10957     21914
## model2   12 21592 21673 -10784     21568 346.09  1  < 2.2e-16 ***
## ---
## Signif. codes:  0 '***' 0.001 '**' 0.01 '*' 0.05 '.' 0.1 ' ' 1
```

Sex also affects model fit. And now we will test if arrangement affects model fit.

```
{ temp.anova <- anova (
    lmer(time ~ Sex + arr + temp + (1|arr:isogen) + (1|batch_temp),
         data=dev_temp),
    lmer(time ~ Sex + temp + (1|arr:isogen) + (1|batch_temp),
         data=dev_temp)
  )
    rownames(temp.anova) <- c("model1", "model2")
    temp.anova
}
```

```
## refitting model(s) with ML (instead of REML)

## Data: dev_temp
## Models:
## lmer(time ~ Sex + temp + (1 | arr:isogen) + (1 | batch_temp), data = dev_temp): time ~ Sex + temp +
## lmer(time ~ Sex + arr + temp + (1 | arr:isogen) + (1 | batch_temp), data = dev_temp): time ~ Sex + a
##        npar   AIC   BIC logLik -2*log(L)  Chisq Df Pr(>Chisq)
## model1    7 21586 21633 -10786     21572
## model2   12 21592 21673 -10784     21568 4.0778  5     0.5383
```

Adding arrangement does not improve model fit. However, we will include arrangement in the analysis below because we will test if interactions with arrangement affect the fit of the model.

To test for interaction effects, we will evaluate if each possible interaction between sex, arrangement, and temperature improves the model fit. First, here is the sex-by-arrangement interaction.

```
{ temp.anova <- anova (
    lmer(time ~ Sex + arr + temp + Sex*arr + (1|arr:isogen) + (1|batch_temp),
         data=dev_temp),
    lmer(time ~ Sex + arr + temp + (1|arr:isogen) + (1|batch_temp),
         data=dev_temp)
```

```
  )
    rownames(temp.anova) <- c("model1", "model2")
    temp.anova
}
```

```
## refitting model(s) with ML (instead of REML)

## Data: dev_temp
## Models:
## lmer(time ~ Sex + arr + temp + (1 | arr:isogen) + (1 | batch_temp), data = dev_temp): time ~ Sex + a
## lmer(time ~ Sex + arr + temp + Sex * arr + (1 | arr:isogen) + (1 | batch_temp), data = dev_temp): ti
##        npar   AIC   BIC logLik -2*log(L) Chisq Df Pr(>Chisq)
## model1   12 21592 21673 -10784     21568
## model2   17 21596 21712 -10781     21562 5.447  5     0.3638
```

The sex-by-arrangement interaction does not improve model fit. Next, we will test the arrangement-by-temperature interaction.

```
{ temp.anova <- anova (
    lmer(time ~ Sex + arr + temp + arr*temp + (1|arr:isogen) + (1|batch_temp),
         data=dev_temp),
    lmer(time ~ Sex + arr + temp + (1|arr:isogen) + (1|batch_temp),
         data=dev_temp)
  )
    rownames(temp.anova) <- c("model1", "model2")
    temp.anova
}
```

```
## refitting model(s) with ML (instead of REML)

## Data: dev_temp
## Models:
## lmer(time ~ Sex + arr + temp + (1 | arr:isogen) + (1 | batch_temp), data = dev_temp): time ~ Sex + a
## lmer(time ~ Sex + arr + temp + arr * temp + (1 | arr:isogen) + (1 | batch_temp), data = dev_temp): t
##        npar   AIC   BIC logLik -2*log(L)  Chisq Df Pr(>Chisq)
## model1   12 21592 21673 -10784     21568
## model2   22 21590 21740 -10773     21546 21.233 10    0.01953 *
## ---
## Signif. codes:  0 '***' 0.001 '**' 0.01 '*' 0.05 '.' 0.1 ' ' 1
```

The arrangement-by-temperature interaction improves model fit. Lastly, we test the sex-by-temperature interaction.

```
{ temp.anova <- anova (
    lmer(time ~ Sex + arr + temp + Sex*temp + (1|arr:isogen) + (1|batch_temp),
         data=dev_temp),
    lmer(time ~ Sex + arr + temp + (1|arr:isogen) + (1|batch_temp),
         data=dev_temp)
  )
    rownames(temp.anova) <- c("model1", "model2")
    temp.anova
}
```

```
## Warning in checkConv(attr(opt, "derivs"), opt$par, ctrl = control$checkConv, :
## Model failed to converge with max|grad| = 0.00527962 (tol = 0.002, component 1)

## refitting model(s) with ML (instead of REML)
```

```
## Data: dev_temp
## Models:
## lmer(time ~ Sex + arr + temp + (1 | arr:isogen) + (1 | batch_temp), data = dev_temp): time ~ Sex + a
## lmer(time ~ Sex + arr + temp + Sex * temp + (1 | arr:isogen) + (1 | batch_temp), data = dev_temp): t
##        npar   AIC   BIC logLik -2*log(L)  Chisq Df Pr(>Chisq)
## model1   12 21592 21673 -10784     21568
## model2   14 21592 21687 -10782     21564 3.5483  2     0.1696
```

The sex-by-temperature interaction does not improve model fit. Therefore, our best fitting model includes the following predictors of development time: temperature, sex, and the arrangement-by-temperature interaction.

Next we will use the `lme()` function from the `nlme` package to construct our best fitting model.

```
anova(
    lme(fixed = time ~ Sex + arr + temp + arr*temp,
        data = dev_temp,
            random = list(~1|isogen, ~1|batch_temp)
    )
)
```

```
##             numDF denDF  F-value p-value
## (Intercept)     1  5750 41087.93  <.0001
## Sex             1  5750   384.70  <.0001
## arr             5    21     3.29  0.0238
## temp            2   960  1542.72  <.0001
## arr:temp       10   960     2.12  0.0204
```

This analysis suggests that sex and temperature have highly significant effects on egg-to-adult development, while arrangement and the arrangement-by-temperature interaction also affect development.

To quantify the effect of temperature on development time, we will run `lme()` on a model with sex, arrangement, and temperature as predictors, but not any interaction terms.

```
lme(fixed = time ~ Sex + arr + temp,
    data = dev_temp,
        random = list(~1|isogen, ~1|batch_temp)
)
```

```
## Linear mixed-effects model fit by REML
##   Data: dev_temp
##   Log-restricted-likelihood: -10792.36
##   Fixed: time ~ Sex + arr + temp
##  (Intercept)      Sexmale        arrCH        arrCU        arrPP        arrST
## 21.471121164  0.500272851  0.489449264  0.158103767  0.003589843  0.082922861
##        arrTL      temp22C      temp25C
##  0.165721348 -4.655709410 -7.053179621
##
## Random effects:
##  Formula: ~1 | isogen
##         (Intercept)
## StdDev:   0.3979685
##
##  Formula: ~1 | batch_temp %in% isogen
##         (Intercept) Residual
## StdDev:    1.381738 1.012576
##
## Number of Observations: 6750
## Number of Groups:
```

```
##                  isogen batch_temp %in% isogen
##                      27                      999
```

When we analyze data from all three temperatures, males take half a day longer to complete egg-to-adult development than females, similar to what we saw above when we analyzed each temperature separately. In addition, flies develop from egg-to-adult 4.7 days faster at 22°C than 18°C, 7 days faster at 25°C than 18°C, and therefore 2.3 days faster at 25°C than 22°C, Therefore, each 1°C leads to 0.77–1 days faster development.

We will now use a model with an interaction term to see how each arrangement modulates the effect of temperature on egg-to-adult development time.

```
lme(fixed = time ~ Sex + arr + temp + arr*temp,
    data = dev_temp,
        random = list(~1|isogen, ~1|batch_temp)
)
```

```
## Linear mixed-effects model fit by REML
##   Data: dev_temp
##   Log-restricted-likelihood: -10782.37
##   Fixed: time ~ Sex + arr + temp + arr * temp
##   (Intercept)       Sexmale         arrCH         arrCU         arrPP
##    21.1875081     0.5001983     0.9070593     1.0149834     0.2037090
##         arrST         arrTL      temp22C       temp25C arrCH:temp22C
##     0.3057118     0.7193379    -4.2374628    -6.3928712    -0.7516539
## arrCU:temp22C arrPP:temp22C arrST:temp22C arrTL:temp22C arrCH:temp25C
##    -1.2429827    -0.2503728    -0.2414159    -0.7651267    -0.3930257
## arrCU:temp25C arrPP:temp25C arrST:temp25C arrTL:temp25C
##    -1.8554885    -0.3472744    -0.6792362    -1.4347426
##
## Random effects:
##  Formula: ~1 | isogen
##         (Intercept)
## StdDev:   0.4048839
##
##  Formula: ~1 | batch_temp %in% isogen
##         (Intercept) Residual
## StdDev:     1.37177 1.012603
##
## Number of Observations: 6750
## Number of Groups:
##                  isogen batch_temp %in% isogen
##                      27                      999
```

This analysis reveals that flies carrying the CU arrangement have the most accelerated development time as temperature increases. In contrast, development of AR flies is least accelerated by the increase in temperature because the interaction terms for all other arrangements are negative.

Let's repeat the analysis looking at males and females separately.

```
lme(fixed = time ~ arr + temp + arr*temp,
    data = subset(dev_temp, Sex=="female"),
        random = list(~1|isogen, ~1|batch_temp)
)
```

```
## Linear mixed-effects model fit by REML
##   Data: subset(dev_temp, Sex == "female")
##   Log-restricted-likelihood: -5714.837
##   Fixed: time ~ arr + temp + arr * temp
```

```
##   (Intercept)         arrCH         arrCU         arrPP         arrST
##   21.30572293    1.14938662    0.75914982    0.07265277    0.34638985
##         arrTL       temp22C       temp25C arrCH:temp22C arrCU:temp22C
##    0.68754448   -4.27708767   -6.51656194   -1.30351940   -1.15091980
## arrPP:temp22C arrST:temp22C arrTL:temp22C arrCH:temp25C arrCU:temp25C
##   -0.21238209   -0.37061437   -0.71679040   -0.44192078   -1.80913828
## arrPP:temp25C arrST:temp25C arrTL:temp25C
##   -0.34196677   -0.74630090   -1.42253681
##
## Random effects:
##  Formula: ~1 | isogen
##         (Intercept)
## StdDev:   0.4425806
##
##  Formula: ~1 | batch_temp %in% isogen
##         (Intercept) Residual
## StdDev:    1.399822 1.027201
##
## Number of Observations: 3399
## Number of Groups:
##                   isogen batch_temp %in% isogen
##                       27                    850
```

We see that females with the CU arrangement have the most accelerated development as temperature increases, followed by TL.

```r
lme(fixed = time ~ arr + temp + arr*temp,
    data = subset(dev_temp, Sex=="male"),
        random = list(~1|isogen, ~1|batch_temp)
)
```

```
## Linear mixed-effects model fit by REML
##   Data: subset(dev_temp, Sex == "male")
##   Log-restricted-likelihood: -5476.167
##   Fixed: time ~ arr + temp + arr * temp
##   (Intercept)         arrCH         arrCU         arrPP         arrST
##   21.66858844    0.92065688    1.27912805    0.37221901    0.30217164
##         arrTL       temp22C       temp25C arrCH:temp22C arrCU:temp22C
##    0.92295913   -4.22988393   -6.40767138   -0.64813724   -1.48370230
## arrPP:temp22C arrST:temp22C arrTL:temp22C arrCH:temp25C arrCU:temp25C
##   -0.37552241   -0.06895331   -1.05761175   -0.66648156   -1.54004511
## arrPP:temp25C arrST:temp25C arrTL:temp25C
##   -0.31860269   -0.63619417   -1.41201476
##
## Random effects:
##  Formula: ~1 | isogen
##         (Intercept)
## StdDev:   0.3595858
##
##  Formula: ~1 | batch_temp %in% isogen
##         (Intercept)  Residual
## StdDev:    1.382973 0.9708654
##
## Number of Observations: 3351
## Number of Groups:
```

```
##                 isogen batch_temp %in% isogen
##                     27                     842
```

Males with CU or TL have the greatest acceleration in development as temperature increases.

Let's now write a function to extract the average development time for each arrangement at each temperature. We can also make a graph of each estimate.

```
arr.temp.effects <- function(dev_temp_sex){
  lme_arr_temp <- lme(fixed = time ~ arr + temp + arr*temp,
      data = dev_temp_sex,
        random = list(~1|isogen, ~1|batch_temp)
  )$coefficients$fixed

  data.frame(
    arr = c("AR", "CH", "CU", "PP", "ST", "TL"),
    temp = c(rep("18C", 6), rep("22C", 6), rep("25C", 6)),
    dev_time = c(
      lme_arr_temp[1],
      lme_arr_temp[1] + lme_arr_temp[2],
      lme_arr_temp[1] + lme_arr_temp[3],
      lme_arr_temp[1] + lme_arr_temp[4],
      lme_arr_temp[1] + lme_arr_temp[5],
      lme_arr_temp[1] + lme_arr_temp[6],
      lme_arr_temp[1] + lme_arr_temp[7],
      lme_arr_temp[1] + lme_arr_temp[7] + lme_arr_temp[2] + lme_arr_temp[9],
      lme_arr_temp[1] + lme_arr_temp[7] + lme_arr_temp[3] + lme_arr_temp[10],
      lme_arr_temp[1] + lme_arr_temp[7] + lme_arr_temp[4] + lme_arr_temp[11],
      lme_arr_temp[1] + lme_arr_temp[7] + lme_arr_temp[5] + lme_arr_temp[12],
      lme_arr_temp[1] + lme_arr_temp[7] + lme_arr_temp[6] + lme_arr_temp[13],
      lme_arr_temp[1] + lme_arr_temp[8],
      lme_arr_temp[1] + lme_arr_temp[8] + lme_arr_temp[2] + lme_arr_temp[14],
      lme_arr_temp[1] + lme_arr_temp[8] + lme_arr_temp[3] + lme_arr_temp[15],
      lme_arr_temp[1] + lme_arr_temp[8] + lme_arr_temp[4] + lme_arr_temp[16],
      lme_arr_temp[1] + lme_arr_temp[8] + lme_arr_temp[5] + lme_arr_temp[17],
      lme_arr_temp[1] + lme_arr_temp[8] + lme_arr_temp[6] + lme_arr_temp[18]
    )
  )
}
sex_arr_temp <- rbind(
  cbind(sex="female", arr.temp.effects(subset(dev_temp, Sex=="female"))),
  cbind(sex="male", arr.temp.effects(subset(dev_temp, Sex=="male")))
)

arrcols <- c( "AR"="blue", "CH"="orange", "CU"="black",
                "PP"="red", "ST"="darkgreen", "TL"="purple")
ggplot(sex_arr_temp, aes(x=temp, y=dev_time, color=arr, group=arr)) +
  geom_point() +
  geom_line() +
  facet_wrap(~sex, nrow=1) +
  scale_color_manual(values=arrcols, name="") +
  scale_x_discrete("temperature") +
  scale_y_continuous("mean development time (days)") +
  theme_bw()
```

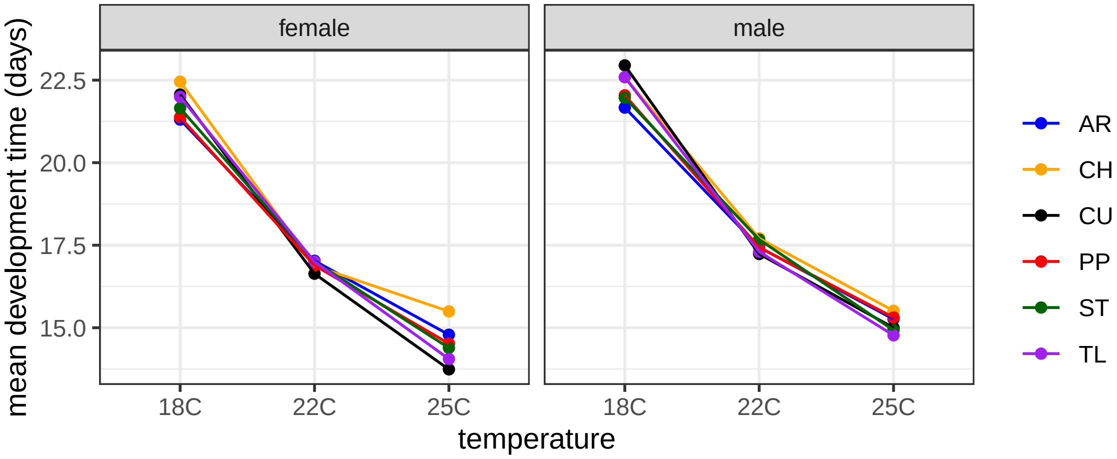


The graph shows us that the rank order of arrangement effects on development time differs across temperatures, when we look within each sex. The graph confirms the statistical test result that flies with the CU and TL arrangements have the most accelerated development as temperature increases. In contrast, AR has the least accelerated development.

To explor how the rank order of each arrangement differs between sexes, we can plot the male-female differences for each arrangement at each temperature.

```
sex_arr_temp_diff <- sex_arr_temp %>% pivot_wider(names_from = sex, values_from = dev_time)
sex_arr_temp_diff$sex_diff <- sex_arr_temp_diff$male - sex_arr_temp_diff$female

ggplot(sex_arr_temp_diff, aes(x=temp, y=sex_diff, color=arr, group=arr)) +
  geom_point() +
  geom_line() +
  scale_color_manual(values=arrcols, name="") +
  scale_x_discrete("temperature") +
  scale_y_continuous("sex difference in dev time (days)") +
  theme_bw()
```

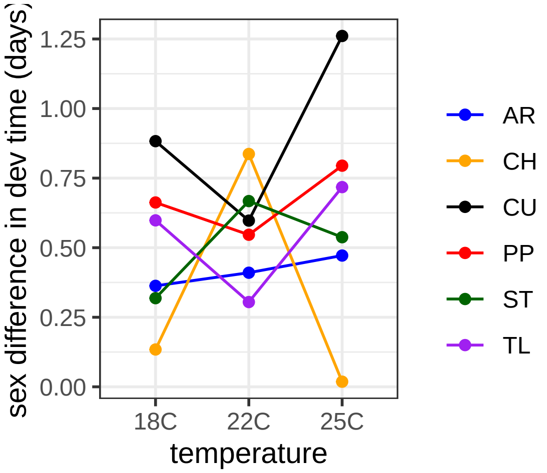


Notice how the sex difference is greatest for CH at the intermediate temperature (22C), while the sex difference is smallest for CU and TL at the intermediate temperature.

#### Graph the data

We will next make graphs of the development time data. To do, we define a color scheme for sexes, and then use `ggplot()` to make boxplots. We will plot the data points on top of the box plots. Each point represents a single fly.

```
sexcols <- c("female" = "magenta", "male" = "cornflowerblue")
ggplot(dev_temp, aes(x=arr, y=time, color=Sex)) +
    geom_boxplot(outlier.shape=NA)+
    geom_point(position=position_jitterdodge(), size=0.05, alpha=0.1)    +
    scale_color_manual(values = sexcols) +
    scale_y_continuous("developmental time (days)") +
    scale_x_discrete("") +
    facet_wrap(~temp, ncol=3) +
    theme_bw() +
    theme(legend.position = "top", legend.title=element_blank())
```

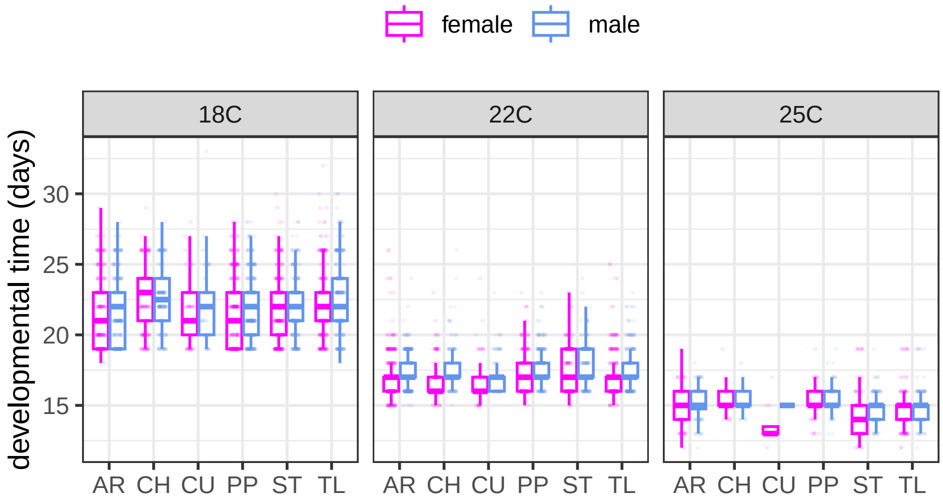


There are a lot of points in the plot, and each point is hard to see. We will therefore make a version of the boxplot without points shown.

```
ggplot(dev_temp, aes(x=arr, y=time, color=Sex)) +
    geom_boxplot(outlier.shape=NA)+
    scale_color_manual(values = sexcols) +
    scale_y_continuous("developmental time (days)") +
    scale_x_discrete("") +
    facet_wrap(~temp, ncol=3) +
    theme_bw() +
    theme(legend.position = "top", legend.title=element_blank())
```

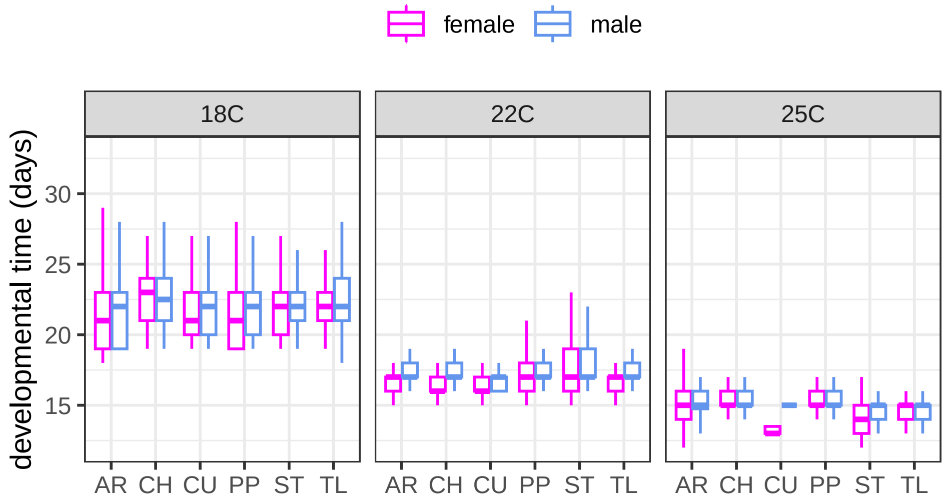


Looking at the boxplots, we can observed patterns that are consistent with the statistical analysis. First, the blue boxes tend to be higher than the pink, consistent with the statistical analysis that found males develop slower than females. The statistical tests additionally found that at 25°C, the sex difference is especially pronounced in flies with the CU arrangement, which can be seen in the graph. Second, the graphs clearly show that flies develop faster at warmer temperatures, and flies carrying the CU arrangement have the most accelerated development time as temperature increases. The acceleration of CU development time appears to be driven by faster female development.

An alternaitve way to represente these same data is with a ridgeplot.

```
dev_temp$ylabs = paste(dev_temp$arr, dev_temp$Sex, sep=" ")

ggplot(dev_temp, aes(y=ylabs, x=time, color=Sex, fill=Sex)) +
    geom_density_ridges(quantile_lines = TRUE, quantiles = 2) +
    scale_color_manual(values = sexcols) +
    scale_fill_manual(values = c("female"="pink", "male"="lightblue")) +
    scale_x_continuous("developmental time (days)") +
    scale_y_discrete("") +
    facet_wrap(~temp, ncol=3) +
    theme_bw() +
    theme(legend.position = "top", legend.title=element_blank())
```

```
## Picking joint bandwidth of 0.661

## Picking joint bandwidth of 0.265

## Picking joint bandwidth of 1.15
```

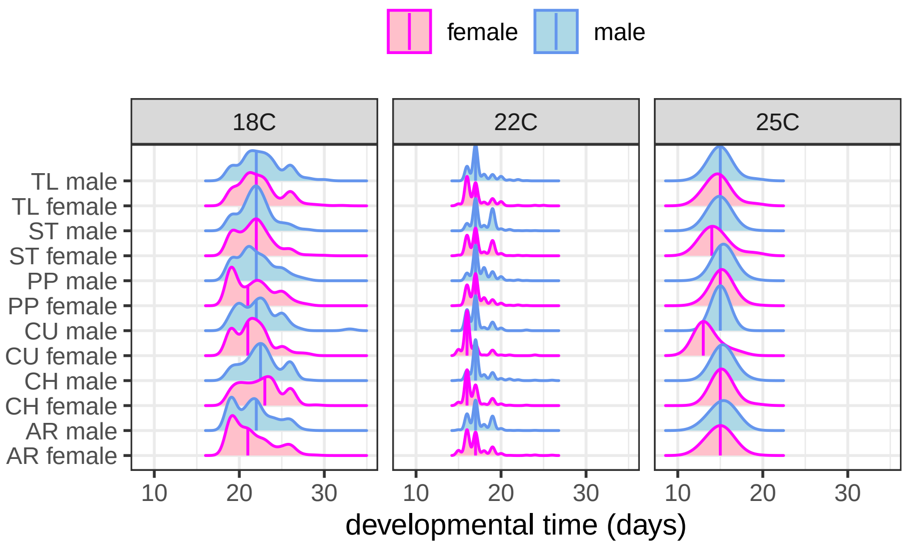


#### Compare development time and lifespan across strains

We will compare development time and lifespan across strains and arrangements, to test if they are correlated. First, we will load two different measures of the average lifespan for each strain: the median time and the mean time.

```
# load median lifespan
sex_aging_18 <- read.delim("sex_median_18.tsv")
sex_aging_22 <- read.delim("sex_median_22.tsv")
sex_aging_25 <- read.delim("sex_median_25.tsv")

# load mean lifespan
sex_agingInt_18 <- read.delim("sex_intercept_18.tsv")
sex_agingInt_22 <- read.delim("sex_intercept_22.tsv")
sex_agingInt_25 <- read.delim("sex_intercept_25.tsv")
```

Next, we calculate median developmental time for each strain at each temperature. The following code defines a function to calculate the median development time, and then it invokes the function for data from each temperature. We calculate the median for each sex separately from each strain at each temperature.

```
sex.median <- function(datas){
    strains <- levels(factor(datas$Strain))
    female.df <- data.frame(
        strain = strains,
        median = NA,
        obs = NA
    )
    male.df <- female.df

    for(i in 1:length(strains)){
        female.df$median[i] <- median( # calculate the median for females
          subset(datas, Strain==strains[i] & Sex=="female")$time)
        female.df$obs[i] <- length( # count the number of observations per strain
          subset(datas, Strain==strains[i] & Sex=="female")[,1])
    }
    female.df$sex <- "female"
```

```
    for(i in 1:length(strains)){
        male.df$median[i] <- median( # calculate the median for males
          subset(datas, Strain==strains[i] & Sex=="male")$time)
        male.df$obs[i] <- length( # count the number of observations for each strain
          subset(datas, Strain==strains[i] & Sex=="male")[,1])
    }
    male.df$sex <- "male"
    rbind(female.df, male.df)
}
sex_dev_18 <- sex.median(dev_18)
sex_dev_22 <- sex.median(dev_22)
sex_dev_25 <- sex.median(dev_25)
```

Next, we will use the intercept of a linear model to calculate the mean development time for each strain. We do this for each sex from each strain at each temperature. The linear model includes the date laid (batch) as a random effect. As above, we will define a function to perform the calculations, and then invoke the function for data from each temperature.

```
sex.intercept <- function(datas){
  # get an array of all strains
    strains <- levels(factor(datas$Strain))

    # make a data frame for females
    female.df <- data.frame(
        strain = strains,
        effect = NA,
        obs = NA
    )

    # copy the data frame from females to make the male version
    male.df <- female.df

    for(i in 1:length(strains)){
      # calcular mean development time for females for each strain
      # only consider if have >9 samples per sex from strain
        if(length(subset(datas, Strain==strains[i] & Sex=="female")[,1]) > 9){
            if(length(levels(factor( # if there are more than one egg laying dates
              subset(datas, Sex=="female" & Strain==strains[i])$Date.Laid))) > 1){
                # calculate the effect using batch as a random effect
              female.df$effect[i] <- summary(
                lmer(time ~ 1 + (1|Date.Laid),
                     data=subset(datas, Strain==strains[i] & Sex=="female"))
                )$coefficients[1]
            }else{ # if there is only one egg laying date
                # calculate the effect without a random effect
                female.df$effect[i] <- summary(
                  lm(time ~ 1, data=subset(datas, Strain==strains[i] & Sex=="female"))
                  )$coefficients[1]
            }
        } else{ # if <10 females from strain
            female.df$effect[i] <- NA
        }
      # count the number of observations for females per strain
        female.df$obs[i] <- length(subset(datas, Strain==strains[i] & Sex=="female")[,1])
```

```
    }
    female.df$sex <- "female"

    for(i in 1:length(strains)){
      # calcular mean development time for males for each strain
      # only consider if have >9 samples per sex from strain
        if(length(subset(datas, Strain==strains[i] & Sex=="male")[,1]) > 9){
            if(length(levels(factor( # if there are more than one egg laying date
              subset(datas, Sex=="male" & Strain==strains[i])$Date.Laid))) > 1){
                male.df$effect[i] <- summary(
                  lmer(time ~ 1 + (1|Date.Laid),
                       data=subset(datas, Strain==strains[i] & Sex=="male"))
                  )$coefficients[1]
            } else{ # if there is only one egg laying date
                # calculate the effect without a random effect
                male.df$effect[i] <- summary(
                  lm(time ~ 1, data=subset(datas, Strain==strains[i] & Sex=="male"))
                  )$coefficients[1]
            }
        } else{ # if <10 males from strain
            male.df$effect[i] <- NA
        }
      # count the number of observations for males per strain
        male.df$obs[i] <- length(subset(datas, Strain==strains[i] & Sex=="male")[,1])
    }
    male.df$sex <- "male"

    rbind(female.df, male.df)
}
sex_intercept_18 <- sex.intercept(dev_18)
sex_intercept_22 <- sex.intercept(dev_22)
sex_intercept_25 <- sex.intercept(dev_25)
```

Write these average development time data to file so that they can be used elsewhere if desired.

```
write.table(sex_dev_18, "sex_dev_18.tsv", quote=FALSE, sep="\t", row.names=FALSE)
write.table(sex_dev_22, "sex_dev_22.tsv", quote=FALSE, sep="\t", row.names=FALSE)
write.table(sex_dev_25, "sex_dev_25.tsv", quote=FALSE, sep="\t", row.names=FALSE)

write.table(sex_intercept_18, "sex_dev_intercept_18.tsv", quote=FALSE, sep="\t", row.names=FALSE)
write.table(sex_intercept_22, "sex_dev_intercept_22.tsv", quote=FALSE, sep="\t", row.names=FALSE)
write.table(sex_intercept_25, "sex_dev_intercept_25.tsv", quote=FALSE, sep="\t", row.names=FALSE)
```

The next code block is a function that makes graphs comparing lifespan and development time at each temperature. The function also returns a data frame with the lifespan and development time data for each strain in a single object. Send the function the lifespan data, development time data, and an object containing the strains used in the experiments.

```
aging.dev <- function(aging, dev, strains){
    arrcols <- c( "AR"="blue", "CH"="orange", "CU"="black",
                  "PP"="red", "ST"="darkgreen", "TL"="purple")

    female.df <- data.frame(
        strain = intersect(
          subset(aging, sex=="female")$strain,
```

```
      subset(dev, sex=="female")$strain),
    sex = "female",
    devtime = NA,
    survival = NA
)
female.df$devtime <- subset(dev, sex=="female")[
  match(female.df$strain,
        subset(dev, sex=="female")$strain),
  2]
female.df$survival <- subset(aging, sex=="female")[
  match(female.df$strain,
        subset(aging, sex=="female")$strain),
  2]
female.df$arr <- strains$arrangement[match(female.df$strain, strains$ID)]

male.df <- data.frame(
    strain = intersect(
      subset(aging, sex=="male")$strain,
      subset(dev, sex=="male")$strain),
    sex = "male",
    devtime = NA,
    survival = NA
)
male.df$devtime <- subset(dev, sex=="male")[
  match(male.df$strain,
        subset(dev, sex=="male")$strain),
  2]
male.df$survival <- subset(aging, sex=="male")[
  match(male.df$strain,
        subset(aging, sex=="male")$strain),
  2]
male.df$arr <- strains$arrangement[match(male.df$strain, strains$ID)]

print(
    ggplot(na.omit(rbind(female.df, male.df)),
           aes(x=devtime, y=survival, color=arr)) +
    geom_point() +
    scale_color_manual(values=arrcols) +
    scale_x_continuous("egg-to-adult development time (days)") +
    scale_y_continuous("lifespan (days)") +
    facet_wrap(~sex, ncol=2) +
    theme_bw() +
    theme(legend.title=element_blank())
)
na.omit(rbind(female.df, male.df))
}
```

Call the function to make graphs comparing lifespan and development time. Consider both the mean (intercept of model) and median measures separately. Analyze data from each temperature separately. Here are the three graphs for median time.

```
agingMed_devMed_18 <- aging.dev(sex_aging_18, sex_dev_18, strainIDs)
```

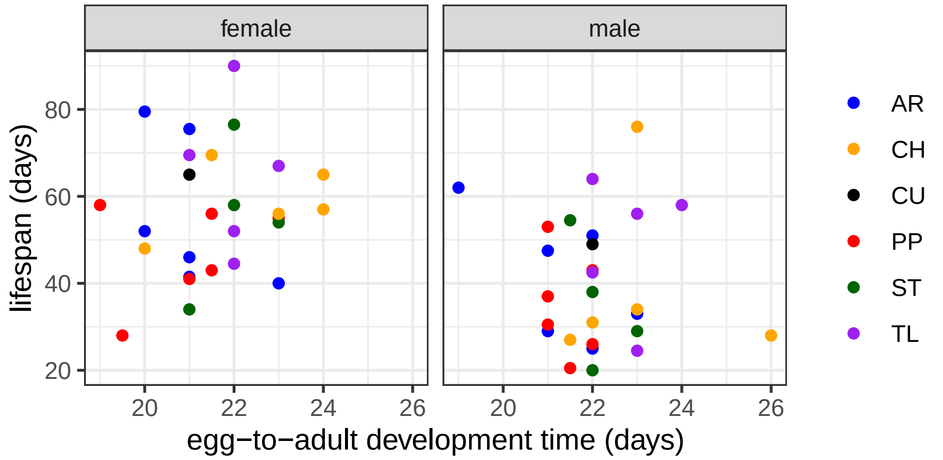


```r
agingMed_devMed_22 <- aging.dev(sex_aging_22, sex_dev_22, strainIDs)
```

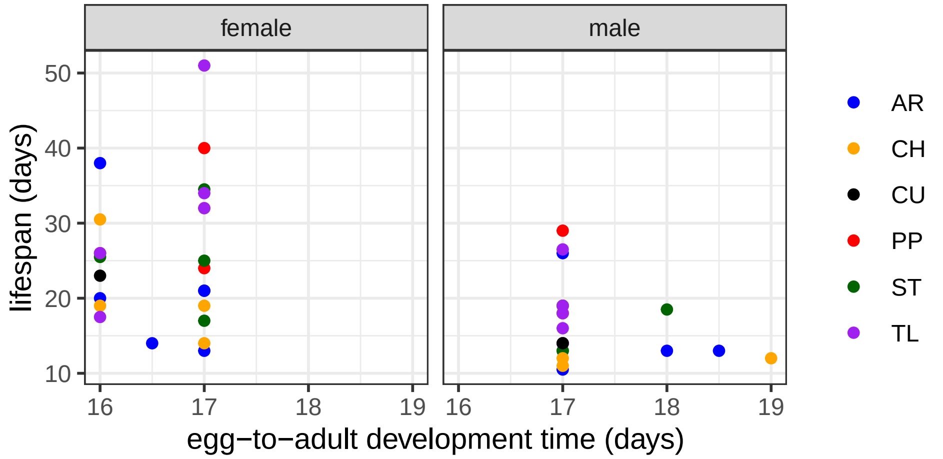


```r
agingMed_devMed_25 <- aging.dev(sex_aging_25, sex_dev_25, strainIDs)
```

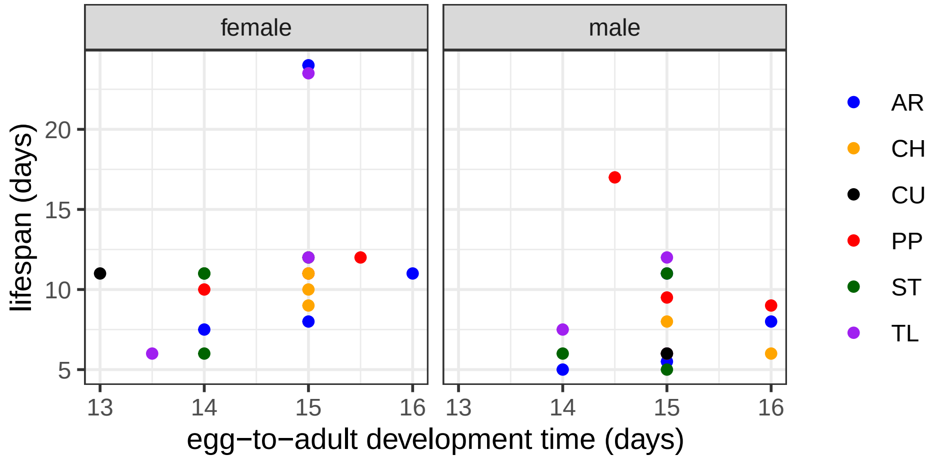


And here are the three graphs for mean time.

```r
agingInt_devInt_18 <- aging.dev(sex_agingInt_18, sex_intercept_18, strainIDs)
```

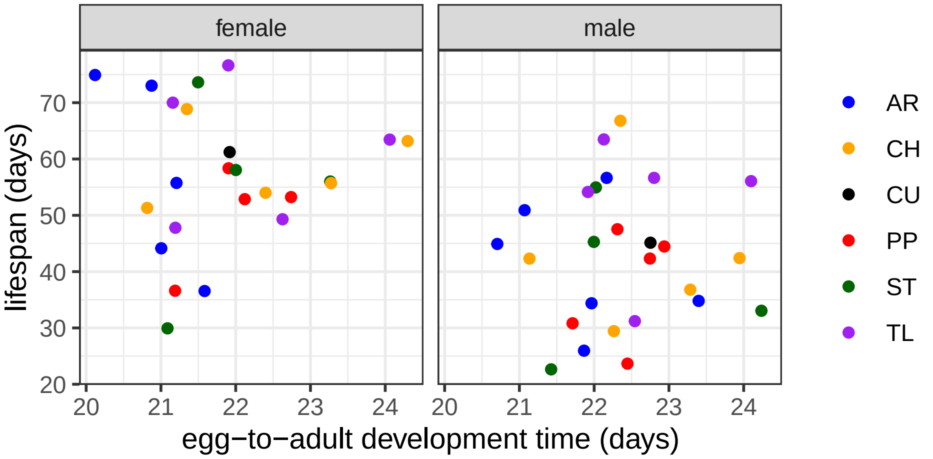


```
agingInt_devInt_22 <- aging.dev(sex_agingInt_22, sex_intercept_22, strainIDs)
```

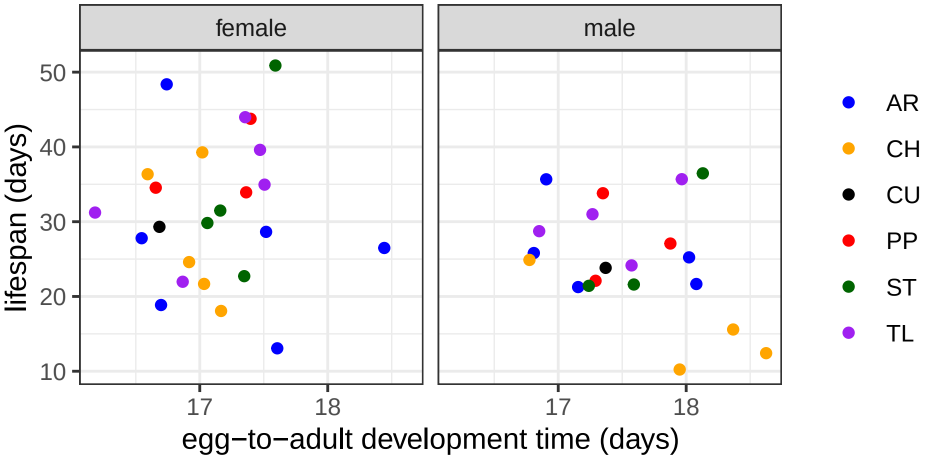


```
agingInt_devInt_25 <- aging.dev(sex_agingInt_25, sex_intercept_25, strainIDs)
```

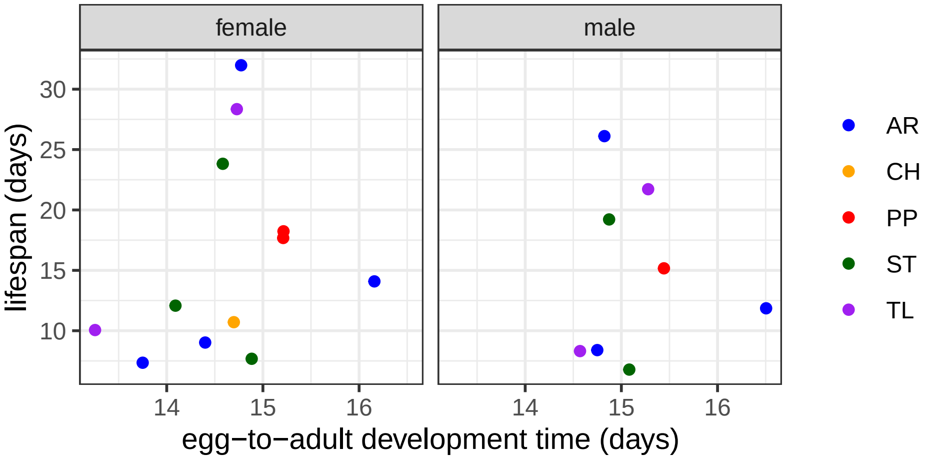


Make single graph for all temperatures, using the mean lifespan and development time for each strain. To do so, first make a single dataframe with mean times at all three temperatures. Then, make a graph using that data frame.

```
agingInt_devInt <- rbind(
    cbind(agingInt_devInt_18, temp="18C"),
```

```r
    cbind(agingInt_devInt_22, temp="22C"),
    cbind(agingInt_devInt_25, temp="25C")
)

arrcols <- c( "AR"="blue", "CH"="orange", "CU"="black",
              "PP"="red", "ST"="darkgreen", "TL"="purple")
ggplot(agingInt_devInt, aes(x=devtime, y=survival, color=arr)) +
    geom_point() +
    scale_color_manual(values=arrcols) +
    scale_x_continuous("egg-to-adult development time (days)") +
    scale_y_continuous("lifespan (days)") +
    facet_grid(sex~temp) +
    theme_bw() +
    theme(legend.title=element_blank())
```

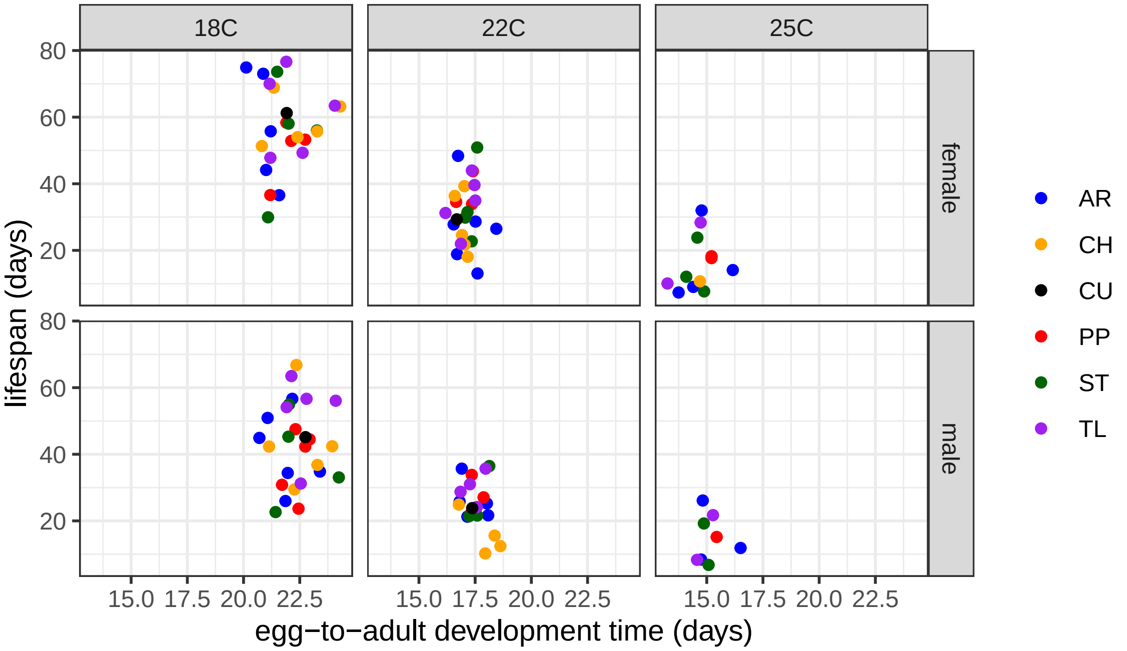


Lastly, we will perform statistical tests to determine if there is a correlation between mean development time and mean lifespan across arrangements. We will consider lifespan as the response, and development time as a fixed effect, along with sex and arrangement. Therefore, each strain within arrangement is treated as a replicate measure of lifespan for that arrangement. We will do this separately at each temperature.

```r
anova(lm(survival ~ devtime + sex + arr, data= agingInt_devInt_18))
```

```
## Analysis of Variance Table
##
## Response: survival
##           Df Sum Sq Mean Sq F value    Pr(>F)
## devtime    1  106.5  106.48  0.6986 0.4079959
## sex        1 2309.7 2309.74 15.1532 0.0003486 ***
## arr        5  919.5  183.90  1.2065 0.3228151
## Residuals 42 6401.9  152.43
## ---
## Signif. codes:  0 '***' 0.001 '**' 0.01 '*' 0.05 '.' 0.1 ' ' 1
```

```r
anova(lm(survival ~ devtime + sex + arr, data= agingInt_devInt_22))
```

```
## Analysis of Variance Table
##
## Response: survival
##           Df  Sum Sq Mean Sq F value  Pr(>F)
## devtime    1  227.92 227.917  3.2404 0.08023 .
## sex        1  267.98 267.980  3.8100 0.05876 .
## arr        5  621.76 124.352  1.7679 0.14437
## Residuals 36 2532.13  70.337
## ---
## Signif. codes:  0 '***' 0.001 '**' 0.01 '*' 0.05 '.' 0.1 ' ' 1
```

```r
anova(lm(survival ~ devtime + sex + arr, data= agingInt_devInt_25))
```

```
## Analysis of Variance Table
##
## Response: survival
##           Df Sum Sq Mean Sq F value Pr(>F)
## devtime    1  23.84  23.841  0.3121 0.5859
## sex        1  22.62  22.623  0.2961 0.5955
## arr        4  71.05  17.764  0.2325 0.9151
## Residuals 13 993.08  76.391
```

Sex is a significant predictor of lifespan at 18°C, and both sex and development time are almost significantly associated with life span at 22°C. Extract correlation of development time and lifespan at 22°C.

```r
lm(survival ~ devtime + sex + arr, data= agingInt_devInt_22)
```

```
##
## Call:
## lm(formula = survival ~ devtime + sex + arr, data = agingInt_devInt_22)
##
## Coefficients:
## (Intercept)      devtime      sexmale        arrCH        arrCU        arrPP
##     61.7892      -1.8829      -5.6219      -3.9974      -0.3494       6.1728
##       arrST        arrTL
##      4.0963       5.5102
```

As an alternative approach, we will will consider development time, sex, and arrangement as interacting predictors of lifespan.

```r
anova(lm(survival ~ devtime*sex*arr, data= agingInt_devInt_18))
```

```
## Analysis of Variance Table
##
## Response: survival
##                 Df Sum Sq Mean Sq F value   Pr(>F)
## devtime          1  106.5  106.48  0.5764 0.454084
## sex              1 2309.7 2309.74 12.5025 0.001436 **
## arr              5  919.5  183.90  0.9954 0.438460
## devtime:sex      1    2.4    2.40  0.0130 0.910049
## devtime:arr      5  569.5  113.91  0.6166 0.688139
## sex:arr          4  143.8   35.95  0.1946 0.939168
## devtime:sex:arr  4  513.4  128.34  0.6947 0.601932
## Residuals       28 5172.8  184.74
## ---
```

```
## Signif. codes:  0 '***' 0.001 '**' 0.01 '*' 0.05 '.' 0.1 ' ' 1
```

```
anova(lm(survival ~ devtime*sex*arr, data= agingInt_devInt_22))
```

```
## Analysis of Variance Table
##
## Response: survival
##                 Df  Sum Sq Mean Sq F value  Pr(>F)
## devtime          1  227.92 227.917  3.0124 0.09662 .
## sex              1  267.98 267.980  3.5419 0.07313 .
## arr              5  621.76 124.352  1.6436 0.19030
## devtime:sex      1   23.12  23.117  0.3055 0.58600
## devtime:arr      5  519.07 103.814  1.3721 0.27280
## sex:arr          4  218.46  54.614  0.7218 0.58626
## devtime:sex:arr  4  106.96  26.740  0.3534 0.83884
## Residuals       22 1664.52  75.660
## ---
## Signif. codes:  0 '***' 0.001 '**' 0.01 '*' 0.05 '.' 0.1 ' ' 1
```

```
anova(lm(survival ~ devtime*sex*arr, data= agingInt_devInt_25))
```

```
## Analysis of Variance Table
##
## Response: survival
##                 Df Sum Sq Mean Sq F value Pr(>F)
## devtime          1  23.84  23.841  0.1456 0.7222
## sex              1  22.62  22.623  0.1382 0.7290
## arr              4  71.05  17.764  0.1085 0.9731
## devtime:sex      1  26.29  26.295  0.1606 0.7091
## devtime:arr      3  71.88  23.959  0.1463 0.9269
## sex:arr          3 157.78  52.594  0.3212 0.8111
## devtime:sex:arr  2  82.17  41.085  0.2509 0.7895
## Residuals        4 654.96 163.739
```

Including the predictors as interaction terms does not affect our results.

As another alternative approach, we will consider data from all three temperatures at once. For this analysis, we treat development time, sex, arrangement, temperature, and their four-way interaction as predictors of lifespan.

```
anova(lm(survival ~ devtime*sex*arr*temp, data= agingInt_devInt))
```

```
## Analysis of Variance Table
##
## Response: survival
##                      Df  Sum Sq Mean Sq  F value    Pr(>F)
## devtime               1 17884.1 17884.1 128.8985 6.331e-16 ***
## sex                   1  3212.4  3212.4  23.1534 1.243e-05 ***
## arr                   5   850.7   170.1   1.2263   0.30955
## temp                  2  1279.6   639.8   4.6113   0.01416 *
## devtime:sex           1   668.5   668.5   4.8179   0.03248 *
## devtime:arr           5   622.6   124.5   0.8975   0.48950
## sex:arr               5   127.2    25.4   0.1833   0.96772
## devtime:temp          2    63.6    31.8   0.2294   0.79582
## sex:temp              2     8.3     4.2   0.0300   0.97047
## arr:temp              9  1090.0   121.1   0.8729   0.55473
## devtime:sex:arr       4    68.8    17.2   0.1240   0.97323
```

```
## devtime:sex:temp      2    38.6    19.3   0.1390   0.87054
## devtime:arr:temp      7   533.7    76.2   0.5496   0.79295
## sex:arr:temp          7   446.2    63.7   0.4594   0.85936
## devtime:sex:arr:temp  6   354.3    59.0   0.4255   0.85876
## Residuals            54  7492.3   138.7
## ---
## Signif. codes:  0 '***' 0.001 '**' 0.01 '*' 0.05 '.' 0.1 ' ' 1
```

When we consider all three temperatures together, sex, temperature, and development time are significant predictors of lifespan, as is the development time by sex interactions. We can extract the effects of these predictors from a model. First, we will test which effect is significant when we consider a model with only the significant interaction identified above.

```
anova(lm(survival ~ sex + temp + devtime + arr + devtime*sex, data= agingInt_devInt))
```

```
## Analysis of Variance Table
##
## Response: survival
##              Df  Sum Sq Mean Sq  F value    Pr(>F)
## sex           1  1148.6  1148.6  10.9080  0.001317 **
## temp          2 21292.7 10646.3 101.1074 < 2.2e-16 ***
## devtime       1     1.0     1.0   0.0097  0.921903
## arr           5   784.6   156.9   1.4902  0.199448
## sex:devtime   1   668.5   668.5   6.3484  0.013285 *
## Residuals   103 10845.6   105.3
## ---
## Signif. codes:  0 '***' 0.001 '**' 0.01 '*' 0.05 '.' 0.1 ' ' 1
```

Here, we see that sex, temperature, and the sex by development time interaction are all significant predictors of lifespan. We can extract the effects of each predictor like so:

```
lm(survival ~ sex + temp + devtime + arr + devtime*sex, data= agingInt_devInt)
```

```
##
## Call:
## lm(formula = survival ~ sex + temp + devtime + arr + devtime *
##     sex, data = agingInt_devInt)
##
## Coefficients:
##     (Intercept)          sexmale          temp22C          temp25C
##         35.6815          22.2422         -21.3185         -34.0307
##         devtime            arrCH            arrCU            arrPP
##          0.9266          -1.4117           1.9095          -0.3193
##           arrST            arrTL  sexmale:devtime
##         -0.0122           5.8466          -1.6365
```

These results show that flies live shorter at warmer temperatures. The effect is quite strong, with flies at 22°C living 21 days shorter than flies at 18°C, and flies at 25°C living 34 days shorter than at 18°C. Therefore, each 25°C causes flies to live ~5 days shorter. In addition, when we consider development time, males live significantly longer than females, by an average of >22 days.

Development time is positively correlated with lifespan, with each day of egg-to-adult-development adding almost a full day to lifespan. However, the negative interaction between sex and development time demonstrates that the effect is reversed for males, i.e., development time and lifespan is only positively correlated for female *D. pseudoobscura*.

The results above show that there are sex differences in the relationships between development time and lifespan. We will therefore analyze sex-specific models relating development time, temperature, and arrangement with lifespan. First, here is the model for females.

```
anova(
  lm(survival ~ temp + arr + devtime,
   data= subset(agingInt_devInt, sex=="female"))
)
```

```
## Analysis of Variance Table
##
## Response: survival
##           Df  Sum Sq Mean Sq F value    Pr(>F)
## temp       2 15444.0  7722.0 62.7085 1.804e-14 ***
## arr        5   232.8    46.6  0.3781    0.8615
## devtime    1    29.5    29.5  0.2399    0.6264
## Residuals 51  6280.2   123.1
## ---
## Signif. codes:  0 '***' 0.001 '**' 0.01 '*' 0.05 '.' 0.1 ' ' 1
```

In females, only temperature, and not development time, is significantly associated with lifespan.

```
lm(survival ~ temp + arr + devtime,
   data= subset(agingInt_devInt, sex=="female"))
```

```
##
## Call:
## lm(formula = survival ~ temp + arr + devtime, data = subset(agingInt_devInt,
##     sex == "female"))
##
## Coefficients:
## (Intercept)      temp22C      temp25C        arrCH        arrCU        arrPP
##     35.1540     -21.1231     -33.9932       0.2262       2.9601       0.9265
##       arrST        arrTL      devtime
##      1.0717       5.2657       0.9178
```

In females, the association between temperature and lifespan is negative, consistent with what we saw when we analyzed lifespan data previously.

Next, we can examine the relationships in males.

```
anova(
  lm(survival ~ temp + arr + devtime,
   data= subset(agingInt_devInt, sex=="male"))
)
```

```
## Analysis of Variance Table
##
## Response: survival
##           Df Sum Sq Mean Sq F value    Pr(>F)
## temp       2 6513.0  3256.5 32.7257 1.682e-09 ***
## arr        5  601.8   120.4  1.2095    0.3202
## devtime    1   13.1    13.1  0.1318    0.7183
## Residuals 45 4477.9    99.5
## ---
## Signif. codes:  0 '***' 0.001 '**' 0.01 '*' 0.05 '.' 0.1 ' ' 1
```

In males, only temperature, and not development time, is significantly associated with lifespan. This is the same as what we saw in females.

```r
lm(survival ~ temp + arr + devtime,
   data= subset(agingInt_devInt, sex=="male"))
```

```
##
## Call:
## lm(formula = survival ~ temp + arr + devtime, data = subset(agingInt_devInt,
##     sex == "male"))
##
## Coefficients:
## (Intercept)      temp22C      temp25C        arrCH        arrCU        arrPP
##     58.5504     -21.5221     -34.3139      -3.3692       0.7789      -1.6267
##       arrST        arrTL      devtime
##     -1.2393       6.4892      -0.7022
```

In males, the association between temperature and lifespan is negative, consistent with what we saw in females and when we analyzed lifespan data previously.

Recall that, when we analyzed data from each temperature separately, the association between development time and lifespan was only observed at 22°C. We can extract the effects from the model for the 22°C data to see if they are consistent with the analysis from all temperatures.

```r
lm(survival ~ devtime + sex + arr, data= agingInt_devInt_22)
```

```
##
## Call:
## lm(formula = survival ~ devtime + sex + arr, data = agingInt_devInt_22)
##
## Coefficients:
## (Intercept)      devtime      sexmale        arrCH        arrCU        arrPP
##     61.7892      -1.8829      -5.6219      -3.9974      -0.3494       6.1728
##       arrST        arrTL
##      4.0963       5.5102
```

In contrast to what we observed across all temperatures, at 22°C, there is a negative correlation between development time and lifespan. In addition, males live shorter than females at 22°C. We therefore conclude that, while there may be associations between development time and lifespan, they are not consistent across temperatures or between sexes.

Lastly, we will test for male-female correlations in development time and lifespan. We will create a new version of the objects so that we can extract compare between sexes from each strain.

```r
AgingDevIntSex_18 <- pivot_wider(agingInt_devInt_18,
            names_from=sex, values_from=c(devtime, survival))
AgingDevIntSex_22 <- pivot_wider(agingInt_devInt_22,
            names_from=sex, values_from=c(devtime, survival))
AgingDevIntSex_25 <- pivot_wider(agingInt_devInt_25,
            names_from=sex, values_from=c(devtime, survival))
AgingDevIntSex_temp <- pivot_wider(agingInt_devInt,
            names_from=sex, values_from=c(devtime, survival))
```

Test for male:female correlations using a linear model with male development or lifespan as a response and female development or lifespan as a predictor, along with arrangement as a predictor (and temp when considering all temperatures).

```r
anova(lm(devtime_male ~ devtime_female + arr, data=AgingDevIntSex_18))
```

```
## Analysis of Variance Table
##
```

```
## Response: devtime_male
##                 Df  Sum Sq Mean Sq F value    Pr(>F)
## devtime_female   1 14.2464 14.2464  64.744 3.374e-07 ***
## arr              5  0.5237  0.1047   0.476    0.7892
## Residuals       17  3.7407  0.2200
## ---
## Signif. codes:  0 '***' 0.001 '**' 0.01 '*' 0.05 '.' 0.1 ' ' 1
```

```r
anova(lm(devtime_male ~ devtime_female + arr, data=AgingDevIntSex_22))
```

```
## Analysis of Variance Table
##
## Response: devtime_male
##                 Df Sum Sq Mean Sq F value   Pr(>F)
## devtime_female   1 1.9427 1.94268 10.8191 0.005868 **
## arr              5 1.3466 0.26933  1.4999 0.256250
## Residuals       13 2.3343 0.17956
## ---
## Signif. codes:  0 '***' 0.001 '**' 0.01 '*' 0.05 '.' 0.1 ' ' 1
```

```r
anova(lm(devtime_male ~ devtime_female + arr, data=AgingDevIntSex_25))
```

```
## Analysis of Variance Table
##
## Response: devtime_male
##                 Df  Sum Sq Mean Sq F value  Pr(>F)
## devtime_female   1 2.00146 2.00146 13.1001 0.03626 *
## arr              3 0.16123 0.05374  0.3518 0.79315
## Residuals        3 0.45835 0.15278
## ---
## Signif. codes:  0 '***' 0.001 '**' 0.01 '*' 0.05 '.' 0.1 ' ' 1
```

```r
anova(lm(devtime_male ~ devtime_female + arr + temp, data=AgingDevIntSex_temp))
```

```
## Analysis of Variance Table
##
## Response: devtime_male
##                 Df Sum Sq Mean Sq   F value    Pr(>F)
## devtime_female   1 439.80  439.80 2385.1206 < 2.2e-16 ***
## arr              5   0.47    0.09    0.5097  0.767363
## temp             2   2.82    1.41    7.6450  0.001443 **
## Residuals       43   7.93    0.18
## ---
## Signif. codes:  0 '***' 0.001 '**' 0.01 '*' 0.05 '.' 0.1 ' ' 1
```

In all three temperatures and when considering the entire data set, there is a significant association between male and female development time. We can extract the effects from each model.

```r
DevIntSex_temp_corr <- data.frame(
  temp = c("18C", "22C", "25C", "all"),
  r_mf = c(
    coefficients(lm(devtime_male ~ devtime_female + arr, data=AgingDevIntSex_18))[2],
    coefficients(lm(devtime_male ~ devtime_female + arr, data=AgingDevIntSex_22))[2],
    coefficients(lm(devtime_male ~ devtime_female + arr, data=AgingDevIntSex_25))[2],
    coefficients(lm(devtime_male ~ devtime_female + arr, data=AgingDevIntSex_temp))[2]
  )
)
```

```r
DevIntSex_temp_corr
```

```
##   temp      r_mf
## 1  18C 0.7370984
## 2  22C 0.7439961
## 3  25C 0.7814284
## 4  all 0.9726493
```

All correlations are >0.7! That's pretty strong inter-sexual correlation for development time. We can do the same for lifespan.

```r
anova(lm(survival_male ~ survival_female + arr, data=AgingDevIntSex_18))
```

```
## Analysis of Variance Table
##
## Response: survival_male
##                 Df  Sum Sq Mean Sq F value   Pr(>F)
## survival_female  1 1286.68 1286.68 11.1068 0.003942 **
## arr              5  237.69   47.54  0.4104 0.835002
## Residuals       17 1969.38  115.85
## ---
## Signif. codes:  0 '***' 0.001 '**' 0.01 '*' 0.05 '.' 0.1 ' ' 1
```

```r
anova(lm(survival_male ~ survival_female + arr, data=AgingDevIntSex_22))
```

```
## Analysis of Variance Table
##
## Response: survival_male
##                 Df Sum Sq Mean Sq F value    Pr(>F)
## survival_female  1 720.58  720.58 57.9819 3.823e-06 ***
## arr              5 134.91   26.98  2.1711    0.1208
## Residuals       13 161.56   12.43
## ---
## Signif. codes:  0 '***' 0.001 '**' 0.01 '*' 0.05 '.' 0.1 ' ' 1
```

```r
anova(lm(survival_male ~ survival_female + arr, data=AgingDevIntSex_25))
```

```
## Analysis of Variance Table
##
## Response: survival_male
##                 Df Sum Sq Mean Sq  F value   Pr(>F)
## survival_female  1 339.80  339.80 181.4016 0.000885 ***
## arr              3   6.00    2.00   1.0679 0.479109
## Residuals        3   5.62    1.87
## ---
## Signif. codes:  0 '***' 0.001 '**' 0.01 '*' 0.05 '.' 0.1 ' ' 1
```

```r
anova(lm(survival_male ~ survival_female + arr + temp, data=AgingDevIntSex_temp))
```

```
## Analysis of Variance Table
##
## Response: survival_male
##                 Df Sum Sq Mean Sq  F value Pr(>F)
## survival_female  1 8992.8  8992.8 168.3739 <2e-16 ***
## arr              5  195.5    39.1   0.7321 0.6034
## temp             2  112.0    56.0   1.0482 0.3594
## Residuals       43 2296.6    53.4
```

```
## ---
## Signif. codes:  0 '***' 0.001 '**' 0.01 '*' 0.05 '.' 0.1 ' ' 1
```

In all three temperatures and when considering the entire data set, there is a signficant association between male and female lifespan. We can extract the effects from each model.

```
AgingIntSex_temp_corr <- data.frame(
  temp = c("18C", "22C", "25C", "all"),
  r_mf = c(
    coefficients(lm(survival_male ~ survival_female + arr, data=AgingDevIntSex_18))[2],
    coefficients(lm(survival_male ~ survival_female + arr, data=AgingDevIntSex_22))[2],
    coefficients(lm(survival_male ~ survival_female + arr, data=AgingDevIntSex_25))[2],
    coefficients(lm(survival_male ~ survival_female + arr, data=AgingDevIntSex_temp))[2]
  )
)
AgingIntSex_temp_corr
```

```
##   temp      r_mf
## 1  18C 0.5494571
## 2  22C 0.5935527
## 3  25C 0.8004913
## 4  all 0.7211242
```

The correlations are strongest at 25C, but still >0.5 at all temperatures. Make figures showing the inter-sexual correlations.

```
intersex_surv_corr <- ggplot(AgingDevIntSex_temp, aes(x=survival_female, y=survival_male, color=arr)) +
    geom_point() +
    scale_color_manual(values=arrcols) +
    scale_x_continuous("female mean lifespan (days)") +
    scale_y_continuous("male mean lifespan (days)") +
    facet_grid(~temp) +
    theme_bw() +
    theme(legend.title=element_blank())
intersex_dev_corr <- ggplot(AgingDevIntSex_temp, aes(x=devtime_female, y=devtime_male, color=arr)) +
    geom_point() +
    scale_color_manual(values=arrcols) +
    scale_x_continuous("female development time (days)", breaks=c(14,17,20,23)) +
    scale_y_continuous("male development time (days)", breaks=c(14,17,20,23)) +
    facet_grid(~temp) +
    theme_bw() +
    theme(legend.title=element_blank())
plot_grid(intersex_surv_corr, intersex_dev_corr, ncol=1)
```

```
## Warning: Removed 10 rows containing missing values or values outside the scale range
## (`geom_point()`).
## Removed 10 rows containing missing values or values outside the scale range
## (`geom_point()`).
```

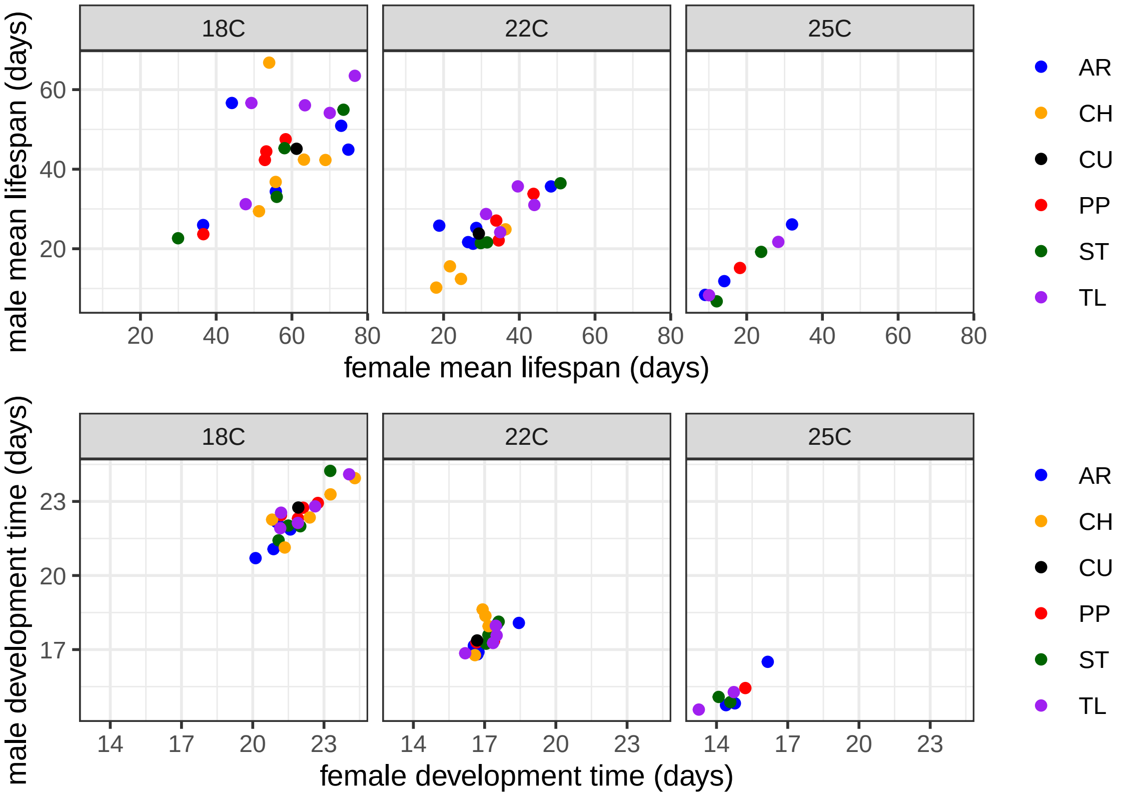
18C
22C
25C
male mean lifespan (days)
female mean lifespan (days)
male development time (days)
female development time (days)
AR
CH
CU
PP
ST
TL
20
40
60
80
14
17
20
23
