## Supplementary Figure S4 for "*Drosophila pseudoobscura* third chromosome inversion arrangements have temperature-dependent and sex-specific effects on life history traits"

### Supplementary File S4: Analyze body size data from *Drosophila pseudoobscura* at 22C

2026-09-03

#### Overview

The analysis described below was performed on body size measurements from *Drosophila pseudoobscura* collected in 2025. Males and females carrying one of six different third chromosome inversions arrangements were sampled. The flies were sampled from 28 different strains, each of which is homozygous for one of six different chromosomal inversion arrangements.

Five mated flies from a given were collected and stored in a vial with standard *Drosophila* medium for 1 day. Once the offspring began to emerge, we waited ~12-24 hours before collecting them and measuring their body size.

The body length of each fly was measured using an Olympus SZX10 stereomicroscope with the Olympus cellSens software.

#### Install and Load Required Packages

```
if (!require("ggplot2")) install.packages("ggplot2")
if (!require("outliers")) install.packages("outliers")
if (!require("lme4")) install.packages("lme4")
if (!require("nlme")) install.packages("nlme")

library(ggplot2)
library(outliers)
library(lme4)
library(nlme)
```

#### Load, Prepare, and QC the Data

Load body size measurements.

```
bodysize <- read.delim("Dpse-body-Gabe_2025_07_31.tsv")
```

Confirm that all dates are formatted the same. The flies were collected from vials that were established at different dates. First, check if the vial setup dates are formatted the same.

```
vialsetup <- levels(factor(bodysize$vial.setup))
vialsetup
```

```
##  [1] "11-Jun-25" "13-Jun-25" "16-Jun-25" "18-Jun-25" "2-Jul-25"  "2-Jun-25"
##  [7] "20-Jun-25" "23-Jun-25" "25-Jun-25" "27-Jun-25" "30-Jun-25" "30-May-25"
## [13] "4-Jul-25"  "4-Jun-25"  "6-Jun-25"  "9-Jun-25"
```

Everything looks good there.

```
emergence <- levels(factor(bodysize$Emergence))
emergence
```

```
##  [1] "1-Jul-25"  "10-Jul-25" "11-Jul-25" "14-Jul-25" "15-Jul-25" "16-Jul-25"
##  [7] "17-Jul-25" "17-Jun-25" "18-Jul-25" "18-Jun-25" "19-Jun-25" "2-Jul-25"
## [13] "20-Jun-25" "21-Jul-25" "22-Jul-25" "23-Jul-25" "23-Jun-25" "24-Jul-25"
## [19] "24-Jun-25" "25-Jul-25" "25-Jun-25" "26-Jun-25" "27-Jun-25" "28-Jul-25"
## [25] "29-Jul-25" "3-Jul-25"  "30-Jul-25" "30-Jun-25" "4-Jul-25"  "7-Jul-25"
## [31] "8-Jul-25"  "9-Jul-25"
```

All of the emergence dates look good.

Check the number of flies that from vials setup on each date. If any of these dates are singletons, they are probably errors that need to be corrected. Create a data frame with the number of flies per vial setup date.

```
vialsetup.df <- data.frame(
    date=vialsetup,
    count=NA
)
for(i in 1:length(vialsetup)){
    vialsetup.df$count[i] <- length(subset(bodysize, vial.setup==vialsetup[i])[,1])
}
vialsetup.df
```

```
##         date count
## 1  11-Jun-25    78
## 2  13-Jun-25   227
## 3  16-Jun-25   152
## 4  18-Jun-25    98
## 5   2-Jul-25    18
## 6   2-Jun-25   117
## 7  20-Jun-25   230
## 8  23-Jun-25    81
## 9  25-Jun-25   195
## 10 27-Jun-25   354
## 11 30-Jun-25   135
## 12 30-May-25   116
## 13  4-Jul-25    52
## 14  4-Jun-25   231
## 15  6-Jun-25   299
## 16  9-Jun-25   234
```

Everything looks good with the vial setup dates.

Next, check the dates when the flies emerged from pupae in the vials.
To do so, count the number of flies that emerged on each date. If any of these "birth dates'' are singletons, they are probably errors that need to be corrected. Create a data frame with the number of emergence per date.

```
emergence.df <- data.frame(
    date=emergence,
    count=NA
)
for(i in 1:length(emergence)){
    emergence.df$count[i] <- length(subset(bodysize, Emergence==emergence[i])[,1])
}
emergence.df
```

```
##         date count
## 1   1-Jul-25    60
```

```
## 2  10-Jul-25    77
## 3  11-Jul-25    42
## 4  14-Jul-25   207
## 5  15-Jul-25    92
## 6  16-Jul-25   100
## 7  17-Jul-25   111
## 8  17-Jun-25    15
## 9  18-Jul-25    81
## 10 18-Jun-25     5
## 11 19-Jun-25    62
## 12  2-Jul-25   124
## 13 20-Jun-25    18
## 14 21-Jul-25   127
## 15 22-Jul-25    16
## 16 23-Jul-25    11
## 17 23-Jun-25   190
## 18 24-Jul-25     6
## 19 24-Jun-25    74
## 20 25-Jul-25     6
## 21 25-Jun-25   130
## 22 26-Jun-25   165
## 23 27-Jun-25    97
## 24 28-Jul-25    11
## 25 29-Jul-25     2
## 26  3-Jul-25    79
## 27 30-Jul-25     3
## 28 30-Jun-25   266
## 29  4-Jul-25    80
## 30  7-Jul-25   236
## 31  8-Jul-25    56
## 32  9-Jul-25    68
```

All looks good on the emergence dates as well.

We will now load a file with the information for each strain. We will match the strain ID numbers from our data object with the information from the strain file. Lastly, we remove the flies from vials set up on July 2, 2025 because only 18 flies were collected from vials set up on that date.

```r
strainIDs <- read.delim("Dpse-strains.tsv")

# function to perform the matching
strain.assign <- function(strains, sizes){
    sizes$arr <- strains$arrangement[match(sizes$Strain, strains$ID)]
    sizes$isogen <- strains$isogen[match(sizes$Strain, strains$ID)]
    sizes$bal <- strains$balancer[match(sizes$Strain, strains$ID)]

    # Omit data with non-existent strain ID
    sizes <- subset(sizes, !is.na(arr))
}

# remove July 2, 2025 because only 18 flies from that date
bodysize <- subset(strain.assign(strainIDs, bodysize), vial.setup != "2-Jul-2025")
head(bodysize)
```

```
##   Strain    Sex vial.setup  len.um Emergence age Notes arr isogen bal
## 1      4   male  30-May-25 2833.36 17-Jun-25  18         AR DM1050   L
```

```
## 2       7   male  30-May-25 3012.69 17-Jun-25  18        AR  KB635   L
## 3      14 female  30-May-25 2696.18 17-Jun-25  18        AR MSH126   L
## 4      19 female  30-May-25 2859.37 17-Jun-25  18        PP DM1038   B
## 5      42 female  30-May-25 2482.82 17-Jun-25  18        CH MSH202   B
## 6      42   male  30-May-25 2439.15 17-Jun-25  18        CH MSH202   B
```

We now have our data with the body size measurement and strain information in single data frame. The `age` variable refers to the time between the vial setup date and the emergence date. The notes column in the data frame indicates if the fly emerged over the weekend, which means the `age` may not be an accurate measurement (because emergence may have occurred on Saturday or Sunday, but the fly was not collected until Monday).

#### Graph the data

Produce box plots showing the body size differences across arrangements. To do so, define a color scheme for male and female samples, then use `ggplot()` to create a boxplot.

```r
cols <- c("female" = "magenta", "male" = "cornflowerblue")

ggplot(bodysize, aes(x=arr, y=len.um, color=Sex)) +
    geom_boxplot(outlier.shape=NA) +
    geom_point(position=position_jitterdodge(), size=0.05, alpha=0.2)   +
    scale_color_manual(values = cols) +
    scale_y_continuous("body length (um)") +
    scale_x_discrete("") +
    theme_bw() +
    theme(legend.position = "top", legend.title=element_blank())
```

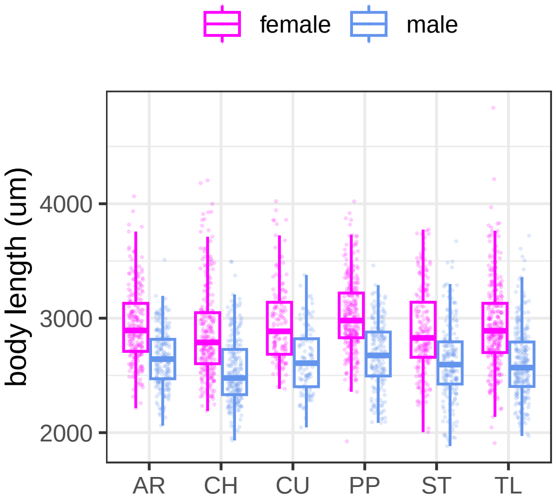


There is at least one individual who appears to be an outlier, with a body size much longer than all other flies. We will use a Grubbs test to perform a formal outlier analysis and decide if we need to exclude that fly.

```r
grubbs.test(bodysize$len.um)
```

```
##
##  Grubbs test for one outlier
##
## data:  bodysize$len.um
## G = 5.54506, U = 0.98824, p-value = 3.51e-05
## alternative hypothesis: highest value 4838.87 is an outlier
```

Grubbs test confirms that we have at least one outlier. However, this method only tests for a single outlier. To test for additional outliers, run Grubbs test iteratively, omitting each outlier that is detected. We will do this with a while loop, exiting when we fail to detect an outlier.

```
# Check for additional outliers
OUTLIERS <- TRUE # set an indicator variable to TRUE
bodysize.out <- bodysize # make a copy of the bodysize data for outlier detection

while(OUTLIERS==TRUE){ # while indicator is TRUE, continue to check
    grubby <- grubbs.test(bodysize.out$len.um) # run Grubbs test
    if(grubby$p.value < 0.05){ # if an outlier is found
        # identify the outlier as high or low
      if(grepl("highest", grubby$alternative)){
            # omit the high outlier
        bodysize.out <- subset(bodysize.out, len.um < max(bodysize.out$len.um))
        } else{
          # omit the low outlier
            bodysize.out <- subset(bodysize.out, len.um > min(bodysize.out$len.um))
        }
    } else{ # if no outlier found, reset indicator to FALSE to exit search
        OUTLIERS <- FALSE
    }
}
grubby # print final Grubbs test result
```

```
##
##  Grubbs test for one outlier
##
## data:  bodysize.out$len.um
## G = 3.88446, U = 0.99423, p-value = 0.1312
## alternative hypothesis: highest value 4215.29 is an outlier
```

The Grubbs test only detected one outlier. Our cleaned data, excluding the outlier, are stored in `bodysize.out`. We will make a new graph of the data with the one outlier excluded.

```
ggplot(bodysize.out, aes(x=arr, y= len.um/1000, color=Sex)) +
    geom_boxplot(outlier.shape=NA)+
    geom_point(position=position_jitterdodge(), size=0.05, alpha=0.2)   +
    scale_color_manual(values = cols) +
    scale_y_continuous("body length (mm)") +
    scale_x_discrete("") +
    theme_bw() +
    theme(legend.position = "top", legend.title=element_blank())
```

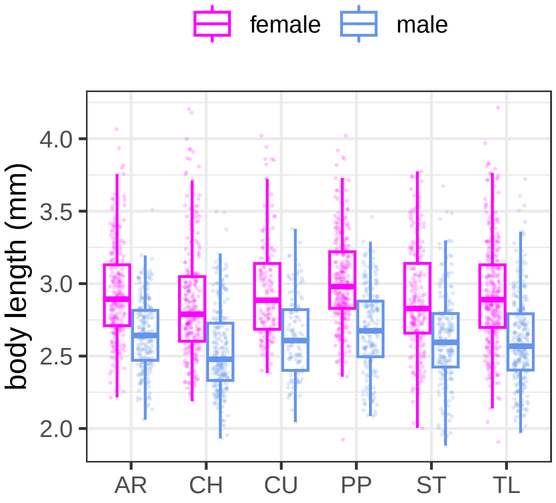


#### Statistical tests to determine which factors affect body size

We will perform ANOVA to test for the effect of arrangement and sex on body size. In the analysis, treat age and vial.setup date as random variables We will use the `lmer()` function from the `lme4` package to construct a mixed effects model. Our analysis will compare nested models to test if factors significantly improve model fit. We will first test if the interaction between sex and arrangement improves model fit.

```
# Define a function to compare models with and without an interaction term.
sex.arr.int.lmer <- function(datas){
    return.anova <- anova(
        lmer(len.um ~ Sex + arr + Sex*arr +
               (1|arr:isogen) + (1|vial.setup)  + (1|age),
             data=datas),
        lmer(len.um ~ Sex + arr +
               (1|arr:isogen) + (1|vial.setup) + (1|age),
             data=datas)
    )
    rownames(return.anova) <- c("model1", "model2")
    return.anova
}
sex.arr.int.lmer(bodysize.out)
```

```
## refitting model(s) with ML (instead of REML)

## Data: datas
## Models:
## lmer(len.um ~ Sex + arr + (1 | arr:isogen) + (1 | vial.setup) + (1 | age), data = datas): len.um ~ S
## lmer(len.um ~ Sex + arr + Sex * arr + (1 | arr:isogen) + (1 | vial.setup) + (1 | age), data = datas)
##        npar   AIC   BIC logLik -2*log(L) Chisq Df Pr(>Chisq)
## model1   11 36873 36938 -18426     36851
## model2   16 36881 36975 -18425     36849 1.833  5     0.8717
```

The model with the interaction term does not improve model fit, and we therefore conclude that there is not a signficant interaction between sex and arrangement.

We will next test if including sex in the model significantly improves the fit by comparing with a model where sex is not included.

```
sex.arr.lmer <- function(devdata){
    return.anova <- anova(
```

```
        lmer(len.um ~ Sex + arr + (1|arr:isogen) + (1|vial.setup)  + (1|age), data=devdata),
        lmer(len.um ~ arr + (1|arr:isogen) + (1|vial.setup) + (1|age), data=devdata)
    )
    rownames(return.anova) <- c("model1", "model2")
    return.anova
}
sex.arr.lmer(bodysize.out)
```

```
## refitting model(s) with ML (instead of REML)

## Data: devdata
## Models:
## lmer(len.um ~ arr + (1 | arr:isogen) + (1 | vial.setup) + (1 | age), data = devdata): len.um ~ arr +
## lmer(len.um ~ Sex + arr + (1 | arr:isogen) + (1 | vial.setup) + (1 | age), data = devdata): len.um ~
##        npar   AIC   BIC logLik -2*log(L)  Chisq Df Pr(>Chisq)
## model1   10 37488 37547 -18734     37468
## model2   11 36873 36938 -18426     36851 617.35  1  < 2.2e-16 ***
## ---
## Signif. codes:  0 '***' 0.001 '**' 0.01 '*' 0.05 '.' 0.1 ' ' 1
```

Including sex in the model does indeed improve model fit, demonstrating that there is a sex effect on body size.

Let's now compare models with and without arrangement as a predictor.

```
arr.sex.lmer <- function(devdata){
    return.anova <- anova(
        lmer(len.um ~ Sex + arr + (1|arr:isogen) + (1|vial.setup)  + (1|age), data=devdata),
        lmer(len.um ~ Sex + (1|arr:isogen) + (1|vial.setup) + (1|age), data=devdata)
    )
    rownames(return.anova) <- c("model1", "model2")
    return.anova
}
arr.sex.lmer(bodysize.out)
```

```
## refitting model(s) with ML (instead of REML)

## Data: devdata
## Models:
## lmer(len.um ~ Sex + (1 | arr:isogen) + (1 | vial.setup) + (1 | age), data = devdata): len.um ~ Sex +
## lmer(len.um ~ Sex + arr + (1 | arr:isogen) + (1 | vial.setup) + (1 | age), data = devdata): len.um ~
##        npar   AIC   BIC logLik -2*log(L) Chisq Df Pr(>Chisq)
## model1    6 36868 36903 -18428     36856
## model2   11 36873 36938 -18426     36851 4.763  5     0.4455
```

Arrangement does not improve model fit, which means that that there is no effect of arrangement on body size.

We will next use the `lme()` function from the `nlme` package to estimate the effects of the predictors that significantly improve the model fit.

```
anova(
    lme(fixed = len.um ~ Sex + arr + Sex*arr,
        data = bodysize.out,
        random = list(~1|isogen, ~1|vial.setup, ~1|age)
    )
)
```

```
##              numDF denDF   F-value p-value
## (Intercept)     1  1876 14204.139  <.0001
## Sex             1  1876   851.044  <.0001
## arr             5    21     0.692  0.6350
## Sex:arr         5  1876     0.743  0.5915
```

This analysis confirms that only sex has a significant effect on body size, and there is not a significant effect of the sex-by-arrangement interaction. We will therefore exclude the interaction term, and rerun `lme()`.

```
anova(
    lme(fixed = len.um ~ Sex + arr,
        data = bodysize.out,
        random = list(~1|isogen, ~1|vial.setup, ~1|age)
    )
)
```

```
##              numDF denDF   F-value p-value
## (Intercept)     1  1881 14175.930  <.0001
## Sex             1  1881   851.603  <.0001
## arr             5    21     0.691  0.6359
```

Once again, on sex effects body size, confirming what we found above.

We will therefore run our model with sex as the fixed effect, and isogen (strain), vial setup, and age as random effects. First, we will use `lmer()` to estimate the sex effect, coding isogen as a nested random effect within arrangement.

```
lmer(len.um ~ Sex + (1|arr:isogen) + (1|vial.setup) + (1|age), data=bodysize.out)
```

```
## Linear mixed model fit by REML ['lmerMod']
## Formula: len.um ~ Sex + (1 | arr:isogen) + (1 | vial.setup) + (1 | age)
##    Data: bodysize.out
## REML criterion at convergence: 36838.9
## Random effects:
##  Groups     Name        Std.Dev.
##  arr:isogen (Intercept) 111.61
##  age        (Intercept) 241.98
##  vial.setup (Intercept)  85.83
##  Residual               269.23
## Number of obs: 2615, groups:  arr:isogen, 27; age, 17; vial.setup, 16
## Fixed Effects:
## (Intercept)      Sexmale
##      2866.3       -282.8
```

Next, we will use `lme()` to perform the same analysis. This function can take isogen as a random effect, but cannot accept it as a nested within arrangement.

```
lme(fixed = len.um ~ Sex,
    data = bodysize.out,
    random = list(~1|isogen, ~1|vial.setup, ~1|age)
)
```

```
## Linear mixed-effects model fit by REML
##   Data: bodysize.out
##   Log-restricted-likelihood: -18341.42
##   Fixed: len.um ~ Sex
## (Intercept)     Sexmale
##   2935.4265   -287.0727
```

```
##
## Random effects:
##  Formula: ~1 | isogen
##         (Intercept)
## StdDev:    98.29588
##
##  Formula: ~1 | vial.setup %in% isogen
##         (Intercept)
## StdDev:   0.1108915
##
##  Formula: ~1 | age %in% vial.setup %in% isogen
##         (Intercept) Residual
## StdDev:    234.0785 219.8535
##
## Number of Observations: 2615
## Number of Groups:
##                         isogen          vial.setup %in% isogen
##                             27                             180
## age %in% vial.setup %in% isogen
##                            733
```

These two functions estimate similar effects of sex on body size: males are 282–287 $\mu$m smaller than females.

### Analyze with weekend emergence excluded

The data from flies that emerged over the weekend are not reliable estimates, as described above. We will therefore perform the same analysis as above, but with the weekend emergences excluded. First, create a copy of the data with the weekend emergence removed.

```
bodysize_wknd <- subset(bodysize.out, !grepl("weekend", Notes))
```

Let's now test if the sex-arrangement interaction improves model fit.

```
sex.arr.int.lmer(bodysize_wknd)
```

```
## refitting model(s) with ML (instead of REML)
```

```
## Data: datas
## Models:
## lmer(len.um ~ Sex + arr + (1 | arr:isogen) + (1 | vial.setup) + (1 | age), data = datas): len.um ~ Se
## lmer(len.um ~ Sex + arr + Sex * arr + (1 | arr:isogen) + (1 | vial.setup) + (1 | age), data = datas)
##        npar   AIC   BIC logLik -2*log(L)  Chisq Df Pr(>Chisq)
## model1   11 22355 22414 -11167     22333
## model2   16 22363 22449 -11166     22331 2.2801  5     0.8092
```

As we found above, the interaction term does not improve model fit, which provides additional evidence that there is no sex-arrangement interaction effect.

Next, test if sex improves model fit.

```
sex.arr.lmer(bodysize_wknd)
```

```
## refitting model(s) with ML (instead of REML)
```

```
## Data: devdata
## Models:
## lmer(len.um ~ arr + (1 | arr:isogen) + (1 | vial.setup) + (1 | age), data = devdata): len.um ~ arr +
## lmer(len.um ~ Sex + arr + (1 | arr:isogen) + (1 | vial.setup) + (1 | age), data = devdata): len.um ~
```

```
##        npar   AIC   BIC logLik -2*log(L) Chisq Df Pr(>Chisq)
## model1   10 22764 22818 -11372     22744
## model2   11 22355 22414 -11167     22333 411.2  1  < 2.2e-16 ***
## ---
## Signif. codes:  0 '***' 0.001 '**' 0.01 '*' 0.05 '.' 0.1 ' ' 1
```

As above, sex significantly improves model fit, which leads us to conclude that there is a sex effect on body size.

Lasty, we will test if arrangement improves model fit.

```r
arr.sex.lmer(bodysize_wknd)
```

```
## refitting model(s) with ML (instead of REML)

## Data: devdata
## Models:
## lmer(len.um ~ Sex + (1 | arr:isogen) + (1 | vial.setup) + (1 | age), data = devdata): len.um ~ Sex +
## lmer(len.um ~ Sex + arr + (1 | arr:isogen) + (1 | vial.setup) + (1 | age), data = devdata): len.um ~
##        npar   AIC   BIC logLik -2*log(L)  Chisq Df Pr(>Chisq)
## model1    6 22349 22381 -11168     22337
## model2   11 22355 22414 -11167     22333 3.5667  5     0.6133
```

Once again, we confirm the results from above that arrangement does not improve model fit. We can estimate that effect with weekend emergence excluded, and compare it to the complete data set.

```r
lmer(len.um ~ Sex + (1|arr:isogen) + (1|vial.setup) + (1|age), data=bodysize_wknd)
```

```
## Linear mixed model fit by REML ['lmerMod']
## Formula: len.um ~ Sex + (1 | arr:isogen) + (1 | vial.setup) + (1 | age)
##    Data: bodysize_wknd
## REML criterion at convergence: 22319.92
## Random effects:
##  Groups     Name        Std.Dev.
##  arr:isogen (Intercept) 128.22
##  vial.setup (Intercept)  76.73
##  age        (Intercept) 161.59
##  Residual               275.07
## Number of obs: 1578, groups:  arr:isogen, 26; vial.setup, 16; age, 16
## Fixed Effects:
## (Intercept)      Sexmale
##      2976.1       -306.7
```

```r
lme(fixed = len.um ~ Sex,
    data = bodysize_wknd,
    random = list(~1|isogen, ~1|vial.setup, ~1|age)
)
```

```
## Linear mixed-effects model fit by REML
##   Data: bodysize_wknd
##   Log-restricted-likelihood: -11132.55
##   Fixed: len.um ~ Sex
## (Intercept)     Sexmale
##   3024.0158   -306.6624
##
## Random effects:
##  Formula: ~1 | isogen
##          (Intercept)
```

```
## StdDev:    119.6432
##
##  Formula: ~1 | vial.setup %in% isogen
##         (Intercept)
## StdDev:    40.41346
##
##  Formula: ~1 | age %in% vial.setup %in% isogen
##         (Intercept) Residual
## StdDev:    179.9947 239.1489
##
## Number of Observations: 1578
## Number of Groups:
##                         isogen          vial.setup %in% isogen
##                             26                             171
## age %in% vial.setup %in% isogen
##                            523
```

Excluding the weekend emergence reveals a larger effect of sex on body size:
males are 306 $\mu$m smaller than females when we exclude weekend data.

We therefore conclude that only sex affects body size, and arrangement (i.e., genotype) does not affect body size.
